# Transcriptomics supports local sensory regulation in the antenna of the kissing bug *Rhodnius prolixus*

**DOI:** 10.1101/515353

**Authors:** Jose Manuel Latorre-Estivalis, Marcos Sterkel, Sheila Ons, Marcelo Gustavo Lorenzo

## Abstract

*Rhodnius prolixus* has become a model for revealing the molecular bases of insect sensory biology due to the publication of its genome, its well characterized behavioural repertoire and the advent of NGS technologies. Gene expression modulation underlies behaviour-triggering processes at peripheral and central levels. Still, the regulation of sensory-related gene transcription in sensory organs is poorly understood. Here we study the genetic bases of plasticity in antennal sensory function, using *R. prolixus* as an insect model. Antennal expression of neuromodulatory genes such as those coding for neuropeptides, neurohormones and their receptors was characterized by means of RNA-Seq. New nuclear receptor and *takeout* gene sequences were identified for this species, as well as those of enzymes involved in the biosynthesis and processing of neuropeptides and biogenic amines. We report a broad repertoire of neuromodulatory and endocrine genes expressed in antennae and suggest that they modulate sensory neuron function locally. Diverse neuropeptide-coding genes showed consistent expression in the antennae of all stages studied. Future studies should characterize the contribution of these modulatory components acting over antennal sensory processes to assess the relative contribution of peripheral and central regulatory systems on the plastic expression of insect behaviour.

## 1. INTRODUCTION

*Rhodnius prolixus* has been an important insect model for neuroethological studies for many decades (Barrozo et al., 2016; Wigglesworth and Gillett, 1934). Relevant aspects of its neuroethology, such as host odour-mediated behaviour (Guerenstein and Lazzari, 2009; Manrique and Lorenzo, 2012), circadian modulation (Lazzari, 1992), the action of biogenic amines and neuropeptides (Orchard, 2006) or the expression of behavioural plasticity (Bodin et al., 2009a; Vinauger et al., 2016) have been thoroughly studied. Recently, molecular processes related to sensory function have been characterized for *R. prolixus*, such as the tissue-specific expression profiles of odorant receptor genes (Latorre-Estivalis et al., 2015a) and related changes associated to development and nutrition (Latorre-Estivalis et al., 2015b). Additionally, neuropeptide precursor genes were described for *R. prolixus* (Ons et al., 2009) and the dynamics of neuropeptide expression or release at diverse physiological conditions were characterized for processes such as feeding or ecdysis (Sterkel et al., 2011; Wulff et al., 2017). Based on the current knowledge on its behaviour and physiology, and the recent publication of its genome sequence, it is reasonable to suggest that *R. prolixus* has become an appropriate model for revealing the molecular bases of neuroethological processes in insects. Furthermore, neuroethological research in kissing-bug insects is of medical interest given their role as vectors of *Trypanosoma cruzi*, the causative agent of Chagas’ disease, which is considered a neglected disease affecting over 8 million people worldwide (http://www.who.int/chagas/disease/en/).

Kissing-bug antennae are multimodal sensory organs dedicated to detect diverse stimuli associated to hosts (Guerenstein and Lazzari, 2009), microenvironmental features and intraspecific communication (Barrozo et al., 2016). The physiological bases of sensory processes sit on receptor neurons that express specific membrane proteins that confer them an ability to react to specific stimuli present in the environment. These neurons are mostly found in tiny hair-like structures called sensilla, which can house from one to several dozen sensory cells (Carey and Carlson, 2011). In a recent study, the expression of sensory-receptor coding genes was characterized in *R. prolixus* antennae by means of RNA-Seq (Latorre-Estivalis et al., 2017). Therefore, the antennal expression of a large set of genes related to diverse stimulus transduction processes was reported (chemoreceptors, odorant binding proteins - OBPs, chemosensory proteins - CSPs, transient receptor potential - TRP channels and pick pockets - PPKs receptors) (Latorre-Estivalis et al., 2017).

The triggering of behaviors as a response to relevant external stimuli can be modulated at peripheral and central levels (Gadenne et al., 2016). Insect behavior shows plasticity depending on age, physiological status (i.e., phase of daily cycle, nutritional or reproductive status) and experience (Gadenne et al., 2016). For instance, mature kissing-bugs seek host cues promptly, but they do not express proper host-seeking behaviour during the first week after ecdysis (Bodin et al., 2009b) or after engorgement (Bodin et al., 2009a). Electroantennography and single sensillum recordings performed on different insect species have reported a high degree of physiological plasticity at sensory levels (Kromann et al., 2015; Qiu et al., 2006), at least partially explaining behavioral changes triggered by feeding or development. Similar changes have been documented at the molecular level, with altered gene expression associated to feeding (Bonizzoni et al., 2011) or age (Bohbot et al., 2013). In fact, variations in gene expression depending on nutritional status or development have been described for olfactory correceptors in the antennae of *R. prolixus* (Latorre-Estivalis et al., 2015b). Nevertheless, information about elements regulating sensory gene transcription and the abundance of the corresponding proteins in insect peripheral organs is very limited (Farhan et al., 2013; Jung et al., 2013; Kwon et al., 2016; Lee et al., 2017). Physiological mechanisms modulating peripheral responses to sensory stimuli involve signaling controlled by biogenic amines, hormones, and neuropeptides, as well as their target G-protein coupled receptors (GPCRs) and nuclear receptors, overall controlling the functional status of sensory processes (Gadenne et al., 2016). The main objective of this study is to characterize modulatory components potentially involved in the local regulation of antennal sensory function using *R. prolixus* as a model insect. For this purpose, we characterized the expression of a diverse set of genes known for their neuromodulatory and endocrine roles in the antennae of 5^th^ instar larvae and adults of *R. prolixus* by means of RNA-Seq.

## 2. MATERIAL AND METHODS

### 2.1 Transcriptomic data analysis

Read sequences and *de novo* assemblies were obtained from Latorre-Estivalis et al. (2017). In this study, three antennal transcriptomes of unfed 21 day-old 5^th^ instar larvae and female and male adults from *R. prolixus* (colony originated from Honduras and held at the Centro de Pesquisas René Rachou – FIOCRUZ) were obtained. A total of sixty antennae were collected per sample and used for RNA extraction for subsequent RNA-Seq library preparation and sequencing as described in Latorre-Estivalis et al. (2017). Briefly, sequencing was performed at the W. M. Keck Centre for Comparative and Functional Genomics (University of Illinois at Urbana-Champaign, IL, USA) on an Illumina HiSeq2000 from both ends to a total read length of 100 nucleotides. Read sequences were obtained from PRJNA281760/SRP057515 project at NCBI, which contains data from the three conditions analysed: SRS923612/SRX1011796/SRR2001242 (antennal library from larvae); SRS923595/SRX1011769/SRR2001240 (antennal library from female adults); and SRS923599/SRX1011778/SRR2001241 (antennal library from male adults). Reads were mapped to the *R. prolixus* genome assembly (version RproC3) by means of STAR v.2.6.0 (Dobin et al., 2013) and an edited genome GFF file. Raw read counts were used for differential expression analyses among stages and between sexes using the edgeR (v3.6.8). The FDR adjusted p-value (False Discovery Rate) <0.1 was set as threshold to define the significance level. Heat maps showing gene expression (expressed as Fragments Per Kilobase Million - FPKM value +1 following by Log10 transformation) of the different protein families in the conditions tested were prepared using the gplot package in R.

### 2.2 Manual gene curation

Manual curation of genome project databases by means of the inclusion and correction of gene models, using transcriptomic data and published studies, is fundamental for increasing database quality. The use of reliable genome databases, which need to be as complete and validated as possible, is especially relevant for performing adequate quantitative transcriptomic and functional genetic studies. Most of the target sequences curated herein were obtained from Ons et al. (2011); Ons et al. (2016); Ons (2017); Mesquita et al. (2015); and Yeoh et al. (2017) (details in Supplementary Tables S1 and S2). Therefore, all sequences were compared to the SOAPdenovo and Trinity generated antennal assemblies from Latorre-Estivalis et al. (2017). The discrepancies observed between target gene models from the *R. prolixus* genome (Gene set: RproC3.3, available on 24 Oct 2017) and the transcripts from the *de novo* antennal assemblies are reported in Supplementary Tables S1-S6. In the case of neuropeptide precursor and GPCRs genes that were manually corrected/extended, new Generic Feature Format (GFF) files were created and included in the RproC3.3 version of the *R. prolixus* genome GFF file. In case of the other gene families, new gene models were created only for those genes that were absent from the VectorBase gene prediction database or those whose gene models were partially constructed. The modified GFF file of the genome was used for read mapping. The protein sequences of all genes analysed and the edited GFF file are included in the Supplementary Material (Database S1 and Database S2, respectively).

**Table 1.** Antennal expression (represented as Fragments Per Kilobase Million - FPKM - values) of neuropeptides and their corresponding receptors with FPKM values higher than 1 in at least two of the analysed conditions. Complete names are detailed in Supplementary Table S1 and S2.

| Neuropeptide | Larvae | Female | Male | Receptor | Larvae | Female | Male |
| --- | --- | --- | --- | --- | --- | --- | --- |
| CCAP | 11.3 | 9.8 | 4.5 | CCAP r | 12.1 | 9.8 | 14.1 |
| Dh31 | 3.4 | 3.3 | 4.8 | CT/DH r1 | 41.8 | 81.1 | 116.7 |
|  |  |  |  | CT/DH r3 | 7.5 | 79.3 | 131.6 |
| Dh44 | 7.7 | 10.7 | 12.4 | CRF/DH r1 | 6.5 | 6.1 | 10.9 |
|  |  |  |  | CRF/DH r2 | 4.1 | 14.8 | 29.2 |
| GPA2 | 10.5 | 14 | 17.5 | GPA2/GPB5 r | 87.9 | 54.1 | 32.5 |
| GPB5 | 1.37 | 1.06 | 1.02 |  |  |  |  |
| LK | 8.7 | 7.2 | 6 | Kinin r 1 | 1.3 | 1.5 | 2.2 |
|  |  |  |  | Kinin r 2 | 0.7 | 7.4 | 14.1 |
| ITP | 7.5 | 20.1 | 12 | ITP r | 11.3 | 5.2 | 5.9 |
| LNPF | 2.5 | 5.1 | 2.6 | LNPF r1 | 13.5 | 31.8 | 39.8 |
| Ntl | 2.4 | 2.3 | 2.1 | Ntl r | 1.7 | 2.9 | 5.4 |
| NP | 5.1 | 0.7 | 11.5 | NP r | 6.2 | 1.6 | 8.2 |
| NPLP1 | 7.7 | 7.1 | 7.9 | NPLP r | 1.3 | 0.7 | 1.3 |
| PDF | 9.6 | 5.8 | 8.1 | PDF r | 1.2 | 1.7 | 6.1 |
| RYa | 3.9 | 8.9 | 8 | RYa r | 3.25 | 3.2 | 3.87 |
| sNPF | 0.2 | 2.5 | 1.9 | sNPF r | 5.3 | 1.6 | 1.5 |
| TK | 3.8 | 8.1 | 4.8 | TK 86C-like r | 2.1 | 7.2 | 5.8 |
|  |  |  |  | TK 99D-like r | 0.2 | 1.3 | 1.4 |

### 2.3 Identification of new genes

Orthologous sequences from *D. melanogaster* (Pauls et al., 2014; Velarde et al., 2006) were used in tBLASTn searches in the *R. prolixus* genomic database (www.vectorbase.org) to identify nuclear receptor genes and enzymes related to prepropeptide/preproprotein processing. Sequences of *takeout* (*to*) genes previously annotated for *R. prolixus* (Mesquita et al., 2015) were used as query to search for new sequences in the genome. Subsequently, all sequences were manually corrected/extended according to our *de novo* antennal transcriptomes and annotated based on their phylogenetic relations to other insect sequences. In addition, the structural characteristics of *to* genes, such as the presence of a signal peptide (detected by means of SignalP 4.0 (Petersen et al., 2011)); of two conserved cysteine residues in the amino terminal region implicated in disulfide bond formation and ligand binding (Touhara et al., 1993); and of two conserved motifs (So et al., 2000) were confirmed in *R. prolixus to* sequences

### 2.4 Phylogenetic analysis

For building the phylogenetic trees, protein sequences of *R. prolixus* and other insect species were aligned using G-INS-I strategy in MAFFT v.7 (mafft.cbrc.jp/alignment/server), and manually edited in Jalview v2.6.1. Finally, maximum likelihood trees were built in PhyML v.3.0. Branch support was determined using the approximate Likelihood Ratio Test (aLRT). Non-parametric branch support based on the Shimodaira-Hasegawa-like (SH) procedure

## 3. RESULTS

### 3.1 Manual gene curation

#### 3.1.1 Neuropeptide and neurohormone precursor genes

A total of 17 neuropeptide precursor gene models that were absent from the RproC3.3 version of the *R. prolixus* genome annotation were included in the genome GFF file (Supplementary Table S1). The long neuropeptide F (LNPF) and orcokinin (OK) predictions were corrected according to Sedra and Lange (2016) and Sterkel et al. (2012), respectively. The RYamide gene model was fixed based on our antennal transcriptomes. Besides, IDLSRF-like peptide, glycoprotein hormones alpha-2 (GPA2) and beta-5 (GPB5), and bursicon-beta (also known as partner of bursicon) genes were identified in the *R. prolixus* genome. A new isoform of the *R. prolixus* adipokinetic hormone (AKH) gene, originated through alternative splicing, was identified in the antennal assemblies (Supplementary Table S1). Both AKH isoforms share the signal peptide and the active conserved peptide, but differ in the C-terminal region. Whereas the previously reported isoform encodes the core peptide and a single spacer peptide, the isoform presented here encodes the core peptide and two non-conserved spacer peptides. The gene models of eclosion hormone (EH); ion transport peptide (ITP) isoform A; NVP-like; orcokinin-B; and orcokinin-C remained incomplete because it was impossible to fix them due to problems in the genome assembly, e.g. some fragments were located in the opposite strand or were absent from the genome assembly (Supplementary Table S1).

#### 3.1.2 G-protein coupled receptors

Most of the biogenic amine-related GPCR gene models were edited (Supplementary Table S2). However, many of these genes models are still incomplete. In the case of Family A neuropeptide receptor genes, a total of 15 gene models based on Ons [40] were included in the GFF file of the *R. prolixus* genome (Supplementary Table S2). Besides, 11 gene models of this receptor family were edited in the existing GFF file of the genome. Two isoforms (alfa and beta) of the Corazonin (CZ) receptor gene were described Hamoudi et al. (2016). Nevertheless, our antennal transcriptome only presented the alfa isoform (GenBank Acc. N° AND99324). A second kinin receptor (previously described as an orphan receptor by Ons et al. (2016)) and a Tachykinin 86C-like receptor were identified. Most of the Family B neuropeptide receptor gene models were also fixed (Supplementary Table S2). A phylogenetic tree was built to annotate both the calcitonin-like (CT) and the corticotropin-releasing factor-related like (CRF) diuretic hormone (DH) receptors (Supplementary Fig. 1S). Two CT/DH-like receptors were previously described in *R. prolixus* by Zandawala et al. (2013): receptor 1 and receptor 2, the ortholog of *D. melanogaster hector* gene (FlyBase Acc. Number CG4395). The resulting phylogenetic tree suggested that a third CT/DH like-receptor previously described by Ons et al. (2016) seems to be exclusive of heteropteran insects (Supplementary Fig. S1). The CRF/DH-like receptors 1 and 2 (including isoforms 2A and 2B) were grouped in a different clade as shown in Zandawala et al. (2013).

#### 3.1.3 Biogenic amine biosynthesis enzymes

All enzymes known to mediate biogenic amine biosynthesis in other insects were annotated in the last version of the *R. prolixus* genome (Mesquita et al., 2015); however, minor changes would be needed to fix some of them (Supplementary Table S3). These models include: 1) Tyrosine 3-monooxigenase (ple), which synthesizes dopamine from L-tyrosine; 2) DOPA decarboxylase (Ddc), involved in the synthesis of dopamine from L-DOPA; 3) Tyrosine decarboxylase-2 (Tdc2), which participates on synthesis of tyramine from L-tyrosine and; 4) Tryptophan hydroxylase (Trh), which synthesizes serotonin from L-tryptophan.

#### 3.1.4 Neuropeptide processing enzymes

The neuropeptide processing enzymes were not previously annotated in the *R. prolixus* genome (Mesquita et al., 2015). Using sequences from *Drosophila* as queries, we were able to identify a total of 9 enzyme genes that seem to correspond to *R. prolixus* orthologues (Supplementary Table S4). The processing of neuropeptides involves the following enzymes: 1) signal peptidase (SP), which cleaves the signal peptides from their N-terminals; 2) three members of the furin subfamily (dFUR1, dFUR2a and dFUR2b), which are Subtilisin-like endoproteases that cleave the propeptide at monobasic (Arg) and dibasic (Arg-Arg/Lys-Arg) sites; 3) prohormone convertase 2 (*amontillado* or PC2), which cleaves mono (Arg) and dibasic (Arg-Arg; Lys-Arg; Arg-Lys; Lys-Lys) sites; 4) the carboxypeptidase M (two new isoforms were identified in the antennal assemblies with differences in the 3’ region) and D (known as *silver*, which trims C-terminal Arg and Lys after Furins/PC2 cleavage reaction); 5) the PHM (Peptidylglycine alfa-hydroxylating mono-oxygenase) amidating enzyme, which is responsible for the alpha-amidation of the peptide C-terminal; 6) a prolyl endoprotease belonging to the Peptidase 9 protein family, for which no functional information is available for insects (Supplementary Table S4); 7) the amidating enzymes, the peptidyl alfa-hydroxyglycine alfa-amidating lyase (PAL) 1 and 2.

#### 3.1.5 Nuclear receptors

The ecdysone receptor (*Eip75B*) gene was the only annotated nuclear receptor in the *R. prolixus* genome so far (Mesquita et al., 2015); however, no information about isoforms was included in the annotation. In the antennal assemblies, the sequence of the *RproEip75B* was identified using *DmelEip75B* and posteriorly compared to the VectorBase prediction. This comparison allowed correcting the VectorBase prediction and identifying it as isoform A (by means of the two distinctive exons in the N-terminal-region) and the antennal sequence as isoform B (with the first exon located in the second intron of the A isoform)(Segraves and Hogness, 1990). Besides *RproEi75B*, a total of 20 nuclear receptor genes were identified (Supplementary Table S5) and annotated based on their phylogenetic relations to those of *Cimex lectularius*; *Pediculus humanus*; and *D. melanogaster* nuclear receptor sequences (Supplementary Fig. S2). The orthologues of *D. melanogaster eagle* and hormone receptor like-83 genes were not identified either in the *R. prolixus, C. lectularius* or *P. humanus* genomes (Supplementary Fig. S2).

#### 3.1.6 takeout genes

Three *takeout* (*to*) genes had been previously identified in the *R. prolixus* genome: *to1* (RPRC010098); *to2* (RPRC002313); and *to3* (RPRC01009) (Mesquita et al., 2015). A total of 12 new *to* gene sequences were identified in our assemblies (Supplementary Table S6) and annotated based on their phylogenetic relations (Figure 4). Considering this analysis, RPRC002313 and RPRC010096 were annotated as *to6* and *to2*, respectively. *R. prolixus to* genes were separated into two different clades: *to1*-*to9* and *to10*-*to15*. All the structural characteristics of *to* genes were identified in *R. prolixus to* sequences: presence of signal peptide; two conserved cysteine residues in the N-terminal region and two conserved motifs (So et al., 2000). As expected, the length of all *to* sequences was close to 250 amino acids (Supplementary Fig. S3). Finally, it was observed that 11 out of 15 *to* genes clustered in KQ034137 and KQO34102 supercontigs, with 8 and 3 genes each (Supplementary Fig. S4).

### 3.2 Antennal expression profiles

#### 3.2.1 Neuropeptide and neurohormone precursor genes

A total of 31 neuropeptide precursor genes were found to be expressed in *R. prolixus* antennae, considering a value of >1 Fragments Per Kilobase Million (FPKM) in at least one library as an exclusion threshold (see Supplementary Database 3). Fifteen out of 44 *R. prolixus* neuropeptide genes showed FPKM values higher than 10 in at least one library. Allatostatin-CC (AstCC), allatostatin-CCC (AstCCC), ITG-like, IDLSRF-like peptide and OK were the most highly expressed neuropeptide genes in the antennae of *R. prolixus* (Fig. 1a and Supplementary Database 3). The gene encoding for AstCC was the one showing highest expression in our database, especially in larval antennae (larvae FPKM value = 888; female FPKM value = 98.5 and male FPKM value = 55). Indeed, the lower expression of this gene in male antennae was statistically significant (FDR<0.05) when compared to that observed in larval antennae (Table S7). For AstA and myoinhibitory peptide (MIP), a significant lower expression (FDR<0.05) was also observed in the antennae of both adult stages when compared to larvae (Table S7). The antennal expression of allatotropin (AT); OK and IDLSRF-like peptide seems to increase after imaginal moult (Fig. 1a). The expression reported for OK; Dh31; CAPA; AKH and ITP is the sum of their different isoforms or splicing variants.

**Figure 1.**
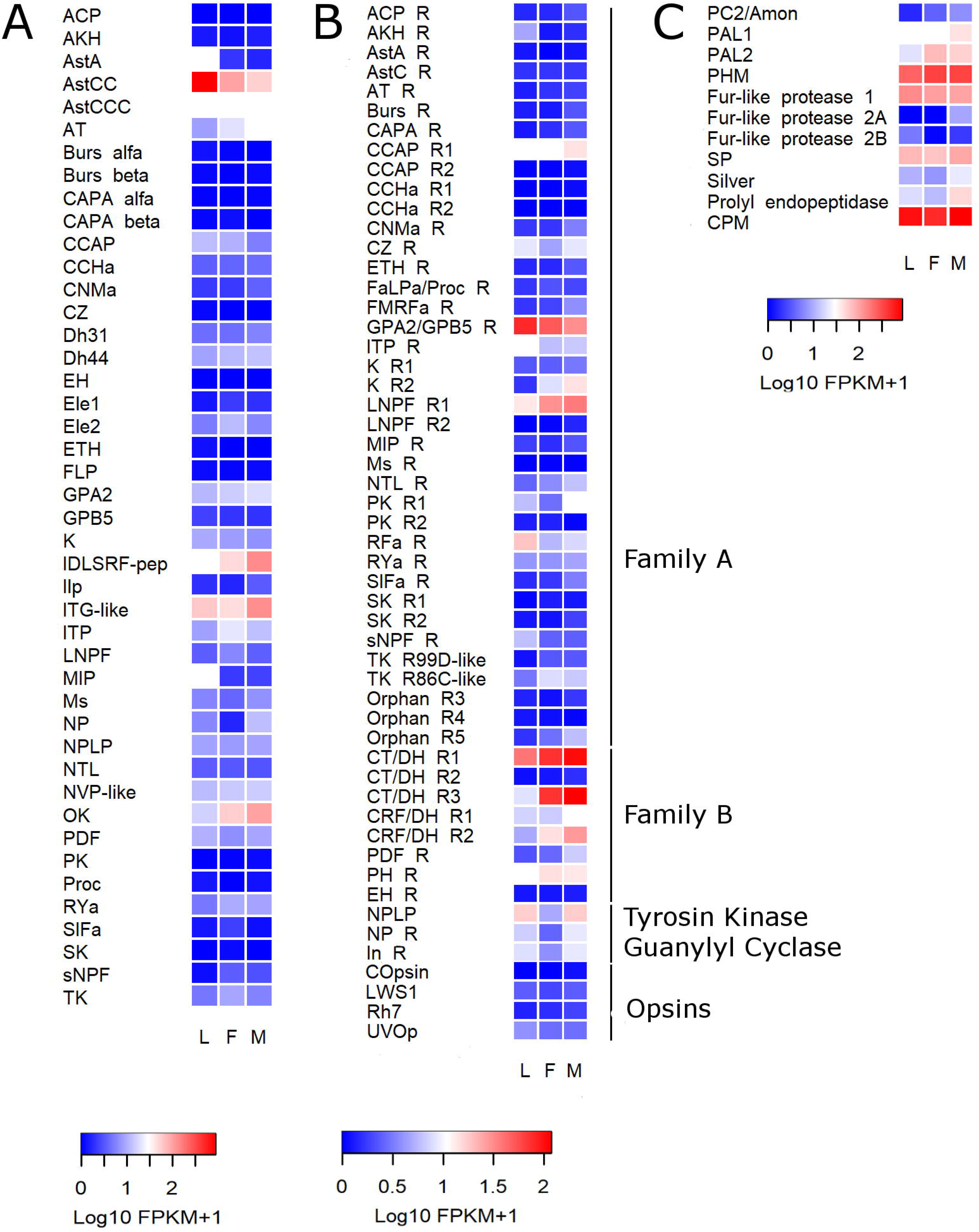
Heat map comparing the expression levels of (a) neuropeptide precursor genes, (b) G protein-coupled receptor genes, and (c) neuropeptide processing enzymes in the antennae of *R. prolixus* larvae (L), female (F) and male (M) adults. Expression levels (displayed as Log10 FPKM +1) represented by means of a colour scale, in which blue/red represent lowest/highest expression. Abbreviations: R, receptor; H, hormone. The complete names of neuropeptide precursor genes, their receptors and enzymes are detailed in Supplementary Table S1-3.

#### 3.2.2 GPCRs

Data suggest that more than half of Family A neuropeptide receptor genes (25 out of 38 genes) were expressed in the antennae (FPKM values >1 in at least one library; Supplementary Database 3). Crustacean cardioactive peptide (CCAP) receptor 1; NPF receptor 1; ITP; GPA2/GPB5 receptor and RFamide peptide receptor were the most highly expressed Family A receptor-coding genes (Fig. 1b and Supplementary Database 3). The expression of the AKH receptor was significantly lower (FDR=0.06) in females, as compared to larval, antennae (Table S7). Interestingly, the expression of kinin receptor 2 increased significantly in the antennae of adults (FDR=0.058 and FDR=0.014 for female and male, respectively; Fig.1b and Table S7). The antennal expression reported for ACP/CZ related peptide, Capability (CAPA) and CZ receptors, as well as for Pyrokinin receptor 2 was the sum of their different isoforms.

In the case of Family B neuropeptide receptor genes, only calcitonin-like diuretic hormone (CT-DH) receptor 2 showed FPKM values lower than 1 (Supplementary Database 3). Five out of seven receptor genes belonging to this family presented FPKM values higher than 10 in at least one library (Supplementary Database 3). CT/DH receptor 3, which according to our phylogenetic analysis seems to be exclusive of heteropterans, showed the highest expression for this family. In fact, its expression showed a significant increase in the antennae of adults (FDR=0.0978 and FDR=0.041 for female and male, respectively) when compared to those from larvae (Fig. 1b and Table S7). A similar expression pattern was observed for CT/DH receptor 1 gene (isoforms B and C included) and for the corticotropin releasing factor like diuretic hormone (CRF/DH) receptor 2 (isoforms A and B included) (Fig. 1b). Regarding opsin expression, transcripts of UV opsin and long wave sensitive opsin 1 (LWS1) were detected in all three libraries (Fig. 1b).

#### 3.2.3 Tyrosine kinase and guanylyl cyclase type receptors

The neuropeptide-like precursor 1 (NPLP1) putative receptor (tyrosine kinase-type) and the potential neuroparsin (guanylyl cyclase receptor) seem to be expressed in the antennae of *R. prolixus* (Fig. 1b).

#### 3.2.4 Neuropeptide processing enzymes

All enzymes involved in neuropeptide processing, except prohormone convertase 1, seem to be expressed in the antennae of *R. prolixus*, presenting values higher than 10 FPKM in at least one library (Supplementary Table S7). The peptidyl-amidating monooxigenase, signal peptidase and furin-like protease 1 genes showed the highest expression (Fig. 1c).

#### 3.2.5 Biogenic amine related genes

Expression of at least 16 out of 20 biogenic amine receptor genes was detected in the antennae of *R. prolixus* (FPKM value >1 in at least one library). Dopamine ecdysone receptor, muscarinic acetylcholine receptor type C; orphan receptor 1; serotonin receptors 1b and 2b presented the highest antennal transcription within this group (Fig. 2a and Supplementary Database 3). The expression of the octopamine (Oct) beta receptor 3 showed a significant increase (FDR=0.071) in male antennae compared to larvae (Fig. 2a), while octopamine beta receptors 1 and 2 showed a similar trend.

**Figure 2.**
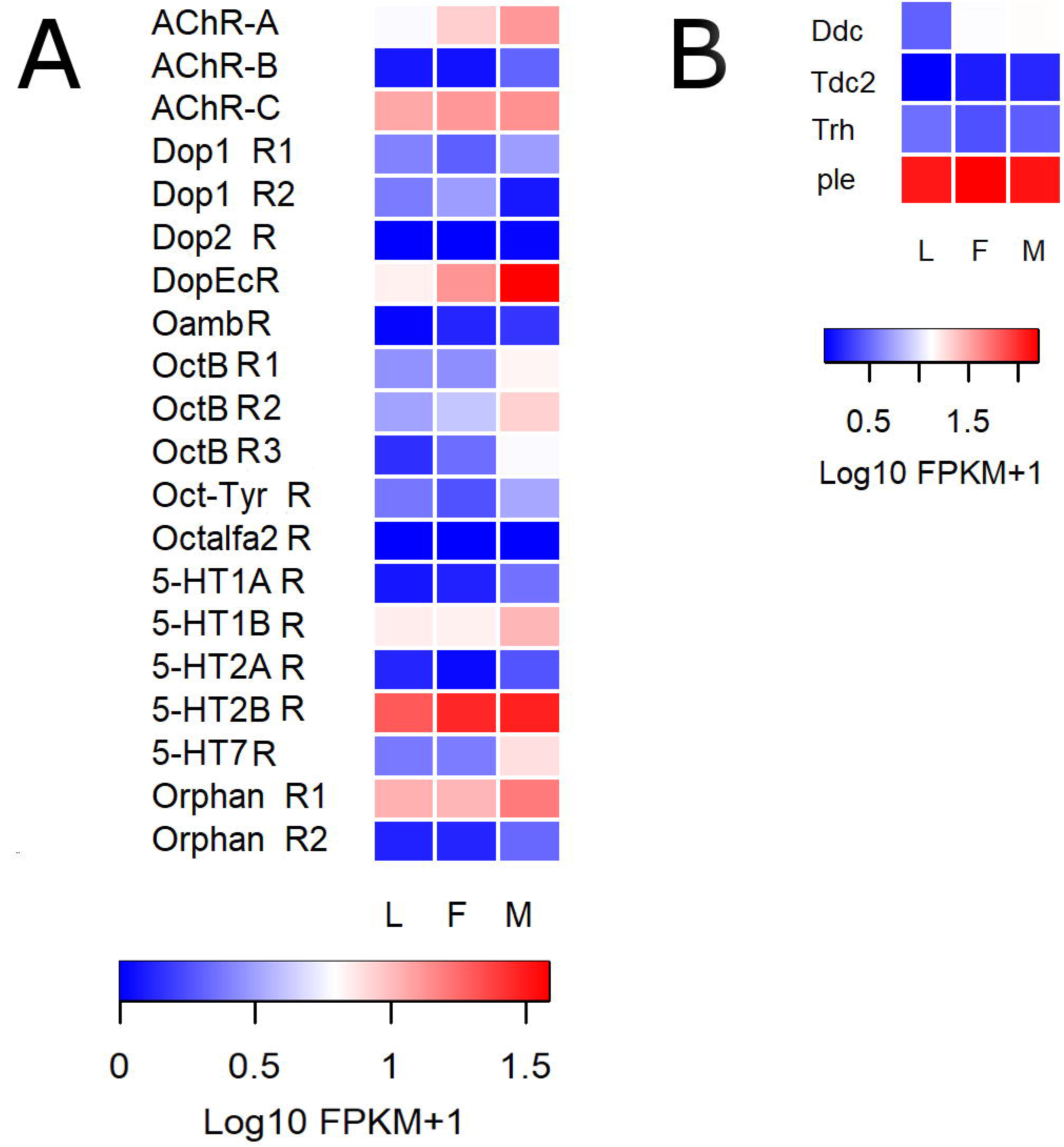
Heat map comparing antennal expression levels of *R. prolixus* genes coding for putative (a) BA-detecting GPCRs and for (b) enzymes involved in BA synthesis in the antennae of larvae (L), female (F) and male (M) adults. Expression levels (displayed as Log10 FPKM +1) represented by means of a colour scale, in which red/red represent lowest/highest expression. Abbreviations: AC, acetylcholine; R, receptor; Dop, Dopamine, M-Ach, Muscarinic Acetylcholine; Oct, Octopamine; Tyr, Tyramine; Ser, Serotonine; AADC, Amino acid decarboxylase. Complete names of biogenic amine receptors and enzymes are detailed in Supplementary Table S4-5.

All genes encoding for enzymes involved in the biosynthetic pathway of biogenic amines were detected in the antennae of *R. prolixus* (Fig. 2b). The gene that encodes for Tyrosine 3-monoxigenase, which synthesizes DOPA from L-tyrosine, was the most highly expressed of this enzyme group (Fig. 2b).

#### 3.2.6 Nuclear receptor genes

Ecdysone-induced protein 75, hepatocyte nuclear factor 4, hormone receptor-like in 96 and *ultraspiracle* were the genes with the highest expression, with FPKM values >10 in the three libraries (Fig. 3; Supplementary Database 3). The expression of hormone receptor-like in 3 increased significantly after imaginal moult in male antennae (FDR= 0.017; Table S7). Six nuclear receptor genes had no expression (FPKM value < 1 in the three libraries) in the *R. prolixus* antennal transcriptomes, these were: *Dissatisfaction*; Ecdysone-induced protein (EIP) 78C; Hormone receptor (HR) like in 51; *Knirps-like2*; *Tailless* and *Seven up* (Fig. 3; Supplementary Database 3).

**Figure 3.**
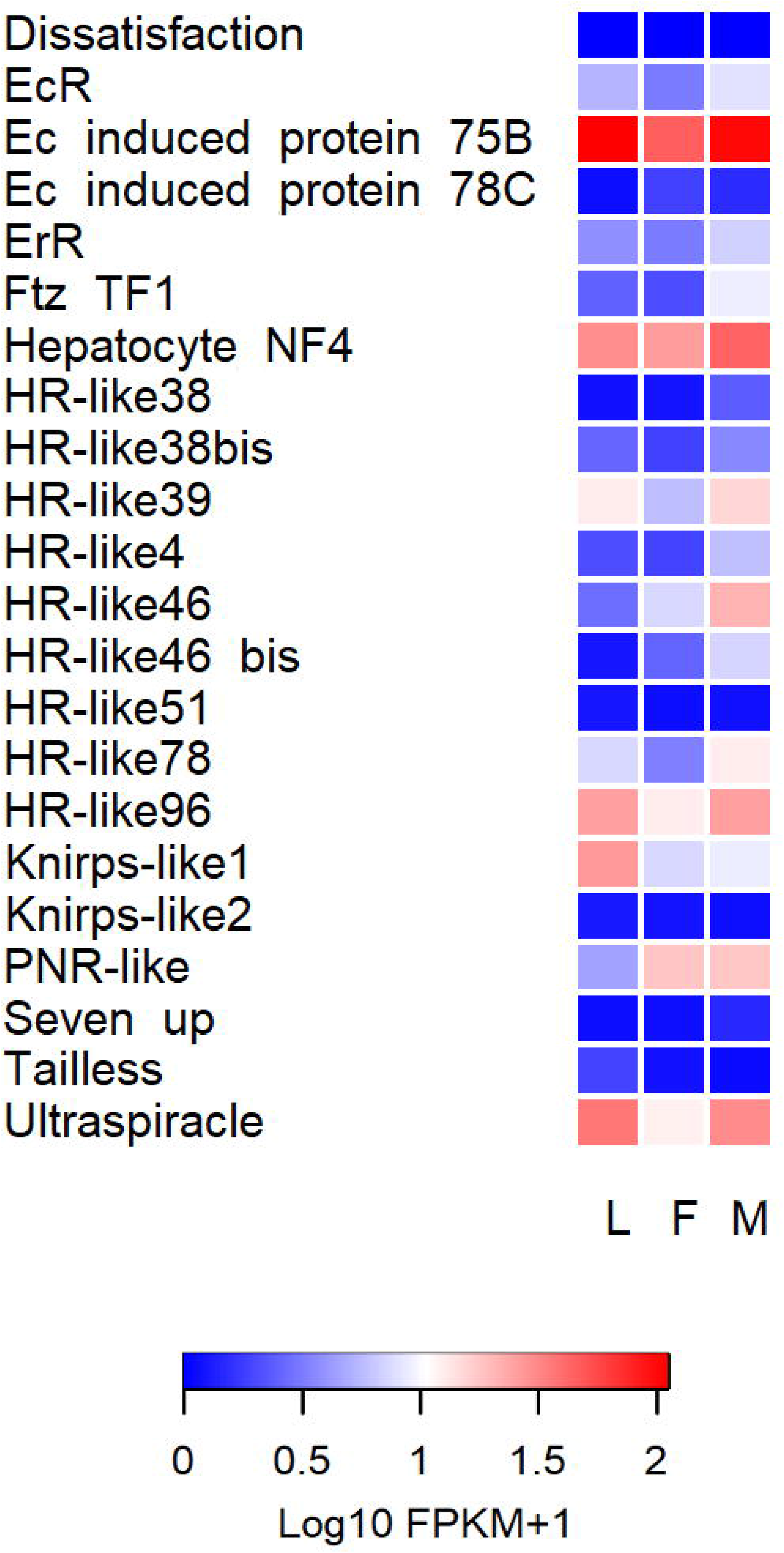
Heat map comparing the expression levels of *R. prolixus* nuclear receptors in the antennae of larvae (L), female (F) and male (M) adults. Expression levels (displayed as Log10 FPKM +1) represented by means of a colour scale, in which blue/red represent lowest/highest expression. Abbreviations: R, receptor; Eip, Ecdysone-induced protein; TF, transcription factor; NF, nuclear factor; HR, hormone receptor; PNR, photoreceptor-specific nuclear receptor. Complete names of these genes are detailed in Supplementary Table S6.

#### 3.2.7 takeout genes

These genes were highly expressed in *R. prolixus* antennae (Fig. 4), 6 out 15 presenting FPKM values higher than 1000 in at least one library (Supplementary Database 3). While most *to* genes tended to present an increased expression in adult antennae, a few seemed to follow the opposite pattern. For example, *to11* gen showed a significant decrease after imaginal molt (FDR <0.05 in both sexes; Table S7), while *to2*, decreased its expression significantly only for male adults (FDR=0.012; Table S7). Nevertheless, the expression of *to3* showed a significant increase in both adult stages after molting (FDR<0.05; Table S7), and those of *to4*, *to7*, *to8*, *to10*, *to12*, *to14* and *to15* followed a similar profile. The genes included in the clade of *to1*-*to9* tended to present higher expression level in antennae.

**Figure 4.**
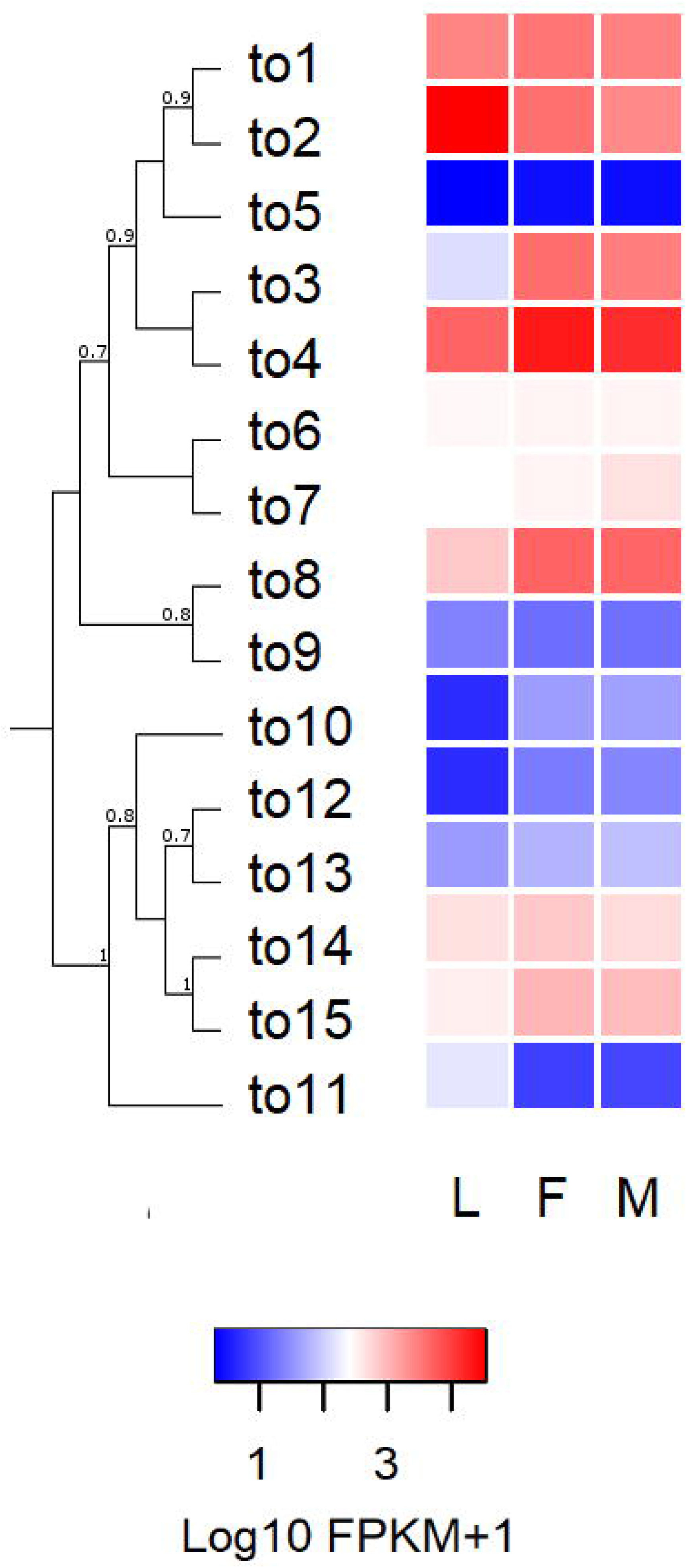
Heat map comparing the expression levels of *R. prolixus takeout* (*to*) genes in the antennae of larvae (L), female (F) and male (M) adults. Expression levels (displayed as Log10 FPKM +1) represented by means of a colour scale, in which blue/red represent lowest/highest expression. The evolutionary history of R. *prolixus takeouts* was inferred by using the Maximum Likelihood method in PhyML v3.0. The support values on the bipartitions correspond to SH-like P values, which were calculated by means of aLRT SH-like test. The LG substitution amino-acid model was used

## 4. DISCUSSION

The molecular bases of sensory plasticity at the local antennal level have been sparsely analysed (revised by Gadenne et al. 2016). Our study has characterized the expression profile of a diverse set of genes encoding different modulatory elements (neuropeptides, GPCRs, nuclear receptors and *takeout* genes) in the antenna of *R. prolixus*. The antennal transcription of a broad repertoire of these genes suggests that diverse local systems may be dedicated to the modulation of antennal functions, such as the detection of host cues and communication signals (Barrozo et al., 2016).

Our results have proven that neuropeptide gene transcripts are produced in the antennae of kissing-bugs (a total of 31 neuropeptide genes seem to be expressed). The production of neuropeptide gene transcripts has already been reported in the antennae of a few insect species (Jung et al., 2013; Matthews et al., 2016; Rinker et al., 2013). The expression of neuropeptide processing enzyme genes was also detected in bug antennae, and as far as we know, this is the first report on the expression of this type of enzyme-coding genes in insect antennae (Figure 1c). The results presented herein add evidence supporting the antennal production of neuropeptides. However, immunohistochemistry and microscopy experiments would be necessary to identify the types of cells producing neuropeptide transcripts in insect antennae. The presence of neurosecretory cells in insect antennae has only been described for mosquitoes (Meola et al., 2000). The authors showed that these cells form synaptoid sites on the dendrites of sensory neurons (Meola and Sittertz-Bhatkar, 2002; Meola et al., 2000).

In recent years, modulatory action by different neuropeptides have been shown for both antennal and labellar chemosensory neurons (Farhan et al., 2013; Jung et al., 2013; Kwon et al., 2016; Lee et al., 2017). Nevertheless, the source of these neuropeptides, whether local or central, was not reported. The current study shows that *R. prolixus* antennae produce a diversity of neuropeptide-coding transcripts, among them high levels of AstCC and ITG-like peptide transcripts in all three libraries (Fig. 1a). Functional RNAi or CRISPR/CAS9 studies should be performed in order to elucidate their role. Orcokinin and IDLSRF-like peptide presented increased antennal expression after the imaginal moult, suggesting that these peptides may modulate adult-specific sensory processes underlying dispersion by flight and mating in kissing-bugs. On the other hand, the decreased antennal expression of AstA and MIP in adults, when compared to 5^th^ instar larvae, suggests an augmented role in immature instars. Instar-specific functional studies with both allatostatins and orcokinin will be necessary in order to understand their antennal function. The significantly lower expression of AstCC in male antennae may suggest a sex-specific antennal role.

The expression of 33 out of 49 neuropeptide and neurohormone receptor genes (FPKM value >1, Supplementary Database 3), the other fundamental component of the neuropeptidergic system, suggests that diverse local regulatory processes can react to a similarly complex set of modulatory signals. Indeed, 14 neuropeptides/neurohormones and their corresponding receptors presented expression higher than 1 FPKM value in at least two conditions (Table 1), reinforcing that parallel local regulatory systems may modulate diverse components of antennal sensory function. The expression of neuropeptide receptor genes in antennae has been already described in other insects (Matthews et al., 2016; Rinker et al., 2013). The high expression shown in all conditions by LNPF receptor 1, GPA2/GPB5 receptor (also known as leucine-rich repeat-containing G protein-coupled receptor 1 - LGR1) and CT/DH receptor 1 (Fig. 1b), suggests important regulatory roles on antennal function. Interestingly, a LNPF-based system modulates responsiveness to food odours of a specific class of OSN in *D. melanogaster* (Lee et al., 2017). Whether this could also be the case for OSNs in *R. prolixus* antennae deserves consideration. The significantly augmented expression of CT/DH receptor 3 and Kinin receptor 2 observed in the antennae of adults (Fig. 1b and Supplementary Table S7) suggests a regulatory function of adult-specific sensory processes. A similar increased adult expression profile was previously observed in the antennae of *R. prolixus* for several chemoreceptors (Latorre-Estivalis et al., 2017). Therefore, it would be interesting to study whether these are functionally connected in the adult phase. The significant decrease observed on the expression of the AKH receptor gene in female antennae may suggest a relation to the modulation of pheromone perception and production as observed for *D. melanogaster* in a sex-specific and starvation dependent manner (Lebreton et al., 2016). Again, it would be interesting to analyse its functional role in kissing-bugs.

Peripheral effects of biogenic amines and their antennal production in insects have been reviewed by Zhukovskaya and Polyanovsky (2017). As observed for neuropeptides (Jung et al., 2013; Kwon et al., 2016; Lee et al., 2017), the modulation of chemosensation and other sensory modalities by biogenic amines (Andres et al., 2016; Inagaki et al., 2012) depends on their levels (Zhukovskaya and Polyanovsky, 2017), as well as the abundance of their receptors (McQuillan et al., 2012). Actually, *in situ* hybridization allowed detecting octopamine and tyramine receptor gene transcripts in the vicinity of sensory receptor neurons of different insects (Jung et al., 2013; Kutsukake et al., 2000). Furthermore, the presence of dopamine ecdysone receptor has been shown for the labellar cells expressing Gr5 in *D. melanogaster* (Inagaki et al., 2012). This supports the existence of direct modulatory effects of biogenic amines on peripheral sensory processes. Biogenic amines such as octopamine have been proposed to directly affect signal transduction and spike generation on OSNs (Grosmaitre et al., 2001). Consistent with these findings, a diverse set of transcripts of biogenic amine receptors was identified in the antennal transcriptome of *R. prolixus* (a total of 16 biogenic amine receptor genes seem to be expressed in them) and in those from other insects (Farhan et al., 2013; Matthews et al., 2016; Rinker et al., 2013). As observed for neuropeptides, most of the genes coding for enzymes involved in the biosynthesis of biogenic amines seem to be expressed in *R. prolixus* antennae (Fig. 2b). Serotonergic nerve fibres innervate the antennae of mosquitoes (Siju et al., 2008) which could relate to the high antennal expression observed for 5-HT receptors in *R. prolixus*, (Fig. 2a). The dopamine ecdysone receptor, which binds dopamine and ecdysone, showed a high expression on adult antennae, especially in those from males (Fig. 2a). Interestingly, this receptor modulates sex pheromone sensitivity in the antennal lobe of male moths (Abrieux et al., 2014). Our results suggest a similar modulation could also occur at peripheral level in *R. prolixus* male antennae. Octopamine receptors may also have a modulatory role on male sensory processes, as they showed increased expression, this being significant in the case of beta receptor 3, in the antennae of male adults (Fig. 2a; Supplementary Table S7). A role of octopamine receptors in the modulation of male sensory physiology was observed in male moths in which this molecule enhances OSN sensitivity to specific sexual pheromone components (Grosmaitre et al., 2001).

Hormonal regulation on insect sensory systems has been poorly studied at the peripheral organs (Bigot et al., 2012). Here we show that most described nuclear receptors are expressed in the antennae of an insect (Fig. 3 and Supplementary Database S3), suggesting that these organs have broad capacity to respond to endocrine signals. It is worth mentioning that *Eip75B* and hepatocyte nuclear factor 4 (*Hnf4*) genes are the most expressed nuclear receptor in *R. prolixus* antennae (Fig. 3). Considering ecdysteroid signalling, the detection of *Eip75B* transcripts indicates a potential capacity of kissing-bug antennae to respond to the EcR-*USP* complex (Ecdysone receptor + *Ultraspiracle*), as observed for *Spodoptera litoralis* (Bigot et al., 2012). Besides, *Eip75B* and hormone receptor-like in 51 transcripts (also known as *unfulfilled*) have been identified in central clock cells of *D. melanogaster* and control the expression of clock genes, playing an important role in the maintenance of locomotor rhythms (Jaumouillé et al., 2015; Kumar et al., 2014). Therefore, we suggest that these nuclear receptors may have a similar regulatory role at the periphery, considering that the presence of a peripheral circadian clock has been reported for insect antennae (Tanoue et al., 2004). The *Hnf4* gene, which induces the expression of enzymes that drive lipid mobilization and β-oxidation as a response to starvation in *D. melanogaster* (Palanker et al., 2009), also showed high expression in antennae. The relatively low nutritional status of the insects used in our studies could relate to its high expression in *R. prolixus* antennae. Functional studies would need to be performed in order to evaluate the potential role of this gene as a nutritional sensor in insect antennae. An increased expression of the hormone receptor-like in 3, which is the heterodimer partner of Eip75B, in male specimens suggest a sex-specific role in antennae.

Fifteen *takeout* genes were identified in the *R. prolixus* genome, while Ribeiro et al. (2014) identified 18 potential takeout transcripts in a midgut transcriptome of this species and Marchant et al. (2016) identified 25 takeout transcripts in the transcriptome of the kissing-bug *Triatoma brasiliensis*. Consistently, these numbers match the scale of those found in *Anopheles gambiae* (10); *Acyrthosiphon pisum* (17); and *Bombyx mori* (14) genomes (Vanaphan et al., 2012). *R. prolixus to* genes present a cluster organization (Supplementary Fig.4S), probably due to gene duplication events, as it was previously observed in other insects (Vanaphan et al., 2012). The antennal expression of *takeout* genes has already been reported in Dipterans (Bohbot and Vogt, 2005; Sarov-Blat et al., 2000). Furthermore, it has been shown that starvation induces the expression of these genes (Sarov-Blat et al., 2000) that have also been related to foraging activity (Meunier et al., 2007). This putative function could explain the high expression observed in the three antennal libraries (Fig. 4), however, functional studies need to be performed to be able to confirm these roles in the antennae of kissing-bugs. Two *to* genes presented significant differences between larval and adult antennal transcriptomes (*to11* and *to3*, with an up and downregulation, respectively) and *to2* is significantly down-regulated when male antennae are compared to those of larvae (Supplementary Table S7). Results suggest that these *to* genes may be related to sex, as observed in *D. melanogaster* (Dauwalder et al., 2002) but experiments are necessary to test this hypothesis.

Antennal cells are bathed by haemolymph but not so the dendrites of sensory neurons (bathed by sensillar lymph). Therefore, it is certain that central signals, i.e., circulating hormones, biogenic amines and neuropeptides can modulate the function of most cells in insect antennae (Gadenne et al., 2016). However, the antennal detection of neuropeptide transcripts (and those of enzymes involved in their biosynthesis and that of biogenic amines) suggests the existence of local regulatory systems that could represent additional sources of modulation of the sensitivity of peripheral neurons. Future RNA-seq, peptidomics, in *situ* hybridisation and other functional genetic experiments should test whether these regulatory components are also present in the antennae of other insects and unveil the interaction between central and peripheral regulatory systems to understand their relative contribution to the control of antennal sensory physiology.

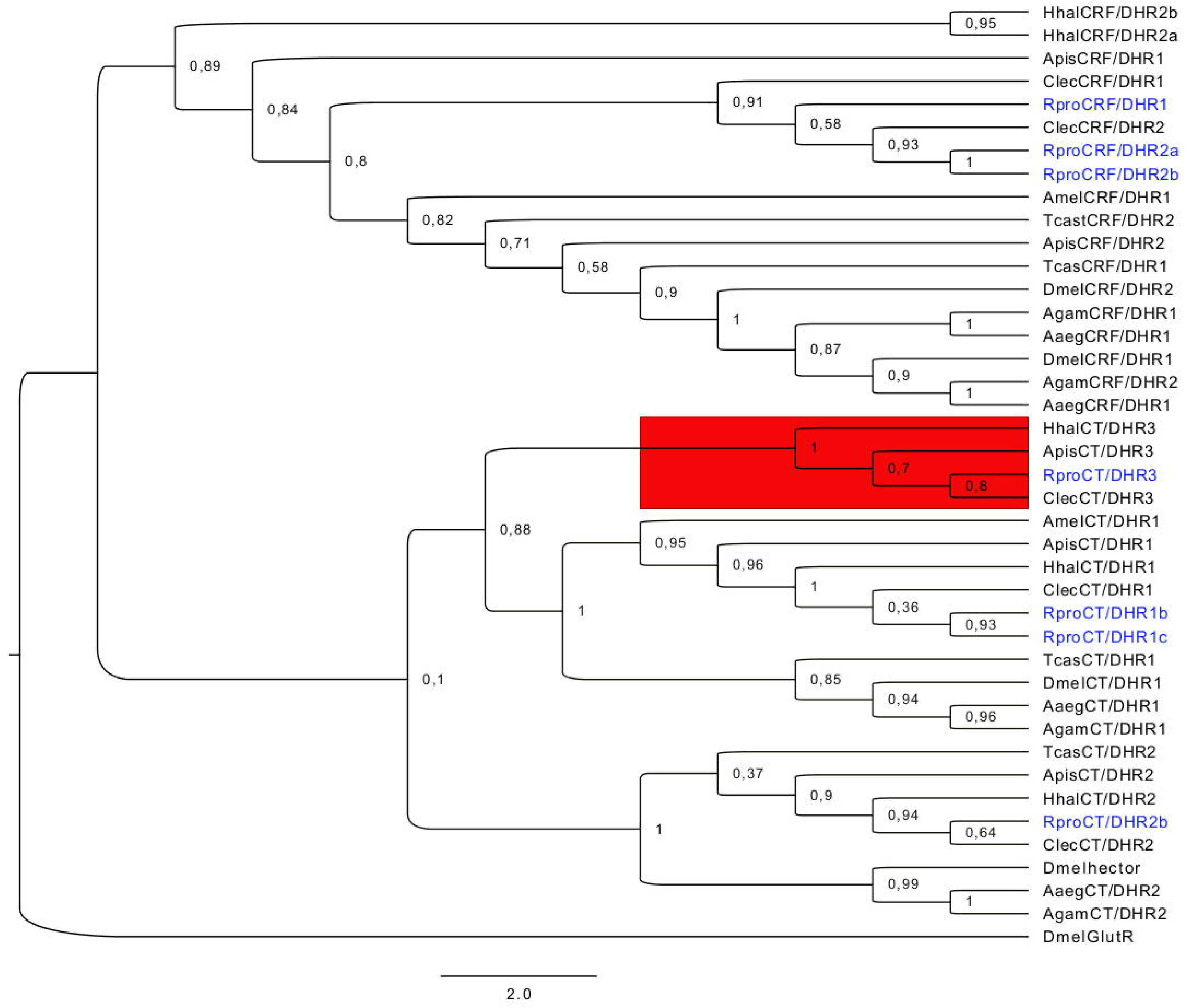

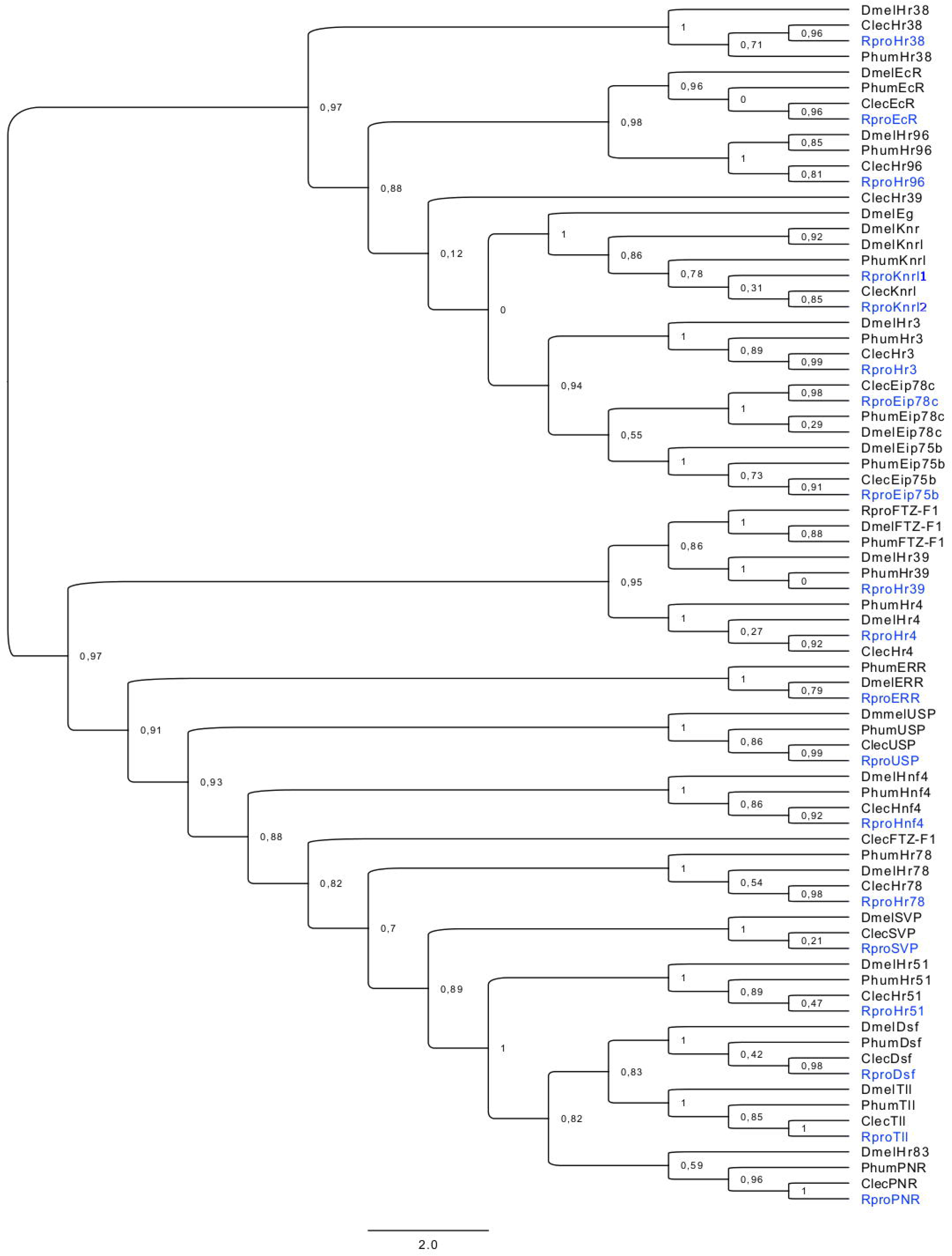

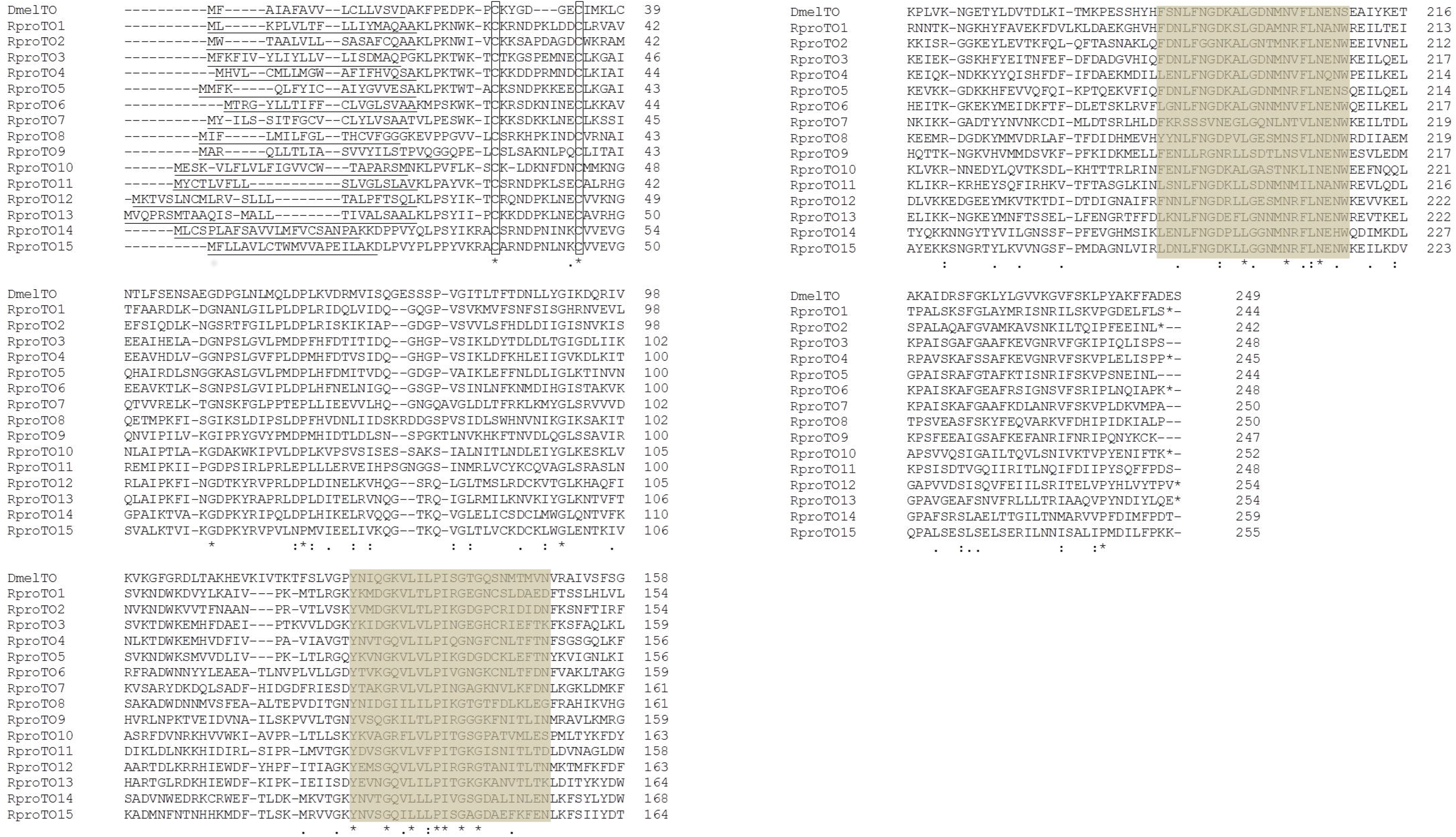

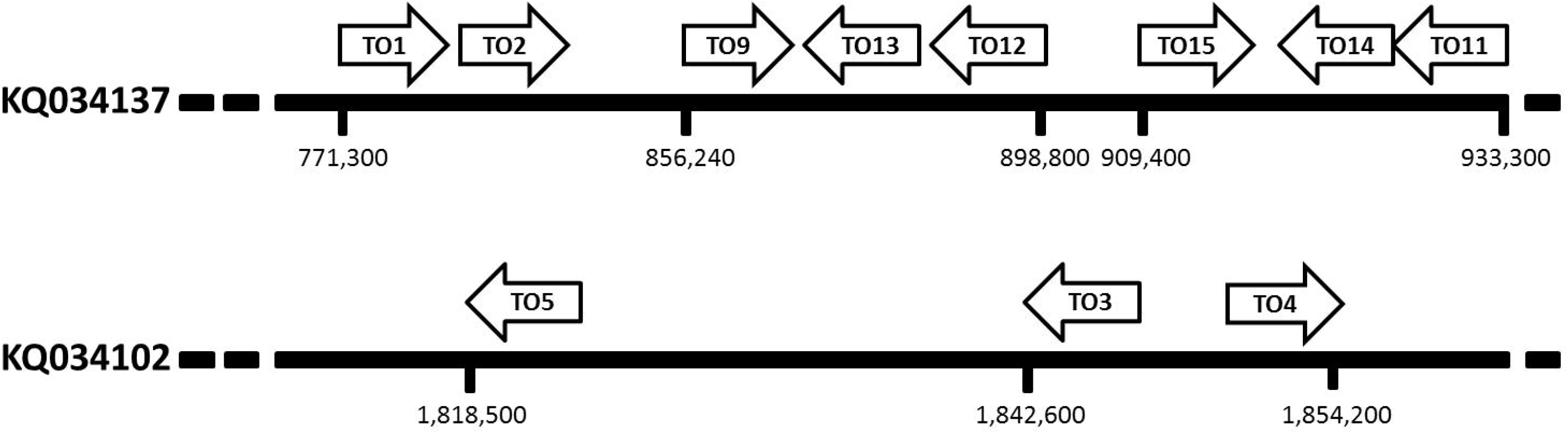

## Supporting information

Supplementary database 1

Supplementary database 3

Supplementary database 4

## ACKNOWLEDGEMENTS

The authors wish to thank the Program for Technological Development in Tools for Health-PDTIS-FIOCRUZ for having facilitated the use of its facilities. We wish to thank Ivana Helena Rocha Oliveira for her help in processing samples used in our experiments, Prof. Claudio Lazzari for kindly providing comments on an earlier version of the manuscript, and Dr. Gabriel da Rocha Fernandes and Fausto Gonçalves dos Santos for their help and support in the bioinformatic analysis.

## SPONSORSHIPS

Authors are indebted to INCTEM (Project number: 573959/2008-0), FAPEMIG (Project number: APQ-01359-11), PROEP-FIOCRUZ (Project number: 401973/2012-3), CNPq (Project number: 483805/2013-0), Le Studium for a granting research fellowship to M.G.L (Short Term Contract of Employment N° 2017-2001-179 - Y17F16), FIOCRUZ Visiting researcher fellowship program (fellowship to J.M.L.E 550017/2015-17), Consejo Nacional de Investigaciones Científicas y Técnicas (CONICET) postdoctoral fellowship (fellowship to J.M.L.E 2015-2017) and Agencia Nacional de Promoción Científica y Tecnológica (Project number: PICT 2016-3103). J.M.L.E., M.S and S.O are researchers from CONICET.

