## Supplementary database 1 for "Transcriptomics supports local sensory regulation in the antenna of the kissing bug *Rhodnius prolixus*"

Abbreviations:

NTE, Initial methionine is missed in the transcript sequence;
CTE, Stop codon is missed in the transcript sequence;
INT, the transcript sequence presents internal problems.

Some GenBanK accesion numbers are included in the header of the transcript sequences

============================
NEUROPEPTIDE PRECURSOR GENES
============================
>Adipokinetic_hormone/Corazonin_related_Peptide-KM975505
MGEEDSSQDHHFSIYTGQMDYSSIIFEMDRSKTVRRLSTVALVYLIFINIFLVEAQVTFSRDWNAGKRSNNIPDCAIAIKSAA
AICQMLLNELRAIATCEMHSLTSQRLNEDVENSQDVFGSQHNG
>Adipokinetic_Hormone_Isoform_A-KM283242
MATNLFITSVLVLLTFHYTLAQLTFSTDWGKRSVRHNAPDCTPNPDTVIFLYKYLQNEFYKMIECGKTEGL
>Adipokinetic_Hormone_Isoform_B
MATNLFITSVLVLLTFHYTLAQLTFSTDWGKRSVRHNAPDCTPNPDTVIFLYKYLQVRRLETRLLGVFIC
>Allatostatin_A-GQ856315
MMLPFIVLLVVDVFALGAQAINDREDDFNKKLTELGLGKRAAYSYVSEYKRLPVYNFGLGKRAHNEGRLYSFGLGKRDYDS
GEEMEYLDDELAIRDELAKRAAKMYSFGLGKRLPSIKYPEGKMYSFGLGKRVPFADQAYFLDDNDSSEESKRSNPNGHRFSF
GLGKRDEQEMNEKRKGERSMQYSFGLGKRTQLDPANNLHN
>Allatostatin-B
MQYVFHIRMSWCYKILLATTLTAIVQGQNPGTVTPMDNVIAEDEYLIPSNPALIDDKRSWKDLQSSGWGKRGWKDMQT
VGWGKRAWTDLPSSGWGKKRAWSDLQSSGWGKRGWKDMQSSGWGKRAWSDLQSSGWGKRAWSDLQSSGWGKR
DWKDMQSSGWGKRAWSDLQSSGWGKRSKGDDQDEDIEENLAEEDKRAWNSLHGGWGKRTADWGSFRGSWGKREP
AWQNLKGLWGKRSPVNLLQDNFQTGYVEPY
>Allatostatin-CC
MWLPSAYISRLSWWCFALVFTTLLAATCVLTSPAPSYNDYQQVGVSYDEYPVVVPKRAALLLDRIMVALQKAVDEENSVKS
TKNRIPIEETMDLQRRGQQKGGRIYWRCYFNAVTCF
>Allatostatin-CCC
MKVFAKMSSSQICAGVLFCLSMLIMLGMSQPTPDKEKLLNELSQELVEDDGSIDRAVIDYLYAKQLFNRLRAQAGAAEIQQ
GKRSYWKQCAFNAVSCFGK
>Allatotropin-GQ162783
MRWSSLLVLVALASIINCIKAGSPSSALYSSAARASGRTRTIRGFKNVQLSTARGFGKRTYPDSQLQPDLIPADWMAEELSSN
PELARFIIRRFIDVDQDGLVSPVELLRNTVCQEPN
>Bursicon_alfa
MASALSSSEHKAQQIDECQVTPVIHVLQYPGCVPKPIPSFACTGRCSSYIQPFKFGIYSCILKLLLSVRLEDLADGEIVHVLSGE
WRARGQCCLVLSESQAWRKKVQKGMKVNLCIKRNVCACLQVTTKAPLECMCRPCTSVEESAIIPQEIAAYADEGPLNNHF
LKPQ
>Bursicon_beta
MPSEIHLVK
EEFDELGRLHRTCSGDIAVNKCEGACSSQVQPSVITPTGFLKECYCCRESYLRERLITLTHCYDPDGMRLTQNGHSTMEIKLK
EPSDCKCFKCGDYTR
>DH31_Isoform_A-GQ856316/AEA51300
MVTNIAVVGVSLMLGTLIVLSAASENIPYIGHRASYFGDMDNEPDSEVMLEILAKLGRTIMRANDLEKPMIYSREASNPWT
AVNKLRPSNLPYNIELAENPDSIYSKRGLDLGLSRGFSGSRAAKHLMGLAAANYAGGPGRRRQA
>DH31_Isoform_B-GQ856317/AEA51302
MVTNIAVVGVSLMLGTLIVLSAASENIPYIGHRASYFGDMDNEPDSEVMLEILAKLGRTIMRANDLENSKRGLDLGLSRGFS
GSQAAKHLMGLAAANYAGGPGRRRQA
>DH31_Isoform_C-AEA51301
MVTNIAVVGVSLMLGTLIVLSAASENIPYIGHRASYFGDMDNEPDSEVMLEILAKLGQTIMRANDLEKPMIYSREASNPWT
AVNKLSSKRGLDLGLSRGFSGSQAAKHLMGLAAANYAGGPGRRRQA
>CAPAalfa-ABS17680

MSYTVGTILVTAVLVSTCAVINSAEQTNEDKTNTTLRIKRSPISSVGLFPFLRAGRARNFPATWGMLVGDDKNKREGGFISFP
RVGRSGPKRNGGGGNGGGLWFGPRLGRNQKRGDSWTLEQLQPNLLPGYPAYNEEEKENQFSEELSDESVSNKIV
>CAPAbeta-ACH70295
MSYTVGTILVTAILVSICSVIISADQTNEDKTDASLRFKRSPITSIGLLPFLRAARARNFPITWGMLVGDDKYKREGGFISFPRV
GRSGPKRIGGGGNGGGLWFGPRLGRNQKRGDSWSLEELQPNLLPGYPAYNDEEKENQFSEESSNESMSNKVV
>CCHAmide
MICSRKMIVTLLLVSLLLTVHGAAFKGARDGDASFRKKPLRRVFYFLLLSGGCSAFGHSCFGGHGKRSDDYMAQIQSRQLQ
RLPPADIVRQWMKSLHNVPPLLE
>CNMamide
MNGKVWLQFLYEGVSRQVMACAVLLGLLTQVPASSPSSTRTQELQESSPSLQQLISLFDYLQNRELEALASDGTQNDSGG
NNGLLESNSNKNRLRELANAYDELLRQNMLQQSAAQHLSAEKRASYMSLCHFKICNMGRKRTPYEAVHKK
>Corazonin
MNFRSSCLLIFIIYSIVHVFGQTFQYSRGWTNGKRAGIPSKEVTACQLQRIKSLLEGKTIPQLYWPCEWSPFMEAALSRQMK
TSELTSLPVVAPLTPEIEENCC
>DH44-ADM26617/HM153808
MWIKCVWVACLVGAVHGSLQDTRAHNLLQDKWEPPLPPAHNTRYYFIDQPLRPQPPLQESEIDDEVSWRGKRMQRPQ
GPSLSVANPIEVLRSRLLLEIARRRMKEQDASRVSKNRQYLQQIGKRHTNRLTNEDKTVNMDYYNSGDFDWDSA
>Crustacean_cardioactive_peptide-GQ888668
MQLLVPCFLLFTALVFAVLTDDVFLQKRVYFPGEIAEPIDPKMKKPFCNAFTGCGKKRSDESMATLVDLNSEPAVEELSRQIL
SEAKLWEAIQEARMELLNRKQQQSDRIPLQPLPLTTIRKRSHYLYT
>Ecdysis_triggering_hormone
MIAILKYILCIECFLTIVATTAKASDEISSSPILNPAFVRDPNEDRQIVFIPLGNDQDKSATNIVRRNDFFYAKNLKTVPRIGRRN
GFTATLAADGRSGARVDRRDEFHEMDKDRLWPWPKAGKYIPIVVKKKSGYFHGDSSADHS
>Eclosyon_hormone-NTE
NIINSKLLCRYTMKKLLLVILLTSFLAEISGRQIGVCIRNCAQCKKMFGVYFEGQMCADTCLKYKGKLIPDCEDIASIGPFLNKLI
>Elevin_1
MGRKYLLTIFVWLCLITLPLALAQGENENGTVVDCRKYPYAPVCRGAGAKRLYLYSPQEINYEDMA
>Elevin_2
MCRKDLLNIFVWLCLITLLLALTSKSQVIGKNGQIVDCKKYVFAPVCRGAGAKRSYPPRFSPQEINYEDLAEEILDSYPIGKRKL
YL
>FaLP
MDRYMMFGGMLVGAWLLIHGQCCVNAGTDSRIRSPLVDPLIRRSPLEKNFMRFGRSSPALQQFPTAYNSNYLDLENSKRS
GRFDRARDNFMRFGRDNEKIALSNRAKDNFIRFGRSKDNFMRFGRIKDNFIRFGRGNDNFMRFGRSGDEIEEILPKDRRDK
ALNRLGRQRLSDKSDNFIRFGRASSDDNEMGDMSKSKRSPIYMDEQKIQEECHEGVCQQTFTKEDTHQHQQDFCPPELD
PNIIVAPEFSLLPSISMKPRIKPKISSNFLRLG
>GPA2
MSMVKILCMFLILTCLDNVFCKDAWGKPGCHKVGHTRKVSIPDCVEFHVNTNACRGYCESWSVPSPLETVLHNPRQAVTS
IGQCCNIMDTEDVEVSVLCLDGTKDLVFKSAKSCSCYHCKKD
>GP5B-NTE
AVLLMVIVGCVTTKVKGTLDPSSTLDCHRRIYSYKVTQADSEGRICWDTINVMSCWGRCDSNEISDWRFPYKRSYHPVCIH
GVRELRRVVLKHCEEGAEPGTEQYDYQHATTCRCHVCRSSEASCQGLRDVVNDQSDLYTESGGNAWNATLVPNGI
>Kinin-BK007870
MILLWMVWTIAVICKTSQGNDIISTSAEGHNLTTAPSPLPTAKDNSKGIRDKRGTGHLLEQLLKENELAAEDLEDEEDLVND
KIKRTNNRGNFAGNPRMRFSSWAGKRAKFSSWGGKRVDDELISGTDGPIEYEIPEDKRANKFSSWAGKRTDEEGVNWM
GNSPADLDSFIQQLEQKRAKFSSWAGKRDEDRQKFSHWAGKKFDDSLNMNDVLLEEEKRGAKFSSWAGKRAKFNSWGG
KRFSNEFMNDNNDIEKNIVEEKRLSINPWKKIDDNGKRAKFSSWGGKRADDDWLKKARFNSWGGKRNSFNANITNSVD
DLFLDHEDALIKRSAAAYTPLSWKRKPIFSSWGGKRTARSTQPQRRLIFPSNLFRDHSTWGALLRPIRRGPDFYAWGGKRST
>ITG-like
MLSVILLLTIGVQAAFGWGGLFNRFSPEMLSNLGYGGHGSSYRVQPFLQNNGAMETLQELQEAEMEGPCYGKRCTANEH
CCPGSVCVDVDGMGSCLFPYGLGQGELCRRHSDCDTGLICSDTGDGAKTCQPPFTAPKQYSEECTMSSECDIHRGLCCQF
QRRHRQAARKVICSYFKDPMVCIGPVASDQVKDDIERTAGEKRITGKTAAFNHLRR
>Insuline_like_peptide
MQALFWKVLLLAAVICCVTCQQWYLYSDEPRQIEAKRGAQKYCGRILDDTLKFICRGKYNERFPSGKKRATESNKGLEDLES
YYDNWYKSLVMPSYYTARAIKRGVHTECCVRPCTFGDLEKYCAE

>Ion_transport_peptide_Isoform_A-GQ253921
MHQERRALAGLVVASTLLSWAVAGPSSRLVLSHPLNKRSFFDLQCKGVYDKSIFARLDRICEDCYNLFREPQLHSLCRSDCFA
SKYFAGCLEALLLREEENKFFQMVEFLG
>Ion_transport_peptide_Isoform_B-GU207866
MHQERRALAGLVVASTLLSWAVAGPSSRLVLSHPLNKRSFFDLQCKGVYDKSIFARLDRICEDCYNLFREPQLHSLCRKNCFT
TDYFKGCLEVLLLQDEMENIQTWIKQLHGAEPEV
>Long_Neuropeptide_F
MNCWLLWLWTGLMACNAAMAMAQPIPADAMARPARPKSFASPDDLRTYLDQLGQYYAVAGRPRFGKRAGGINPRLH
LAVDGVNYRYPLADASDLYDLLFQQQSTE
>Myosuppressin
MILAWMCVTLLAGVLAAPGPDCSPSALQVQSSRVRNMCALYQISSALQAYLDEQNNYQTALRDTNIPYNIPEKRQDIDHV
FMRFGRRR
>Natalisin
MNPVHTLLVALISVGLQETSSVGEKCDKAVCGEERRSDVRAVLGSSEAEPGFWPTRGRRGDSSSTEEVQPPFWAHRGREE
RPCDSSSVNSLNHLYAQEPKFLLIHRRDTMEQDPFWVSRGRKRSQKSNFAYLEETSLTAGEGGGELITVTNSFAGEMRGLW
SLGRLPVQVKSHLNIQYKWHLSLANITNTQPEQVP
>Neuroparsin-GU207864
MSSQSSKTATTALAVLTIFCMVALVSGVFYGCVPCIGDECNLNPGNCPYGIVRDPCGRLVCAAGPGERCGGRDFHLGKCGE
GLSCKCGKCRGCSIKQIMNGRIDCDTTNPMCQ
>Neuropeptide_like_precursor_1-GU207865
MIATSLAPLILTLLLLSKASGENNDNTKSSSKHALVTDTGEEHNLEKRHVSSLLGNRASGPYQTGKRSSPSKSSLDELAERLEEA
AQEDKRYLGALARSGDLRVVARDRQEKREDLDSLIDELASTEEMRRMQFDALRDDLFEEEPDKRGVVSLARAGYLKPTTHD
FLEDDEESSFYPAEDEDKRGGIASLARNGYYQKRTVDAELEQLMSEVYGIGEKRSVASLARSYNLPNAVKGGYENDDEKRNI
PSLLRDRTSPLGEGKRHIGSFVANHGIPFVNNKEGGKRSVGSLARNRDFPYAVKFGKRDAPDEEGEEMSKRYVATLLRQGR
LPIGIDSPDHGEMSSMKEDNDDHDVDKDEMSQVMQDEASKLSIRKKKSVPTVEGPTRIKREAAADEFGGVRDDDSLAQF
ADDAADSPINKRYFGVRGGGKMPGGRLPKVGGRSRNRHENSGRRRH
>NVP-like
MLLLKILLNITTLSAVVLAIPTSILEDLKNVQLQSPIRSNKVKRAQEFIMFGNQQNRAPSFSNIRNDKRASEIDDNSLPDDEGPL
PQVVPQADEATYENNHLTSGNVYDKAYPYTSRDLYYNMLLRNLELTQNLHNDPVFNYDFPSYSIMDGRFKRDTKSTKESHK
PLYRSKREHVLNPEEFLALMKMADSNKDINNKYYDRSSIGWPVYESEIEDFPGVDDDTYETESNDENGAWYNNGMMYQS
RFGHPKDGFRTNRPHKRFMVSKRRMGAENAAQYNMPYGVYDPSDGIPLHRRFLL
>Orcokinin_Isoform_A-FJ167860
MINMLSLTILAMAVAVTSAFPRGELGVEEGNLYPGLYRDQTMEDKEGRNLDTLGSGNLLRDLEAVLRAHPNLFYGRPARN
HDTLDSLSGITFGSQKRFDPLSSAYAADKRNFDEIDRSGFNSFIKKKKNFDEIDRSGFDGFVKRNFDEIDRVGFGSFIKKSEPH
H
>Orcokinin_Isoform_B-JF761320
MINMLSLTILAMAVAVTSAFPRGELGVEEGNLYPGLYRDQTMEDKEGRNLDTLDGVIVEPIYGLRPDKKEGGTLDSLGGGH
LIRNLKENEDNYEHEIKKEALKTKFHTDTSKAFLPNKLSNTNHFISKSLESVNGRSEIMDQHLLRNLVYGKGRRFKRDFLDGLG
GGHLLRDNIDPFVDVHPVKGTSKIPYTGPISSDGDQQVSVNFSNGRHFVREFLDPLGGGHLIRGIDSIGGGHLLRGLDAKEN
GNLGKELTDGFRSGYSISGGPLVREFLDPLGGGHLIRGLDSKSGDHFKREFLDPLGGGHLIRGLDSIGGGHLVREFLDPLGGG
HLIRGLDSKGGDHFKREFLDPLGGGHLIRGLDSIGGGHLVREFLDPLGGGHLIRGLDSEGDSHP
>Orcokinin_Isoform_C-AGW15565
MINMLSLTILAMAVAVTSAFPRGELGVEEGNLYPGLYRDQTMEDKEGRNLDTLDSLQNNRLNAARVKRNSPEIQKSNWSK
RDGVIVEPIYGLRPDKKEGGTLDSLGGGHLIRNLKENEDNYEHEIKKEALKTKFHTDTSKAFLPNKLSNTNHFISKSLESVNGR
SEIMDQHLLRNLVYGKGRRFKRDFLDGLGGGHLLRDNIDPFVDVHPVKGTSKIPYTGPISSDGDQQVSVNFSNGRHFVREF
LDPLGGGHLIRGIDSIGGGHLLRGLDAKENGNLGKELTDGFRSGYSISGGPLVREFLDPLGGGHLIRGLDSKSGDHFKREFLD
PLGGGHLIRGLDSIGGGHLVREFLDPLGGGHLIRGLDSKGGDHFKREFLDPLGGGHLIRGLDSIGGGHLVREFLDPLGGGHLI
RGLDSEGDSHP
>Pirokinin-GU230851
MVSVSLVGLLLVALQLITNGCTQGTRRTLSKWHGDNDLTRQEETVMELLKDNPWAHFSVREGGRNTVNFSPRLGRDEEV
VFTETSRSPPFAPRLGRIVFRPRFGRLTLAAQH
>PDF
MNSGSVSVKELASWLLYLSQHHDQPHKRNSEIINSLLGIPKVLIDAGR
>Proctolin-JN543225

MATTTQSKVMSREVIVVAVLMMVLLSSSMVQSRYLPTRGADDRILRLRQLLKDLMENDLDPIMEHPAAPNGQYDPRLYK
RAAPPVQWDAVGAQFAGN
>IDLSRF-peptide
MFHNILLVLAEACHPYEPFKCPGDGACISIQYLCDGAPDCLDGYDEDSRLCTAAKRPPVEETSSFLQSLLASHGPNYLEKLFGS
KARDALAPLGGVDKVAITLSESQTIEDFGAALHLMRSDLEHLRSVFMAVENGDLGMLKSLGIKDSELGDVKFFLEKLVNTGF
LD
>Ryamide
MWLCQGLSLLLLLVSMLGTYGMEFYAAGGRYGKRPVQPPAIMRSREIRSGIFWTGSRYNRRGSDTRVAKQRKKDSFFMN
GRYGKRTGNTEDLCKYTNYQTLYPCNSR
>Short_neuropeptide_F-GQ452380
MKIALSALCCLIAVALMFTPETTSAPAIQDYDS
MRDLYELLLQREALPDSWAHKVVRKNNRSPQLRLRFGRRNDPTFLQGDHLMDNSMIDTL
>SIFamide-GQ253922
MSRTLFVCCFTLVVALIFLDAAMATYKKPPFNGSIFGKRAGPSSDYETAGKALSTMCEIAAEACSAWFPVQDNN
>Sulfakinin-GQ162784
MGSSFLITLLLAIGVYMFIENSHFMCLAEPAERRSLIRIRPEPALFAAEDDPLDIVDKRQFNEYGHMRFGKRGGSDEKFDDYG
YMRFGRSRP
>Tachykinin-GQ162785
MPVGSLLVMSCVLAACLAQERRAMGFVGMRGKKDTPDMEEYKRAPSTMGFQGVRGKKDDLIGEPDDTFLEEFKRAPAA
MGFQGMRGKKTPAMGFMGMRGKKDSDYGWWEEDKRAPASGFFGMRGKKAPASGFFGMRGKKGPSSSAFFGMRGK
KGPSGFMGVRGKKDSPDDLNHLLQLLRESALKQEMEEMLEDGRGLKRFAGLSDSFEDYPAELL
================================
NP, hormone receptors and GPCRs
================================
> Muscarinic_Acetylcholine _receptorA
MVSVSNATELLNLSTVQPLPEKVNYYTVTQIVLIAIVAGTLSIVTVVGNIMVMISFKIDKQLQTISNYFLFSLAVADFAIGLISM
PLFTISTLMGKWPLGPHICDTWLALDYLASNASVLNLLIISFDRYFSVTRPLTYRARRTTNKAAIMITCAWGISLLLWPPWIYS
WPYIEGQRTVPDNECYIQFIETNHYITFGTAIAAFYVPVTVMCFLYYRIWMETEKRQKDLPNLQAGKKDSSKRSNSSDEAVD
LEDWKRARGEGGGAKTVSGSTEQGNMEYGVVLEPHHYHKWRWVREWCIAWWHSGREEVDNDCELEDTPSDMGYAT
PVSVETPLQSSVSRCTSMNVIRDPYAGAGVPQTPRLAHHHVHYDSRSLPPTTRLQVYTILIRLPPDNSCDGVTDQQPSIKMI
SEDMSSTPVRAGVAGGVRGRVENSTADYNLSLHPAGGYHAAVNRRPSHISDIRIPLNAKIIPKQLAAGAKHPPPKKKKKTQE
KKADKKAAKTLSAILLTFIITWTPYNILVLLKPLTACTDCIPQGLWDFFYYLCYINSTINPVNQL
> Muscarinic_Acetylcholine _receptorB-NTE
FIATLCHYLLFSQFYYLFFSTTNCSFVYFIFRKLVKQKERLCRYVYQRLFTEQCRYTFNLSPAAYNSIWDSADSRYSAWNGEPL
GNGEIAPTNSTLVDTVANTSELYDNLTTSTADVLTPLPPFELWQTVLIAICLGLCIILTVGGNILVLMAFIVDRTIRQPSNYFIAS
LAATDMLIGTVSMPFYTVYVLMGYWNLGPILCDLWLSVDYTVCLVSQYTVLLITIDRFCSVKIATKYRGWRTKNRVIWMVTI
TWIIPALLFFISIFGWEHFVGFRDLLPGQCAVQFLKDPVFNTALIIGYYWTTLVVLFVLYGGIYKVAYDMQKKSEAKQRKMQS
MVALSAGAMSGMAGRAAGIGLSKTQSTLLSQDKPKTTTAPSTTTSTTTAPTSLTVAASTHSGGDSSSSQKTSSKTGTSNTEI
TSSTEKKKHQSGEGDKSERSSSPAFESDEDSTTCTTEKNTTYSKKKPSLAGMVAQTGALNMFAPSKVNGVIPKTTEQVKPST
PVDVPPLPQIPEATLLDTDTPGAQSTKTLTPILGNRDLSKSDKSSDKSLSEMSTPPISIQSPPSIIPPPSMFQSSTPASGCSPLSV
GRPRSLIMIKNTYDILTGMDNSDLRYMDESSVIVPTPNFESPPSTMSFPNTSSTAPPSPVHNSMTVSNTSLLQAALIRAAAQ
QVANNTPPKVSTPPITKVNTEVSLSGEDKAITSIVETAPDPETTVVDKLVGTPATLYSTASNSIATVQNVITAPPTPDPRKEST
RTTRSSSDRKDSSRRDFVKTIGKRLKGKGRSSGGSRQKSKSENRARKAFRTISFILGAFVACWTPYHILALVEGFCANPPCTNE
HLFMFSYFLCYANSPMNPFCYALANHQFKKTFTRLLKGDFHMT
> Muscarinic_Acetylcholine _receptorC
MCNKKDAVCLGGHHTPEQWAWVAADSILFVTILCGNLFTVYVLRSSRLMSNRFVLSLAVTDTLVGLTLPYHIAFTVFPQLSV
NKTTCILRFVFVLLACCCSILNVTAIAFDRYLAIVHPFKYECLMTKRVVRRVLLVALLNAIVISTVPIYWNTWEEASVCELGQVL
PRYYTIAVLLPAFLIIWSVMIAIYTRIWHEAVIQAEKLRRTNVCHQCKPSFQVVLLVMGCFTICWLPFLAVAAAQAVGYRDRI
SSNAYKAVLCLALSGSALNPLIYAWKNAEFKEAVRELMRCKNKEQQGSVMTITTGKMLAHPQTNV
>Dopamine1_receptor1-NTE
VPIEYNETTPILETDARGSDALATLTLLLIGIFLSILIFLSIAGNILVCVAIYTDRGLRKIGNLFLASLAIADLFVAALVMTFAVMND
LLGYWLFGTQFCDTWIAFDVMCSTASILNLCAISLDRYIHIKDPLRYSRWVTKRVAIISIACIWLLAGLISFVPISMGLHRPLSPP
VYYDGHPTCALDLSPAYAVVSSCISFYVPCIVMLGIYARLYCYAQKHVKSIKAVTRPLTSPSLPPMHHTTSSPYHVSDHKAAIT

VGVIMGTFLLCWVPFFCVNIVAAFCKTCIPGMAFKILTWLGYSNSAFNPIIYSIFNTEFREAFKRILTVRYAGCCCCRGYSSVVL
SLAETKPRTTAEIKQNGRKDFPTPRSSVGSIRQIRVNSTDKVIFDNEEISAI
>Dopamine1_receptor2-NTE
FGNPYENLSNQTYDNTTELEIWQQFSTMSQSKAFLASILLLFSLATVFGNSLVIAAVIRERYLHTATNYFVTSLALADCLVGLV
VMPFSAIYEVFQHNWFFGLNWCDIWRSLDVLFSTASILNLCVISLDRYWAITDPFAYPHRMSDRRACMLIAAIWICSMAISF
PAILWWRAVRTETVPGNKCPFTDNLGYLIFSSTISFYVPLFVMVFTYYRIYRAAVEQTKSLKLGSKQVMMTRGEIELTLRIHR
GRGMVYESPSLSDEVAELDRPLQNGAPRQTVANLLHPRHLGKNFSLARKLSKFAKEKKAAKTLGIVMGVFIICWLPFFVVNL
LSGFCVYCIYQEELVSAIVTWLGWINSGMNPVIYVCWSKDFRRAFARILCFCESRKLDQRHRGPLRAPYLGAHNMVLAPAS
VTYSTVCQHEMYSI
>Dopamine2_receptor-NTE/CTE
MNLSSGDLIEPLENWSIAVNWTTSNITGWLDEPNPPEKNYWALVLVLFPFLTLFGNVLVILAVYRERSLQSATNYFIVSLALA
DLLVAVLVMPFAVYVL
>Serotonin_receptor1A-NTE
NDSWLEENISTWNSSLSLDKDQYYDSVLMIVLSVVLGVVILITVVGNAIVIAAIVLERNLQNVANYLIVSLAVADLMVACLVM
PLGAVYEISQGWILGPELCDMWTSSDVLCCTASILHLVAIAVDRYWAVTNVDYIHTRNRGRIVGMIVVVWSVALVVSLAPQ
FGWKDPDYLDRITIQQRCLVSQDVGYQIFATCSTFYVPLLVILVLYWKIFQTARKRIRRRRTKQHTAQMKIDGKQKSSGPFKF
LSKKKLSLICVNWFRLVQEWETKRTGEVALVEGNVETDKEGSDAGATTAFTISTNHGHTPSNVSPERSTAELTANSKPPPPL
QPPSTPVVQTKPKRDAKKESIEAKRERKAAKTLAIITGAFVVCWLPFFIMALLMPLCHTCYINENLASLFLWLGYFNSTLNPVI
YTIFSPDFRQAFKRILCGVSHSRPRNLR
>Serotonin_receptor1B
MNSTVSQLVKRSARRPWRWPWKHRHLTTRLYEAWSTAATTAAAAVTAAHWPVGGGRNILPTISLSHATVANDINTSHAI
TVPTFGSPSSDIVLASSIQSPISTDIVALSGTVSDSPLLSWPDLRLNETLWFAELELANTSTTFIFGPSKELSVATIVVLAFILGLIIL
ATVIGNVFVIAAIVLERHLQNVANYLILSLAVADLLVACLVMPLSAVYEVSSEWTLGPELCDMWTSSDVLCCTASILHLVAIAL
DRYWAVTNIDYIHQRTGKRIGMMILIIWSVAFLVCIAPVLGWKDPGWEARISENRTCIVSQDIGYQIFATASSFYLPLLVILGL
YWRIFQTARKRIRRRFQATSQAALSSGRNPASGGVIAGLCAAGAAGGNGGIAAAVVGVIGRPLPTISEATTTTTTTAFTNVS
STNTSPEKGSYGNGIEADPTTMDIGSTSYQQYSSSHHHHAHHHQQHHHHPPKRKSKEAADSKRERKAAKTLAIITGAFVIC
WLPFFIMAILLPVCNMYQVCVLNEYLFAFFLWLGYFNSTLNPIIYTIFSPEFRHAFKRILCGRRNSLKRRTRQFQVITHL
> Serotonin_receptor 2A-NTE
MEWENTSNILWNITTILGNNSNITADDLERWSKHLQPTTAVTTIQQPTQQHSSHRDWTFLFVLLFVAAGGLGNILVCLAVC
LDRRLQNVTNYFLLSLAVADLLVSLFVMPLGAIPGFLGKQICKCFYLPQQYKDLEDGIVSLTGFGVSSDIMNGYNKKFRPVLT
PGQSILLETHNFDFNTFVVVSRIMDSTAIQNIIFHDTRFKLITLGTFYNNMAANVVATEQKASKVLGLVFFTFVLCWSPFFVL
NIIFAACPSCPVPTHVVDVCLWLGYVSSTINPIIYTIFNRTFRAAFIRLLMCKCSRWNRLARYRSVNEHRTGGGVNTPSTQTG
MGSAVPLSLSLQGTPILSPGTASTYLRTPSTFPDSFTVGEQQT
> Serotonin_receptor 2B
MCSDTTGQELCPLNWWALGAGVLVLATAAGNILVCLAIAWERRLQNVTNYFLMSLAITDLMVAILVMPLGILTLVRGYFPL
PPVYCLAWICLDVLFCTASIMHLCTISVDRYLSLRYPMKFGRNKTRRRVTLKIIFVWLLSIAMSLPLSLMYSQDYNSLVVGGIC
QIPDPLYKLIGSIVSFYIPLGVMLLTYALTVRLLAEQRQNLGGGGKAGSSGGTGWASGWLGQPTPGLERRGTWKRLLGKPG
PLVIRGQHTSSAGSTDTELTTLDTHDLWIPEPDPKPYATSALQQFGAEMLKLSRGFESVAVAIESKPLNQRTSKYVSIQRGGH
SSIDDALRLDNKKQRIRKRRKANEEDRWPIKARRRASTMELIRRDSEKVIKRAVSYHERVCSEGSSSSSRSSGDSASSEEDEG
QSPSVQQQEHQGHASLSPQSPPPQININTVQKSQQQAQADDEDPLSLLPPPCTCPYFGDSENKKPIRNNEVVIITSEEKPAF
LRRDDHKFKYDSSNTIVTWDSPPKRRSRRGSSFSSGSIRTTLASNESSPAIRKPPILRRSATLRTGRANDFKAMEEAAQGVLLR
YGSNQTMGRVIKSPHSRNSSVLSRTSSRHGRIIRLEQKATKVLGVVFFTFVILWAPFFVLNLVPSVCAECERNIDKWVFDFVT
WLGYASSMVNPIFYTIFNKVFRQAFKKVLLCRYRNQRWTPRT
> Serotonin_receptor 7A-NTE
VAGNILVCVAVCLVKKLRRPCNYLLVSLAVSDLCVAILVMPMALLYELMGTWNLGPIMCDIWVSFDVLSCTASILNLCMISV
DRYYAITKPLEYGVKRTPKRMMACVSVVWLGAACISLPPLLILGNTHGEETGQPGTQCLVCQDFGYQIYATLLSFYIPLAVM
MFVYYKIFRAARKIVLEERRAQTHLETSHCFLEINKNGSGPESRLAVSCASSPAARAHHRTSTTSTNTTVSIHFSCSGSVVGVA
RVSNESQCPMLSVAAAAVNAAASTRTRRRLQLQSSAPSRKLRFQLAKERKASTTLGIIMSAFTVCWLPFFVLALVRPFLHDQ
DAIPASLSSLFLWLGYANSLLNPVIYATLNRDFRRPFQQILYFRCGSLNHMMREEFYQSQYGDPAPTHFNEDEPTCDLMPVS
DLKGAGHESFL
>Octopamine alfa 2 receptor
LTKNVLFVSLLQLSKDIGYVLYSALGSFYIPSCIMVFVYIRIYYAAKARARRGIRKAVARPRPAEKVTSFSKKEPLTSPVDRNSNN
SPVAVTVEKPVIPVVTCDFASDISTSDNIEQPDPTAPKDTLNVSKLPTCSLTPNVTFKGSTLSVNGDLAAMSRCRAPSVGIDV

DMVSEFDPSSSDSGVVSRCAVVKPLKLRLCKPIFGRKSSKAKREVIDMGRVISTTGSQEIPQEIPKVQKPRDPEREKRRLARKK
EKRATLILGLIMGSFIACWLPFFFLYILTAICSACQIPDFAFAVAFWLGYMNSALNPVIYTIFNKDFRRAFRRILFK
>Octopamine Beta 1 receptor-NTE
VNFTENQQEFATEFQTWQLIFVIKSTLMGFIILAALFGNLLVIVSVMRHRKLRVITNYFVVSLALADMLVAIWAMCFNASVE
LTDGKWLFGYFMCDVWNSLDVYFSTVSILHLCCISVDRYYAIVQPLDYPLIMTFKKLVIMLAVVWISPALVSFLPIFMGWYTT
EKHLEYRRSNPDLCEFLVNQAYALISSCVSFWVPGVIMLFMYYRIYIEADRQERMLYRSKVATALLNKHLQINGIAAGMEVG
AAEQLRPLATTSKMKRERKAARTLGIIMSAFLACWLPFFLWYLITALCGPDICYSPHLVVAAVFWVGYFNSALNPIIYAYFNR
EFRAAFKKTLESCCHQMTMCFRSSETQRRRALRGTNAVNESRSNEAVRAQMVGSSNVSSAAEIHYLNSACVVRTTTADD
MAKLSTAVNNLNLEQGI
> Octopamine Beta 2 receptor
MESMEAGDVLKGLRGGHMVTSSSTSSTTVLPSLVAQIDFNGTNVTMQDFEDVDQSWAVILSTLIKGTIMGAIIIAAIFGNLL
VIVSVMRHRKLRIITNYFVVSLAFADMLVALAVMTFNASVQLTGRWMFGPFMCDAWNSLDVYFSTASILHLCCISVDRYYA
IVKPLRYPINMTKQVVAIMLLSTWIAPAVISFVPILCGWYTTNENQEFRNEHPDICHFKVTAIYAVVSSLISFWIPCTIMVFTYL
AIFREAIRQEKQLHARMGNQMLLNNSSRDCNGDTLGSSGGSSKALALNEVGLHSTPTKDRNIIKMKREHKAARTLGIIMGT
FILCWLPFFLWYVIVSVCGEEACPTPEIVVSTLFWIGYFNSTLNPIIYAYFNRDFREAFKNTIQCAFCSLCRRPPSDLESLEVRRP
SLRYDERTRSIYSETYLKHIDRRRSSEVGSSL
> Octopamine B3 receptor-NTE
MFNDMADSTPSAYLEAYPNSTGAELVEGVSVAPFLLKGFLMGLIIVGAVLGNALVIISVVRHRKLRIITNYYVVSLALADLLVA
LCAMTFNASVTLTGSWLFGPFMCDVWNSLDVYFSTASILHLCCISVDRYYAIVRPLQYPITMTHRTVSFMLANVWVWPAII
SFTPIFLGWYTTSEHQEFRKTHPHICIFVVNKYYAIISSCVSFWIPGVVMVTMYFRIYKEAVRQRKALSRTSSNIILNSIHQHRR
GHPHYAPRLLPPSNEGTTMRTATSWRSEHKAARTLGIIMGVFMICWLPFFLWYVITTLCGQVCYCPDVVVAVLFWIGYFN
SALNPLIYAYFNRDFREAFKNTLQCAFPCCFTCCPKEDTSPMQYV
>Oamb-like-CTE
MVWIFGDFWCSAWLALDVWMCTASILNLCAISLDRYVAVTRPVTYPSIMSNRRAKILIAAVWVLSFLICFPPLVGWKDTTQ
NTVIPEGPGRPLTPGGATVIQWPTAPPTEEPPCPWRCELTNDAGYVIYSALGSFYLPMFVMLFFYWRIYRAAVQTTRAINQ
GFKLTKGNRGIGNRFDEQRLTLRIHRGRGSVMQGSSGSNSTTTTNSAANSVTNSPGGCGGGGGGGGGSGGGIGGGGKS
PEKKNTRRHERIKISVSYPSSDAISAVNNNSPPPTSPKSSISSNSPPPSGQLFAVHYTGNETSSVYRKDPNCHLRVSGNRLASH
RRARRTSSEGQPTRPRLLGDPLSVTIQKDLSPSPTYDENNPAKPKLISRMGKRNIKAQVKRFRMETKAAKTLGIIVGGFIFCW
LPFFTIGKFY
>Octopamine-Tyramine receptor
MKMTDWDYTERYYNASNITGELAGCPEPERVFYDTTLGISFAVPVWEAIATAILLSLIIILTIIGNILVILSVFTYKPLRIVQNFFIV
SLAVADLTVALLVLPFNVAYSILGRWEFGIHVCKMWLTCDVMCCTASILNLCAIALDRYWAITDPINYAQKRTLKRVLIMIAG
VWLLSMLISSPPLAGWNDWPDVFDSSTPCQLTSQQGYVIYSSLGSFYIPLFIMTIVYIEIFIATRRRLRERARASKLNAVKQNY
QQNSIKEKHSPVDGESVSSEANEHEHKEKKKANKKKRKKSDQSGTYLAPAQVAEDSFTDVHDISSVNSPQKQNKKNNDEY
RDSSKKDVHVPVMAVVNQTKRAVQVNQFIEEKQRISLSKERRAARTLGIIMGVFVVCWLPFFLMYVILPFCSVCCPSDKFIN
FITWLGYINSALNPIIYTIFNLDFRRAFKKLLHIKSNT
>Dopamine Ecdysone receptor-NTE
VHLITLEPYTQAAALTSIAVAIVLANLTIIAAFLNSRGPGEVINCYLLSLAVADLLCGLLVVPLSVYPALVQRWVYGDLVCRLVG
YLEVTLWAVSVYTFMWISVDRYLAVRKPLRYDTVQTKIRCQCWMALTWVSAAMMCCPPLLGFNQPVFDNDSLVCLLDW
RDMAAYSATLAVLVLGPSVITITYTYSYIFMMMRKVKSGAPIHDKEYATALSENLTNPSHVMSFVLVMMFCVSWTPFIVVR
AYEGTTGSRMSLIPHLHFAVFWLGVLNSVWKAVVLVFLSPQFRLMLRILGFTLCCRHRARMQMELIGMEDE
>RproOrphan1
MKNLNSLSGIEKKELFDSIDTVLFDCDGVIWLSSEVIPGALDAINGLKRVGKRIFFVTNNSTKSRADLLKKSLKLGFQVTVMVT
HERRLSTNASRTMAAMSLGFIVMVTPWTIQEVVVACTGTKMPEFLEFTCTWLALSNSFWNPFLYWLLNNNFRRISKELFM
TKILCRGKKTPPSNSLQCCSVPGSVPAPLTRDIEGLSEKYWGEILERTVSSNSLQMQKMYPPPPPPPPRSTK
>RproOrphan2
MIRRRLFVFLISSKTSGKQMVVYHSELTVLRSSKKHCCHMTSFSMLIALMYNGQIEVNFEFYYKFITIPLLVFCAFSVLANALIL
ASLRWIRRPLSPTLHISLSLAGADMFTSLVIGIGLIVNSLLPQVFSIQIDKCSQLVIEALRMGGMYTSIGHLLTLAVNHYLGIKKP
LHYPSLMTTRNITVIVLALWIIPPSSFAIYFSLLEQDGFAIIGCDYEFITRAEFRSSCCYIFVVCLITMAAIYSHIYLLVRQHQANRN
RFRRVGSSYSRNPTSANDLRQSRNEKALRTTLLIVGTVVLGWLPATIHFYLVCDNCFIKLNWGTPVLRIITSALVNFLILLKTAL
NSCIYAARMHEIKAAIRLMRDRFITCCKFGTDQNEHNGLNSESSRHFVSRASIRPSRGTVLCRLQSLPRNEGPPRQHAHYTT
NL

>AKH_corazonin_related_peptide_receptor1-KM975506
MDPLFSNFTIEFNYTESIYYSPTNNTLYELPKFDDNALIVVIAYSLLFIIAAIGNLTVFITLVRGRHRKSRISLMITHLAAADLFVTF
IMIPLEIGWRLTTQWVAGNIACKLFLFLRAFGLYLSSNVLVCVSVDRYFAILHPLRVSDARRRGKMMLTMAWIFSLICALPQS
VVFHVSQHPQHPDFWQCVTFGFFGSRTQEIAYNLFCVMAMYFVPLLVIVIAYTCILLEISKKTKETRGPTSVPDLTNGDRCE
GITSKNKRTSGEQVTKKEPEGGCASEEVICQILKGPELAHLG*
>AKH_corazonin_related_peptide_receptor2-KM975507
MDPLFSNFTIEFNYTESIYYSPTNNTLYELPKFDDNALIVVIAYSLLFIIAAIGNLTVFITLVRGRHRKSRISLMITHLAAADLFVTF
IMIPLEIGWRLTTQWVAGNIACKLFLFLRAFGLYLSSNVLVCVSVDRYFAILHPLRVSDARRRGKMMLTMAWIFSLICALPQS
VVFHVSQHPQHPDFWQCVTFGFFGSRTQEIAYNLFCVMAMYFVPLLVIVIAYTCILLEISKKTKETRADQWRTGHEERTRG
RMRLRRSDMSNIERARARTLRMTVTIVLAFIWCWTPYVVMTLWYMFDRESAEKVDPRLQDALFIMAVSNSCMNPLVYG
SYALNFRRECTTCFCYLFSSHQQLDRRSTDAAAHTSRVLRVPKCVLTFVRGSGITRSTAVTGYGGTLGSRNHLTVPRKNVMR
PASAEHLVIRGRMIETAPLNPEEFHSDPGTNTGIYLVTS*
>AKH_corazonin_related_peptide_receptor3-KM975508
MDPLFSNFTIEFNYTESIYYSPTNNTLYELPKFDDNALIVVIAYSLLFIIAAIGNLTVFITLVRGRHRKSRISLMITHLAAADLFVTF
IMIPLEIGWRLTTQWVAGNIACKLFLFLRAFGLYLSSNVLVCVSVDRYFAILHPLRVSDARRRGKMMLTMAWIFSLICALPQS
VVFHVSQHPQHPDFWQCVTFGFFGSRTQEIAYNLFCVMAMYFVPLLVIVIAYTCILLEISKKTKETRADQWRTGHEERTRG
RMRLRRSDMSNIERARARTLRMTVTIVLAFIWCWTPYVVMTLWYMFDRESAEKVDPRLQDALFIMAVSNSCMNPLVYG
SYALNFRRECTTCFCYLFSSHQQLDRRSTGSGITRSTAVTGYGGTLGSRNHLTVPRKNVMRPASAEHLVIRGRMIETAPLNP
EEFHSDPGTNTGIYLVTS*
>Adipokinetic_hormone_receptor-AIJ49751
MTRTEEVFSFTFFPEWSEIRTEKNETYVIPPDMRFNEGHKLALAVYSLLMLISGVGNVWVLVRLAKSRRSRTNRMLTHLAIA
DLFVAFLMMPAEILSAATVAWWFGDIPCRIFAFFKTFGLYQSSFVLVCIGIDRYYAIVKPLSIKDTYCRGKGIVALAWVISGICS
LPQVVVFREQEHYTFTGYKQCCTFNAFPTSSHEIAYSMYNMAMMYMLPLVVIIFCYGSIFIEIYRRTSAQNSGKLRRSTLGFL
GRAKNRTLKLTITIILAFFICWTPYYIMALWYWLDRSTAEGVDVRVKRALFLFACTNSSINPLVYGVYQRTGCGPNNSRTSHN
TCITELRQQRTNNHVRENLG
>AstA_receptor_KM283241
MNGSPATAIVEAIGSMPPIKYDPNNNFTNNTINFNNNIHNFNNNNIDMRNFYSNVTDTEIFMEEISPELTEKIVAIVVPVLF
GIIVILGLFGNALVVIVVAVNQQMRSTTNILIINLAIADLLFIVFCVPFTATDYIFRFWPFGDTWCKMVQYLIVVTAYASVYTLV
LMSLDRFLAVVHPIASMSIRTEKNAISAILVTWIVIVISNIPVFLCHGEVTFNYSSSEHTVCIFLEMDPLIRPDGFNKVAFQVSFF
ATAYVIPLALICGLYLVMLVRLWGGAAPGGRCSAESRRGKRRVTRMVLVVVAIFAICWCPIQVILVMKSIGQYEITPTSVMV
QIVSHVLAYMNSCVNPILYAFLSENFRKAFRKVIYCGPEGGSHPPHLNGRQIDAEKSALTKNTRTTDIL
>Allatostatin_C_receptor
MSKDTTVSWLADSLENGSIESLYNSSYGNETQFCGSTDQPTLHIFTQVLYAFVCIVGLLGNTLVIYVVLRFSKMQTVTNLYIV
NLAVADECFLIGIPFLIATMSLQLWPFGNVMCKLYMASTSINQFTSSIFLTIMSADRYVAVCHPITAPKMRTPFISKIVSLSAW
TASAIFMIPIFMYANIMDDDQVKSCNILWPEGENLSGQTAFTLYSFVLGFAVPVVLIFCFYFMVIRKLQTVGPKNKSKEKKKS
HRKVTKLVLTVITVYVLCWLPYWITQMALIFTPPKQCQSKFTVTVFLFAGFFSYSNSAMNPILYAFLSDNFKKSFVKACTCAA
GKEVNATLHLENSVFPRRTQRGGSERARAGKNRADHTDEGAETGPLVSRGEHSTTALTSRSNITVTSDTTTPVKNGVKINLT
PTEL
>Allatotropin_receptor_1_KF740716
MTEHIFPTVYEWILIGMHAVVFAVGLTGNFLVCLVVHRNPAMRTVTNYFIVNLAVADFLVILICLPPTLIWDTTETWFLGHV
LCKLVLYLQTVSVAVSVLTLTFISLDRWYAICFPLKFKSTTSRAKTAILIIWIIALLYDIPELITLRTASRKKFHVETVLFTQCIASWD
DVAERHYTTSKIVFLYLLPLTITSAAYFQIVRVLWKSDNIPGHRYQREVCYISGSSVDSRRYMAVSRGPTSGGTQAQIRSRRKA
AKMLVCVVLMFALCYFPVHLLSILRYTVDIPQNDITVALAMLSHWLCYANS
>Lutropin-choriogonadotropic hormone receptor
MIKNNNPLLLTAYILVHRMMYVCDRTSAENLSKKFGTQINQLWDNFGTDFTYPGNLPAYVEEYFEEQEYNNKQTDPPPAKI
QCLPTPGPFLPCVDLFDWWTLRCGVWIVFLLAMLGNGTVVFVLIFSRSKIDVPRFLVCNLAAADFFMGVYLGLVTLTLVDA
STLGEFEMYAIPWQMSAGCQLAGFLGVLSSELSVYTLAVITLERNYAITHAMHLNKRLSLKHAGYIMLCGWSFATIMATLPL
LGVSDYRKFATCLPFETSTTWSLTYVVFLMFINGVAFLILMGCYLKMYCAIRGSQAWNSNDSRIAKRMALLVFTDFLCWSPI
AFFSLTAAFGLQLVSLEQAKVFTVFVLPLNSCCNPFLYAILTKQFKKDCVLICKAIEESRVTRGIGRCRHSSNFSNRQTPANTNS
LMDRSSRDQGHQPCSCNTKLLGDSSSKRTPKRWVATKLYWLSSCMNRESNSRHRTRSDQYAYQIAEIQQKQHKRASSVSS
SENFSSSRSDSWRQNHYHCGIPMRLLDPKKRASSWIVTRKTSQDSNLSSSRNDSSGSATTNSTTMSRVSRSSNSSDIRPKPR
LTRQSAVVDETEGPPGSPARLTVRFLTTIPSAAESSMQMDEESASACYAILHTGPEPGTNAEQTSSPKEDEIKETNQRSNTIY
AYTVRCRCRSDFIIHLHNFNRPQLHRCTGHRR
>CT/DHR1b-AHB86317.1

MSDETGNQSFLDPHAELVNSRYLQCLTTINESLSRSLQGLQCEATFDGWSCWPATSAGETAFAPCPHFITGFDPNRLAHKE
CTENGTWFRHPESGQIWSNYTTCVNLDDLNLRQQVNNIYQAGYFISLLALLLSLFILSYFKSLRCPRNTLHMNLFTAFAFNNF
LWLLWYRLVIPFPEVILENGVWCQCLHVILHYFLLSCYAWMLAEGVYLHTLLVSAFTSEQKLVKVLTVLSWFVPIVFITLYTTL
RLASGHTDQCWIDESDSNTVLIILVATSMGLNFIFLCNIMRVVVGKLRAGPAQSSRPSRALLQALRATLLLLPLLGLNYLLTPF
RPPNNHPWETYYELISAVTASFQGLCVATLFCFCNGEVIAQIKRKWQYAMFRTRANSYTATTVSFVRSNAAPVAEEENV
>CT/DHR1c-AHB86318
MSDETGNQSFLDPHAELVNSRYLQCLTTINESLSRSLQGLQCEATFDGWSCWPATSAGETAFAPCPHFITGFDPNRLAHKE
CTENGTWFRHPESGQIWSNYTTCVNLDDLNLRQQVNNIYQAGYFISLLALLLSLFILSYFKSLRCPRNTLHMNLFTAFAFNNF
LWLLWYRLVIPFPEVILENGVWCQCLHVILHYFLLSCYAWMLAEGVYLHTLLVSAFTSEQKLVKVLTVLSWFVPIVFITLYTTL
RLASGHTDQCWIDESDSNTVLIILVATSMGLNFIFLCNIMRVVVGKLRAGPAQSSRPSRALLQALRATLLLLPLLGLNYLLTPF
RPPNNHPWETYYELISAVTASFQGLCVATLFCFCNGEVIAQIKRKWQYAMFRTRANSYTATTFVRSNAAPVAEEENV
>CT/DHR2-GC4395-Ortholog-AHB86571.1
MGNVDNLTTQSNLHSNILRYLKGLQRECDLRKRAQFQYLSTILPNVSDLKVYCPATFDGWSCWNTTPSGEIALAPCPNFVT
GFDINRFAFRKCLENGTWFRHPDTGQPWSNYTTCIDMDDLKFRKAVNTIYVVGYYISFAALVLSLIIFLMFRSLRCTRIAIHV
QLFSSFAANNLMWIIWYKTVVGNTSVVQENQLICQVLHVILQYFMVANYLWMFCEGLHLHLALVVVFVKDDSAMRWFY
CIGWFLPAILTAIYAWVRSANPDDTRQCWMNESYTQWILIVPVCLSLFASLGFLINVVRVLLTKLHCNSANPAPVGLRKAVR
AALILVPLFGIHHILIPFRPEPKAPGERAYEIFSALLVSLQGFCVSVLFCFVNVDVHCAFKAMVRRIQRRAADNGNLTATQTRE
VM
>CT/DH
MLNPAEQGLSVEEVIALRRSDCLLLDNQTDVLECPKEFDGWTCINSTAAGTVAHFPCPYFIFGFDPTRFGHRTCLEDGTWF
RHPASNKTWSNYTTCVDLEDLKMRTQVNMIYKGGYAISLAALTLSIFIFFYFKSLTCTRIQIHKSLFISLAVNNLLWLIWYEAVA
DNLPVLFANGFGCQLLHILVQYFLVATYLWMFCEGLYLHTLLVVTFVTESKVMPFLHLIGWGVPAILVTIYAILRMSNKEDSV
HCWIHESLYSWTLSGPVLVSMIANFVFLINIVRLLLTKLHTTAQTSPRSSFKESKNASIRSKRSTISTQSAPSGRTKKAVRATLILI
PLLGLQYIVTPFRPNQGTPWEYAYQVTSALVASCQGLCVALLFCFCNGEVVAVMRKKWRQCRISKKRPWHSCSGVTSVSR
HRRI
>CRF/DHR1
MEANYTKSNARLLEEAEKCLEQLEADGIPPADYCPRSWDGILCWPPSPSATIVYLPCFEELHGIKYDTSQNASRWCLWNGS
WANYSDYDSCSHLQIPPFAADPGLVVVTMIYLIGYGISLIALCVAVAILIYYKDLRCLRNTIHTNLMCTYILAAFMWILNFTLQ
MSMDTDMVSCIILVILLYYFHLTNFFWMFVEGLYLYMLVVETFNRENIKLRAYLAIGWGSPVAVVFIWAISRSFIGDESNSES
KSNVQRGCAWMSPNSSDWINQAPAIIVLAVNLIFLVMIMWVLITKLRSANNVETQQYRKAAKALLVLIPLLGITYILFIVGPT
EGQYAVIYSYIRALLLSTQGLTVALFYCFLNTDVQNTVRHHLSRWREARDIDARRYTHTKDWSPNTRTESVRLCAKKGRTPR
HYKKRESTVSETTTLTIVGYSSTNRVSNGSMIATTLTVRAPGLHPSEAENAV
>CRF/DH-R2A-ANJ03339.1-NTE
GLLCWPNTPPGVTAYLPCVAEIDNVKYDTNQNASRICYENGTWANQTDYGLCSELHTLTSNQILSDEGIIVQSTIYAVGYGF
SLTALGLAVWIFLYYKDLRCLRNTIHTNLMCTYILADLMWILSSIQVYVKTDPAICMVLFILLHYLILTNYFWMFVEGLYLYML
VVETFTRENINLRAYLAIGWGIPVIIVIPSCLARAFISDDYEYVLITKLRSSNNAETQQYRKATKALLVLIPLLGVTYILFIAGPTEG
PYAYLFSYIRAFLLSTQGLMVALLYCFLNTEVQNTVRHHFTRWKESRNLGARRYTCSKDWSPNTRTESVRLCSKHDVMPYR
KRESVASENTTMTLVGGSTNLARLSNGSTGQTVRTPVSLYLEPNNSNNAL
>CRF/DH-R2B-ANJ04995
MSTDGNFTDPTIKLGEEQEMSIDVNFTDSIIKLREEVEKCFNLSITEVPPSEEYCTTTWDGLLCWPNTPPGVTAYLPCVAEID
NVKYDTNQNASRICYENGTWANQTDYGLCSELHTLTSNQILSDEGIIVQSTIYAVGYGFSLTALGLAVWIFLYYKDLRCLRNTI
HTNLMCTYILADLMWILSSIQVYVKTDPAICMVLFILLHYLILTNYFWMFVEGLYLYMLVVETFTRENINLRAYLAIGWGIPVII
VIPSCLARAFISDDYEYGLIGHHEGCTWVVSNSSDWIYMTSSIIVLAVNVIFLIMIMWVLITKLRSSNNAETQQYRKATKALLV
LIPLLGVTYILFIAGPTEGPYAYLFSYIRAFLLSTQGLMVALLYCFLNTEVQNTVRHHFTRWKESRNLGARRYTCSKDWSPNTR
TESVRLCSKHDVMPYRKRESVASENTTMTLVGGNNSSPQNKTINYEYVRTPVSLYLEPNNSNNAL
>CAPA_receptor_variant_A-ADG27752
MNSFDIIETVTNSTPVNVSLEEYLIIVRGPKFLPLKILLPITFTYGILFISGLFGNLAVCIVIAYNKSMHNATNYYLFSLAMSDLVLL
LLGLPNDLSVFWQQYPWILGLLVCKLRALVSEMSSYVSVLTIVAFSVERYTAICYPLKSYTTDKLNRVIKVIGTLWLISLGFAAP
FAIYTTIDYVDFPPGSGKAVIESAFCAMLKQNVPADVPLYELSCTLFFICPAVILIFLYVRIGLTIKNNTKLRGNVHGELQSIQSK
KSIVSMLMAVVVAFFICWAPFHMQRLIYVYMSDYPWYGIVNVWLYYISGIFYYFSATINPILYNLMSLKYRKAFKQTLWCRK
YNRIIKTPGLRETNSTSRQVNKSIKSMNMQHNQSLLANNIEDIT
>CAPA_receptor_variant_B-ADG27753
MNSFDIIETVTNSTPVNVSLEEYLIIVRGPKFLPLKILLPITFTYGILFISGLFVLLLLGLPNDLSVFWQQYPWILGLLVCKLRALVS
EMSSYVSVLTIVAFSVERYTAICYPLKSYTTDKLNRVIKVIGTLWLISLGFAAPFAIYTTIDYVDFPPGSGKAVIESAFCAMLKQN

VPADVPLYELSCTLFFICPAVILIFLYVRIGLTIKNNTKLRGNVHGELQSIQSKKSIVSMLMAVVVAFFICWAPFHMQRLIYVY
MSDYPWYGIVNVWLYYISGIFYYFSATINPILYNLMSLKYRKAFKQTLWCRKYNRIIKTPGLRETNSTSRQVNKSIKSMNMQ
HNQSLLANNIEDIT
>CCHamide_receptor_1
MEYKDQLNNDSLVNTTENAITVIPYSERPETYIVPIVFAVIFLVGVLGNGTLVLIFIRHRTMRNVPNTYILSLALGDLLVIISCVP
FTSTIYTVNSWPYGLFICKLSEATKDVSIGVTVFTLTALSADRFFAIVDPMRKLYSSIGGRGATRCTIMIACAIWLLAIACAIPGA
LFSYIRIFKQGNHTLFEICYPYPEELGSVYPRGLVMAKFLIYYAIPLTVIGCFYILMARHLVLSTKNMPGELQGQARQVRARKK
VAKTVLAFVLVFAVCFLPQHVFLLWFYNNPNSDRDYNEFWHVFKIVGYCLSFINSCINPIALYCVSGTFRKHFDR
>CCHamide_receptor_2
MDPETVEDFKYVSSVNSNISNVEEYTPYPERPATYIVPIVFAVMFLVGVLGNGTLVLIFIRHRTMRNVPNTYILSLALGDLLVII
SCVPFTSTLYTIESWPYGGFVCKLCEATKEISIGVSVFTLTVLSAERYCAIVNPIRRHISTKPLTIVTVFCIWVISFLLALPAAIFTHV
SKANITNGRTIEFCSPFPEEYGPTYRKLNVLLRFIIYYAGPLLIIAWFYILMARHLLLSTKNMPGELQGQSNQIRARKKVAKVVL
VFVIIFIICFLPHHFFMLWFHFNPDSDEEYNLFWHVLRIVGFCLSYLNSCINPIALYCISKAFRKHFNRYLLCSFVRDSTLDEISLG
NMNSSTKHVRQSSIITSHYTITQSEKT
>CCAP_receptor_1_KC004225
MDWVIRDNYSNPAANITNTTDEINSFYFYQTEQFTVLWLLFAAIVLGNSAVLLALLFNKSSKSRMNFFIMHLAFADLSVGLIS
VLTDIIWRITVEWKAGNVVCKVVRFMQAVVTYSSTYVLVALSLDRLDAITRPMNFSGSWRRARLLVGFSWALSAFFSSPILIL
YKERLIQGSFQCWIELGSTLKWQIYMSLVAVSLFLVPALVITACYTVIVYTIWTKSIHISRDSQQSTPLKNGGDKGDDNDIRRA
SSRGIIPRAKIKTVKMTFVIVFVFILCWSPYIVFDLLQVFGYVPRTQTNIAVATFIQSLAPLNSAANPVIYCLFSTHICRALSKLPP
FSWICCCFANRGAESSSIIDTVTSTLRRATIRNQDNL
>CCAP_receptor_2-NTE/CTE
LIIFFRKIDELIEVKENHVRGLQSWLGQTEQFAVLWLLFLLIVCGNSAVLAALKCAKKPKSRMNFFITQLALADLCVGVLSVLT
DIIWRSTIAWNAGNIACKVIRFSQMAGKTLVFIVCWSPYFIFDLLQVYGYVPTTQTNIAVASFVQSLAPLNSAANPLIYCLFST
RICRGIRVSTPQKIGKGITE
>CNMamide_receptor-NTE/CTE
MTKSSSQFARNLASTNSATIVGNMAAGTSNLVTTRTRTPSSQTKVTEMLLVVSTVFIILNLPSYVVRVWIYLTDTHNVGTEQ
KVTMYVLQQYCNILFNTNFGINFALYCISGQNFRRALLSLFRPEIQRRSGETTQTTED
>Corazonin_receptor_VectorBase_isoform
MQTLFPNISDETTLRQLQDHLINSDDGHRFILPLELCDLWNITVSSSSRIQCLEHAPQLTSSARTRAIVLGVMAVISFIGNVLTI
ISIRSSRRRRRRNQNWSAVYALILHLSVSDLLVTIFCIAGEALWSYTVAWTADNVTCKLFKFSEMFALYLSTFILVLIGLDRFVAV
RYPIKAISTAKRCGRFVAGAWFLSFLLSLPQVFIFHLSKGPFYEEFYQCVTYGFYTEPWQEQLYTTFSFVCMFMLPLLILIISYVS
TIITISRNDKMFRDESNNTSATRKLDINRRRLIHRAKMKSFRISLVIVVTFIVWWTPYYTMMIIFMFLNPDKHLSEELQKGIFFF
GMSNSLVNPLIYGAFHLWRPSKKTGSARSVSVYFLLLLIFFL
>Corazonin_receptor_isoform_alfa-AND99324
MQTLFPNISDETTLRQLQDHLINSDDGHRFILPLELCDLWNITVSSSSRIQCLEHAPQLTSSARTRAIVLGVMAVISFIGNVLTI
ISIRSSRRRRRRNQNWSAVYALILHLSVSDLLVTIFCIAGEALWSYTVAWTADNVTCKLFKFSEMFALYLSTFILVLIGLDRFVAV
RYPIKAISTAKRCGRFVAGAWFLSFLLSLPQVFIFHLSKGPFYEEFYQCVTYGFYTEPWQEQLYTTFSFVCMFMLPLLILIISYVS
TIITISQSDKMFRDESNNTSATRKLDINRRRLIHRAKMKSFRISLVIVVTFIVWWTPYYTMMIIFMFLNPDKHLSEELQKGIFFF
GMSNSLVNPLIYGAFHLWRPSKKTGSARSGREHATYSLLKRTRTWHTPGGRDYLCLNRTNGDSKQFQQISVLLPNNKDPITI
TNKVSKLGLSRSMYRIAS
>Corazonin_receptor_isoform beta-AND99325
MQTLFPNISDETTLRQLQDHLINSDDGHRFILPLELCDLWNITVSSSSRIQCLEHAPQLTSSARTRAIVLGVMAVISFIGNVLTI
ISIRSSRRRRRRNQNWSAVYALILHLSVSDLLVTIFCIAGEALWSYTVAWTADNVTCKLFKFSEMFALYLSTFILVLIGLDRFVAV
RYPIKAISTAKRCGRFVAGAWFLSFLLSLPQVFIFHLSKGPFYEEFYQCVTYGFYTEPWQEQLYTTFSFVCMFMLPLLILIISYVS
TIITISQSDKMFRDESNNTSATRKLDINRRRLIHRAKMKSFRISLVIVVTFIVWWTPYYTMMIIFMFLNPDKHLSEELQKGIFFF
GMSNSLVNPLIYGAFHLWRPSKKTGSAREGSTLHTPCSRGLGRGTLREEETTFVLIEQTATANNFNRSASFCQTTRIQ
>Ecdysis_triggering_hormone_receptor
MISGLALVGDDHNDTAAYLFNINLTSFYTNLTVNLTNETGFVNGSYGEQTLVFPSYIRTASMVLCIVILGIGVVGNMMVPLVI
IKTKDMRNSTNIFLMNLSIADLMVLLVCTPTVMVEVNSKPETWVLGEEMCKAVPFVEMTVAHASVLTILAISFERYYAICEP
LRAGYVCTKTRAMLICLKAWVFAALFTSPVLVLADYRDEEYKDGSIEKVCLLQVDTFWKSFYFVMSITLFFVLPLGILVILYSIIA
RHLMSNACLAASSSHISNLRYRRQVVMMLGAVVLSFFIFLLPFRALTLCIILAPPGFIFSLGMEKFYNILYFSRIMLYLNSAVNPI
LYNLMSSKFRDGFKRLLGINHKDGLARKGTVTTTTLTSSRKCSTDHHLIKRSVVRVISVEEKLDNGITTTINLIGHNGITKGDES
YV

>FMRFamide_receptor
MNFTNTNFSLDDNETAYISTDNENDDKSEILFEFITNGVLLNLVGILGIMGNIISMVILSRPQMRSSINYLLTGLARSDTVLIITS
ILIFGLPALFKYTNSQLLFSYYYRVYPFLAPVVYPLAVIAQTVSVYLTLTVTLERFVAVCHPLQARSLCTYGRARLYVLLIIIFSILYN
LSRFWEVKLEQEYLVQYNVTVYIPLPSSLRSNQIYISVYIHWLYLLFIYFLPFSCLAVLNAAIYRQVRKANQERQRLSRLQKKEIG
LATMLLCVVVVFFICNILALVSNVLEAFYGILLTKMVKTSNLLVTINSSVNFIIYVIYGEKFKRLFLKLFCSHSPRICDGGVRESPD
CATLHEDSVMLSNGDARHSVRGNRANESVKRAARALPCVYYPARHNSKWNQDDTTTTTLNQI
>FaLP/Proctolin_receptor-NTE
MGNVVTVIVMTRKRMKSSTNTYLTALAVSDLLFLIFNMILSFEHQPAIRQSQYVTYWHLHKWTIWLVDATGACSNWLTVS
FTLERYIAVKHPLRGKVLCTESRARKVIXTCNITSLPLNNYYSIMECRRDRRVALQSTWLGQHPTYKSVFYWFSSITVTAIPLAS
LSVLNYLLVAAVRRSTKGRNQLTEDARGVRGSFRPTCNAGNSIYSRCGSERSPPIQGMSQRRVLKERQENKVTIVLISVVFLF
LICQAPSAITVIVKVFYEPESDTSGDYLLRSAGNICNFLMVINAASNFFLYCALSDTYQRTLTTTFCRRERRWNERNDTLSTAA
SFRNSSVKQLRHENETQ
>GPA2/GPB2_receptor-NTE
RIVHTFLRRLPKIKDLGKIRPLHIVDFESNMIETLESNNIAITTEQLFLDYNRITTIQGWAFEGSQIGKLSFRGNRFLKDLSKDAF
SGLKSLRDLDLSETAIQYLPTSGLEELEVLVITNTPSLKTIPSIYDLTHLKKAYLTYFFHCCAFHYPERHDPARHQKYLEAMRKFC
SNSVERKARSIDDGGFREFEHDYIEAGIHNHTIFKQNYTEGKHYMWRFHFDNDEEHMIHEMFHNQSAELPNTKVHVTCG
NLTKRQVKVRCWPVADPLNPCEDMLSWVWLRISVWIVLTTALLGNTTVLLVLATTSHDRSLPRMLMSHLATADLSMAIYL
LLLAIMDLVSVDEYFNYAAAWQLGAGCQIAGFLTVFSSQLSLFTLSLLTIERWFAIRHALYSPLLNISRACRIMSVGWLYSIIM
AALPLLGISSYSATSICLPMDTHDYLSVGYILILLGGAALAFIVMCVCYIQIYMSLSYETRHSRSEGNVARKITVLVLTNLACWA
PVAFFSLTAVSGYPLINVTQSKILLVFIYPINSCANPYLYAILTKQFRRDFILLLSRHGLCTRYAQRYKVGYSRPTCNGTPATPLTS
DTCL
>Ion_transport_receptor
MTVLYSLIFLTGVIGNVSTCIVIARNRHMHTATNYYLFSLAISDLLLLISGLPQEMYQLWSKYPYVFGQALCVLLGLAAETSSN
ATVLTITAFTVERYVAICHPFFSHTVSKLSRAIKFVIAIWIMAILCATPQAMQFGLIYATDRRGNLIDPDEFNMCGLKEHYPYSF
EVSTFLFFFAPMSAITILYILIGVRLRKSTSRKAGQRLRDSRRSHGQAKSTTRVVKMLVVVVVAFFICWAPFQTQRLYALYFSP
SGNTSPRTILIYKFITYASGLLYYLSTTINPFLYNIMSLKFREAFKNTLTKCLNGREVLQSEGGGGSSGGVGAGGVFPRFNYSVL
SHRSVRHNNSTNNSLNNNNSCSSSNSYIDPVDLNRRPLVVSFRKKNTRTSSLEFSGNSPEKRELLQVPRMETFMPGAPKSPS
AQTRTAIIRTISTKYSPVRIKSGKQNGIQKIPRIEDIEIFQESQLKILQTKHAQHHQQQQQQHYHRHHHHHHNYHHQSHHK
EMENDDENHSCSVHSDDKQHTSCVDENDEDHVNYVKKSFTVPCLMMN
>Kinin_receptor1
MNCSFLEDELGPLPPSANCSWLLHNQSVYFYEESLYEVPAGVIVLLSVFYGTISVVAVGGNFLVMWIVATSRRMQNVTNCF
IANLALADIVIGLFAIPFQFQAALLQRWNLPNFMCPFCPFVQVLSVNVSVFTLTAIAVDRHRAVLNPLSAPPSKLRAKALLGAI
WILAAILATPMAVALNVTYVEENDHVGHVYTKPFCINTKLSNNHMMAYRMILVSVQYLTPLCVISYAYAKMALRLWGSRA
PGNAQHSRDANLMRNKKKVIIYYVIFNLLCI
>Kinin_receptor2
MMNLSNGSWPDEEEETLYDPPVSLVVFLSLCYGSISIAAVVGNGLVIWVILTSRRMRNVTNYYIANLALADIVIGLFAIPFEF
QAALLQRWVLPHFLCPFCPFIKVLSISVSVLTLSAIALDRYRAIIHPLTARVSRFQFRLVVSIIWIASASMAAPMAYALRVIPHPY
IKNIENESIYFCANEKLSSEAMQWYHSVLVLLQYFIPLTVIIFAYARMGLTLWGATAPGNAQSERDANIMRNKKKVIKMLVIV
VVLFALCWLPLQTYNVLQNITAINEYKYINILWFSFDWLAMSNSCYNPFIYAIYNEKFKREFQVRLQTPCLKKRHRSAPLTREL
SGFESSRSEWKRKSTMKNGMLINPATITLH
>Long_Neuropeptide_F_receptor_1_IsoformA-AKO62910
MVCRLVGTFRNKIELCVIMELNDTFNFSLNEVYRILIEHKNDDHNVDPVAEAILIALYALLIVVGILANLIVSFVVARRPQMHT
ARNLYIVNLTVSDMTLCLVCMPFTLVNILRRAWTLGIVLCKLVPALQGTNIMVSIGTITVIALDRYFTIVRGQDSATTRRRVIIS
IALVWFFSFLATLPVVFFQIVEPFKFEAVILYETCIERWPSQELKVAYAVCVLMIQAVIPALVVGCIHAKIASYLNAHAKTQRDS
KRAQRELQRNKRTTLLLSAVAVLFAVSWLPLGLFSLMADLLYPPGSETHISSQSLYITLAACHLLAMSSAISNPVVYGWLNSNI
RRELVQLLPSRCTSRQQQSQQQTTNAPSPTIMLCQNGQNIPHQQPATTYTAL
>Long_Neuropeptide_F_receptor_1_IsoformB-AKO62911
MVCRLVGTFRNKIELCVIMELNDTFNFSLNEVYRILIEHKNDDHNVDPVAEAILIALYALLIVVGILANLIVSFVVARRPQMHT
ARNLYIVNLTVSDMTLCLVCMPFTLVNILRRAWTLGIVLCKLVPALQGTNIMVSIGTITVIALDRYFTIVRGQDSATTRRRVIIS
IALVWFFSFLATLPVVFFQIVEPFKFEAVILYETCIERWPSQELKVAYAVCVLMIQAVIPALVVGCIHAKIASYLNAHAKTQRDS
KRAQRELQRNKRTTLLLSAVAVLFAVSWLPLGLFSLMADLLYPPGSETHISSQSLYITLAACHLLAMSSAISNPVVYGWLNSNI
RRELVQLLPSRCTSRQQQSQQQTTNAPSPTIMLCQNGQSIPHQQPATTYTAL

>Long_Neuropeptide_F_receptor_2-NTE
MLTGGLPTMDHEYNLLPPDASGLLSSVPGLHLTGGNTTTEFNQRSLDPIFNFSYHEAIEILREHQRTEKVLVPDTEIVIIIVYSV
LMTSGVVSNALVCFVVARQCARKHHQAGPSPRNMYIVNLAVADLALCLVCMPFTLVSLLKRRWTLGLVLCKLVPAVQGA
NIMVSAGTITAIALDR
>MIP_receptor_IsoformA-KF958188
MVEEMEEIWPTLGGPYFQCVNCTGGGVYDVSILNTTATSITMPNSSIWHNATEEEESDYLNVTKEFPIDYAVPMYGYAVPF
LLLITIVANTLIVVVLSKRHMRTPTNAVLMAMALSDMFTLLFPSPWLFYMFTFGNHYKPLSPVSACFAWDIMHEVIPSLFHT
ASIWLTLALAVQRYIYVCHAPVARTWCTMPRVLKCIGLIICLAILHQSTRFLDRVYIPVTITWRGQPQVPVCKVDMAPWVQ
WLTPTVYFTTYFAFRVIFVHMVPCILLVGLNLLLFRALRRAQRKRDKLFKENRKSECKRLRDSNCTTMMLIVVVTVFLITEIPLA
VLTVLHVISSSIKEILDYSVANVLVLFTNFFIIISYPINFAIYCGMSRQFRETFKELFIRGAVQVTRRNGGGSSKYSLVNGPRTCTN
ETVL
>MIP_receptor_IsoformB-KF958189
MVEEMEEIWPTLGGPYFQCVNCTGGGVYDVSILNTTATSITMPNSSIWHNATEEEESDYLNVTKEFPIDYAVPMYGYAVPF
LLLITIVANTLIVVVLSKRHMRTPTNAVLMAMALSDMFTLLFPSPWLFYMFTFGNHYKPLSPVSACFAWDIMHEVIPSLFHT
ASIWLTLALAVQRYIYVCHAPVARTWCTMPRVLKCIGLIICLAILHQSTRFLDRVYIPVTITWRGQPQVPVCKVLHVISSSIKEIL
DYSVANVLVLFTNFFIIISYPINFAIYCGMSRQFRETFKELFIRGAVQVTRRNGGGSSKYSLVNGPRTCTNETVL
>Myosuppressin_receptor-AGT02812
MDIMNSTALPEHQPYCGEGFDTFRNVYREVHGYLSLWVCLFGSVANLLNIVVLTRREMTSPTNAILTGLAVADLLVMLEYIP
FVWLMYLSPSSRSDRYTYGWSFFVLFHSNFTQVCHTISIWLTVTLAVWRYIAVAYPQRNREWCGMQRTIVAIFLGYVICPIL
CIPLYLAFNIQSKSPPLDDNTWTQNQTLYIVHFSELGSANHNLLVDLNFWVYSVVIKIIPCIALTVLSLRLICALMDAKRRREAL
TSGSKKTPRNLENERQTDRTTKMLLAVLLLFLITEFPQGILGMMTIILGRGFFKDCYGKLGDVMDILALINSAINFMIYCAMS
RQFRNTFSLLFRPRWLPVPQVENGVNHNRTTTMVTQV
>Natalisin_receptor
MRTTTNYFLVNLSISDLLMSLFNCIFNFTYMLDSHWPFGAIYCTINTSNFVANDSVAASVFTLVAITLDRYMAIVRPLKHRMS
RRKARIALLIIWAASSLLAIPCLLYSTTKSRTINGQTSTVCYMMWPDGHYPKSMSEHVYNLIFLVVTYLGPVVAMAICYTLMG
RELWGSKSIGEQTQRQLDNIKSKRKVVRMFITVISIFTICWLPYHGYFIYVFHHESVAVSSYVPHLYLSFYWLAMSNAMVNPI
IYYWMNNRFRVYFRQIICLCCCVRPSMHPELQSTPNNRLVRSELLLRSKSCKPQGQFSEIEFSGGGISGSLSGGSSTTMHIET
GRFRSDKSRYKNKPFVDSRYD
>Pyrokinin-1_receptor_variant_A-AFO73269
MDSGIESFTNETVSVRFPKRDPLYIVIPITILYSTIFVTGIVGNVSTCVVIARNRHMHTATNYYLFSLAVSDLLLLITGLPQEMYYI
WSRYPYVFGEAFCLLRGLAAETSANATVLTITAFTIERYVAICHPFLAHTMSKLSRAVKFILAIWVVALAFAIPQALQFGLIYVN
EPLQIQCNLKKILIVHSFEVSTLLFFIAPMTLITILYALIGLRLRRSALLTRNSGSFGHGDSGRKGTNSCRHHSSQRVLKMLVAVV
VAFFICWAPFHTQRLVAIYITNMDGLGQHVYSVVTYISGILYYVSTTINPILYHIMSLKFREAFKTALCCRTHTRSVVGRRGSR
VHT
>Pyrokinin-1_receptor_variant_B-AFO7320
MDSGIESFTNETVSVRFPKRDPLYIVIPITILYSTIFVTGIVGNVSTCVVIARNRHMHTATNYYLFSLAVSDLLLLITGLPQEMYYI
WSRYPYVFGEAFCLLRGLAAETSANATVLTITAFTIERYVAICHPFLAHTMSKLSRAVKFILAIWVVALAFAIPQALQFGLIYVN
EPLQIQCNLKKILIVHSFEVSTLLFFIAPMTLITILYALIGLRLRRSALLTRNSGSFGHGDSGRKGTNSCRHHSSQRVLKMLVAVV
VAFFICWAPFHTQRLVAIYITNMDGLGQHVYSVVTYISGILYYVSTTINPILYHIMSLKFREAFKDTYSKCCWPKRKSRPYLILS
RGRAEPESGRSLTDSSAQHSLTNQQTNRPNDSFSQPETQPIMVMECKSTQLPPLCTRCQQHPQKPLVDGDISNSSLRDVD
KAALEDELTAYMNELARRQLS
>Pyrokinin-1_receptor_variant_C-AFO73271
MDSGIESFTNETVSVRFPKRDPLYIVIPITILYSTIFVTGIVGNVSTCVVIARNRHMHTATNYYLFSLAVSDLLLLITGLPQEMYYI
WSRYPYVFGEAFCLLRGLAAETSANATVLTITAFTIERYVAICHPFLAHTMSKLSRAVKFILAIWVVALAFAIPQALQFGLIYVN
EPLQIQCNLKKILIVHSFEVSTLLFFIAPMTLITILYALIGLRLRRSALLTRNSGSFGHGDSGRKGTNSCRHHSSQRVLKMLVAVV
VAFFICWAPFHTQRLVAIYITNMDGLGQHVYSVVTYISGILYYVSTTINPILYHIMSLKFREAFKDTYSKCCWPKRKSRPYLILS
RGRAEPESGRSLTDSSAQHSLTNQQTNRPNDSFSQPETQPIMVMECKSTQLPPLCTRCQQHPQKPLIGMNE
>Pyrokinin_receptor-NTE/CTE
KHSSLLVNAYTILTHISGVLYYVSTTVNPVLYNIMSLKFRDAFKCTLSQMCGGRGRSGKPRWTYSMLSRGGHQTTVGSGPH
RVPSEVRRLRLANKPATISNSSLQDVDEAEYTGPELANYMGQLNSR
>Pyroglutamylate_RFamide_peptide_receptor
MDQPIKSSGITKPTKMELQENLTDDYDYDVQESFNTYIWEELVPTLVVYILTLVIGVTGNFLIIFTIARYRRMKSITNVFLASLA
SADLLLILVCIPVKLAKLFSFTWTMGVFLCKMMHYMQSVSAICSVFTLTAMSVERYYAIVHPMKAKYVCTISQARKIIFTTWV
ASFFLAVPILFVQVQMPVGGRIKAYWCVRDWDWVVAWRCHEVYMLVLVLLLPASVMTVTYSAICREIFRVMQRRFHMT

SGKATMNCESFPLSTKEKNPRTFKPKIRSEEENNTVKQMSNYKSERQVIKMLVAVVIVFILCWAPVLVDNVLTSYDILPHIRE
GTLKQLATYFQLLAYFNSCVNPLVYGFMSKNFRESFSKALCCRRKVPRRQLSVSHTRTTSLINYVN
>RYamideR-INT
MTNEEDQSTFEWTFNNETYGKQNDSFNETIDCDKIGPGGLLSSGYFQSAVYILYILIFVTALSGNGLVCYVVQSSPRMRTVT
NYFIGNLAVGDILMALFCVPFSCVPSLLQHWPFGLHMCRLVSYTQGVSVLVVSAYTLVAISIDRYIAILWPLKPRMSKKMAK
LTILTVWTVALTTALPIAAVSALGQPSVWHVQCHRCLCVELWRSESAKHIYSYTLLSVQYMLPLAVLLYTYTSIAVVVWGKST
PGEAETSRDMRMARSKRKLIWEANPALSDWHGLPYIYFALHWLAMSHSCYNPLIYCWLNARFRTAFCVALKKLSCKRKPP
PDQLLHRINTCTTYISMHRAKHRSKTSFLEQQI
>Short_neuropeptide_F_receptor
MNNTTIEEDLSMIVDCIVTQYNISQYKGQWPNCTEIGFRKDIIDDKIVQAIFCLLYTSIFVLGLFGNILVCYVVGRNRAMHTVT
NCFITNLALSDILLCTLAVPFTPLYSFLGRWIFGNALCHLVVYAQSTSVYISTLTLTSIAVDRFFVIIYPFKPRMRLSTCLAVIFFIW
TFSLIATIPFGLFMDHKSIAGRFYCEEKWPSENFRQVFGGMTSTIQFVLPFIVVTFCYVRVSVKLNDRARSKPGAKTSRKEEV
DRERKKRTNRMLIAMVTIFGVSWLPINLINVINDLYMHTSSWTYYNLFFFLSHAVAMSSTCYNPFLYAWLNDNFRKEFKQV
LPCFGAQTGPQASGRLGNWRSERTCNGNETCQETLLPTSVVISTTKPSPPLLNTSSNEGRGGRTDGGGARLRSTDSVEVVL
VAYTAAEDAVHIDKLDKGTIEAQKNRPVQVV
>Sulfakin_receptor_1
MIHIISFILLQIKMLPTESWWEAGKVQIPTYSIIFLLGLVGNILVILVLVKNKGMRTVTNVFLLNLAVSDILLGVLCMPFTLVGSL
LKDFVFGHFMCRLIPYMQGFFSAVSVSVAVWTLVAISLERYFAICRPLKSRRWQTQFHAYKMIAIVWAMSLVWNSPILFVS
RLLAMGGKGEGRHKCREVWPGRRSEGAYIIFLDIVLLMIPLLIMSLAYSLIVLKLWKGLQRELKHSNSCLKSFSLLLQVIRMLF
VVVAEFFICWAPLHVLNTWYQFRPDLVHQYVGSTGVSLVQLLAYISSCCNPITYCFMNYRFRQAFISLF
>Sulfakin_receptor_2-NTE/CTE
MTGGRIFHLTGNIINFDFAVADILLGVFCMPFTLIGQLLRNFVFGRIMCKLIPYFQAVSVSVAVWTLVAISLERYFAICRPLKSR
RWQTQFHAYKMIAIVWAMSLVWNSPILFVSRLLAMGGKGTRLFTEIVRAVAIK
>SIFamide_receptor
MVSLRLPDSEEYIEARGRALETGLAPTRASNTSSRRLFVNSLLVDFVMDTLVTNTTPSASSSPSSPAAGAVADLPDSTSNQTY
QHLFYRHSIAMTIVFCVAYLIVFIVGLIGNCFVIMVVYRSPRMRNVTNFFIVNLAVADILVIVFCLPATLMSNIFVPWVLGWW
MCKTVPYVQGVSVAASVYSLIAVTLDRFLAIWWPLKCQITTRRARLMILVIWVVALTTTIPWALFFDLVVIFTDNPEVKVCSE
VWPEYLNGSLYFLIANLLFCYILPMILISMCYVLIWIKVCKRHIPSDSKDAQMERMQQKSKVKVVKMLVVVVILFVLSWLPLY
LIFARIKLGGEISGWEEDMLPMATPVAQWLGASNSCINPILYAFFNNKYRRGFAAILKSRKCCGTLRYYDTVVRANSSSTSLR
KSSYYVTNNNNNNNSSTRRQLSQDTNVSYISNNTGV
>Tachykinin_86C
MPLELQLAWAGLFTGMVFVAVAGNLIVIWSVFAHRRMRTVTNYFLVNLSLSDLLMSTFNCLFNFVYMVNNDWTFGSGY
CTVNNYLANVSVAASVFTLTCITVDRYLAIITPLKPRMSKANAQLAILAIWAASLLLATPCLLYSTTITHKPSGQTACTLLWPD
GQPLVSTLDYIYNLVLFLVTYVVPMTAMVCCYSAMGRELWGSRSIGELTQRQLDSIKSKRKVVGMLLVIVLVFGLCWLPYH
GYFLCAHHWPALVYTRHVQHVYLAFYWLAMSNAMLNPLIYYSMNHRFREYFRKAVCEWRCQLWSRQ
>Tachykinin receptor 99D-like
MAYTENSTLGWNELTQNITVNETYDDEDGNQFILPWWRQLIWTFLFGGMVIVATGGNLIVIWIVLAHKRMRTVTNYFLV
NLAIADAMVSSLNVTFNYTYMVNSDWPFGTLYCKISQFVAVLSICASVFTLMAISVDXXMAIMHPLRPRMGRRMTLCIAVS
IWIVGSFFSLPMLIFFTTFVQEFPNGDNRVICYAEWPDGSTNESRQEYLYNVLFMVMTYFIPIASMCFTYVRVGIELWGSQSI
GECTQRQLENIKSKRRVVKMMMVVVSIFAVCWLPFHIYFIITSHMPEITKLPYIQDLYLTIYWLAMSNSMYNPIIYCWMNM
RFRRGFKQFFSWCPYVHVPPEGLTRREAVTSRYNYSCSGSPEAHYRIVRNGKRLLLHK
>Orphan_receptor_3-NTE
ETYTCPTTHFKCNNHYCIPIDLLCNFEDDCGDKSDESKDCNHRQCWNLEFRCENGECIRPGFVCDGRKDCKDGSDEALCAE
DDFVMCRDGSRVHRSYWCDGWPDCPGNHADEWNCEVCDGPNDYKCPNGRCIKKANICDSQCDCAPHNGSLECADEM
NCSKYYRSVHGKVDRCIASKYICDGSNDCHNGKYLSDEYGCQPSENQYSESTFRCLDNRTLPESLLCDYKNDCLDGDDENLC
RALYQCDETMFTCNNSQCIDKNGRCNVTYECLDKSDELGCLDVPCPEGMVKCTYGGQCIPEKLLCDYFIDCPDESDEKNCP
VTECNKLQFQCDNGQCVSIEHHCFISGNQRDGCADNSHLKNCKNFTCMRDHFKCRLGPCLNQSLLCNKKIDCQHTWEDE
DNCTFTCSEKYPECPCKDIYINCTALGLESVPLDTEGEITWFHLGSNKLNASLTNETFSSLDRLLYLDLSNNSITGLPPMMFSNL
WRLTVLNLQNNRIHTLVSSSFYGLASLKGLHLQGNGIRVVRMLAFYGLSSLRNLDLHDQNINLIEPDAFLGLRSLVGLDLSQN
KIEYISDSTFRGMPHLLYLDISNNYIDVIDANAFRMATTLEKLVTDEFRFCCLARHVKSCLPPRDEFSSCEDLMSNMVLRICV
WALAVIATVGNILVIACRARYKHCNQVHSFLITNLALGDLLMGSYLLLIAVVDWHYRGVYFIHDSSWRSSQLCSFAGFISTFS
SELSVFTLTVITLDRFLVIIFPFRVRRLEMTRTRRLMAFGWIVAISISAVPLIHIDYFKNFYGRSGVCLALHITPDKPNGWEYSVF
VFLFLNLVSFTIIAVGYLWMFLVARTTQHAVNKDRRTSESAMAWRMTLLVATDAACWVPIIILGIVSLAGYTVPPQVFAWV
AVFVLPLNAAVNPVLYTLSTAPFLTPARHGFLTFRRSCKMSLSQDQRRTYTSGLNHYAGKSS

>Orphan_receptor_4-NTE
QVRNATAVFIINLSVSDLMSCCFNLPLAASTFWRRSWRHGLLLCRLFPLLRYGLLAVSLFTVLAITINRYVMIGHPTIYPKLYRK
QYLGLMVAATWICGFGALIATWLGRWGKFGLDPKIGSCSILPDSSGRSPKEFLFLVAFVIPCICIVVCYARIFYIVRKTALKSRV
AGRSAASVTSGGTTLTRSGYYGKVKLVRKGNTSSTDDSAFATSSTAQSFSTEKSSTILDNGEMGGNETVIKMVNLAPSPHLL
APQRRSKICAEASSSSGIEEGLREDDEVVSTRSDSPISACSSSPPPAHYSVKVQKIKKRSERVNSTLSHMASVFRRTSHARGVL
SPSRRQSCAPPQPGKMTAKDKKLLKMILVIFASFVTCYLPITLSRHTEI
>Orphan_receptor_5-NTE
MGKGLKEIHKYKGDIPLHVSTLTLLLIAWDRHRFLKDPMKPRIPAFVCATGSWLTAICLVLPYPVYTTYMDLGFSNFSPPMLY
LYFWRSYGQKPDWGFMRTTGFDNDYENYIKNCEK
>PDF_receptor-NTE
LEIGVLPGEYDVNQMGTTVRLDLSNREFCKTKHDNVTFEDTPFCPAVWDQVLCWPPTKGGLTSTQSCPNHHGVDPSRLV
SKRCLENGKWEGIGETGWTNYTTCYSPDLIQLFKKLFTGPYSHDAIKYQIAERTRTLEIYGFSISLAALFISLYIFSHFRVLKNNR
TKIHKNLFAAMVAQAVIRLTLYVDQAIIRARKVQGIDNTPILCEASYVLLEYARTAMFMWMFIEGLYLHNVVSVRVFQETFH
YKLYTSLGWGAPVIMTSAWAVTLAVQMKTECWWGYNLSIYFWILEGPRFAVVILNFLFLLNIIRVLVVKLRQSHTNEIEQVR
KAVRAAVVLLPLLGITNLANMLGAPLDRQVWEFAAWSYATHFLTSFQGFFVAALYCFLNGEVSIQTI
>Parathyroid_hormone like_receptor
MDIKPIPLSKDEQDKLLDSLRSECILKNDTILYPDGCPTIWDGILCWPNTPSNTLASLPCPEYFKGFPSHRNATKYCQLNGTW
YFDYSLNQTWTDYTACMENDPEEDLLPPKSNIMYLEKYLPTLKIISQIGYSVSLISLIFAFILLASFKKLRCPRNVLHMHLFVSFI
MRAGIKLLRDTVFFSGLGFHYEIRALIEYASNYSYSNDLPDNWICKFVTGLWQYCIVANYSWILMEGLYLHNLIFLALFSDTSA
ITVYVLLGWGLPLLFVVPWVIVRAVYENTLCWTVNNNPYYFWIIRAPIASSVVLNFILFINIVRVLMLKLTVSISEEKKRYRRW
AKSTLLLVPLFGVHYALLIGMSSSMNKNQYVEMVWLFCEQLFASFQGFVIAVLYCFMNGEVRTEISKLSWNKRSPQYFQN
QYSVTDRPISSFFRLAKRQKRGSTAESCMTSFMSSVPTDLNVRRPTLPKTVTNSHLVNEIKADSCLLNKGPTICSGMQEIN
>Long_wave_sensitive_opsin1
MAQPIGPSFAAYQWGQSANPSANRSVVDMVPPEMLSMVDAHWYQFPPLNPLWHGILGFVIGVLGIISIVGNGMVIFIFS
STKTLRTPSNLLVVNLAFSDFLMMFTMSPPMVINCYNETWVLGPLMCELYGMLGSLFGCASIWTMTMIALDRYNVIVKGI
SAKPMTNKTAMLRILLVWAFSIMWTVFPFFGWNRYVPEGNMTACGTDYLTKNWVSRSYILVYSVFVYFLPLFTIIYSYFFIL
QAVSAHEKQMREQAKKMNVASLRSAEAANTSAEAKLAKVALMTISLWFMAWTPYLVINYSGIFETISISPLFTIWGSLFAK
ANAVYNPIVYAISHPKYKQALEKKFPSLSCASPQDDTTSVATGVTTSTDDKAPSA
>UV_opsin_-NTE
LLNVLFVINCHRASTSGNIRTLGWNLSPEDLKHIPEHWLSYPEPEPILNYALGVLYIFFMLIALIGNGLVIWIFSTAKTLRTPSNI
FVVNLAICDFLMMSKTPIFIYNSFKLGYALGHRACQIFALLGSFSGIGASATNAVIAYDRYRVIATPFAPKLSRTKAVLYLALVW
AYVTPWALLPLFEQWSRFVPEGFLTSCTFDYLTPTSEIRNFVTVMFFICYVFPMSLIIYFYSQIVSHVIIHEHNLREQAKKMNV
ESLRSNANMHTQSAEIRIAKAAITICFLFVASWTPYAVLALIGAYGNQDLLTPAVTMIPACACKAVACVDPYVYAISHPRYRQ
ELSKKFPWLDIKEAPAPSSVDANSTATEMTLPTQTS
>COpsin2_Pteropsin
MELMLMPSAGFLAASIILFLIGFLGFFGNLIVIIIMCRDKNLWTPVNFILFNVIVSDFSVAALGNPFTLASAIAKRWFFGQSMC
VAYGFFMALLGITSINSLTVLALERYLIVSQPVSHGSLSRPTALTIVGSIWLYSFVITAPPLVGWGEYGLEAANISCSINWETRS
HSSTSYILFLFTFGFFIPIIVISYSYMNIILTMKKSTMNAGRVNKAESRVTWMIFVMIFAFFLAWTPYAILALMIAFFDSNVSPAI
ATIPAIFAKTSICYNPFIYAGLNTQFRQSWRRVLGGKREDSTTMATATSFGLNSKRYKEVSCVIDVKGKDKIKLSALNKSTATE
TAI
>Rh7-NTE
LCSANISTTIYPKYFHLYPIEQWKMHRFFTEEYLKLVNTHWFEYPPPNKQIHYIFAAVYFLVMLVGVSGNLLVIFMILSFRTLRT
SSNILILNLAVSDFLMVAKMPVFIYNSFYFGPVLGEMGCHFYGFIGGLSGTASILTLAAIAMDRYLGIAHPLNFNQGRAKKRTI
VWITFIWVVYSITFASIPLSHIGVKTYVPEGFLTSCSFDYLSTDIQNRCFIFIYFVAAWCLPLLVIITSYVGICREVLRVSLIRKGQER
EQRKREAKLSAILALATFLWFLSWTPYAAVALLGIFGYKNHITQLASMIPALFCKTAACVNPFIYGLNHPRLRQQLLKLCCKKR
YNLEKTHFSRSWRNTSCSFK
>Potential Neuroparsin_receptor
MDITSSTKRMGYLSKLFWLLLLLVQQAFCLTPCLQNEEVNPKRYLMSRFLVQLKLGVSDRILHQLTTQVYQILLTEVLGYASV
GIVQYNSSGNVSEQLRRLWQEDENVPPEYVVDLEVMIPAHENPYEMLGVKDCGNLGPPGRYGWFIPKKLLPHHTQNRKE
HIDHWKIFHDLETAMLFSLSEDDWKYVVQNTLVENAVVKKYHCEKSFCENGLYTPDRCKEGKKCAVLITENPDMTEFVASD
IERLSLYVRVAWVGSAMEVVNYLTDKYYATDPKNRGLIVFLSYTPSVLTLTVDHLSVSFPPCDLFDNTTCSYANQRHVKVAW
PKLDEVAKFALESLQKMEFRSEDLKEMVYDYMTEYKNNNERPVNITQVACAWMRRHHKILTLNTTNWETWLTVFDNNQ
TIYIGGIFPMSGLHAARGAVAGAIMAVNAVNKYSVIKNYSLAMKLDNGQCKADVVMKTFVEYMLFGEHHSLAGVLGPAC
SDTLEPLAGISKHFKTVVISYSAEGSSFSDRTKYPYFFRTIGENKQFKYVYLELFKELNWKRVASLTEDGTKYTEYISLTQDLLQQ

NDITFVANRKFPQDWGKDSAMRQYLEEFKDIDARIIIADVNDEAARVVMCEAYHMKMTAKQGYVWFLPLWLPTDWYNT
TIFNETRGLPLTCDTNQMIEAINGHLAITHSFFAPDDNIMQENITVRQWRDNYEKKCQVGLLTPSNYAGYAYDAVWTYAY
ALNTLLKENQSSVSSLHDDKTVTRMVQIIQNTNFNGVSGHIYFGGGPSRYSIVNVVQWYNMKTKIVGTYYPNIPNVTVSPD
KFVLRKEEIVWLNEGGIIPSDGGSKCVIDSFASLLNVGCQAAIIVANLLGIALLILFLIVGFLIIKHRYDKKVQLTQKYMKSLGIDL
LSVSTFGGLDSWEIPKDRVVINRKLGEGAFGTVYGGEAYFNEKGWVAVAVKTLKVGSTTEEKLDFLSEAEVMKRFDHKNIV
QLLGVCTKNEPVYTIMEFMLYGDLKTFLLARRHLVNEKADDNDEISNKKLTNMALDVARALSYLAQLKYVHRDVASRNCLV
NVSRVVKLADFGMTRLMFENDYYRFNRKGMLPVRWMAPESLALGVFTPASDVWSFGVLLYEIITFGSFPFQGLSNNQVLE
HVKAGNTLTVPAGIKPQLDSLIKSCWNIDHKKRPQASEIVEFLANNPRLISPCLDVPLSSVQLEDTGQLEISITDPDKPRKFSM
TLRQRSPSSCGGTSAAGGGCGTLIGGGCDSKQWNEVAQPLLPSPNRYKPTRTSSISEEAATSFEASDSLL
>Eclosion_hormone_receptor-NTE
LTISGAISLAVNEINAELAANGSEHRFRFIVAETMGDELISIRQTAILWTKNVSVYIGPQETCVHEARLAASFNIPMISYYCTNH
ETSNKLLFPTFARTRPPDTQISKSVAAVLLAFNWTQVTFLYRNSSDLTTIAETIKQVLNSVGIIVTNSHTWKDFYHHGYTENPF
YELIQRTYKQTRIYVILGEDDEHLGLMVALEESNLLDNGEYWVVGVDIVQYDENDPTKYLRGLLQKKNEPIYRNAFRSYLGV
APRPPINITNFAVQVNKYMEESPFNFSNPIGHLGGVKVIKAEAVYLYDAVRLYASVVLELMANGLDHKNGTLIINRLLGKHY
HSAMGYMVYMDRNGDAEGNYTLLATQEVRPGDFGLYPVGGFSYTRSSRWLPKTIQFCSFQELHLTKNIDWVGGRVPVA
VPPCGFSGEKCFSITMEIVGGITGAGLAVLILSALIVYRSWKYEQELDSLIWRLDFRDIKLNEDFFSNEQKANRVIQNIRCSSGK
TMTNPLIRTSQVSLSSNPDADFRYSSILTQVGIYKGRVLAVKKINKKSVDISRKMKKELKLVRDLRHDNLNSFIGACIDPPNVCI
VTEYCTRGSLKDILENGGVKLNNMFIASLVGDILRGLSFLHDSQLRYHGNLKSSNCLVDSRWVVKLADFGLTELKKGSDDYQ
QAWSLKHVFCSGLLYRAPELLRLTLSSYGVGSQKGDIYSFGIILYELHGRKGPFGDGLGLTPKQIIHRIICPPRNSSQAFRPPLN
QLETNFDYVKDCLIECWQENPDDRPDIKGVRAILRPMRKGMKPNIFDNMIAMMETYANNLEALVDERTDQLVEEKKKTE
ALLYEMLPRCVAEQLKRGHKVEAESFDCVTIFFSDIVGFTAMSANSTPLEVVDFLNDLYTCFDSIIVNYDVYKVETIGDAYMV
VSGLPVRNQDQHAAEIASLSLDLLQAILSFKIRHRPQDKLQLRIGIHSGPVCAGVVGQKMPRYCLFGDTVNTASRMESTGLA
QKIHCSAETKNLLDRLGGYNILERGYIDIKGKGEILTYFLESEDITHRTQRRSVREKRRHSKETVYHRAILRSSLKATCSLSRAASF
ESSKRLRFSNKNVEYNNSYEIAKLESVVDSSPSKCLTVDFCDHLSVSCPCIENVQNRVNGEKLIEANSDPLLTVMSDQICT
>NPLP_receptor-NTE
MMCEEIAAFFGPEGSCHVEAIVAQARNIPMISYKCSDYRASEVPTFARTEPPNTQVTKSVISLLRHYQWSKFSIISEETWRPV
ARSLEEEAKKKVNNMTVNHKKIIMDRHKCCELQLPCCQTGIWYQLLQETKNGTRIYVFLGTPVSLIDMMTTMQALQLFEK
GEYIVIHVDMMTYTPREATKYLWKPKTFNEPSSCLDHPGFEKRARSLLVVVSTAPTTNYENFTERVRLYNTKEPFFFPTPTVF
KNVSYVKFVSIYAAYLYDSVWLYARALHELIYGNTSNGETRNITENVIKEVVKNGTKIIETIVRTKTYKSVTGFKIKLDTNGDSE
GNFSVLAYKPHNITEDNFTCSHHLVPIGQFQQGYNDQHGGYPELKVKWTIDWAGNDKPEDEPICGFDYSRCAKEDSQGSV
AAAIVLAIALFCAMVSTLSIYRKWRIEQEIEGLLWKIDPEDLLDPQDLMSSPSKLSLASATSFESRCVPQLFAWTSKYRGIVVRL
KEIKFSKKKDISRDIMKEMRLMRSFIHDNVNSFIGAVVEPMRIVIVTEYCAKGSLYDIVENEDIKLDKMFVASLVHDLIKGMM
FLHGSAIGVHGNLKSANCVVSSRWVLQVADFGLNELRHCPENESIGEHQPYRNLLWTAPELLRDEAHMHRGTQKGDVYA
FAIILHEIIVRKGPFGGCGKDEPGEIVRLVRCPHSDEPFRPNVDLVRDSEVGSDQVISVMVDSWAEDPEMRPDFGTIRSRLKS
LRGGKQRNIMDQMMEMMEKYANNLEELVNQRTLEVYEEKRKTEDLLHRMLPAPVAKRLTSGFGVEPESFDLVTIYFSDIV
GFTAMSAESTPLQVVNFLNDLYTLFDRIIKGYDVYKVETIGDAYMVVSGLPLRNGDNHAGAIASMSLDLLNAVKNYKIAHRP
DETLKLRIGIHTGPVVAGVVGLTMPRYCLFGDTVNTASRMESNGEPLKIHISQQCCAALQKLGGYQVQPRGIINMKGKKPV
QTYWLVGATEKAVKPTEVDLTELPPPLFCRPRKSPKIMGLVDSRRQSSVLRTCAQPVVLRPLVRGAQSLDPLPSTAIPRSPVK
RSCHSLQEKGVVCSSICGNGALVSEPLLQDNKRWHSLETVSAEPCTKKSLATRSSLRSWLFGLFNNTAFNASDASLRKVGYQ
DLQPERESIV
>Insulin_receptor
MIVYWVLLYCTGITFLILGMQPAYTKICPSMDIRNTVSALNKLAGCRVIDGYFSFVLIDYADESEYDNMTFPELREITSFLMVN
KVSGLRSLGRLFPNLSIIRGERLFLDYALVITNMQNLLEIALTSVTILRGSVAIAWNKKLCYAETIDWDQIAPGGDHFLVGNSPI
TDPECPGCACKNNLCWSRNQCQEIRKWVTPDGEPCDDECVGGCTGLGPYNCKACRRFDHDGGCMKSCPSNRYAFENHY
CVTEEECRNPNSTYRIAIERKKGQDDVVWSKPQEWFVWNGTCIQDCPTGLEKTTMSSCERCKDGKCKKECYGSVVDSLEK
AERLRKCTHILGSLEIQIKSGQQSVVAAELEDSLGMIEEIQGQLKITRSFPLVSLDFFKNLRIIQGDRHFYFNSNYSLFIKDNQNL
MTIWNWDKRPAGRNFTINMGRPLFNDNPKLCIKHIRELTTIAGFKDVKDTEVTKQNGVKFACNLVELNISAHLTFSQSIVIHI
HKPDFNNTSVVFTRYIAYYMEEPYGNLTTAIPSDDCEENAWKLNDVAISEEDKSMSNLKMYHHTITKLQPDTQYAIFVKTYT
VDSTGGQSPVLYVRTLPSRPSMPLYLLAHSNSSSEIVVTWEPPEKPNGKLSHYIVKATMHGDDPTYLEARDYCKYPIKKEETT
TPAPRLVMLKEDSDDKSTDDCVDKKPEKKRPGDVCESIDPHLPPKLYDAPTCEKYMYTLVDSTRLTPTAEEHVMTEPSLYER
NIELAEEPADLIRRNIKDEDEDDRLVEDLEQFNSDGTYASFTARYPHNVTLVTLSNLKHYTAYTVEVIACRERHPRDSATTKRC
SLNAFTTLRTLPDPKADNIEGGIKESVVNRTVTITWTPPLANGVIVAYMLERVREGSGADSKLMVECIPVAMARGSFELRGL
ELGSYRIRLRALSLAGAGEFTEPEHFSISEYSSTNIIIITFFIVITILLIGIVAGFVYYHRRKMNLQEVLIASVNPEYFGLPTVDEEWE
LPRDRVRLIRELKRGNFGVVCEGILSPQGTTVAVKMSIDDEPSDRDAMQFLNEAVVMKQFTEAQHIVKLIGIVSRDRPFMV

VMEMMAKGDLKSYLRECRNGIPPSPAGMILMAAQIADGMAYLESAKFVHRDLAARNCMVSDKLIVKIGDFGMTRDIYET
DYYRKGNKGLLPIRWMAPESLNDGVFTSKSDAWSYGVVLW

=====================
Biogenic amine enzymes
=====================
>Tyrosine_3-monooxigenase (ple)
MMAVAAAQKNREMFAIKKSYSIENGYPARRRSLVDDARFESVVVKQTKQSVLEEARIKHNDKADTPSNQTNLKEEIKKSSE
SHEPKIPVEIDDENKENVCDETLQHQNGHSDAGLTEEEVILSNAASESKEAEQAIQRAALILKLKEGMGSLARTLKTIENFKGS
IVHLESRPSKDAGVQFDVLVKVDMSRQSLLLLIRSLRQSAALGGVDLLADNKISLKNPWFPRHARDLDNCNHLMTKYEPEL
DMAHPGFADKVYRERRRVIANIAFEYKYGDPIPYIEYTEVERATWEAVYNTVVELMPKHFCKEYKEKFAQMEEEGIFTPKRI
PQLEEVSNFLKKNTGFTLRPAAGLLTSRDFLASLAFRVFQSTQYVRHSTTPFHTPEPDCIHELLGHMPLLADPAFAQFSQEIGL
ASLGASDEEIEKLSTVYWFTVEFGMCKEHEQLKAYGAGLLSSYGELLHAISDKPEHRPFDPSTTALQPYQDQEYQPIYYVAES
FEDAKEKFRRWVSTMSRPFEVRYNPHTQEVEVLDSVDRLDNLVSHLNLEMQHLTTAINKLRATFG
> DOPA decarboxylase (Ddc)
MDHKSFKDFAKAMAEYIAEYLENIRNRPVLPTVEPGYLRPLIPSAAPESPESWQQIMGDLESVIMPGVTHWHSPRFHAYFP
TANSYPAIVADMLSDAIACIGFSWIASPACTELEVVMLDWMGKLIGLPKEFLACSGGKGGGVIQGTASEATLVALLGAKAKII
HQVKEQHPEWSDYEIVSKLVAYASKQAHSSVERAGLLGGVKFRLLPTDEKHRMRGSSLQEAIEKDKKDGLIPFYVVGTLGTT
SSCAFDVITEIAPICNKEEVWLHIDAAYAGSAFICPEYRYLMEGVELADSFNFNPHKWLLITFDCSTMWLKDPSWVVNAFN
VDPLYLKHEHQGAAPDYRHWQIPLGRRFRALKMWFVMRLYGAENLRAHIRKQIGLAHQFEQLVQSDKRFEIIAEVLMGLV
CFRLKGSNEKNEELLKMINGRGKIHLVPSKIKDTYFLRMAVCSRYSDPDDINFSWNEIKST
> Tyrosine decarboxylase 2 (Tdc2)
MDTMEFRRRGKEMVDYICEYMTSLSKRRVTPSVEPGYLRSLLPEQAPQQPESWDQIMADVENYIMPGVTHWQHPRFHA
YFPSGNSYPSILGDMLSDAIGCIGFSWAASPACTELETIVLDWLAGKAIGLPDEFLAFTEGSKGGGVIQTSASECVLVTMLAA
RAQAIKRLKQLHPFVEEGMLLSKLMAYCSKEAHSCVEKAAMICFVKLRILEPDDRCSLRGATLRQAMEEDEAMGLIPFFVST
TLGTTSCCSFDSLNEIGPVCRLFSSVWLHVDAAYAGNAFILPELKSLLDGIEYADSFNTNPNKWLLVNFDCSTMWVRDRIRL
TSALVVDPLYLQHGYSHSAIDYRHWGVPLSRRFRSLKLWFVLRSYGISGLQEYIRHHCRLAKFFESLVKSDNRFLVCNDVKLG
LVCFRLKGTDKLNEKLLSNINGSGKLHMVPANVNDRYTIRFCAVAQNASQKDVDYAWGVITDHATELLEVQQIEKEQVFEL
LERKRKETLAYKRSFFVRMVSDPKIYNPKIAKSLPGSRRHTSHVTDDEDDIESTPDNGVSAHTPIASWISWPLAFLFQDTGAD
ALPNSFVSFNRFRHLDTMVRLPAKKETGSGGSRRNSHSPSSSAPVLGSIDNTVNHQL
>Tryptophan_5-hydroxylase_1-CTE (Trh)
MSGSGKSLLGLWLYRSGEKWSLKHTDSEVRRSRVQPHIQNTGRNSVVFSLKNQIGGLARVLQVFQDMGVNVVHIESRPS
DRHDSEYEILVEVECDNRKMEQVVSLLRREVAAINLTTYESGSDLPPPTPLSATTSFDFEEMPWFPKRIQDLDKAQKVLMYG
SELDADHPGFKDPVYRKRRETFANIANSYKYGQPIPRVQYTEEEIKTWGTVFTELHKLYAKYACREYLENWPELVKYCGYRQ
DNIPQLQDLNLFLKRKTGFQLRPVAGYLSPRDFLAGLAFRVFHCTQYIRHSSDPYYTPEPDCCHELLGHMPLLANPSFAQFS
QELGLTSLGASDEDVEKLATVTISRLNMIYHGELRVYGAGLLSSIAELQHAIQATEKIKKFDPELTCHEECIITSYQNAYYYTDSF
EEAKEQMRAFAKCIQRPFGVRYNPYTQSVDVLSNAQKIAALVSELRGDLCIVSNALKQIDAREDTMENIAHMLQDGIDL

================================
Neuropeptide processing enzymes
================================

>Signal_peptidase
MAEVNKILSDIKMNLTENVTDTLHRTPATPEGMAVAYISIIIMAILPIFFGSMKSVKYHKEMIQFQKESGEMLETMSHKDAA
MFPFIASAALFGLYIFFQIFSKEYINLLLTGYFFILGVIALCYLSSPVICSLVPAAIPNTPFHLKFTKGKGAHSEVIINYEFTLHDVICL
MCCSLIGVWYIINKHWIANNLFGIAFAINGVELLHLNKVVTGCILLCGLFVYDVFWVFGTNVMVTVAKSFEAPIKLVFPQDLL
ENGLTANNFAMLGLGDIVIPGIFIALLLRFDCSLKRNSKVYFYTTLVAYFLGLVTTLCVMHLFNHAQPALLYLVPACLGAPLILA
HIRGDLSAMFQYEDHPSNATDEKKDSTSNSSASNAGKVSPKTKKNK
>Silver-NTE-CTE
LGDCLKYVSKSYPNLASVFEVGKSTSGLSIWGIQLTENITGPQDLKPSVKLVANLHGDETVGRELLANLTRHLVNNYGSDNRI
TKLLHSTRIFIVPSLNPDGFSASQEGHCESLKNFVGRTNANGVDLNRDFPERLEEGRKGTGLLIGRQKETQVIMRWISMSDF
VLSAFHGGSVVASYPYDSGKGNEYSKSPDDDVFVQLAATYSQAHLTMPLGNNCPDDNFRNGITNGNNWYTVKGSGMQ
DFNYLYSNCFEVTFELSCCKYPLRETLSGEWENNIESILKYVESSHWGVKGVVSSVTGDRIAGSIIQVQGVNHNITTNKLGEY
WRLLLPGAYTIKVHAPGYMISSLASQIIFVAQRTVVSVRILRNIETSQVAPLRTYPISLCPVVVEPVLEEDIKNEYGFHHVVKFR

HHNYVQMTKELQDISKEYPNITRLYTIGKSVRGRDLFVLEIAEKPGVHIPGKPEVKYVANMHGNEVVGREMLLILARYLCEN
YGSDERVTHIVKSMRTHLLPSLNPDGYEVSHEGDYNGLDGRNNANNIDLNRNFPDQYGVSKVGFKMKQAHRKDCEVHR
GIKGFVWSSSGNPIKNAVVEVDGILKSVKTYKDGDFWRLLPPGKYVISAYAEGYDRQTVNVDLTHQQDGPMQNAYWVNF
TLQRDGLKSWSIENDFSQKDSVEQLHQYLNDDQIAEILKSLDAASPSIVRYYEKYGYHYLAISHEVSLGIEHKFHILVIGGLYGS
EPSGRELVLRLSRHLSAGHHLRDPNIRTLLQRSVVTLLPVVDTYIEAPCQVTDVRNNPLALEIINNGNTIGKRFEQFLHEQHFD
LIVSVPTSCCNYVEPNKIPRMWRHFLRPLMEVISSSVQGIRLTVLNSVHTPLRMAVVSVNGTTYSVSPNQAIAKIMLPPGSY
VAKISCKYHEPQLLPVSIEEGQLLDMKVILRETTIGASLAHAGNGIAGYVVDFNHHPVDNAIILVEGTNISQKVNDQGAFWV
PLQEGDFVMVATAKGYAPSTKLVKVVHMEPTQVLFTIMKDQDVIGLPRMGFIFLISLAIMLVLGGGLLGCMLCCNDNKSTK
GFALLREKSSFFSYSKGNHMQLPVQEDGLKTTAYYDESDIDEESSLSDEEVIDVHKLQKQAAILQQ
>PHM
MLATLLILIFSIIAAESFTTKKYSLVMPNVQPNVDELYLCTPVKVNYTKNFYIVGFEPKASMGTVHHMLLYGCQLPGSSQEVW
NCGEMVRDVEEDQHNPCKAGSQIIYAWARDAPKLNLPESVGFKVGGDSSIQYLVLQVHYAHSLHGATDNSGVYLFYTEKL
MPKLAGVLLLGTAGYIRPLSVEHMETACTIDEDKVLHPFAFRTHTHQLGKVVAGYRVRLENGRNEWTLLGKKNPQDPQMF
YPIEKNLTVRQGDQLAARCTMESHLYTTTFIGATNKDEMCNFYLMYWVEQSEPLSKKYCFTSGPPNYYWNNPGNLNNIPD
GASSLN
>Prohoromone convertase 2-Amontillado
MIRPGGWSLILGAWILVLITSYSCTALVFTNSFYVRLRGDGGQEAASLVAKRTGFDNFGPVLGSQNEFHFVHRGLQEARTK
RSIPHTRRLKVDPLVHMAVQMAGFKRVKRGYKPLVVENLVGEMKPGRDPTDPYFQLQWYLKNTGQNGGKARLDLNVEA
AWAQGVTGKNVTTAIMDDGVDYMHQDLKFNYNAKASYDFSSNDQFPYPRYTDDWFNSHGTRCAGEVAAARDNDICG
VGVAYDSKIAGIRMLDQPYMTDLIEANSMGHEPNLIDIYSASWGPTDDGKTVDGPRNATMRAIVKGVNEGRNGLGNIYV
WASGDGGEEDDCNCDGYAASMWTISINSAINDGQNAHYDESCSSTLASTFSNGAKDPNTGVATTDLYGKCTTTHSGTSA
AAPEAAGVFALALEANTQLTWRDIQHLTVLTSKRNSLFDAKGRFHWTMNGVGLEFNHLFGFGVLDAGAMVALAKQWRT
VPARYHCEAGSINRLQKISSNQPLFLKIETTACQGTDTQVNFLEHVQAVITLNSTRRGDVELFLTSPMGTRSMILSRRVNDN
DHRDGFTKWPFMTTHTWGEYPQGTWTLEVSFNSENVQSGFIKEWRLMLHGTSDPPYTGLPVSDHHSKLAIVKKAHEERN
SRSKHVYY
>CPM-IsoformA
MLTGGWLWLLLLCIASAEENRIKKFIDANDKSRFSADEPSSRVIAGGYQPSSYIRDTNSLEMKYHDFEQMTKFLRTTSSKYPN
LTALYSIGKSVQGRDLWVMVVSSSPYEHMIGKPDVKYVANMHGNEAVGRELMLHLIQYLVNSYSVDPYIKWLLDNTRIHIL
PSMNPDGFEVAREGQCDGGQGRYNARGFDLNRNFPDYFKQNNKRGQPETDAVKEWTSKIQFVLSGGLHGGALVASYPF
DNTPNSMFQSFSSAPSLTPDEDVFRHLALTYSQNHPTMHKGRACKSGSPAFTDGITNGAAWYPLTGGMQDFNYVWYGC
MEITLELSCCKYPPSTELPKYWEDNRLPLVKFLAEAHRGIHGFVLDENGNPIEKASLKIKSRDVGFQTTKYGEFWRILLPGVYK
LEVYADGYTPKDMDFMVVEEHPTLLNVTLYPAKVEGAGERGDRPYHIYTGANSGFYRPPHQHYYHHQIPPPKKPGSTDSGI
FSSLTSGFNNFVNNIFG
>CPM-IsoformB
MLTGGWLWLLLLCIASAEENRIKKFIDANDKSRFSADEPSSRVIAGGYQPSSYIRDTNSLEMKYHDFEQMTKFLRTTSSKYPN
LTALYSIGKSVQGRDLWVMVVSSSPYEHMIGKPDVKYVANMHGNEAVGRELMLHLIQYLVNSYSVDPYIKWLLDNTRIHIL
PSMNPDGFEVAREGQCDGGQGRYNARGFDLNRNFPDYFKQNNKRGQPETDAVKEWTSKIQFVLSGGLHGGALVASYPF
DNTPNSMFQSFSSAPSLTPDEDVFRHLALTYSQNHPTMHKGRACKSGSPAFTDGITNGAAWYPLTGGMQDFNYVWYGC
MEITLELSCCKYPPSTELPKYWEDNRLPLVKFLAEAHRGIHGFVLDENGNPIEKASLKIKSRDVGFQTTKYGEFWRILLPGVYK
LEVYADGYTPKDMDFMVVEEHPTLLNVTLYPAKNESFIVHKQVTSSPISTVQSTNGQNRESIMFPEY
>CPM-IsoformC
MLTGGWLWLLLLCIASAEENRIKKFIDANDKSRFSADEPSSRVIAGGYQPSSYIRDTNSLEMKYHDFEQMTKFLRTTSSKYPN
LTALYSIGKSVQGRDLWVMVVSSSPYEHMIGKPDVKYVANMHGNEAVGRELMLHLIQYLVNSYSVDPYIKWLLDNTRIHIL
PSMNPDGFEVAREGQCDGGQGRYNARGFDLNRNFPDYFKQNNKRGQPETDAVKEWTSKIQFVLSGGLHGGALVASYPF
DNTPNSMFQSFSSAPSLTPDEDVFRHLALTYSQNHPTMHKGRACKSGSPAFTDGITNGAAWYPLTGGMQDFNYVWYGC
MEITLELSCCKYPPSTELPKYWEDNRLPLVKFLAEAHRGIHGFVLDENGNPIEKASLKIKSRDVGFQTTKYGEFWRILLPGVYK
LEVYADGYTPKDMDFMVVEEHPTLLNVTLYPAKVGGRNRDDMTLYQKPWHPPWLVSIHKPPQSAPGMEHNEEGAGER
GDRPYHIYTGASTHSLTTATFITCALPITIVSWPALTNHLSLPI
>Prolyl endopeptidase
MSNSFYRFAHISSIVSSRIISTSYLSKFTKLSVNYFQTRRLLSKEKLSRPQEMKFNYPSARRDENIKENLFGVQISDPYRWLEDP
DSEETKKFVDAQNNISVPYLHEAKDREKINAKLTEMWDFPKYGCPFRRGDKYFYHMNTGLQNQNVLYKLDTLDAEPQVFL
DPNLLSPDGTISLAGTAFSEDGKIMGYALSESGADWVTFHFKKVDDGENLPEVLEKTKYTSLSWTHDNKGAFYGCFPDVEG
NKAVGSESGVVKNQKLYYHRIGTPQSEDILCVEFPEDSGYILDATVSDCGRWLVLEPRKMCHFNLLYFADLSALPNGITGPIQ
LTEVVSTLEADYQYITNTGSKFVFRTNKNAPNYKLIVIDFDNYTRENWITLVPEHETDVLDWAACVATDKLVLAYVHDVKSV

LHVHNLADGSFIQELPLPMGTVSGFSGKKKYPEIFYDFTSFFTPGTIYRCDLSQSPIPQPTVFRETTINGLDPEMYELEQVFYPS
KDGTKVPMFVMYKKGTVKDGSNPCLLYGYGGFNVNLQPVYSTFRLVFMKHLNGVVAVANLRGGGEYGEKWHDAGRLL
NKQNVFDDFHSAAEYLIQNKFTNNKMLAIQGGSNGGLLTAACSNQRPDLYGATISMVGVLDMLRYHKFTIGAMWISDYG
NPDEEEHFKNVLKYSPLHNIKIPEGQYPATLLITADHDDRVVPAHSLKFIATLQHTLENHIGQINPLLVRVDTKAGHGAGKPT
SKKIEECTDILSFLQRTLNLTYVG
>RPRC006957-Furin-like protease 1
MVRGEMIRVWTVLLLAVEVCCHYTSQWAAHIEGGYSVASNIAHAHDFTILAEIFPDTYHLEHRKVAKRSIKPNTQYHDNLIS
EKQVKWAKQQRARRRKKRDYLTRKDLGFSDSHVYLNDQRWPQMWYLNRGSGLDMSVEGAWKEGVTGKGIVVTILDDG
LEKDHPDLRENYDPQASFDVNSHDDNPMPRYDMIDSNRHGTRCAGEVAAVANNTICSVGIAYKASVGGVRMLDGDVTD
AVEARSLSFNPQHIDIYSASWGPDDDGKTVDGPGELAMRAFIEGVTKGRGGKGSIFVWASGNGGRDHDNCNCDGYTNSI
WTLSISSATENGLVPWYSEACSSTLATTYSSGSTGEKQIVTTDLHHQCTSTHTGTSASAPLAAGICALALEANTQLTWRDMQ
HIVVATAKPANLRAPDWTTNGVGRNVSHSFGYGLMDAAAMVQLARNWPTVPQQHKCEVSAPHIDKVIPAKSHVSLSLN
VKECAGVNYLEHVQAKISLLSQRRGDIEIQLTSPAGTKTTLLARRPHDISRSGFRSWPFMSVHTWGENPIGVWTLEIHNEAR
FLAQLSDWNLIFYGTETAPGTDLDGFSSSKKKGDVLASPVSGPVDNNVETSRGFTGQGSPVSSGSGEDSSPPGWPDLAQR
TSSQQSLEGLGIGHGDMDLVPISSTACRVTSRACLECSQGYYLWQGHCEAKCPPGTYATESICTSCHYSCEHCTGTNDYECT
QCYPDAEFYNVSSFESYCYPKSILPWLNYTKWYYRTCFGLVINGILLSFVGILFYFRRGCSFNKKKDHLQPTIINSVRYKLAQNE
SDSEN
>RPRC002472-Furin-like protease 2
MKAVVLIVSLFWIKTAVTSNRPQPIYTNQFAVHVPKGKEYADAIAERHGFINIGQIGSLKEYYLLEHRRLHKRSLSHSQEHHD
LLEAEEHVHWYQQQTEKRRVKREYAKRDIFSGHPFIPSYDSDFSSSLFTHRHSANRNHYRSGGGLSSFSFPDPLYKEEWYLN
GGAKDGYDMNVAPAWAKGYTGRGVVVSILDDGIQTNHPDLAANYDPAASTDINDNDDDPMPRDNGDNKHGTRCAGE
VAAVAFNSFCGIGVAYNASIGGVRMLDGSVNDAVEARALSLNPDHIDIYSASWGPEDDGKTVDGPGPLARRAFINGVTTG
RKGRGSIFVWASGNGGRHTDSCNCDGYTNSIFTLSISSATQGGYKPWYLEECSSTLATTYSSGTPGHDKSIATVDMDGKLRS
DHICTVEHTGTSASAPLASGLCALALEANPDLSWRDMQHIVVMTSNPAPLLKESGWITNGVNRKVSHKFGYGLMDGAA
MVTLAEQWTSVPPQHICKSHEVIEDRPIDPSFSSVLTVAVDASGCPGTVNEVRYVEHVQCKISLRFFPRGNLRLVLTSPKGTP
STLLMERPRDVVSSNFDDWPFLSVHYWGENPRGRWTLQVINAGNRHVNQPGILRKWQLIFYGTVADPVRLRLPQSSTTN
QDFTFPSVAPPLNSFFPSAPSQDIFSGFRNLPNIFTASGSENKKMKVLMDLNELAEGENCHESCATCAGTTQDSCLTCAPGH
LYMTDLGLCLQRCPDGYYEDQVSNSCIGCYGNCASCPEHPSVCGSCDHHLILFNGTCVTGCPKGTFETDDYRCGECDSSCE
TCLHDGTTGCVTCKGGIRAGSSGKCHNKSFECPPHCVLCTLQEDICTECSAGYIIDINGDCVIDEKSCKPGEYTTTNGCEPCE
NLSNCTKCPPGLNIINNTCVAFCGTGYFSDSGWCRPCAEHCAECLGDRQDQCLSCVDEYKFASGYCMESCPGGTYKTQW
GCLKCHHFCNECSSEGPYACTSCSSGRFLDAATKLCMPCHPCNGTNTHQCCHCDPTTGKCHLPAGKRRITSEQAAKAHED
ELLAEASAPHNFFTTLTSNNHRNAFTGTTLVTIASCAIVVALFGLIFTVLQLRSRDRDRRGGYMKVPIQDMNFSTLGENPRNS
KIRFSRGDSRITDSDEEEEEQVALNMTAEKS
>RPRC013490-Furin-like protease 2
MEAKLRPDHLCTLEHTGTSASAPLAAGMCALALEANPNLTWRDMQHIIVMSANPAPLEKEAGWTINGAQRKVSHKFGF
GLMDGEAMVNLAEQWTSVPAQHICKTPVIAENRILDKTYSSETKFHTNVTGCEGTESEVRYLEHVQCKITLNFQPRGNLRL
VLTSPQGTPSTLLSQRPRDEVSAALNDWPFLSVHFWGEDPRGIWALTVINDGAKKVSANGVFTKWQLIFYGTDTKPVNLR
PPSLQMYRKKQDLSDPAFYKEKVEGEAGYDDNVVDDAVEVLYNCHLECDSKGCYGPANNQCISCAHYKLNKSCVASCPEE
GYYASGGLCLPCHVDCRTCTGPGYHNCLTCAPHLYYITDLALCIHACPHTYYQDVDTKRCISCHETCASCQDGPTLCSSCDSH
LVHHSNSCLASCPAGTYLNHHQRCAPCHPSCDTCIGGKITDCLPPSRSATCSDGQCSECPPGLFLQNSTCVSGCDPGWYHI
GKSCKRCWPGCLGCYGGRKDECVSCVAGRMLARGQCRLHCPRNMYSTSTGCANCHHFCLTCNGGGAYSCTACSSGRYL
DEESGLCYSCHPTCLTCSSAAQNACTSCPQNLILDSGQCISYDDTDGSNCINCIENQDTKIHSFGAGKRRISEAVSWSGIVDE
PLQAFHRGKPPEYSPFVTVTVIAVIACLAIVVLFALLFATLQVR
>PAL1
MIFNNHILFITTWINLISSKISKVGALHDPHKETYLDLTTSKLAVPEKWLPKLWLDPSIKLGQVSGLCVDRDNYVYLFHRADRV
WSMNSFTADNIFSQRNLGPIRNHTVLVLGSDGTIHRKWGAEMFYLPHGITVDNEYNVWVTDVALHQVMKFSASSWTTP
LLSVGVAFTPGSDSRHLCKPTSVAVTSNGDFFVADGYCNSRILKFNSDGDKILEWGRPTVGGGFRVPTPGEFLVPHALTIAE
DLGIICVADRENGRVQCFNLLNASFAFQLKSEHIGPRLFSVAYCKSKGLFFLVNGEAFQRNIPVQGFVMTAKGDIVGKFGPQ
LKTPHDLAISSNCDTLYVGEIDPYIAWKFQLKGNLSSTSVAAPVMPSSSASTMDSNVSRGEVEGLAGAIMVTGACLVFAASL
LIAALVYSRSRRTGTSDTIRLLPDSTISDY
>PAL2
MAPVFTSLTALFVCCIVSNHVDATSDIRQDFYYKLRSMLQTTPFPEVPKGVILRPVEVQGWGRNLTELGQVSGVSVNARGN
PVIFHRGPRIWDEQSFNSSHYYQQLEKGAIEVDTVLTLDKTTGEVISSWGKDMFYMPHGITIDHHSNTWITDVAMHQVFK
FPRGWITPSLILGELFTPGHDESRFCKPTSVAVASTGEIFVADGYCNSRVLKFNHKGVLQRIFPQYEEFLSLLVPHSLALIEKQD

LLCVADRENMRVACWAAETGYKSAPNLPFSPFTTHQPDLGRVFGIAALGDLVYAVNGPTSSHIPIQGFTINPMAEVILDHW
QPKSEELKNPHGIGVSPLGDALYICEIGPNRVWKFALEAKHI
=============
Takeout genes
=============

>RproTo1
MLKPLVLTFLLIYMAQAAKLPKNWKKCKRNDPKLDDCLRVAVTFAARDLKDGNANLGILPLDPLRIDQLVIDQGQGPVSVK
MVFSNFSISGHRNVEVLSVKNDWKDVYLKAIVPKMTLRGKYKMDGKVLTLPIRGEGNCSLDAEDFTSSLHLVLRNNTKNGK
HYFAVEKFDVLKLDAEKGHVHFDNLFNGDKSLGDAMNRFLNANWREILTEITPALSKSFGLAYMRISNRILSKVPGDELFLS*
>RproTo2
MWTAALVLLSASAFCQAAKLPKNWIVCKKSAPDAGDCWKRAMEFSIQDLKNGSRTFGILPLDPLRISKIKIAPGDGPVSVVL
SFHDLDIIGISNVKISNVKNDWKVVTFNAANPRVTLVSKYVMDGKVLTLPIKGDGPCRIDIDNFKSNFTIRFKKISRGGKEYLE
VTKFQLQFTASNAKLQFDNLFGGNKALGNTMNKFLNENWEEIVNELSPALAQAFGVAMKAVSNKILTQIPFEEINL
>RproTo4
MHVLCMLLMGWAFIFHVQSAKLPKTWKTCKKDDPRMNDCLKIAIEEAVHDLVGGNPSLGVFPLDPMHFDTVSIDQGHG
PVSIKLDFKHLEIIGVKDLKITNLKTDWKEMHVDFIVPAVIAVGTYNVTGQVLILPIQGNGFCNLTFTNFSGSGQLKFKEIQKN
DKKYYQISHFDFIFDAEKMDILLENLFNGDKALGDNMNVFLNQNWPEILKELRPAVSKAFSSAFKEVGNRVFSKVPLELISPP
>RproTo3
MFKFIVYLIYLLVLISDMAQPGKLPKTWKTCTKGSPEMNECLKGAIEEAIHELADGNPSLGVLPMDPFHFDTITIDQGHGPVS
IKLDYTDLDLTGIGDLIIKSVKTDWKEMHFDAEIPTKVVLDGKYKIDGKVLVLPINGEGHCRIEFTKFKSFAQLKLKEIEKGSKHF
YEITNFEFDFDADGVHIQFDNLFNGDKALGDNMNVFLNENWKEILQELKPAISGAFGAAFKEVGNRVFGKIPIQLISPS
>RproTo5
MMFKQLFYICAIYGVVESAKLPKTWTACKSNDPKKEECLKGAIQHAIRDLSNGGKASLGVLPMDPLHFDMITVDQGDGPV
AIKLEFFNLDLIGLKTINVNSVKNDWKSMVVDLIVPKLTLRGQYKVNGKVLVLPIKGDGDCKLEFTNYKVIGNLKIKEVKKGDK
KHFEVVQFQIKPTQEKVFIQFDNLFNGDKALGDNMNRFLNENSQEILQELGPAISRAFGTAFKTISNRIFSKVPSNEINL
>RproTo6
MTRGYLLTIFFCLVGLSVAAKMPSKWKTCKRSDKNINECLKKAVEEAVKTLKSGNPSLGVIPLDPLHFNELNIGQGSGPVSIN
LNFKNMDIHGISTAKVKRFRADWNNYYLEAEATLNVPLVLLGDYTVKGQVLVLPIVGNGKCNLTFDNFVAKLTAKGHEITK
GKEKYMEIDKFTFDLETSKLRVFLGNLFNGDKALGNNMNVFLNENWQEILKELKPAISKAFGEAFRSIGNSVFSRIPLNQIAP
K
>RproTo7
MYILSSITFGCVCLYLVSAATVLPESWKICKKSDKKLNECLKSSIQTVVRELKTGNSKFGLPPTEPLLIEEVVLHQGNGQAVGLD
LTFRKLKMYGLSRVVVDKVSARYDKDQLSADFHIDGDFRIESDYTAKGRVLVLPINGAGKNVLKFDNLKGKLDMKFNKIKKG
ADTYYNVNKCDIMLDTSRLHLDFKRSSSVNEGLGQNLNTVLNENWKEILTDLKPAISKAFGAAFKDLANRVFSKVPLDKVM
PA
>RproTo8
MIFLMILFGLTHCVFGGGKEVPPGVVLCSRKHPKINDCVRNAIQETMPKFISGIKSLDIPSLDPFHVDNLIIDSKRDDGSPVSI
DLSWHNVNIKGIKSAKITSAKADWDNNMVSFEAALTEPVDITGNYNIDGIILILPIKGTGTFDLKLEGFRAHIKVHGKEEMRD
GDKYMMVDRLAFTFDIDHMEVHYYNLFNGDPVLGESMNSFLNDNWRDIIAEMTPSVEASFSKYFEQVARKVFDHIPIDKI
ALP
>RproTo9
MARQLLTLIASVVYILSTPVQGGQPELCSLSAKNLPQCLITAIQNVIPILVKGIPRYGVYPMDPMHIDTLDLSNSPGKTLNVKH
KFTNVDLQGLSSAVIRHVRLNPKTVEIDVNAILSKPVVLTGNYVSQGKILTLPIRGGGKFNITLINMRAVLKMRGHQTTKNGK
VHVMMDSVKFPFKIDKMELLFENLLRGNRLLSDTLNSVLNENWESVLEDMKPSFEEAIGSAFKEFANRIFNRIPQNYKCK
>RproTo10
MESKVLFLVLFIGVVCWTAPARSMNKLPVFLKSCKLDKNFDNCMMKNGNLAIPTLAKGDAKWKIPVLDPLKVPSVSISESS
AKSIALNITLNDLEIYGLKESKLVASRFDVNRKHVVWKIAVPRLTLLSKYKVAGRFLVLPITGSGPATVMLESPMLTYKFDYKLV
KRNNEDYLQVTKSDLKHTTTRLRINFENLFNGDKALGASTNKLINENWEEFNQQLAPSVVQSIGAILTQVLSNIVKTVPYENI
FTK
>RproTo11
MYCTLVFLLSLVGLSLAVKLPAYVKTCSRNDPKLSECALRHGREMIPKIIPGDPSIRLPRLEPLLLERVEIHPSGNGGSINMRLV
CYKCQVAGLSRASLNDIKLDLNKKHIDIRLSIPRLMVTGKYDVSGKVLVFPITGKGISNITLTDLDVNAGLDWKLIKRKRHEYS
QFIRHKVTFTASGLKINLSNLFNGDKLLSDNMNMILNANWREVLQDLKPSISDTVGQIIRITLNQIFDIIPYSQFFPDS

>RproTo12
MKTVSLNCMLRVSLLLTALPFTSQLKLPSYIKTCRQNDPKLNECVVKNGRLAIPKFINGDTKYRVPRLDPLDINELKVHQGSR
QLGLTMSLRDCKVTGLKHAQFIAARTDLKRRHIEWDFYHPFITIAGKYEMSGQVLVLPIRGRGTANITLTNMKTMFKFDFDL
VKKEDGEEYMKVTKTDIDTDIGNAIFRFNNLFNGDRLLGESMNRFLNENWKEVVKELGAPVVDSISQVFEIILSRITELVPYH
LVYTPV
>RproTo13
MVQPRSMTAAQISMALLTIVALSAALKLPSYIIPCKKDDPKLNECAVRHGQLAIPKFINGDPKYRAPRLDPLDITELRVNQGT
RQIGLRMILKNVKIYGLKNTVFTHARTGLRDKHIEWDFKIPKIEIISDYEVNGQVLILPITGKGKANVTLTKLDITYKYDWELIKK
NGKEYMNFTSSELLFENGRTFFDLKNLFNGDEFLGNNMNRFLNENWREVTKELGPAVGEAFSNVFRLLLTRIAAQVPYNDI
YLQE
>RproTo14
MLCSPLAFSAVVLMFVCSANPAKKDPPVYQLPSYIKRACSRNDPNINKCVVEVGGPAIKTVAKGDPKYRIPQLDPLHIKELRV
QQGTKQVGLELICSDCLMWGLQNTVFKSADVNWEDRKCRWEFTLDKMKVTGKYNVTGQVLLLPIVGSGDALINLENLKF
SYLYDWTYQKKNNGYTYVILGNSSFPFEVGHMSIKLENLFNGDPLLGGNMNRFLNEHWQDIMKDLGPAFSRSLAELTTGIL
TNMARVVPFDIMFPDT
>RproTo15
MFLLAVLCTWMVVAPEILAKDLPVYPLPPYVKRACARNDPNLNKCVVEVGSVALKTVIKGDPKYRVPVLNPMVIEELIVKQ
GTKQVGLTLVCKDCKLWGLENTKIVKADMNFNTNHHKMDFTLSKMRVVGKYNVSGQILLLPISGAGDAEFKFENLKFSIIY
DTAYEKKSNGRTYLKVVNGSFPMDAGNLVIRLDNLFNGDKLLGGNMNRFLNENWKEILKDVQPALSESLSELSERILNNISA
LIPMDILFPKK

==================
Nuclear receptors
==================
>Estrogen_related_receptor
MSHGDGEMGAVPVIKKEVELSGFHAPSSPSSTVYSPTITIPPSDLKFCETSGFSSEIQCSPGSPDHQYCSSTTHSLSNPVVQQ
MGEEMKEEEDMPRRLCLVCGDIASGFHYGVASCEACKAFFKRTIQGNIEYTCPGANDCEINKRRRKACQACRFQKCLNKG
MLKEGVRLDRVRGGRQKYRRNTEVSYLSSTHPNKPLLEDNKLLEILIGCEPEVLSILSEEDSCRAQGVSTLKILSDLYDKELVNII
GWAKQIPGFTELSLPNQMRLLQSTWTEILTLSLAFRSLPPTGKLNFAVDFILDEMLARECGAIEIYHLCLTVMDKLQRSLICKE
EYILLKALVLANSDARVDEASALKHLRDSIVSSLMDCVAVIRGDRSETATVLLCLPVLRQADHVLRTFWTGLLARGAVSINKLF
IEMLEPTLR
>Hepatocyte_NF4-NTE/CTE
QLCSICGDRATGKHYGAYSCDGCKGFFRRSVRKNHVYNCRFNRHCLVNKDKRNQCRYCRLRKCFKAGMKKETVQNERDRI
SGTRASFEEHQIVNGLSVGSLLNADVLTRHQTSGTVELGFDYNLSSKQVATINDVCDSMKQQLLYLVEWAKYIPAFCDLHLD
DQVALLKAHAGEHLLLGVARRSLHLRDVLLLGNNCIIPRYCGETAIAPDLDISKVGTRVMDELVVPLNEVQIDDTEFACLKAIV
FFDPNAKGLSNRLKIKYLRHQIQINLEDYISDRQYESRGRFGELLLTLPALQSITWQMIEQIQFAKLFGVAKIDNLLQEMLLGG
TTIENTNGQGTSLCNYQSSGGSPDSQNAISPVGSPEVCNPGTVVLRDIEVSMGTQEYMAFKQEPGI
>Hormone_receptor_like78
MEKGGSESNGIMTTTAEKIHSICLGVEICVVCGDRASGRHYGAISCEGCKGFFKRSIRKRLGYQCRGNQTCEVTKHHRNRC
QYCRLQKCLTMGMRSDSVQHERKPISVKKEYPTATNHYSLSTLHNNSVKLFMKREIGVDAYTVPPATIGQTNFGMYNPLSY
NIYSDNYNFKQESQNHHLTSCFDDTNFDESSDSDVIDTLCTTQDSKSMINSAIEIATKLGLNGNPCRSDDEESENVQVDGRL
VEDNTVQFEIQSPTPMPAYSDVHYIYESASRLLFLSVHWVRNVPAFQLFSSEAQISLVRGCWSELFALGLSQCAHTLSLPTIIL
SICNHLQNSVAQQKISPAKVKIVTDHICSLQDYVNSMVALSVDEHEYAYLKLITLFSPDNPGVHARRQATELQEKALQELRE
QIGENNYDRFAKLLLRLPPLRSLNRHIIEQIFFPGLGDQCDIDNIIPFILKMDNCDFIADQGSIQNRNMDMIYIKSENRPEEANI
>Hormone_receptor_like3
MDAASAALEKLFGGGGGGDTGRESGRNSKTENWPIKGEDVSSTTPPPASPTNQEQSQPQTTNSIRAQIEIIPCKVCGDKSS
GVHYGVITCEGCKGFFRRSQSSVVNYQCPRSKTCLVDRVNRNRCQYCRLQKCLRLGMSRDAVKFGRMSKKQREKVEDEV
RYHRAQLRAQVEQTPDSSVFDHSQTPSSTDQLHYTGYTYGNEVGSYTYNYSGQVTAAATMQYDISADFVDSTTTAYDPRP
SIDHVSDNSLMGNVVNTGGGGGGASGKVVESGGQSQRLAIKLEAEPLAGDGVVNTFVVDSTTSRQTSDQPSSKDSESNT
HTYTTSHIDPAQISELLSKTIADAHARTCLYTTEHIHNMFRKTQDISKLIFYKNMAHEELWLECAQKLTTVIQQIIEFAKMVPG
FMKLSQDDQIVLLKAGSFELAILRMSRYIDLSSGCVLYGDTMLPQDAFYTTDTSEVKLVTLAFEVSRGVAELKLTETELALYSA
CVLLSSDRAGLKGLAEIGRLGQAVLRALRIELDRNHALPIKGDVTVYDALLAKIPTLRELSMLHMETLGKFKRSTPHLDFPALH
KELFSVDS
>Hormone_receptor_like39

MEPGGGAAALSSLKWPESRGVVTSGQVTVTTINIHHSPHDKEDKYYPPHYNGGPSSKCSVLVSGEQMHVDEDSSAADAE
VSNFTSEPVNLRNKKREECRNSVERPMSWEGELSDSEVTAIKPEPEETAVENAESSVDNKTANKVGSHAEDEKPQSLKISEG
DKSVENVRTNQFNPQSELPLLMDKLLSSSSGGGYNGVLKGPGPTSSPDSAVYSCYSPAASPVTSRHILSSSSGISSPFTPSLSR
NNSDASQYGGSQHSSCYSHSYSSVSVSPTQFSPTHSPIQGRHMHRPGFPSPVLNARGVPEYPMQEEGNTLEDKFSSLTAD
MVQHSTGLASPGISRQQLINSPCPICGDKISGFHYGIFSCESCKGFFKRTVQNRKNYVCLRGSSCPVTIATRKKCPACRFDKCL
NMGMKLEAIREDRTRGGRSTYQCSYTVPAGLVEQKCAELTVQPRLEQQPATNLVPPLLQEIMEVEHLWQYNEADPKLVG
RIPKPPSGNPTDLMANLCNIADHRLYKIVKWCKSLPLFKNISIDDQTSLLINAWCELLLFSCCFRSMSSPGEIRVSLGKCISLTE
AKNLGLGPPIERMLNFTDHLRRLRVDRYEYVAMKVIVLLSSDTSELREPEKVRASQEKALQALQHYTLAHYPDIPSKFGELLLR
IPDLQRTCQVGKEMLSIKSKEGEGPSFNLLMELLRGDH
>Ftz_TF1
MLLEMEQQALSSLSMSHFNLSPGGGGGNDHTGGGGGGGGGSPNCPQQLYAGSPPLYPIQSSLSAGHSPSMAYNLDACF
FSSPGAGGGGGGGGGGGGGGGGGMEAVAAASSFQQQLAQVPPSDMPDTKEGIEELCPVCGDKVSGYHYGLLTCESCK
GFFKRTVQNKKVYTCVAERSCHIDKTQRKRCPFCRFQKCLEVGMKLEAVRADRMRGGRNKFGPMYKRDRARKLQIMRQR
QMAVQTLRNSGYSAVATTAGLGDSVALYQGASNFMHIKQEIQIPQVSSLTSSPDSSPSPITTVGLGQGNICPVLPTTVANPT
SQAGGPAALQLAGQEQRVGGGATPQATHPQVFSNDKLWTAASPPPTKNFQYEGSSQSGGTTPTTGGTKVSPMIRDFVQ
AIDDPEWQNSLYSLLQNQTYNQCEVDLFELMCKVLDQNLFSQVDWARNSIFFKDLKVDDQMKLLQHSWSDMLVLDHM
HQRMHNSLPDETTLPNGQKFDLLSLGLLGVPALSEHFTEISGKLTDLKFDPSDYICIKFLLLLNPDVRGLMNRKHVQEGHEQ
VQKALHDYCLTSYPQIQDKFNKLLLILPEIHVLASRGEEHLYMKHCSGGAPTQTLLMEMLHAKRK
>Hormone_receptor_like38-NTE
MDPLGFGFFKRTVQKGSKYVCLAEKSCPVDKRRRNRCQFCRFQKCLSVMVKEVVRTDSLKGRRGRLPSKPKSPQESPPSPP
VSLITALVRAHVDTSPDLSNLDYSQYREPNEEEIPMSETEKTQQFYNLLTSSVDVIRHFAEKIPGFSELCREDQDLLFQSASLEL
FVLRLAYRTRVMDMKLTFCNGVVLSRAQCQRSLGDWLHAILEFCQSLHAMDVDISSFACLCALTLITERHGLREPHKVEQL
QMKIIGSLRDHMTYNAEAQRKSHYFSRLLGKLPELRSLSV
>Ecdysone_induced_protein_75B-IsoformA-NTE
QFIFIVFTEFDGTTVLCRVCGDKASGFHYGVHSCEGCKGFFRRSIQQKIQYRPCTKNQQCSILRINRNRCQYCRLKKCIAVGM
SRDAVRFGRVPKREKARILAAMQQSTNSKCQEKALAAELEDDQRLLRTVIRAHLDTCEYTREKVEPMIIRAREQPSFTASPPT
LACPLNPNPQPLTGQQELLQDFSKRFSPAIRGVVEFAKRIPGFGLLSQDDQVTLLKAGVFEVLLVRLACMFDTQNNSMICLN
GQVLKRESIHSGSNARFLMDSMFDFAERLNNLRLTDPEIGLFSSIVVIAPDRPGLRNTELIERMQNKLKAGLQLMMSQNHP
NQPNLAQELMKKIPDLRTLNTLHSEKLLAFKMTEQQHLAEQNHQMWGNMVGEHEESKSPGGSTWSSSSDVAMEEVKS
PLGSVSSTESMCSGEVEYHTSSHAASAPLLAATLAGGCPVRHNRTINDDNKDIKPIIIHRTFRKLDSPSDSGIESGTEKVDKLTS
SAPTSVCSSPRSSVEDKEEQHQIEDMPVLKRVLQAPPLYDTNSLMDEAYKPHKKFRALRKECGSEIEPDNTTSTSTLSSTHST
LAKSLMEGPRMTAEQMKRTDIIHNYIMRGEACAWSTGQQQHHQQQQQQQPHQQQPHQTVHQQSVITTSTPNRGA
YVIVSNSTGGSTGASPVLYHHHHSNSPATSPCPSSTSLVELQVGTQPLNLSKKTPPPSPRPKALSLEA
>Ecdysone_induced_protein_75B-IsoformB
MMTEELPILKGILNGVVNYHNAPVRFGRVPKREKARILAAMQQSTNSKCQEKALAAELEDDQRLLRTVIRAHLDTCEYTRE
KVEPMIIRAREQPSFTASPPTLACPLNPNPQPLTGQQELLQDFSKRFSPAIRGVVEFAKRIPGFGLLSQDDQVTLLKAGVFEV
LLVRLACMFDTQNNSMICLNGQVLKRESIHSGSNARFLMDSMFDFAERLNNLRLTDPEIGLFSSIVVIAPDRPGLRNTELIER
MQNKLKAGLQLMMSQNHPNQPNLAQELMKKIPDLRTLNTLHSEKLLAFKMTEQQHLAEQNHQMWGNMVGEHEESKS
PGGSTWSSSSDVAMEEVKSPLGSVSSTESMCSGEVEYHTSSHAASAPLLAATLAGGCPVRHNRTINDDNKDIKPIIIHRTFRK
LDSPSDSGIESGTEKVDKLTSSAPTSVCSSPRSSVEDKEEQHQIEDMPVLKRVLQAPPLYDTNSLMDEAYKPHKKFRALRKEC
GSEIEPDNTTSTSTLSSTHSTLAKSLMEGPRMTAEQMKRTDIIHNYIMRGEACAWSTGQQQHHQQQQQQQPHQQQP
HQTVHQQSVITTSTPNRGAYVIVSNSTGGSTGASPVLYHHHHSNSPATSPCPSSTSLVELQVGTQPLNLSKKTPPPSPRPKAL
SLEA
>Knirps-like1
MNQLCRVCGEPAAGFHFGAFTCEGCKSFFGRTYNNLSSISECKNNGECVINKKNRTSCKACRLRKCLLVGMSKSGSRYGRR
SNWFKIHCLLQEQGNQTQQLKWDEDNNNVKGSPSKEPATHAHSPLWRLPPGLHPGAPLPPLFGFPPPPLHPFYLARPPAP
PPVPVPTLPPTLHPAPPPVLHHHHHHQPPSLPPQQPPPPPLPPPAPTTTNNTTTAAPPQTSIATQSPLELLRNLGPVQDSP
MDLSVKVQQQQQKGSGGIGSASSCSGAPAGSNSMAKLDDAQESEDDEDNQNENDDDLSSEELHHFQQHEDKKTPLDLT
CVKT*
>Knirps-like2
MNQQCKVCGEPAAGFHFGAFTCEGCKSFFGRSYNNLSSISECKNNGECVINKKNRTSCKACRLRKCLLVGMSKSGSRYGRR
SNWFKIHCLLQEQKQKNEESRLGSQIATLNRNKDELLLLGLDDYKTVSSPSISPHGSDSSDDKYNVHRHAATMAAAAAAAA
AAQLQHQQQQHHHHHHHHHQHQQNHHQQQQQQQHSTPPLYNLLPPLLVPHHYPLPYHPAFLPPSPSPNAFHHLYPHHL
NLGLSSLHDSPLDLSTKSQADTASSSASCSEDEDDEEQEISVDCLPDQPPITRTPLDLTTKVV

>Ecdysone receptor
MMGEEKRADEDWLGSGGGGGGVSPRPYNAQQPNGGGYPSPTMSSNSYDPYSPNSKIGREDLSPPNSLNGYSVDSCDGS
KKKKGTSARQQEELCLVCGDRASGYHYNALTCEGCKGFFRRSITKNAVYQCKYGNNCEIDMYMRRKCQECRLKKCLNVGM
RPECVVPEYQCAVKRKEKKLQKDKDKPVSTTNGSPEAIKSESEPIRVSFSSSLISLLKESQTLPPREGEMASKVPVNGVKPVSP
EQEELIHRLVYFQNEYEHPSEEEVRKINAGNDDEEQSDLRFRHITEITILTVQLIVEFAKRLPGFDKLLREDQIALLKACSSEVM
MLRMARRYDAQSDSILFANNQPYTRDSYSMAGMGDVVEDLLRFCRQMFNMKVDNAEYALLTAIVIFSERPSLIEGWKVE
KIQEIYLEALKSYVDNRPKPRPPTIFAKLLSVLTELRTLGNQNSEMCISFKLQNKKLPPFLAEIWDVNA
>Hormone_receptor_like4-CTE
MFHQLYPFVYLTPTSCTAYLFTGESVAETSTSSPDLGGPGSPCKMDQPSITPPPSSIIRGVSPDCGGRKSGEVRVKEELVFDG
GGPSGGSGGGGGGRSNTSSAVVGSGSSSSGSSNYWPPGPLPSPVRINGVRPELIGGAEMRPPRPCATVPRAAPTVIMGEA
GGVRTMVWSQPPPQEQPTTSNWPPPPNQEETAAQLLLTLGQESSGGCGSAGSGSSASSASSAPAGPSGNSGRALNMER
LWAGDLSQLPGAQQITALNLSWAKPLGQHLLDPSAKSSLPASTDEQEDDDQPMICMICEDKATGLHYGIITCEGCKGFFKR
TVQNRRVYTCVADGVCEITKAQRNRCQYCRFKKCIEQGMVLQAVREDRMPGGRNSGAVYNLYKVKYKKHKKSVRNGQLK
GLAGEKGKPAISPEHGIPPHLVNGTILKTALTNPSEVVHLRQRLDNAVSSSRDRTLSIDATLAMIQTLIDCDEFQDIATLRNLE
DLLEHKSDLSDKLMQIGDSIVYKLVQWTKRLPFYLELPVEVRIPFSFKLTCLDFFNL
>PNR_like-INT
MRQSTLNTPMEQTLLQNDHLTNKETACRVCGDKASGKHYGVPSCDGCRGFFKRSIRRDLKYVCKEKGSCVVDVTRRNQC
QACRFSKCLQVNMKKDAVQHERAPRSSNHQQSQLQNQQQSYPASVPVIYPSAAFLQPLHRPYLPAATVGLNYIPNVYLR
MDQHHHVNNISHSSLQTSSAQPPCSENEVSSSQEPTFNSDLKPSALTTLSGLSSENVHDSAIKLLALTVSCVRAIPSYQQLTC
QDRITLLEDSWKDLFLLTVAQWSLPIEEGRLIDECPVDDPDNQLSCDAHRVSTALTRLAQLRADHTEFACLKALLLFKPDMV
ENSRHEVEMLQEQTHLMLREYSGPRFSKLVLALLTISRVSLRSVLALFFRQKSYVNLETLLASPCLDRNTT
>Ultraspiracle
MYLQTNSTKMIKKDKPMMSVAAIIQSHWGRGLSLVENNLSLVGPQSPIDMKPDTATLHCSSFSPTPTSGPTSPQGFNIVPS
SVLGNGNKSSGNVYPPNHPLSGSKHLCSICGDRASGKHYGVYSCEGCKGFFKRTVRKDLSYACREDKQCLVDKRQRNRCQY
CRYQKCLSMGMKREAVQEERQRTKERDQNEVESTSSFHTDMPVERILEAEQRVDCKLEMKEEFNLGPMSDIICQATYNQL
WQLIDWAKHIPHFTSLPIEDQVTLLSAGWNELLIAGFSHRSILAKEGLVLGPGVIVNRNNAHQIGVGPIYDRVLTELVSKMRE
MKMDKTELGCLRTIILFNPEVRRLKSVQEVELLREKVYASLEEYTRISHPNEPGRFAKLLLRLPSLRSIGLKCLEHLFFCRVVGEA
PVDTFLAQLLESPEVSNRI
>Ecydose_induced_protein_78C-NTE
GFFRRSIQKQIEYRCLRDGKCLVIRLNRNRCQYCRFKKCLAVGMSRDSVRYGRVPKRSRERSSGEERVSTSDNSSSAATPPDP
ETALSPVYDLIVSVSQAHIANSDYSEENTRTLVRKPLPSPPSPVCAGPEVASSTAESLEQQRIWLWQQFATHVTPSVQRVVE
FAKRVPGFCELSQDDQLILIKVGFFELWLSHASRLTTDTTLTFSDGTFVTRQQMELMYSIEFVQSMFEFTAGFNSLLLGDME
LGLFSAVVLLSPDRPGVTDVKAVEQHQDRLVEALKLQVSCNLCGDSPSVLAKLPELRVVGAKHALILDWFRLNWDKLRLPPL
FAEIFDIPKCEEDLQ
>Hormone_receptor_like51-NTE
QSGGKLKGLGLSCVVCGDTSSGKHYGILACNGCSGFFKRSVRRKLIYRCQAGTGRCVVDKAHRNQCQACRLKKCLNMGM
NKDAVQNERQPRNTATIRPEALVDMDHERALREAAVAVGVFGDPSLIKMPVRVGKGIQPVPDRVPMPGSRNLLARTNIS
NKVIVLCMYSKHRKYLYMGAIFRGNDSPKYSALALRRSPEKNEEEQANQESLQSNQYPSSPRQESSKDDEDTGQNYLVFPE
CSLNLILKWFTEDSIDVTNEEPSSDSRIAGGPLIPPPDHPIYPPGQETVYETSARLLFMAVKWAKNLPSFASLPFRDQVILLEES
WSELFLLNAVQWCLPLEGSPLFSVADHLAVTQPNGKGCQVASEVRTLSDTLHRFRAVGVDPAEFACLKAIVLFRAETRGLK
DPSQVENLQDQAQVMLCQHTRGRQPGLGPRFGRLLLMLPLLRAVPTHRVEAIFFQRTIGSTPMEKVLCDMYKN
>Tailless-NTE
TGRILYDIPCKVCQDHSSGKHYGIFACDGCAGFFKRSIRRNRQYVCKAKGDGTCLVDKTHRNQCRACRLRKCLEAGMNRD
AVQHERGPRNSTLRRQILMMKEEPSSSASPTDLTMPNLSASPQRYFYPPPSIVVPSIMAPAGPEVRFPHSPLPSSCSLPLPLR
LINALSEPESICETAARLLFMNVRWARHVPAFTTLNMKDQVTLLEESWRELFLLGWAQLLPPTDLTQLIALRSTSVDAQPSSL
IRQAALFQECLAKLRSLTLDHHEFACLRAVLLFKTGNYKPVSLEGEGKSLVDVAGVAALQDQTSFALSKYISTSYPDQPYRLGK
LLLALPELRSVSPRTIEEMFFRRTIGPVTIERIICDMYKS
>Dissatisfaction-NTE
LFQETTARLLFMAVRWVRCLAPFQTLSKRDQLLLLQDSWKELFLVHLAQWSIPWDLSPVLGGPKARERLSQEDPLVPLEIHT
IQDILARFRQLSPDGSECGCIKAVILFTPETPGLVDVQPVEMLQDQAQCILGDYVRGKYARQPTRFGRFLLMIPGLRSVRQA
TVERLFFRETIGDIPIQRLLGDMYLMEKTYT
>Seven_up

MDENPKPVRNACYKHAYRNSVQRGRVPPTQPPSLPGQYALTNGDAMTAAAAAAAAAAGFNGHSYLSSYISLLLRAEPYPT
SRYGQCMQPNNIMGIDNMCELAARLLFSAVEWARNIPFFPDLQVTDQVALLRLVWSELFVLNASQCSMPLHVAPLLAAA
GLHASPMAADRVVAFMDHIRIFQEQVEKLKALHVDSAEYSCLKAIVLFTTDKQIATVVSHRSREHVKL
>Hormone_receptor_like96
MDAVPSDIFKCSDNKTCAVCGDVALGYNFNAVTCESCKAFFRRNALKDKELRCPFKESCQITSITRRFCQRCRLAKCYVVGM
KRELIMTEADKERKRRKIEANKAKLGCFNSTTKKVNSPTGNNTTVNEGTQTTVDTVDCGIQTEPLSQNSDCACIMSCFLQP
NVYLTPAVSPSFGLNLNSPTSPWLLGKIDGCIPLNSAHRTLLEDLVIANKALDAPVDQEISNLLGEEFKSCGSKNLLDVINLTAL
AIRRLIKMCKRINAFRTLCQEDQLSLLKQGCTQMMILRSVATFDADRNSWKIPHTEDRMSQIKVEVLKEARGNIYETHEAFL
RSFDTRASRDTAVICLLIAIALFDPTRNHLEDKVIIAQHQGTYYELL
