## Supplementary database 3 for "Transcriptomics supports local sensory regulation in the antenna of the kissing bug *Rhodnius prolixus*"

| | ANTENNAL FPKM VALUES | | |
|---|---|---|---|
| **ANNOTATION** | **LARVAE** | **FEMALE** | **MALE** |
| ENZYMES INVOLVED IN BIOGENIC AMINE SYNTHESIS | | | |
| DOPA decarboxylase (Ddc) | 3.4 | 14.1 | 18.7 |
| Tyrosine decarboxylase-2 (Tdc2) | 0.9 | 1.5 | 1.7 |
| Tryptophan hydroxylase (Trh) | 4.0 | 2.9 | 3.3 |
| Tyrosine 3-monooxigenase (ple) | 125.5 | 156.4 | 129.8 |
| BIOGENIC AMINE RECEPTORS | | | |
| **Muscarinic Acetylcholine receptor type A** | 5.8 | 9.3 | 14.4 |
| Muscarinic Acetylcholine receptor type B | 0.2 | 0.2 | 1.2 |
| Muscarinic Acetylcholine receptor type C | 12.8 | 14.6 | 15.4 |
| Dopamine 1-like receptor 1 | 1.7 | 1.1 | 2.4 |
| Dopamine 1-like receptor 2 | 1.6 | 2.4 | 0.2 |
| Dopamine 2-like receptor | 0.0 | 0.0 | 0.1 |
| Octopamine receptor in mushroom bodies | 0.1 | 0.3 | 0.5 |
| Dopamine ecdysone receptor | 6.6 | 14.9 | 45.0 |
| **Octopamine beta receptor 1** | 2.1 | 2.0 | 6.8 |
| Octopamine beta receptor 2 | 2.5 | 3.5 | 9.0 |
| Octopamine beta receptor 3 | 0.5 | 1.3 | 5.8 |
| Octopamine-Tyramine receptor | 1.5 | 0.9 | 2.6 |
| α2-adrenergic-like octopamine receptor | 0.0 | 0.0 | 0.0 |
| **Serotonin receptor 1A** | 0.2 | 0.3 | 1.4 |
| **Serotonin receptor 1B** | 7.2 | 7.0 | 11.4 |
| **Serotonin receptor 2A** | 0.3 | 0.1 | 0.9 |
| Serotonin receptor 2B | 23.7 | 35.8 | 37.6 |
| **Serotonin like receptor 7** | 1.5 | 1.6 | 8.1 |
| Orphan receptor 1 | 12.0 | 11.5 | 18.2 |
| **Orphan receptor 2** | 0.3 | 0.3 | 1.2 |
| OPSINS | | | |
| COpsin/Pteropsin | 0.0 | 0.0 | 0.2 |
| Long wave sensitive opsin 1 | 1.5 | 1.0 | 1.4 |
| Rh7 | 0.4 | 0.6 | 0.9 |
| UV opsin | 3.0 | 1.9 | 1.9 |
| NEUROPEPTIDE PRECURSOR GENES | | | |
| Adipokinetic hormone/corazonin-related peptide | 0.0 | 0.0 | 0.0 |
| Adipokinetic hormone (3 variants included) | 0.4 | 0.3 | 0.5 |
| Allatotropin | 7.3 | 19.2 | 22.9 |
| Allatostatin CCC | 36.5 | 33.7 | 22.0 |
| Allatostatin CC | 888.4 | 98.5 | 54.9 |
| Allatostatin A | 22.1 | 1.1 | 0.7 |
| Bursicon alfa | 0.3 | 0.1 | 0.0 |
| Bursicon beta | 0.1 | 0.0 | 0.2 |
| **Diuretic hormone 31 (3 isoforms included)** | 3.4 | 3.3 | 4.8 |
| Cardioacceleratory peptide alfa | 0.1 | 0.0 | 0.1 |
| Cardioacceleratory peptide beta | 0.03 | 0.1 | 0 |

| | ANTENNAL FPKM VALUES | | |
|---|---|---|---|
| **ANNOTATION** | **LARVAE** | **FEMALE** | **MALE** |
| Crustacen cardiactive peptide | 11.3 | 9.8 | 4.5 |
| CCHamide | 2.6 | 2.9 | 3.4 |
| CNMamide | 1.2 | 1.2 | 2.7 |
| Corazonin | 0.1 | 0.0 | 0.0 |
| Diuretic hormone 44 | 7.7 | 10.7 | 12.4 |
| Ecdysis triggering hormone | 0.3 | 0.0 | 0.0 |
| Eclosion hormone | 0.0 | 0.0 | 0.0 |
| Elevenin-1 | 0.4 | 1.2 | 1.0 |
| Elevenin-2 | 4.3 | 11.0 | 5.1 |
| FLP | 0.2 | 0.1 | 0.2 |
| GPA2 | 10.5 | 14.0 | 17.5 |
| GPB5 | 1.4. | 1.1 | 1 |
| kinin (Leucokinin) | 8.7 | 7.2 | 6.0 |
| Ion Transport Peptide (ITP) | 7.5 | 20.1 | 12.0 |
| Insulin-like peptide | 0.9 | 0.7 | 2.4 |
| ITG-like | 60.4 | 47.3 | 129.1 |
| LNPF | 2.5 | 5.1 | 2.6 |
| **MIP** | 24.8 | 1.2 | 1.4 |
| Myosuppressin | 5.0 | 3.0 | 6.0 |
| Natalisin | 2.4 | 2.3 | 2.1 |
| Neuroparsin | 5.1 | 0.7 | 11.5 |
| Neuropeptide like precursor 1 | 7.7 | 7.1 | 7.9 |
| NVP-Like | 11.0 | 13.4 | 14.2 |
| Orcokinin (3 isoforms included) | 15.3 | 56.4 | 101.0 |
| **Pyrokinin** | 0.0 | 0.1 | 0.2 |
| PDF | 9.6 | 5.8 | 8.1 |
| Proctolin | 0.4 | 0.0 | 0.3 |
| IDLSRF-peptide | 26.9 | 47.6 | 136.2 |
| RYamida | 3.9 | 8.9 | 8.0 |
| SIFa | 0.4 | 1.3 | 0.2 |
| **sNPF** | 0.2 | 2.5 | 1.9 |
| Sulfakinin | 0.0 | 0.2 | 0.0 |
| Tachykinin | 3.8 | 8.1 | 4.8 |
| **NEUROPEPTIDE PROCESSING ENZYMES** | | | |
| **Peptidylglycine alfa-hydroxylating mono-oxygenase (PHM)** | 159.5 | 213.0 | 210.7 |
| Amontillado (Prohormone convertase 2) | 5.9 | 10.1 | 15.6 |
| Signal peptidase | 76.9 | 71.1 | 89.6 |
| **Silver** | 21.1 | 16.6 | 34.4 |
| Prolyl endopeptidase | 30.5 | 22.7 | 59.0 |
| Carboxypeptidase M (CPM) (3 isoforms included) | 335.6 | 264.8 | 383.4 |
| Furin like protease 1 | 113.7 | 95.7 | 92.4 |
| Furin like protease 2A | 3.9 | 3.8 | 18.9 |
| Furin like protease 2B | 12.6 | 4.0 | 6.9 |

| | ANTENNAL FPKM VALUES | | |
|---|---|---|---|
| ANNOTATION | LARVAE | FEMALE | MALE |
| **PAL1** | 35.9 | 49.8 | 54.1 |
| **PAL2** | 32.5 | 75.6 | 62.4 |
| NEUROPEPTIDE AND NEUROHORMONE RECEPTORS FAMILY A | | | |
| **AKH Corazonin related peptide receptor** | 0.5 | 0.5 | 1.3 |
| Adipokinetic hormone receptor (two isoforms included) | 3.9 | 0.3 | 0.6 |
| Allatotropin receptor | 0.4 | 0.7 | 0.9 |
| AstA receptor | 0.2 | 0.0 | 0.3 |
| AstC receptor | 0.7 | 0.7 | 0.7 |
| Cardioacceleratory peptide receptor (isoforms B and C) | 0.3 | 0.6 | 1.3 |
| CCHamide receptor 1 | 0.0 | 0.0 | 0.1 |
| CCHamide receptor 2 | 0.0 | 0.0 | 0.0 |
| CNMamide receptor | 0.7 | 0.7 | 2.4 |
| **Crustacean cardioactive peptide receptor 1** | 12.1 | 9.8 | 14.1 |
| **Crustacean cardioactive peptide receptor 2** | 0.0 | 0.0 | 0.2 |
| Corazonin receptor (3 isoforms included) | 8.0 | 3.7 | 8.0 |
| Ecdysis triggering hormone receptor | 0.5 | 0.5 | 1.4 |
| FaLPa/Proc receptor | 0.7 | 1.2 | 1.0 |
| FMRFamide receptor | 0.7 | 0.9 | 3.0 |
| Ion Transport peptide receptor | 11.3 | 5.2 | 5.9 |
| GPA2/GPB5 receptor | 87.9 | 54.1 | 32.5 |
| Kinin receptor 1 | 1.3 | 1.5 | 2.2 |
| **Kinin receptor 2** | 0.7 | 7.4 | 14.1 |
| LNPF receptor 1 (only isoform 1 included in the GFF)* | 13.5 | 31.8 | 39.8 |
| LNPF receptor 2 | 0.0 | 0.1 | 0.4 |
| Lutropin-choriogonadotropic hormone receptor-Bursicon receptor | 0.3 | 0.3 | 1.3 |
| Myoinhibitory peptide receptor (2 isoforms included) | 0.8 | 0.6 | 1.3 |
| Myosuppressin receptor | 0.0 | 0.0 | 0.0 |
| Natalisin receptor | 1.7 | 2.9 | 5.4 |
| Orphan receptor 3 | 0.4 | 0.2 | 0.7 |
| Orphan receptor 4 | 0.2 | 0.2 | 0.1 |
| Orphan receptor 5 | 0.6 | 1.9 | 5.2 |
| Pyrokinin 2 receptor (3 isoforms included) | 5.2 | 1.9 | 9.9 |
| Pyrokinin 1 receptor | 0.3 | 0.4 | 0.1 |
| CRF receptor | 19.5 | 4.8 | 6.9 |
| **RYamide receptor** | 3.2 | 3.3 | 3.9 |
| SIFamide receptor | 0.5 | 0.7 | 2.4 |
| **SNPF receptor** | 5.3 | 1.6 | 1.5 |
| **Sulfakinin receptor 1** | 0.1 | 0.3 | 0.2 |
| Sulfakinin receptor 2 | 0.2 | 0.2 | 0.9 |
| Tachykinin 86C receptor | 2.1 | 7.2 | 5.8 |
| **Tachykinin 99D receptor** | 0.2 | 1.3 | 1.4 |

| | ANTENNAL FPKM VALUES | | |
|---|---|---|---|
| **ANNOTATION** | **LARVAE** | **FEMALE** | **MALE** |
| **NEUROPEPTIDE AND NEUROHORMONE RECEPTORS FAMILY A** | | | |
| **Calcitonin-like diuretic hormone receptor 1 (isoforms B and C)** | 41.8 | 81.1 | 116.7 |
| Calcitonin-like diuretic hormone receptor 2 | 0.2 | 0.3 | 0.6 |
| Calcitonin-like diuretic hormone receptor 3 | 7.5 | 79.3 | 131.6 |
| **Corticotropin-releasing factor-related like diuretic hormone receptor 1** | 6.5 | 6.1 | 10.9 |
| Corticotropin-releasing factor-related like diuretic hormone receptor (isoforms B and C) | 4.1 | 14.8 | 29.2 |
| Parathyroid hormone like receptor | 12.2 | 14.7 | 13.5 |
| PDF receptor | 1.2 | 1.7 | 6.1 |
| **TYROSINE KINASE AND GUANYLYL CYCLASE RECEPTORS** | | | |
| Eclosion hormone receptor | 0.3 | 0.3 | 0.3 |
| Insulin receptor | 0.9 | 0.6 | 0.9 |
| NPLP receptor | 1.3 | 0.7 | 1.3 |
| Potential neuroparsin receptor | 6.2 | 1.6 | 8.2 |
| **NUCLEAR RECEPTOR GENES** | | | |
| Dissatisfaction | 0.0 | 0.0 | 0.0 |
| Ecdysone receptor | 4.4 | 2.2 | 7.1 |
| Ecdysone-induced protein 75B (isoform A and B included) | 111.2 | 44.7 | 101.6 |
| Ecdysone-induced protein 78C | 0.1 | 0.8 | 0.5 |
| Estrogen-related receptor | 2.8 | 2.2 | 5.9 |
| Ftz transcription factor 1 | 1.5 | 1.0 | 8.0 |
| Hepatocyte nuclear factor 4 | 29.0 | 25.2 | 43.2 |
| **Hormone receptor-like in 38** | 1.8 | 1 | 4 |
| Hormone receptor-like in 39 | 11.6 | 4.8 | 14.4 |
| Hormone receptor-like in 4 | 1.1 | 0.9 | 4.9 |
| **Hormone receptor-like in 3** | 2.1 | 7.9 | 26.4 |
| Hormone receptor-like in 51 | 0.2 | 0.1 | 0.2 |
| Hormone receptor-like in 78 | 6.4 | 2.3 | 11.6 |
| Hormone receptor-like in 96 | 24.8 | 11.6 | 24.0 |
| Knirps-like1 | 26.0 | 6.3 | 7.9 |
| Knirps-like2 | 0.3 | 0.2 | 0.2 |
| PNR-like | 3.5 | 17.1 | 16.7 |
| Seven up | 0.2 | 0.2 | 0.5 |
| Tailless | 0.9 | 0.2 | 0.1 |
| Ultraspiracle | 35.8 | 11.2 | 29.5 |
| **TAKEOUT GENES** | | | |
| to1 | 2703.7 | 3575.1 | 2862.7 |
| to2 | 34671.6 | 3919.8 | 2425.5 |
| to3 | 134.7 | 4271 | 3082.8 |
| to4 | 5200 | 20859.5 | 14346.7 |
| to5 | 1.0 | 1.7 | 1.6 |
| to6 | 358.8 | 373.2 | 369.1 |

| | ANTENNAL FPKM VALUES | | |
|---|---|---|---|
| **ANNOTATION** | **LARVAE** | **FEMALE** | **MALE** |
| to7 | 204.2 | 380.5 | 453.8 |
| to8 | 734.8 | 5004.6 | 4869.7 |
| to9 | 23.4 | 15.2 | 16.7 |
| **to10** | 3.7 | 40.1 | 42.2 |
| to11 | 161.4 | 5.8 | 6.8 |
| to12 | 3.8 | 20.4 | 25.1 |
| to13 | 37.5 | 61.1 | 76.7 |
| to14 | 469.8 | 738.0 | 519.4 |
| to15 | 390.0 | 1059.7 | 946.2 |

Note:

**Genes in bold** are divided in two or more fragments on the *R. prolixus* genome assembly, so FPKM values were calculated by means of the sum of the counts from the different fragments divided for length of the complete coding sequence.

* Long Neuropeptide F receptor 1 isoform B was not included in the GFF file due to the sequence of the genome scaffold, which not included the position 387 (N for isoform 1 and S for isoform 2).
