## Supplementary database 4 for "Transcriptomics supports local sensory regulation in the antenna of the kissing bug *Rhodnius prolixus*"

CT and CRF DH receptor sequences
====================================
>ClecCT/DHR1-CLEC000667
MTDDVSNETYIDLNADILNVRRTQCLEFINESKPLEGLHCEPVFDTWTCWPATLAGSVATNPCPHFIVGFDPSRKAHKVCTE
NGTWFRHPESGQIWSNYTTCVNLDDLSLRQQVNNIYQAGYFISLLALLLSLFILSYFKSLRCPRNTLHMNLFTSFAFNNFLWL
LWYRLVIPNTDVIVKNRVWCQCLHVVLHYFLACSYAWMLAEGVYLHTLLVSAFTSEQKLVKILTACSWLVPFFFTALYTTLRL
ASGDTQQCWIDESDSKMVLFVLVATSMLLNFLFLCNIVRVVVTKLRAGPNQSARPSTALLQALRATLLLLPLLGLNYLLTPFR
PPDKHPWETYYEVVSALTASFQGLCVATLFCFCNGEVIAQMKRKWQYSVFRTRANSYTATTVSKSVYLQFVRSTAAPAAEE
EIV
>HhalCT/DHR1-XP_024216452.1
DSDFLQCAIYINESKSKNLEGSYCEATWDGWSCWQETPAGTTAYAHCPKFITGFDPNLLAHKICTENGTWFKHPDSGMV
WSNYTTCVNIEDLTLRQTVNNIYQTGYSISLVALLLSLFILSYFKALRCPRNTLHKNLFTAFAANNFLWLLWYRLIVPFPEVVIE
NGVWCQCLHVILHYFLLSCYAWMLAEGVYLHTLLVSAFTSEQKLVRILTVFSWTAPLFFIFLYSVLRLVFDDTDQCWINDSDY
SSVLVVLVVASMGLNLGFLCNIVRVVVGKLRAGPSQSSRPSQALLQALRATLLLLPLLGLNYLLTPFRPPNDHPLESYYEILSAF
TASFQGLCVATLFCFLNGEVMAQIKRKWQYATFRTRANSYTATTVSFVRSTAAPTAEEENV
>ApisCT/DHR1-XP_016661057.1
IVGTAYKICNKNATWFKHPISGAVWSNYTTCINHEDYNWTQQINTIYQTGYLVSFIALLLSIAILTYFKSLRCARNTLHTHMFT
SFAINNLLWLLWYRLVVEHPSVVLHNGWWCQILHVILHYFLLTNYAWMLAEGFYLHTLLVFAFTSEDTLVRWSWTLAWST
PLVVISLYTLLRTIYDHTSECWINESPFTEVLVVPVCMSMALNLVFLCNIVRVLWVKLQAGPSHLSNSTPSRTLLQAFRATLLL
LPLLGLHYLLTPFRPPKQHPWEPFYEVVSATTSSFQGLCVATLFCFLNGEVVAQIKRRWQFMFFRTRANSYTATTVSVRQFV
RSTAGPGDNEDKV
>TcasCT/DHR1-TC002694
MEQMRQHCLVFYKCEKDGEWNFNEQFNKSWVNYTTCINIEDFEFRQQIILIYCVGYGVSLVALLVSLALLTYFKSLRCARITV
HMNLFSSFAMNNFLWLLWYSLVVNDQDVLHENKLWCRVLHVVLFTFLISNYSWMLCEGIYLHTVLVSAFISERRLLRCMLA
LGWGIPLLTTSIYAPVRSVLGENVDELGRCWTQDGRFNKILMVPVVITVFLNVIFLVNIVRVLLIKLRKGPANGGSGSGASRT
SLQALRATMLLVPLLGLNFLLTPFRPEANHPWEYVYEVVSALTASLQGIQPFLMFFWYRDQNHSW
>DmelCT/DHR1-CG32843
MSDQIGNPNATFSGSGSGSGTNVASIAESVAESGPDFDALRAACETRLNASGQLAGSGGPGAEAGTHCAGTFDGWLCW
PDTAVGTSAYELCPDFITGFDPARYAHKECGLDGEWFKHPLTNKTWSNYTTCVNLEDLNWRHTVNLISEVGYGTSLLAILLS
LAILGYFKSLKCARITLHMNLFASFAANNSLWLVWYLLVMPNSELLHQSPMRCVALHITLHYFLLSNYSWMLCEGFYLHTVL
VAAFISEKRLVKWLIAFGWGSPAIVIFVYSMARGLGGTPEDNRHCWMNQTNYQNILMVPVCISMFLNLLFLCNIVRVVLLK
LNAPASIQGSCGPSRTVLQAFRATLLLVPLLGLQYILTPFRPAPKHPWENTYEIISAFTASFQGLCVAILFCFCNGEVIAQMKR
KWRMMCFSNRPRTNSYTATQVSFVRCGPPLPGEEKV
>AmelCT/DHR-LOC412591
MILSMRPSPTWLKMDVQTEDIVLLAGEQGRIRITITIVQISRLISEVANLISNMNASLKRCGHLASNFTKTLYCLMPANATAG
STLPDEEIHRIIMEREQECLQLEAMNSTPPPGPYCRLTFDGWSCWPNTPAGATAYVPCPNFITGFDASLRAHKYCELNGTW
FRHPESGQVWSNYTTCVNLKDLSWQQGINGLYEAGYAISLIALLLSLGILTYFRSLRCARITLHMNLFASFAVNNALWLVWY
RCIVANTDLLLNNGMTCRLLHIVLHYFLLTNYAWMLCEGFYLHTLLVSAFTSEQKLVKWLMILGWPVPAIIVTIYACLRATSN
DLTDTEQCWINEGNYMNVLVYPVCVSTLLNVLFLFNIVRVLLMKLRAGPSIGTQPSRSMRQAFRATLLLVPLLGLHYLVIPFR
PPKNHPWEHFYEVLSAITASFQGLCVAILFCFCNGEVIAQFKRKWEGSAFLRNRANSCTATTVSVRQIHKT
>AgamCT/DHR1-AGAP009770
AKHRYKPTRGTLQVTNPRTIPFTALNVEPHIRFPTKLSSSRVKTTSSPQPRNSMALSTTTDASVDEIAQKRLECLETLNETVDT
TISGFAHKVCSENGEWFRHPETNRSWSNYTTCINLDDFEAGCITLHLVLHYFLLTNYSWMLCEGFYLHTVLVSAFVSEKKLLK
WLLALGWGSPAFFIVLYGFLRGYASPPNDTIECWMNDSSFNKVFVGPVCISMLLNLVFLFNIMRVLLLKLKAPAGPQGAGP
SRTILQAFRATLLLVPLLGLQYILTPFRPGNGHPYERTYETMLACTSSFQGLFVAVLFCFLNGEVIAQVKRKWRTVFLRTRANS
YTATQVSVSTLTVLRV
>AaegCT/DHR1-AAEL010043
MTSSTTTDATDSEFDQRKLQCLEMLNATTEFTTTRSGPFCRGTWDGWLCWPDTAAGESALLPCPDFMDGFDPTRFAHKD
CDEDGEWFRHPLTNRTWSNYTTCVNLDKLEWMEQVRTIYETGYSISLIALILSLGILSYFRSLKCARITLHMNLFASFASNNTL
WLLWYRMVLADPEVLKHNGASCITLHLVLHYFLITNYAWMLCEGFYLHTVLVSAFVSEKKLVNWLVVLGWTTPGIVIMAY
GFLRGYAGTPEDTIECWMNESVYDNVFKAPVCISMLLNLLFLCNIIRVVLLKLKAPAGPQGTGPSRTILQAFRATLLLVPLLGL
QYILTPFRPDPGHSYERTYEIISAFTASFQGLFVAVLFCFFNGEVIAQVKRKWRTVFLRTRTNSYTATQVSPVRLQFIRSGPPIP
GEEKV
>Dmelhector-CG4395

MATTSSDSESQNVDVASQAQTQDNLRIFLKHLYAECVFRYQNVTYDTDDPSFSLGPATDYDSDLPENFSPVPRYLENAAM
NEGVIDMRNVDEELAEKEELMATVVSATMATNQKENRLFCPLNFDGYLCWPRTPAGTVLSQYCPDFVEGFNRKFLAHKT
CLENGSWYRHPVSNQTWSNYTNCVDYEDLEFRQFINELYVKGYALSLLALLISIIIFLGFKSLRCTRIRIHVHLFASLACTCVAW
ILWYRLVVERSETIAENPLWCIGLHLVVHYFMLVNYFWMFCEGLHLHLVLVVVFVKDTIVMRWFIVISWFSPIPIAIVYGLAR
HFSSPDNKHCWITDSLYLWIFSVPITLSLLASFIFLINVLRVIVRKLHPQSAQPAPLAIRKAVRATIILVPLFGLQHFLLPYRPDAG
TQLDHFYQMLSVVLVSLQGFVVSFLFCFANHDVTFAIRTLLNKLLPSLVTPPPAGSNTGQMATTTPSRELGV
>ClecCT/DHR2-CLEC025204
MRAYCPAAFDGWSCWNTTPSGETALAPCPYFVTGFDIKRFAFRKCLDNGSWFRHPDTGQPWSNYTTCIDMDDLEVREIV
NTIYVVGYYVSFAALVLSLLIFLTFRSLRCTRIAIHIQLFSSFAANNLMWIIWYKLVVGNTSVVQRNQPFCQILHIVLQYLMVA
NYLWMFCEGLHLHLALVVVFVKDNSAMRWFYCIGWVFPGILTAIYASVRYWYTEETRQCWMNESHTQWILTAPVCFSM
LASLGFLINVVRVLLTKLHCNSANPAPIGLRKAVRAALILVPLFGIHHILIPFRPEPNAPGERAYQIFSALLVSLQVIISYRYKNTG
KVLMFKCLSISDRTATIKPNFN
>HhalCT/DHR2-XP_024217979.1
MKLKRECEIRKRHQLEMFPTKVADRTGLRPYCAATFDGWSCWNTTPSGEIALAPCPNFVTGFDTKRFAFRKCMDNGSWF
RHPDTGQPWSNYTTCVDMDDLEFRKIVNIIYVVGYSVSFAAIVISLIIFLTFRSLRCTRIAIHVQLFSSFAANDLMWIVWYKM
VIGNPIVVQENRFMCQALHVLLQYLMVANYLWMFCEGLHLHLALVVVFVKDDNAMRWFYFIGWFLPAILAGIYALVRSSY
PDETSQCWLSESHTQWILTVPVLLSMLASLGFLINVVRVLLTKLHCNSANPAPIGLRKAVRAALILVPLFGIHHILIPFRPEPNG
PGERVYQVFSALLVSLQGFCVSLLFCFANVDVHAAFKAMARRIRRRATDNGNLTATQTREVV
>ApisCT/DHR2-XP_016661373.1
MYCRSLHVALQFFMVANYMWMFCEGLHLHMALVVVFVNDVNAMRWFYAIGWGVPVALTVLYVSWRSNSEDTAQCW
MHESHCQWVLTVPVFASILTSLMFLMNVMRVLLTKLHRNSTNPAPIGVRKAVRAALILVPLFGIHYILIPYRPKHKTTVETVY
QIFSVVLVSTQVSDARLPSTVRACISIATVKNFAKIIYYYCTNVWIPLRCTSMDIGRDPRKVHLIHRSRSATSIS
>AaegCT/DHR2-AAEL006490
MIETIEEPQRNQTGLFANDNQSNWFPMAMEERPTSGIAASADGIQYDRLKYADPEALMRHLYYACMRNVTNSGSFPAE
QAVTVLRSISASNARHDQQTQLFCPRRFDGWTCWESQPAGTIAQNFCPNFVLGFDASRLAYRICHANGSWFTHPESGRE
WSNYTNCIDVDDMKFRRLVNDLYIGGYTISLVTLIVSLCVFFSFRTLKCTRIRIHINLFTSLALSCAFWILWYKFVVEDPDVTNR
NGNWCIALHILLHYLMLVNYFWMFCEGLQLHLVLVIVFIKDAIAMRWFFTIGWILPVAFVSIYASVRNKYTLDTEHCWMNE
SHAMWLLTIPVCFSLVASLVFLINVVRVLLTKLNSTSPNPAPLGLKKATRATLILIPLFGLQHILLPFRPDKGCELERYYQVVSAV
LISLQGACVSCLFCFANHDVIFAIKCQLSRFFPTLVHHPFRESYNGGQPATQSRDMVV
>TcasCT/DHR2-TC013321
MDVTNGTNCGGKYTRPGYCPEIFDEMLCWPETLGGTTVNQSCPKKMGYDSRRFAYKDCLENGSWFKHPKSGKIWTNYT
TCVDHEDLAFRTHINHLFVIGYSISLAALVISLAIFFTFRTLKCTRIRIHIQLFISFALNNLMWIIWYKEVVPNPFVTIRNELWCQ
ALHLVVHYLMLANYMWMFCEGLHLHLALVVVFVRDAETMKWFFALGWGAPFIIVLIYSVVRIFILKDNYMCWMADSYYS
SWILTAPVCISLLVSLIFLINVLRVILTIMHPNSANPAPMGLRRAARAALILIPLFGLQHILIPFRPDMYDPYEHLYQYVTVVVVT
LQGLCVSCLFCFANQDVHQAIRGFMHRKVYRTTRWSNYHYTGAADSAGVYVVNGSSHCNNVGLLSLKRKSTTTVKL
>AgamCT/DHR2-AGAP001175
MTIREDAAELVRVDLAKANGSAYFGQTGMIDAGGRVGDGQDGQIQFDRLEFESPEDLMRHLYYTCMRSNTSEPPWTTD
GSTVFCPRRFDGWSCWEPTLAGTVAENWCPKFVLGFDPRRLAYRTCHENGSWFVHPHSGREWSNYTNCIDTEDMQLRR
LVNDIYIGGYTVSFLTLIISLCIFHSFRTLKCTRIRIHIHLFTSLALSCLFWIVWYKFVVEDPDVLNANTPLTSICVTVPGNSCAYKQ
SSLQGWCVGLHILLHYLMLVNYFWMFCEGLHLHLVLVIVSIRAHPGDIPRGRLECHRATISQGWCRLLPFSVRFNSIRLIFPLF
HRRHRHHIRTDAFFSVFFRVLLTAGKKVFIKDIVAMRWFVTIGWILPMALASLVFLINVVRVLLTKLSSTSPHPAPLGLRKATR
ATLILIPLFGLQHILLPFRPEKGCSLERYYQIGAALLISLQGLCVSCLFCFANHDVIFAVQCYLSRFFPDLVTHPFLESNGGQPAT
QSRDVVV
>ApisCT/DHR3-XP_016661370
MADYQYGLSNIQDNHILHQVIAERRKMCESVMDDPPMSDSDAWCPGEFDGWTCINRTKAGEVAKFPCPYFILGFDTKRF
GQKTCLPDGSWFKHPDSNKTWSNYTTCVDLEDLKLRNQVNMIYKWGYTVSLAALAVSIFIFFYFRSLTCTRIQIHKNLFISLA
VNNCLWLVWYEAVVDNLPVLMTNGLGCKLLHVLVQYFLVATYFWMFCEGLYLHTLLVVTFLTESKVMPFLHTIGWGIPAL
LVSTYAALRTATPGETLHCWIHESLYSWILSGPVCLSMLANLVFLINIVRLLLVKLHARQITMSSPKHVAPRERAQSFSLRSFK
RNNCSVDSYDKAPASSSRTGKAVRATLILIPLLGLQYIVTPFRPEPGTSWEPVYQVTSAVVASCQGLCVALLFCFCNGEVLSE
MKKRWKHCWINKNGSWNPCMRGTSVSSPQHHPRLLVTEDQQTQRTGSIQVLPRKDSAECLQNVQL
>HhalCT/DHR3-XP_014287171
MIYTREEIDNITAQIKKECFAVAENYTEGLFCPREFDGWTCLNATPAGTVLHFPCPYFQLGFDPKRTAHRPCLENGTWFRHP
ETNKPWSNYTTCVDLEDFEMRNQVNFIYKAGYMISLAALLLSLFIFFYFRSLTCTRIQIHKNFFISLAINNLLWLIWYEAVLDN
HTVIYENGVGCQILHVILQYFLVTTYFWGFCEGLYLYTLLVVTFLTESKVMCLYLIGWGVPALIVSAYASLRISTNKDTDYCW

IQESIYRWTLIIPVGLSMIANLIFLITIVRVVLTKLHAAQKTTPSNSFKDKNASIRSKRSTVLSDTAFSERTKKAVRATLILIPLIGLQ
YVVMIVRPEQKTTWEYTYELTEAIVASSQGLCVALLFCFCNGEVTAAVRKKWRQCRLSKKRPWNSCSGVTSVSFSRSSFVQ
EMAGVVPPPHIDGLATGNCQL
>ClecCT/DHR3-CLEC013307
MIITQFMFTDGFGCPKEFDGWTCINATSPGAVIHVPCPYFIFGFDPKRLGHRTCLEDGTWFRHPDSNKTWTNYTTCVDTD
DLKMRTQVNLIYKAGYTFSLAALLLSIIIFFYFKSLTCTRIQIHKNLFISLAVNNLLWLVWYEAVVDNIPVLLSNGIGCQILHVLV
QYFMVATYLWMFCEGLYLHTLLVVTFVTESKVMPVLHLIGWGVPALLVTVYASLRVSTKDDTLHCWIHESLYSWSLSGPVC
LSMIANLVFLINIVRLLLTKLRLPRRSTISSQGLPSGRTKKAVRATLILIPLLGLQYIVTPFRPEQGTSWEYPYQVTSALVASCQGL
CVALLFCFFNGEVFLFNLNFLKEENSNFALVLGCSGHEEKVATVQAE
>ClecCRF/DHR1-CLEC025139
MCITKKNFQEIRCFKLMEEERDQKDVCERSWDGLLCWPPSRPGAAYLPCFEELHGIKYDTSQNATRWCHPNGTWANYSN
YTKCSDLGKPAIEFEDGVEVTTMIYSIGYGLSLVALCLAVYIFIYYKELRCLRNTIHTNLMCTYILADFMWILNYSLQISVQTDFV
PCVLLVILLHYFHLTNFFWMFVEGFYLYMLVVKTFTQENIKLRVYLAIGWGNFCLKMNINVLYTMFQIVMDPFQKGCHWM
MPNMTDWIYQTPAIIVLVINLMFLVMIMLVLITKLRSATNVETQQYRKAAKALLVLIPLLGITYILFIVGPSKGPYYANLYEYVR
SILISTQGLMVALFYCFLNTEVQNTVRHHLIRWKEARDIGVRRYTYYCKTRSPNSDIESVR
>ClecCRF/DHR2-CLEC000652
MPTYDEPAENCPWSWDGALCWPSFPPGKNASLPCFEELKGIKYDTSQNASRYCFPNGTWSTYTNYNACQATVPENMVP
EMPWVVGASTIYCIGYSISLVALCVAVWIFIYFKDLRCLRNTIHTNLMCTYILADFLWILNFALQILVPSDFICVVFVLLLKYFILT
NYFWMFVEGLYLYMLVVATFTRENIKLRVYLGIGWGKCFFIGIISTASEELDRNSCSWLTPHWSDWIDQSAALLVLGVNLIF
LFMIMLVLITKLRSSNNVETQQYRKASKALLVLIPLLGVTYIFFLVGPKTDIYEYIRAILLSTQGFSVALFYCFLNTEVQNTFRHHF
LRWKESRDIGHRRYTYSKDWSPNTRTESIR
>HhalCRF/DHR2a-XP_024216523.1
MASNLEVLKNLTTESQFELQSQECATLWSRKEPSGWCRAKFDGALCWGPTAPSHLSTQPCKEVINDVYYDTSKNATKFCD
SNGDWNLTNYDNCQVRDFMVPDAEMTEMSTVIYSVGYTLSLITLSLALFIFIYCKEMRCLRNTIHTNLMFTYVLADFMWIL
SLTVQVSIHTDSISCLVLFTLLHYFLLTNFFWMFVEGLYLYMLVVETFTRENIGLRAYLAIGWGSPIPVIIVWIIARTTVQDPNA
VTIMGVPETMNQCTMMYTSMTDWIYIIPVLVVLVVNLLFLCMIMWVLITKLRSANNAETEQYRKGSKALLVLIPLLGITYILL
IAGPNASVYHNIRALLLSTQGLTVGLLYCFLNTEVQNTLRHRWQRWREERSLPHRTYTKDSSPNTRTESIRLYSKHEVVPYRK
RESTGSETTTMTLVHGNSKISNGPRSSFLQPPSEPV
>HhalCRF/DHR2b-XP_024216523.1
MEDQELTSNFSYHIEREECLQRWSYHYQEGWCPAVFDGALCWGPTGPAVLASQPCREEIHGVLYDTSKNATKYCHESGI
WDNRTDYDQCQERADILATLTADDIEMTTVIYALGYALSLIALSVALFIFIYFKEMRCLRNTIHTNLMLTYVLADFMWILSLTV
SMHTDSVSCLILFTLLHYFILTNFFWMFVEGLYLYMLVVETFTRENIGLRAYLAIGWGLPVAVIIVWVIARFNASDMPEVPPG
TKQCTWMNQSWSDWIYQVPAILVLAINLLFLVRIMWVLITKLRSANNAETEQYRKGSKALLVLIPLLGITYILFIAGPQSAVYS
NIRALLLSTQGLSVGLLYCFLNTEVQNTLRHRWLRWKEERSLATRAYTKDMSPNTRTESIRLYSRHEITPYRKRESTGSESTT
MTLVHSSRLSNGPRSSFLQPPSEPV
>ApisCRF/DHR2-XP_001944842.2
MNITSTNNTCLPEKVLNPGWCPSHWDLLLCWRTSKPGAVVYQACFDELNGMRYDTSQNASRRCKLNGVWENSSDYMN
CRPLDTNNPLYPDPSIVYTSYFYYGGYTISLVALVAAVSIFVYFKDLRCLRNTIHTNLMCTYILSDFTWILTSTLQEWLSASNNA
CVLFTFSLHYFVLTNFFWMFVEGLYLYILVVETFTRENIKLRVYMFIGWGFPLVIMIVWGVSKIITPIEVEERSDSCSWMTPHP
VHDWIYQGPAIIVLIVNLVFLSKIMWVLITKLRSANSAETQQYRKASKALLVLIPLLGVTYILTMVGPTESGTYANYYSYGRATL
LSMQGFMVAIFYCFVNSEVKNTFKHHFIRWNDARNLRTGGSRRFTYSKDWSPNTRSDSVRLARPSIIADTVMNKGKRAST
VSSSTTMTFIVSNGSGGVLLSNACQNRMLNAPPENLV
>ApisCRF/DHR1-XP_008183744.1
MDYFNETWSPNEEAALKCRSWYEWDDRWINGCNASSDSLICWPPTPAGVIVYQPCFQELRGILYDTTKNASRICYENNT
WGLTDFNDCTILGEPKAVTYDEEGTVDTIIYLYIAGHCLSLIMTSLAIFVFCRFKELKCLRNKIHSNLMASYLLAGIMWILNYTN
LTDTGTFKCALLVLPLYYFTMTNYFWAFIEGMYLFILVVDTFFTDRVRLRTYMAIGWGIPLIIIPTWCVTRLLVPTKNDLDLYN
QITYERYCPLMASYVDDWIYQSPIVIVLLINSIFLVKIMRVLITKIRSTKSAETHNYKKATKALLVLIPLLGITFCLDMINPSSTGLL
VNIYKFSKVVIISTQGFTVSLLYCFFNNEVQNTLKYHITRWQTKRKFLASRKKYGRSWLSVKQNNIICDHQLDPKELMPWIPA
NNVRSSSCVSNGTTTSALSNMRLPPDTTIEIPPEQQPLEDSKYVGAKP
>AmelCRF/DHR-LOC413829
MLRIQRELRCIRNNIHTNLMFTYILADFMWILNNVMQVSMQPDVPTCVAFLSLFHYFQLTNYFWMFVEGLYLYLLVVKTFT
GDNIKLRLCLIIGWGIPVLFITMWCIAKSLDQNATSQDIALGKHCPWMVSHAYDWFYQAPAILVLCVNVVFLFMIMWVLIT
KLWSATNVETQQYRKASKALLVLIPLLGVTYVLVLTGPTEGQVANAFSYARAVLLSSQGLFVALFYCFLNTEVRNTVKSHIER
WTTARDLDTERRLYCPPSNPYRKRDSSVSETTTTTVVGLNSTTTFLDKSKKAISEDMNE

>TcasCRF/DHR1-TC007104
MDQGESEVVYARHEDIVRILNRLNETQAELACELKKTLSPPSNGCAVDFDTVLCWPQTAPNSLAVLPCFDQLNGIKYDTRE
NATRLCFANGTWDQYSNYTSCKELSPLEVPEVELTTTIYFIGYTVSLVALLFAVYIFWKFKDLRCLRNTIHMNLMCSYILADFM
WIFVYSLQVPLQTNKAFCIFLIILLHYFHLTNFFWMFVEGLYLYILVVKTFTGENIKPRIYAVIGWGGPILFVLVWGIAKSFTLPL
EDQQAGEMFRSCPWTPHPFDWIYQGPAIAVLIINVIFLCIIMWVLITKLRSANNVETQQYRKAAKALLVLIPLLGVTYILVIVG
PTEGISRRIYDSIRAILLSTQGFTVALFYCFLNAEVKNTVRHHYNSWHTRRTLGSRRTRYSSSKDWSSQARDSMRYGSKRAKL
STLKY
>TcasCRF/DHR2-TC012799
MSWSEPLPQEPEPVDADLWPSSENVLNETEDIKIRLNITLQHCTSLYTNRTTALAETHPDGFCPVTTDGLLCWPPTPINETTY
VKCFAELMNIRYDDTQNATRVCLANGTWTKADYSKCTEIILIPDVETQATIYFVGYVLSLITLSIALGIFTYFKELRCLRNRIHM
NLMWSYMLMYIMWILTLTVLGSKGGTGASIACIFVITLLHYFHISTFFWMFVEGLYLYILVVETLTRENYKLRVYVCIGWGLP
MIFILVWVIVKSFIPAAGDPATCTWFNSHDVDWIFQGPTMLVLLLNLAFLLAIMWVLITKLRSANTVETQQYHKAAKALLVL
MPLLGITYVITIYAPTPDKKSEIIFECVRAVLLSTQPLNEHPPICQSYSVAITGLHCRSILLLLKHGGAEHRPPPLRNVENTTISGPI
QTTFGESQQGLVPQVTHGEYTVREKIFNDSLGTITFGSI
>AgamCRF/DHR1-AGAP005464
MANETMPTTPTAAAIAADDGSGGSPFGEGNVTGISVENPSLVEGLLALAMNASGAERCRLQQQAEELLQDVACPSFFDM
VSCWPRTPPGTLAVLPCFAELKGVQYDSSQNATRFCNVDGTWDNYTDYDRCEHLEQPPPLPSFEPEIELPTLIYFVGYSISLA
ALVLAVAVLVYFKDLRCLRNTIHVNLFLTYIMSSSLWILILSLQITVKLEVAGCIFLVTLFHYFSTTNFFWMLVEGLYLYMLVVQ
TFSGDTLRFRKYAIIGWGGPLIFVGAWAIAKPFFGSVSNLEHPSKLEIECSWMRESHIDWIIQGPSCAVLVINLIFLLRIMWVLI
TKLRSANTVETRQYRKASKALLVLIPLLGITYLIVIYGPVEGVGSHIFAITRAILLSTQGFVVSLLYCFLNSEVRQTLRHHFYRWR
DERNILSGKVNNHHRRPTFSKDNSPRSRTESTRLVLVKECASSVPAKHVLYCRLQVVVAFIHVCLFVFVFWCVCVSLFVFCLY
TLFAPCCHSSLIRSHSTWHTGSLQRFSLLFLNYSKVC
>AgamCRF/DHR2-AGAP005465
MQSCLLSATLQPKPQVKPRTAVRYQTNRDQTIYVPQKFCPTRLWTVQMTGGIISNSIKPSGQRTTPTTLIRLQMDKIQPHA
SGKRLVSVADYDELSSFPPELLGAFENSTEFECLLRAHQDAQSETPSGGCPVDFDRILCWPKTDPGTWAVLPCFEEFKGVHY
DVTQNATRYCHPNGRWDNYSHYAACHHVNEPPPDIVEISSIIYYTGYILSLVALSLAVIVFVYFKDLRCLRNTIHANLFITYILSA
LLWIIILTLQLSSGSSTGMTSCVIFVTLLHYFTLTNFFWMLVEGLYLYMLVVETFSGDNLRFNMYAAIGWGKCSRASLEIECS
WMRESVVDWIFQGPVCAVLIINLVFLIRIMWVLITKLRSANTVETRQYRKASKALLVLIPLLGITYLVVLAAPAEGVVSDIFAIA
RALLLSTQGLSVSLFYCFLNSEVRLALRHRLERWRDERNIRLGQVRQRSRRMEKGPSKPESEPESIHHATESLKGFILPFSSAG
TVPLTYNKRESCASSATTTTLLGPHSYPSAMRGSNGALHMQSIVPRAISPLMQVRRWGEGRKSVTNYYPLRGAGN
>AaegCRF/DHR1-AAEL008292
MNDSSSASVESLLALAGAGGDFAGSGSGDGFPPMILNETLRDLCHQQNDTLFLELSCPPLFDSISCWPRTPPATTAVLPCFS
EFKGVQYDATQNATRYCNIDGTWNSFANYDACKHLELPSSDPTLESFIELPIVIYFVGYTISLLALFCAVTVLVYFKELRCLRNTI
HVNLFVTYIMSSSLWIIILSQQLAAKQGLVDCIFLVTLFHYFSTTNFFWMLVEGLYLYMLVVQTFSGDYLRFWKYSIIGWGGP
LIFVGAWAIAKSFYPYDLMPEHPNKLEIECSWMRESHIDWIIQGPSCAVLVINLIFLLRIMWVLITKLRSANTVETRQYRKASK
ALLVLIPLLGITYLVVIYGPHEGVGSRIFAVTRAVLLSTQGLVVSLLYCFLNYEVRGTLRLHYYRWRDERNIRLGIISKHRRPTISG
TESIRTLEYSVVIMKSLAPPSSRHHHHHNHHNNLGGSN
>AaegCRF/DHR2-AAEL019757
MTEPPPDIVEVTSIIYYSGYIVSLVALSLAVIVFVYFKDLRCLRNTIHANLFITYILSALMWIIILTLQLSGSQSGIASCVILVTLLHYF
TLTNFFWMLVEGLYLYMLVVETFSGDNLRFNMYAAIGWGGPGVFVIMWVIAKGITISGLDQATTLEIDCSWMRESTVDW
IFQGPVCAVLIINLVFLIRIMWVLITKLRSANTVETRQYRKASKALLVLIPLLGITYLVVLAAPSEGVVSDIFAVARALLLSTQGFS
VSLFYCFLNSEVRLALRHRLERWRDERNIRIGQVRQRSRRFTNSGLSKECSPRSRTESFRPLTYSKRESCASSATTTTLLGPPVG
AHGAYPSTMRGSNGALHLHAMAPRAISPLMQGLDENSV
>DmelCRF/DHR1-CG8422
MSDHNHIDSVNASGSDPLLDLHNLDGIGESVELQCLVQEHIEASTYGNDSGHCLTQFDSILCWPRTARGTLAVLQCMDEL
QGIHYDSSKNATRFCHANGTWEKYTNYDACAHLPAPESVPEFEVIVELPTIIYYIGYTLSLVSLSLALIVFAYFKELRCLRNTIHA
NLFFTYIMSALFWILLLSVQISIRSGVGSCIALITLFHFFTLTNFFWMLVEGLYLYMLVVKTFSGDNLRFNIYASIGWGGPALFV
VTWAVAKSLTVTYSTPEKYEINCPWMQETHVDWIYQGPVCAVLIINLTFLLRIMWVLITKLRSANTVETRQYRKAAKALLVLI
PLFGITYLVVLAGPSESGLMGHMFAVLRAVLLSTQGFSVSLFYCFLNSEVRNALRHHISTWRDTRTIQLNQNRRYTTKSFSKG
GGSPRAESMRPLTSYYGRGKRESCVSSATTTTLVGQHAPLSLHRGSNNALHTMPTLAANAMSSGSTLSVMPRAISPLMRQ
GLEENSV
>DmelCRF/DHR2-CG12370
MADDDLRALVDSLDDASQEDLAKVIANFSVDMLQRASALIGAQQGSSGGQLQNRTLQCQQQQQREEEQASLEALASGG
KRILQCPSSFDSVLCWPRTNAGSLAVLPCFEEFKGVHYDTTDNATRFCFPNGTWDHYSDYDRCHQNSGSIPVVPDFSPNVE

LPAIIYAGGYFLSFATLVVALIIFLSFKDLRCLRNTIHANLFLTYITSALLWILTLFLQVITTESSQAGCITLVIMFQYFYLTNFFWM
FVEGLYLYTLVVQTFSSDNISFIIYALIGWGCPAVCILVWSIAKAFAPHLENEHFNGLEIDCAWMRESHIDWIFKVPASLALLV
NLVFLIRIMWVLITKLRSAHTLETRQYYKASKALLVLIPLFGITYLLVLTGPEQGISRNLFEAIRAFLISTQGFFVALFYCFLNSEVR
QTLRHGFTRWRESRNIHRNSSIKNRRHRASKDYSLRSRTESLRLTSTSPIPTGHYE
>DmelGlutR
MKQKNNNGTILVVVMVLSWSRVVDLKSPSNTHTQDSVSVSLPGDIILGGLFPVHEKGEGAPCGPKVYNRGVQRLEAMLY
AIDRVNNDPNILPGITIGVHILDTCSRDTYALNQSLQFVRASLNNLDTSGYECADGSSPQLRKNASSGPVFGVIGGSYSSVSL
QVANLLRLFHIPQVSPASTAKTLSDKTRFDLFARTVPPDTFQSVALVDILKNFNWSYVSTIHSEGSYGEYGIEALHKEATERNV
CIAVAEKVPSAADDKVFDSIISKLQKKPNARGVVLFTRAEDARRILQAAKRANLSQPFHWIASDGWGKQQKLLEGLEDIAEG
AITVELQSEIIADFDRYMMQLTPETNQRNPWFAEYWEDTFNCVLTSLSVKPDTSNSANSTDNKIGVKAKTECDDSYRLSEK
VGYEQESKTQFVVDAVYAFAYALHNLHNDRCNTQSDQTTETRKHLQSESVWYRKISTDTKSQACPDMANYDGKEFYNNY
LLNVSFIDLAGSEVKFDRQGDGLARYDILNYQRQENSSGYQYKVIGKWFNGLQLNSETVVWNKETEQPTSACSLPCEVGMI
KKQQGDTCCWICDSCESFEYVYDEFTCKDCGPGLWPYADKLSCYALDIQYMKWNSLFALIPMAIAIFGIALTSIVIVLFAKNH
DTPLVRASGRELSYTLLFGILVCYCNTFALIAKPTIGSCVLQRFGIGVGFSIIYSALLTKTNRISRIFHSASKSAQRLKYISPQSQVVI
TTSLIAIQVLITMIWMVVEPPGTRFYYPDRREVILKCKIQDMSFLFSQLYNMILITICTIYAIKTRKIPENFNESKFIGFTMYTTCII
WLAFVPIYFGTGNSYEVQTTTLCISISLSASVALVCLYSPKVYILVFHPDKNVRKLTMNSTVYRRSAAAVAQGAPTSSGYSRTH
APGTSALTGGAVGTNASSSTLPTQNSPHLDEASAQTNVAHKTNGEFLPEVGERVEPICHIVNK

=============================
Nuclear receptor sequences
>DmelHr51
MATGRSLLFRVPWYVCLCVCAESAEPGVYWRLRLRLGLPTLAGPHTNTLTLTARTSSCRSIKKERIKASQQANAPPELPLKVS
VDVNIIIAAHSQRRRIGLVRFHQRESEDRPLAVASPRLQINMEPTAMNPKKLHSPQRHCYTPPPAPMHGQAPPPTSTGVAP
PTQPPPPHPAAPNVPNGRLLSWNHSAAAAAAAAAAQAAANSMNHSSAAEGSSMTRIKGQNLGLICVVCGDTSSGKHYG
ILACNGCSGFFKRSVRRKLIYRCQAGTGRCVVDKAHRNQCQACRLKKCLQMGMNKDDDSIDVTNDNEEPHAVSRSDSSFI
MPQFMSPNLYTHQHETVYETSARLLFMAVKWAKNLPSFARLSFRDQVILLEESWSELFLLNAIQWCIPLDPTGCALFSVAE
HCNNLENNANGDTCITKEELAADVRTLHEIFCKYKAVLVDPAEFACLKAIVLFRPETRGLKDPAQIENLQDQAHHTKTQFTA
QIARFGRLLLMLPLLRMISSHKIESIYFQRTIGNTPMEKVLCDMYKN
>Dmeltll
MQSSEGSPDMMDQKYNSVRLSPAASSRILYHVPCKVCRDHSSGKHYGIYACDGCAGFFKRSIRRSRQYVCKSQKQGLCVV
DKTHRNQCRACRLRKCFEVGMNKDAVQHERGPRNSTLRRHMAMYKDAMMGAGEMPQIPAEILMNTAALTGFPGVP
MPMPGLPQRAGHHPAHMAAFQPPPSAAAVLDLSVPRVPHHPVHQGHHGFFSPTAAYMNALATRALPPTPPLMAAEHI
KETAAEHLFKNVNWIKSVRAFTELPMPDQLLLLEESWKEFFILAMAQYLMPMNFAQLLFVYESENANREIMGMVTREVHA
FQEVLNQLCHLNIDSTEYECLRAISLFRKSPPSASSTEDLANSSILTGSGSPNSSASAESRGLLESGKVAAMHNDARSALHNYI
QRTHPSQPMRFQTLLGVVQLMHKVSSFTIEELFFRKTIGDITIVRLISDMYSQRKI
>DmelDsf
MGTAGDRLLDIPCKVCGDRSSGKHYGIYSCDGCSGFFKRSIHRNRIYTCKATGDLKGRCPVDKTHRNQCRACRLAKCFQSA
MNKDAVQHERGPRKPKLHPQLHHHHHHAAAAAAAAHHAAAAHHHHHHHHHAHAAAAHHAAVAAAAASGLHHHHH
AMPVSLVTNVSASFNYTQHISTHPPAPAAPPSGFHLTASGAQQGPAPPAGHLHHGGAGHQHATAFHHPGHGHALPAPH
GGVISNPGGNSSAISGSGPGSTLPFPSHLLHHNLIAEAASKLPGITATAVAAVVSSTSTPYASAAQASSPSSNNHNYSSPSPSN
SIQSISSIGSRSGGGEEGLSLGSESPRVNVETETPSPSNSPPLSAGSISPAPTLTTSSGSPQHRQMSRHSLSEATTPPSHASLMI
CASNNNNNNNNNNNNNNNGEHKQSSYTSGSPTPTTPTPPPPRSGVGSTCNTASSSSGFLELLLSPDKCQELIQYQVQHNTL
LFPQQLLDSRLLSWEMLQETTARLLFMAVRWVKCLMPFQTLSKNDQHLLLQESWKELFLLNLAQWTIPLDLTPILESPLIRE
RVLQDEATQTEMKTIQEILCRFRQITPDGSEVGCMKAIALFAPETAGLCDVQPVEMLQDQAQCILSDHVRLRYPRQATRFG
RLLLLLPSLRTIRAATIEALFFKETIGNVPIARLLRDMYTMEPAQVDK
>DmHr78
MDGVKVETFIKSEENRAMPLIGGGSASGGTPLPGGGVGMGAGASATLSVELCLVCGDRASGRHYGAISCEGCKGFFKRSIR
KQLGYQCRGAMNCEVTKHHRNRCQFCRLQKCLASGMRSDSVQHERKPIVDRKEGIIAAAGGSSTSGGGNGSSTYLSGKSG
YQQGRGKGHSVKAESAATPPVHSAPATAFNLNENIFPMGLNFAELTQTLMFATQQQQQQQQQHQQSGSYSPDIPKADP
EDDEDDSMDNSSTLCLQLLANSASNNNSQHLNFNAGEAPTALPTTSTMGLIQSSLDMRVIHKGLQILQPIQNQLERNGNL
SVKPECDSEAEDSGTEDAVDAELEHMELDFECGGNRSGGSDFAINEAVFEQDLLTDVQCAFHVQPPTLVHSYLNIHYVCET
GSRIIFLTIHTLRKVPVFEQLEAHTQVKLLRGVWPALMAIALAQCQGQLSVPTIIGQFIQSTRQLADIDKIEPLKISKMANLTRT
LHDFVQELQSLDVTDMEFGLLRLILLFNPTLLQQRKERSLRGYVRRVQLYALSSLRRQGGIGGGEERFNVLVARLLPLSSLDAE
AMEELFFANLVGQMQMDALIPFILMTSNTSGL

>DmmelUSP
MDNCDQDASFRLSHIKEEVKPDISQLNDSNNSSFSPKAESPVPFMQAMSMVHVLPGSNSASSNNNSAGDAQMAQAPNS
AGGSAAAAVQQQYPPNHPLSGSKHLCSICGDRASGKHYGVYSCEGCKGFFKRTVRKDLTYACRENRNCIIDKRQRNRCQY
CRYQKCLTCGMKREAVQEERQRGARNAAGRLSASGGGSSGPGSVGGSSSQGGGGGGGVSGGMGSGNGSDDFMTNSV
SRDFSIERIIEAEQRAETQCGDRALTFLRVGPYSTVQPDYKGAVSALCQVVNKQLFQMVEYARMMPHFAQVPLDDQVILLK
AAWIELLIANVAWCSIVSLDDGGAGGGGGGLGHDGSFERRSPGLQPQQLFLNQSFSYHRNSAIKAGVSAIFDRILSELSVK
MKRLNLDRRELSCLKAIILYNPDIRGIKSRAEIEMCREKVYACLDEHCRLEHPGDDGRFAQLLLRLPALRSISLKCQDHLFLFRIT
SDRPLEELFLEQLEAPPPPGLAMKLE
>DmelKnirps
MVFNLMNQTCKVCGEPAAGFHFGAFTCEGCKSFFGRSYNNISTISECKNEGKCIIDKKNRTTCKACRLRKCYNVGMSKGGS
RYGRRSNWFKIHCLLQEHEQAAAAAGKAPPLAGGVSVGGAPSASSPVGSPHTPGFGDMAAHLHHHHQQQQQQVPR
HPHMPLLGYPSYLSDPSAALPFFSMMGGVPHQSPFQLPPHLLFPGYHASAAAAAASAADAAYRQEMYKHRQSVDSVESQ
NRFSPASQPPVVQPTSSARQSPIDVCLEEDVHSVHSHQSSASLLHPIAIRATPTTPTSSSPLSFAAKMQSLSPVSVCSIGGETT
SVVPVHPPTVSAQEGPMDLSMKTSRSSVHSFNDSGSEDQEVEVAPRRKFYQLEAECLTTSSSSSSHSAAHSPNTTTAHAEV
KRQKLGGAEATHFGGFAVAHNAASAMRGIFVCV
>DmelKnrl
MMNQDNPYAMNQTCKVCGEPAAGFHFGAFTCEGCKSFFGRSYNNLSSISDCKNNGECIINKKNRTACKACRLKKCLMVG
MSKSGSRYGRRSNWFKIHCLLQEQQQQAVAAMAAHHNSQQAGGGSSGGSGGGQGMPNGVKGMSGVPPPAAAAAA
LGMLGHPGGYPGLYAVANAGGSSRSKEELMMLGLDGSVEYGSHKHPVVASPSVSSPDSHNSDSSVEVSSVRGNPLLHLG
GKSNSGGSSSGADGSHSGGGGGGGGGVTPGRPPQMRKDLSPFLPLPFPGLASMPVMPPPAFLPPSHLLFPGYHPALYSH
HQGLLKPTPEQQQAAVAAAAVQHLFNSSGAGQRFAPGTSPFANHQQHHKEEDQPAPARSPSTHANNNHLLTNGGAAD
ELTKRFYLDAVLKSQQQSPPPTTKLPPHSKQDYSISALVTPNSESGRERVKSRQNEEDDEARADGIIDGAEHDDEEEDLVVS
MTPPHSPAQQEERTPAGEDPRPSPGQDNPIDLSMKTTGSSLSSKSSSPEIEPETEISSDVEKNDTDDDDEDLKVTPEEEISVR
ETADPEIEEDHSSTTETAKTSIENTHNNNNSISNNNNNNNNNNNNSILSDSEASETIKRKLDELIEASSENGKRLRLEAPVKVAT
SNALDLTTKV
>DmelEg
MNQLCKVCGEPAAGFHFGAFTCEGCKSFFGRTYNNIAAIAGCKHNGDCVINKKNRTACKACRLRKCLLVGMSKSGSRYGR
RSNWFKIHCLLQEQQTTSGLGGGSSVGSGSGGGVSSASLEQLARLQQASNQARQTYQDKTNPCIKSATATTSPRIEGAAV
GTGIGGGASPSFLQAAKLHHQRQLKLDSRLSNTPSDSGASSAGDPNEDGVTSVLGGQIATPSSTNATSLPKLDLRHPNFPAT
SEPDADMQRQRHQELLEIFRSHSEPLYSSFAPFSHLPPVLLAAGVPQLPIFKDQFKAELLFPTTSSPELEEPIDLSFRSRADPAS
PMAHNSNSPSLSEPAAASHCLGESTNFVRKSTPLDLTLVRSQTLTG
>DmelEip75B
MEAVQAAAAATSSGGSSGSVPGSGSGSASKLIKTEPIDFEMLHLEENERQQDIEREPSSSNSNSNSNSLTPQRYTHVQVQT
VPPRQPTGLTTPGGTQKVILTPRVEYVQQRATSSTGGGMKHVYSQQQGTAASRSAPPETTALLTTTSGTPQIIITRTLPSNQ
HLSRRHSASPSALHHYQQQQPQRQQSPPPLHHQQQQQQQHVRVIRDGRLYDEATVVVAARRHSVSPPPLHHHSRSAPV
SPVIARRGGAAAYMDQQYQQRQTPPLAPPPPPPPPPPPPPPPPQQQQQQYISTGVPPPTAAARKFVVSTSTRHVNVIASN
HFQQQQQQHQAQQHQQQHQQHQQHQQHVIASVSSSSSSSAIGSGGSSSSHIFRTPVVSSSSSSNMHHQQQQQQQQS
SLGNSVMRPPPPPPPPPKVKHASSSSSGNSSSSNTNNSSSSSNGEEPSSSIPDLEFDGTTVLCRVCGDKASGFHYGVHSCEGC
KGFFRRSIQQKIQYRPCTKNQQCSILRINRNRCQYCRLKKCIAVGMSRDAVRFGRVPKREKARILAAMQQSTQNRGQQRAL
ATELDDQPRLLAAVLRAHLETCEFTKEKVSAMRQRARDCPSYSMPTLLACPLNPAPELQSEQEFSQRFAHVIRGVIDFAGMI
PGFQLLTQDDKFTLLKAGLFDALFVRLICMFDSSINSIICLNGQVMRRDAIQNGANARFLVDSTFNFAERMNSMNLTDAEIG
LFCAIVLITPDRPGLRNLELIEKMYSRLKGCLQYIVAQNRPDQPEFLAKLLETMPDLRTLSTLHTEKLVVFRTEHKELLRQQM
WSMEDGNNSDGQQNKSPSGSWADAMDVEAAKSPLGSVSSTESADLDYGSPSSSQPQGVSLPSPPQQQPSALASSAPLL
AATLSGGCPLRNRANSGSSGDSGAAEMDIVGSHAHLTQNGLTITPIVRHQQQQQQQQQIGILNNAHSRNLNGGHAMCQ
QQQQHPQLHHHLTAGAARYRKLDSPTDSGIESGNEKNECKAVSSGGSSSCSSPRSSVDDALDCSDAAANHNQVVQHPQL
SVVSVSPVRSPQPSTSSHLKRQIVEDMPVLKRVLQAPPLYDTNSLMDEAYKPHKKFRALRHREFETAEADASSSTSGSNSLS
AGSPRQSPVPNSVATPPPSAASAAAGNPAQSQLHMHLTRSSPKASMASSHSVLAKSLMAEPRMTPEQMKRSDIIQNYLK
RENSTAASSTTNGVGNRSPSSSSTPPPSAVQNQQRWGSSSVITTTCQQRQQSVSPHSNGSSSSSSSSSSSSSSSSSTSSNCSS
SSASSCQYFQSPHSTSNGTSAPASSSSGSNSATPLLELQVDIADSAQPLNLSKKSPTPPPSKLHALVAAANAVQRYPTLSADV
TVTASNGGPPSAAASPAPSSSPPASVGSPNPGLSAAVHKVMLEA
>DmelEip78C
MDVYQIELEEQAQIRSKLLVETCVKHSSSEQQQLQVKQEDLIKDFTRDEEEQPSEEEAEEEDNEEDEEEEGEEEEEDEDEEAL
LPVVNFNANSDFNLHFFDTPEDSSTQGAYSEANSLESEQEEEKQTQQHQQQKQHHRDLEDCLSAIEADPLQLLHCDDFYR
TSALAESVAASLSPQQQQQRQHTHQQQQQQQQQQHPGQQQHQLNCTLSNGGGALYTISSVHQFGPASNHNTSSSSP

SSSAAHSSPDSGCSSASSSGSSRSCGSSSASSSSSAVSSTISSGRSSNNSVVNPAATSSSVAHLNKEQQQQPLPTTQLQQQQ
QHQQQLQHPQQQQSFGLADSSSSNGSSNNNNGVSSKSFVPCKVCGDKASGYHYGVTSCEGCKGFFRRSIQKQIEYRCLRD
GKCLVIRLNRNRCQYCRFKKCLSAGMSRDSVRYGRVPKRSRELNGAAASSAAAGAPASLNVDDSTSSTLHPSHLQQQQQQ
HLLQQQQQQQHQPQLQQHHQLQQQPHVSGVRVKTPSTPQTPQMCSIASSPSELGGCNSANNNNNNNNNNSSSGNASG
GSGVSVGVVVVGGHQQLVGGSMVGMAGMGTDAHQVGMCHDGLAGTANELTVYDVIMCVSQAHRLNCSYTEELTREL
MRRPVTVPQNGIASTVAESLEFQKIWLWQQFSARVTPGVQRIVEFAKRVPGFCDFTQDDQLILIKLGFFEVWLTHVARLIN
EATLTLDDGAYLTRQQLEILYDSDFVNALLNFANTLNAYGLSDTEIGLFSAMVLLASDRAGLSEPKVIGRARELVAEALRVQIL
RSRAGSPQALQLMPALEAKIPELRSLGAKHFSHLDWLRMNWTKLRLPPLFAEIFDIPKADDEL
>DmelHr46
MASLLGSAPPAQILANQPIIVKIEPTQSFHIVDEGDTRVLSLPLSDADKLGASWIDLKDIAGLQAGGGATLLDVCFEQANEDG
TIIATVQPDLENELEAELKAEGEPEDETEPEPPAPKRLATTRPAQSRPQQQQQQQQQQVKFLSDPPALARSSSFSSLSSFSSIS
NISSVCKNMASNTSQEGSLKRTKERTPPPMPPLTTHKPAATTTATSATSAAAATSAATASATATSARENSREHSSSSSGSNG
AMTAQIEIIPCKVCGDKSSGVHYGVITCEGCKGFFRRSQSSVVNYQCPRNKQCVVDRVNRNRCQYCRLQKCLKLGMSRDA
VKFGRMSKKQREKVEDEVRFHRAQMRAQSDAAPDSSVYDTQTPSSSDQLHHNNYNSGGYSNNEVGYGSPYGYSASVTP
QQTMQYDISADYVDSTTYEPRSTIIDPEFISHADGDINDVLIKTLAEAHANTNTKLEAVHDMFRKQPDVSRILYYKNLGQEEL
WLDCAEKLTQMIQNIIEFAKLIPGFMRLSQDDQILLLKTGSFELAIVRMSRLLDLSQNAVLYGDVMLPQEAFYTSDSEEMRL
VSRIFQTAKSIAELKLTETELALYQSLVLLWPERNGVRGNTEIQRLFNLSMNAIRQELETNHAPLKGDVTVLDTLLNNIPNFRD
ISILHMESLSKFKLQHPNVVFPALYKELFSIDSQQDLT
>DmelEcR
MLTTSGQQQSKQKLSTLPSHILLQQQLAASAGPSSSVSLSPSSSAALTLHVASANGGARETTSAAAVKDKLRPTPTAIKIEPM
PDVISVGTVAGGSSVATVVAPAATTTSNKPNSTAAPSTSAAAANGHLVLVPNKRPRLDVTEDWMSTPSPGSVPSSAPPLSP
SPGSQNHSYNMSNGYASPMSAGSYDPYSPTGKTGRDDLSPSSSLNGYSANESCDAKKSKKGPAPRVQEELCLVCGDRASG
YHYNALTCEGCKGFFRRSVTKSAVYCCKFGRACEMDMYMRRKCQECRLKKCLAVGMRPECVVPENQCAMKRREKKAQK
EKDKMTTSPSSQHGGNGSLASGGGQDFVKKEILDLMTCEPPQHATIPLLPDEILAKCQARNIPSLTYNQLAVIYKLIWYQDG
YEQPSEEDLRRIMSQPDENESQTDVSFRHITEITILTVQLIVEFAKGLPAFTKIPQEDQITLLKACSSEVMMLRMARRYDHSSD
SIFFANNRSYTRDSYKMAGMADNIEDLLHFCRQMFSMKVDNVEYALLTAIVIFSDRPGLEKAQLVEAIQSYYIDTLRIYILNR
HCGDSMSLVFYAKLLSILTELRTLGNQNAEMCFSLKLKNRKLPKFLEEIWDVHAIPPSVQSHLQITQEENERLERAERMRASV
GGAITAGIDCDSASTSAAAAAAQHQPQPQPQPQPSSLTQNDSQHQTQPQLQPQLPPQLQGQLQPQLQPQLQTQLQPQI
QPQPQLLPVSAPVPASVTAPGSLSAVSTSSEYMGGSAAIGPITPATTSSITAAVTASSTTSAVPMGNGVGVGVGVGGNVS
MYANAQTAMALMGVALHSHQEQLIGGVAVKSEHSTTA
>DmelHr96
MSPPKNCAVCGDKALGYNFNAVTCESCKAFFRRNALAKKQFTCPFNQNCDITVVTRRFCQKCRLRKCLDIGMKSENIMSEE
DKLIKRRKIETNRAKRRLMENGTDACDADGGEERDHKAPADSSSSNLDHYSGSQDSQSCGSADSGANGCSGRQASSPGT
QVNPLQMTAEKIVDQIVSDPDRASQAINRLMRTQKEAISVMEKVISSQKDALRLVSHLIDYPGDALKIISKFMNSPFNALTVF
TKFMSSPTDGVEIISKIVDSPADVVEFMQNLMHSPEDAIDIMNKFMNTPAEALRILNRILSGGGANAAQQTADRKPLLDKE
PAVKPAAPAERADTVIQSMLGNSPPISPHDAAVDLQYHSPGVGEQPSTSSSHPLPYIANSPDFDLKTFMQTNYNDEPSLDS
DFSINSIESVLSEVIRIEYQAFNSIQQAASRVKEEMSYGTQSTYGGCNSAANNSQPHLQQPICAPSTQQLDRELNEAEQMKL
RELRLASEALYDPVDEDLSALMMGDDRIKPDDTRHNPKLLQLINLTAVAIKRLIKMAKKITAFRDMCQEDQVALLKGGCTE
MMIMRSVMIYDDDRAAWKVPHTKENMGNIRTDLLKFAEGNIYEEHQKFITTFDEKWRMDENIILIMCAIVLFTSARSRVIH
KDVIRLEQNSYYYLLRRYLESVYSGCEARNAFIKLIQKISDVERLNKFIINVYLNVNPSQVEPLLREIFDLKNH
>DmelHnf4
MMKHPQDLSVTDDQQLMKVNKVEKMEQELHDPESESHIMHADALASAYPAASQPHSPIGLALSPNGGGLGLSNSSNQS
SENFALCNGNGNAGSAGGGSASSGSNNNNSMFSPNNNLSGSGSGTNSSQQQLQQQQQQQSPTVCAICGDRATGKHYG
ASSCDGCKGFFRRSVRKNHQYTCRFARNCVVDKDKRNQCRYCRLRKCFKAGMKKEAVQNERDRISCRRTSNDDPDPGNG
LSVISLVKAENESRQSKAGAAMEPNINEDLSNKQFASINDVCESMKQQLLTLVEWAKQIPAFNELQLDDQVALLRAHAGEH
LLLGLSRRSMHLKDVLLLSNNCVITRHCPDPLVSPNLDISRIGARIIDELVTVMKDVGIDDTEFACIKALVFFDPNAKGLNEPH
RIKSLRHQILNNLEDYISDRQYESRGRFGEILLILPVLQSITWQMIEQIQFAKIFGVAHIDSLLQEMLLGGELADNPLPLSPPNQ
SNDYQSPTHTGNMEGGNQVNSSLDSLATSGGPGSHSLDLEVQHIQALIEANSADDSFRAYAASTAAAAAAAVSSSSSAPA
SVAPASISPPLNSPKSQHQHQQHATHQQQQESSYLDMPVKHYNGSRSGPLPTQHSPQRMHPYQRAVASPVEVSSGGGG
LGLRNPADITLNEYNRSEGSSAEELLRRTPLKIRAPEMLTAPAGYGTEPCRMTLKQEPETGY
>DmelHr83
MSNFSACAVCGDQSSGKHYGVSCCDGCSCFFKRSVRRGSSYACIALVGNCVVDKARRNWCPSCRFQRCLAVGMNAAAV
QEERGPRNQQVALYRTGRRQAPPSQAAPSPTPHSQALHFQILAQILVTCLRQAKANEQFALLDRCQQDAIFQVVWSEIFVL

RASHWSLDISAMIDGCGDEQLKRLICEAHQLRADVLELNFMESLILCRKELAINAEYAVILGSHSKAALISLARYTLQQSNYLR
FGQLLLGLRQLCLRRFDCALSCMFRSVVRDILKTL
>DmelSVP
MCASPSTAPGFFNPRPQSGAELSAFDIGLSRSMGLGVPPHSAWHEPPASLGGHLHAASAGPGTTTGSVATGGGGTTPSSV
ASQQSAVIKQDLSCPSLNQAGSGHHPGIKEDLSSSLPSANGGSAGGHHSGSGSGSGSGVNPGHGSDMLPLIKGHGQDML
TSIKGQPTGCGSTTPSSQANSSHSQSSNSGSQIDSKQNIECVVCGDKSSGKHYGQFTCEGCKSFFKRSVRRNLTYSCRGSRN
CPIDQHHRNQCQYCRLKKCLKMGMRREAVQRGRVPPTQPGLAGMHGQYQIANGDPMGIAGFNGHSYLSSYISLLLRAEP
YPTSRYGQCMQPNNIMGIDNICELAARLLFSAVEWAKNIPFFPELQVTDQVALLRLVWSELFVLNASQCSMPLHVAPLLAA
AGLHASPMAADRVVAFMDHIRIFQEQVEKLKALHVDSAEYSCLKAIVLFTTGKLLDILYKDVPALLTKVSALLGKGSTASNDD
VLAVVRDHLDELNRQEQESQAQQQAPLHLAAFMNCVAGVEAAVQQAEQAQVPTSSASASVSAPLVPSAGSAFSSCQAK
SAGSEMDLLASLYAQAQATPPSSGGGDASGHNNSSGLGASLPTQSQSGSSSRNLTASPLSTSLATAPAPASASAPAPVPTSS
VAQVPVPAPVPVTSSASSSSLGGGAYQTPSAAAAAAAMFHYQTPPRAAFGSAFDMFHHSTPFGVGVGHAHALAHSSGS
GSASFGSPSYRYSPYSLAGSRWQL
>DmelERR
MSDGVSILHIKQEVDTPSASCFSPSSKSTATQSGTNGLKSSPSVSPERQLCSSTTSLSCDLHNVSLSNDGDSLKGSGTSGGNG
GGGGGGTSGGNATNASAGAGSGSVRDELRRLCLVCGDVASGFHYGVASCEACKAFFKRTIQGNIEYTCPANNECEINKRR
RKACQACRFQKCLLMGMLKEGVRLDRVRGGRQKYRRNPVSNSYQTMQLLYQSNTTSLCDVKILEVLNSYEPDALSVQTPP
PQVHTTSITNDEASSSSGSIKLESSVVTPNGTCIFQNNNNNDPNEILSVLSDIYDKELVSVIGWAKQIPGFIDLPLNDQMKLLQ
VSWAEILTLQLTFRSLPFNGKLCFATDVWMDEHLAKECGYTEFYYHCVQIAQRMERISPRREEYYLLKALLLANCDILLDDQS
SLRAFRDTILNSLNDVVYLLRHSSAVSHQQQLLLLLPSLRQADDILRRFWRGIARDEVITMKKLFLEMLEPLAR
>DmelHr38
MMRDRLASLIVVKQEGGSNTSISHHQATAIKCEASLYTESSLFQEINNNSCYRQNLNAPTHQQSHTSHLQHAQQHQTHQ
QHPLLPPPLPTLPLIYPCRNLFPDGCDINHLACCSSSNSNSNCNSDSNSTSSSPGNSHFFANGNTCAAALTPAPPATEPRKIKP
LGAGKLKVGKTDSNSDSNSNCDSRAAAAASTSATSATSATTLAATAAATAAAAEAGGAASAAAAAKISQVRLTNQATTSM
LLLQPNSSFSSLSPFDNFSTQTASTTTTTSASAAGHHQHHNHLLHQQHHNQQQQQQQQQQQQQQQQQQQEHLQQQ
HQQQLVSPQQHLLKSETLLSHEEDQLISNLTDSSVVSHSELFSDLFFPSDSNNSLLSPTTSGYPDNPAEDLTSSIENLTKLTCLR
DKRLSSIPEQQLSSEQEQQLCLLSLRSSSDPAIALHAQQQQQQQQQQQQQQQQQHQQQQQHLQLQLISPIGGPLSCGSSL
PSFQETYSLKYNSSSGSSPQQASSSSTAAPTPTDQVLTLKMDEDCFPPLSGGWSASPPAPSQLQQLHTLQSQAQMSHPNS
SNNSSNNAGNSHNNSGGYNYHGHFNAINASANLSPSSSASSLYEYNGVSAADNFYGQQQQQQQQSYQQHNYNSHNGE
RYSLPTFPTISELAAATAAVEAAAAATVSSPSVGGPPPVRRASLPVQRTVSPAGSTAQSPKLAKITLNQRHSHAHAHALQLN
SAPNSAASSPASADLQAGRLLQAPSQLCAVCGDTAACQHYGVRTCEGCKGFFKRTVQKGSKYVCLADKNCPVDKRRRNRC
QFCRFQKCLVVGMVKEVVRTDSLKGRRGRLPSKPKSPQESPPSPPISLITALVRSHVDTTPDPSCLDYSHYEEQSMSEADKV
QQFYQLLTSSVDVIKQFAEKIPGYFDLLPEDQELLFQSASLELFVLRLAYRARIDDTKLIFCNGTVLHRTQCLRSFGEWLNDIM
EFSRSLHNLEIDISAFACLCALTLITERHGLREPKKVEQLQMKIIGSLRDHVTYNAEAQKKQHYFSRLLGKLPELRSLSVQGLQR
IFYLKLEDLVPAPALIENMFVTTLPF
>DmelFtz-F1
MDTFNVPMLAESSNTNYATEATSNHHHLQHQHQQHSHQQQQQQLLMPHHHKDQMLAAGSSPMLPFYSHLQLQQ
KDATATIGPAAAAAAVEAATTSANADNFSSLQTIDASQLDGGISLSGLCDRFFVASPNPHSNSNMTLMGTATAATTTTTNN
NNNNNTNNNNNNNNVEAKTVRPSNGNSVIIESVTMPSFANILFPTHRSANECIDPALLQKNPQNPNGNNSSIIVPPVEYHQL
KPLEVNSSTSVSTSNFLSSTTAQLLDFEVQVGKDDGHISTTTTTGPGSGSASGSGSGSGSGSGSIARTIGTATPTTTTSMSNT
ANPTRSSLHSIEELAASSCAPRAASPNSNHTSSASTTPQQQQQQQHHMQSGNHSGSNLSSDDESMSEDEFGLEIDDNGG
YQDTTSSHSQQSGGGGGGGGGGNLLNGSSGGSSAGGGYMLLPQAASSSGNNGNPNAGHMSSGSVGNGSGGAGNGGA
GGNSGPGNPMGGTSATPGHGGEVIDFKHLFEELCPVCGDKVSGYHYGLLTCESCKGFFKRTVQNKKVYTCVAERSCHIDKT
QRKRCPYCRFQKCLEVGMKLEAVRADRMRGGRNKFGPMYKRDRARKLQVMRQRQLALQALRNSMGPDIKPTPISPGYQ
QAYPNMNIKQEIQIPQVSSLTQSPDSSPSPIAIALGQVNASTGGVIATPMNAGTGGSGGGGLNGPSSVGNGNSSNGSSNG
NNNSSTGNGTSGGGGGNNAGGGGGGTNSNDGLHRNGGNGNSSCHEAGIGSLQNTADSKLCFDSGTHPSSTADALIEPL
RVSPMIREFVQSIDDREWQTQLFALLQKQTYNQVEVDLFELMCKVLDQNLFSQVDWARNTVFFKDLKVDDQMKLLQHS
WSDMLVLDHLHHRIHNGLPDETQLNNGQVFNLMSLGLLGVPQLGDYFNELQNKLQDLKFDMGDYVCMKFLILLNPSVR
GIVNRKTVSEGHDNVQAALLDYTLTCYPSVNDKFRGLVNILPEIHAMAVRGEDHLYTKHCAGSAPTQTLLMEMLHAKRKG
>DmelHr39
MPNMSSIKAEQQSGPLGGSSGYQVPVNMCTTTVANTTTTLGSSAGGATGSRHNVSVTNIKCELDELPSPNGNMVPVIAN
YVHGSLRIPLSGHSNHRESDSEEELASIENLKVRRRTAADKNGPRPMSWEGELSDTEVNGGEELMEMEPTIKSEVVPAVAP
PQPVCALQPIKTELENIAGEMQIQEKCYPQSNTQHHAATKLKVAPTQSDPINLKFEPPLGDNSPLLAARSKSSSGGHLPLPT
NPSPDSAIHSVYTHSSPSQSPLTSRHAPYTPSLSRNNSDASHSSCYSYSSEFSPTHSPIQARHAPPAGTLYGNHHGIYRQMKV

EASSTVPSSGQEAQNLSMDSASSNLDTVGLGSSHPASPAGISRQQLINSPCPICGDKISGFHYGIFSCESCKGFFKRTVQNRK
NYVCVRGGPCQVSISTRKKCPACRFEKCLQKGMKLEAIREDRTRGGRSTYQCSYTLPNSMLSPLLSPDQAAAAAAAAAVAS
QQQPHQRLHQLNGFGGVPIPCSTSLPASPSLAGTSVKSEEMAETGKQSLRTGSVPPLLQEIMDVEHLWQYTDAELARINQ
PLSAFASGSSSSSSSSGTSSGAHAQLTNPLLASAGLSSNGENANPDLIAHLCNVADHRLYKIVKWCKSLPLFKNISIDDQICLLI
NSWCELLLFSCCFRSIDTPGEIKMSQGRKITLSQAKSNGLQVSLCEL
>DmelHr4
MTPLQRDQLNHQQFVHHYLQSQPVNNKIDSFDIKHIRKTQNPTTNPKFRTEDRFKTSSKSKITNPESGNTQLHFHLSETPDL
SASTSLLPSAFVASSAALVNVTCQAEEGQSSAGSHYTVHRGNDAGISFAALDFRFIRRAAQMNAQQPLILEPGSKASSVVAA
AAELEAATTTAALATSSATEASAADSYPAVQQNGGHEEQRLNGNHGCRREQPDDRTDQPEQLDQQDEPEPVKDVANKL
TIIKTPLNKTLLKKCNSNGSLQMPHANNGSSININNRNNTNNTNNNNSNNNNNAASNDVCDSGLQSGNSNEQEPFANN
DRHHHIYHHHHHHYHHHHGGDKSEASGDVASSSSADPNSQVKDEPTAQDSQDNQAAGAAEAGGATAVAATCSPSKCN
KCNSHNNNSNCNNNNNTISNNNNNTSSNTNTNSNSNNNANNNNNNNNNNHYNNYNNYYKKKSRMPPICRIMTLSRG
PYSELDKMSLFQDLKLKRRKIDSRCSSDGESIADTSTSSPDLLAPMSPKLCDSGSAGASLGASLPLPLALPLPMALPLPMSLPL
PLTAASSAVTVSLAAVVAAVAETGGAGAGGAGTAVTASGAGPCVSTSSTTAAAATSSTSSLSSSSSSSSTSSSTSSASPTAG
ASSTATCPASSSSSSGNGSGGKSGSIKQEHTEIHSSSSAISAAAASTVMSPPPAEATRSSPATPEGGGPAGDGSGATGGGNT
SGGSTAGVAINEHQNNGNGSGGSSRASPDSLEEKPSTTTTTGRPTLTPTNGVLSSASAGTGISTGSSAKLSEAGMSVIRSVK
EERLLNVSSKMLVFHQQREQETKAVAAAAAAAAAGHVTVLVTPSRIKSEPPPPASPSSTSSTQRERERERDRERDRERERER
DRDREREREQSISSSQQHLSRVSASPPTQLSHGSLGPNIVQTHHLHQQLTQPLTLRKSSPPTEHLLSQSMQHLTQQQAIHLH
HLLGQQQQQQQASHPQQQQQQQHSPHSLVRVKKEPNVGQRHLSPHHQQQSPLLQHHQQQQQQQQQQQHLHQQ
QQQQQHHQQQPQALALMHPASLALRNSNRDAAILFRVKSEVHQQVAAGLPHLMQSAGGAAAAAAAAVAAQRMVCFS
NARINGVKPEVIGGPLGNLRPVGVGGGNGSGSVQCPSPHPSSSSSSSQLSPQTPSQTPPRGTPTVIMGESCGVRTMVWGY
EPPPPSAGQSHGQHPQQQQQSPHHQPQQQQQQQQQQSQQQQQQQQQQSLGQQQHCLSSPSAGSLTPSSSSGGGS
VSGGGVGGPLTPSSVAPQNNEEAAQLLLSLGQTRIQDMRSRPHPFRTPHALNMERLWAGDYSQLPPGQLQALNLSAQQ
QQWGSSNSTGLGGVGGGMGGRNLEAPHEPTDEDEQPLVCMICEDKATGLHYGIITCEGCKGFFKRTVQNRRVYTCVAD
GTCEITKAQRNRCQYCRFKKCIEQGMVLQAVREDRMPGGRNSGAVYNLYKVKYKKHKKTNQKQQQQAAQQQQQQAA
AQQQHQQQQQHQQHQQHQQQQLHSPLHHHHHQGHQSHHAQQQHHPQLSPHHLLSPQQQQLAAAVAAAAQHQQ
QQQQQQQQQQQAKLMGGVVDMKPMFLGPALKPELLQAPPMHSPAQQQQQQQQQQQQQQQASPHLSLSSPHQQQ
QQQQGQHQNHHQQQGGGGGGAGGGAQLPPHLVNGTILKTALTNPSEIVHLRHRLDSAVSSSKDRQISYEHALGMIQTLI
DCDAMEDIATLPHFSEFLEDKSEISEKLCNIGDSIVHKLVSWTKKLPFYLEIPVEIHTKLLTDKWHEILILTTAAYQALHGKRRGE
GGGSRHGSPASTPLSTPTGTPLSTPIPSPAQPLHKDDPEFVSEVNSHLSTLQTCLTTLMGQPIAMEQLKLDVGHMVDKMT
QITIMFRRIKLKMEEYVCLKVYILLNKAEVELESIQERYVQVLRSYLQNSSPQNPQARLSELLSHIPEIQAAASLLLESKMFYVPF
VLNSASIR
>PHUM516230_retinoid_X_receptor
MDLERRRGEYNMSGDGAPILAISHIKKEVDGSGFLACSPTNSSTSTNIYSPMKTELPNQMDYGNLQEMNCYDVTHSRSPES
PDQQFCSSTTQFVGEASVNNAEGEIREDDLPRRLCLVCGDVASGFHYGVASCEACKAFFKRTIQGNIEYTCPAANDCEINKR
RRKACQACRFQKCLRMGMLKEGVRLDRVRGGRQKYRRNTDTPYQIHSMLPLKPYLPSLEDNKILEALSLNEPDISVMAQDI
SNADPAVRTLNILSDLYDQELVKIITWAKHIPGFLDLTLNDQMRLLQSTWAEILTLTVTYRSLPGTGELKFAADFSFDEKMAR
ECGAMDIYQHCMHIVERQEKLNITKEEYYIIKALVLANSDVRIDDLPPLKKLRDNILSALADCVAVLRPNTTSLHMNHLLLCLP
ALRQADYINRRFWTAVHREGKVYMNKLFIEMLESYIR
>PHUM596250_Orphan_nuclear_receptor_NR6A1
MRINRDRLLDIPCKVCGDRSSGKHYGIYSCDGCSGFFKRSIHRNRVYTCKAQGDLKGRCPIDKTHRNQCRACRLNKCFIAA
MNKDAVQHERGPRKPKHNITSKENNHHHHHHHHHPVLMHHSNPPLSNSAEQPSAKQPPPPPPPPGIVFPQPINPHMHY
LQSPIKTSQELIPASENFVPNITPIPSGPPSAAAMFLAHQPPPPGLLQILMTAEKCQELLWNTKLSTTERVGVPQSSPLALQPL
SPSWEVLQETTARLLFTAVRWVRSLGPFQTLSRHDQLLLLQESWKELFLLYLAQWSIPWDLTTLLNSSKARDRLPQDEITAN
EIKTIQELIGRFRQLSPDLSECGCMKAVILFTPETAGLCDVQPVEMLQDQAQCILGDYIRNRYPRQPTRFGRLLLILPNLRSIRQ
LTVEQLFFKETIGEIPIQRLLGDMYHME*
>PHUM494010_retinoid_X_receptor
MDHKLEALCKVCGDKASGKHYGVISCDGCRGFFKRSIRRGLAYHCKESNSCIVDVTRRNQCQACRFKKCLSVNMKRDAVQ
HERSPRTSFPIHCTGLSSTGRRNNNNNNSNTYSYSGSFLTAATSGPFLDLPTYHFVYPQPSFPIHFDSLVSKQNVWDLTISSQ
QGHPENVYESAAKLLFLTVKWARSIPSFLQLTFHDQSILLENTWNELFILSAAQWTLPVDEEYLVRVTSLPNNKAKEKFEREV
KNFKKIITKFNNLNVDYTEYACLKALTLFKAETSELKDRLQVEMLQDQTHIMLHDYCTSQDSHKARFGKLLMLLPSVNGLSK
DFIEELLFRKTIGIYK*
>PHUM477600_retinoid_X_receptor

MVTESELQMQGTQPGLQGSLQNFNAPTGITQNCAICGDRATGKHYGAASCDGCKGFFRRSVRKNHQYTCRFSRNCVVD
KDKRNQCRYCRLRKCFKAGMKKEAVQNERDRISCRRPSYDETNQNNGLSVTSLLNAEMLSRQQNGSPVNDYDLSNKQIA
HINDICESMRQQLLILVEWAKYIPAFTELHLDDQVALLRAHAGEHLLLGLARRSLHLKDILLLGNNCIITRYCSENARSPDVDIS
RVGIRVMDELVKPLTEVQIDDTEFACLKAIVFFDPNAKGLSDTTRIKHLRYQIQINLEDYISDRQYDTRGRFGEILLILPALQSIT
WQMIEQIQFAKLFGVARIDNLLQEMLLGGKHLSN*
>PHUM475840_Nuclear_hormone_receptor_FTZ-F1
MRGGRNKFGPMYKRDRARKLQMMRQRQIAVQTIRGHPLTDGQVAFSYGPPSHFENLHNIKQEIQIPQVSSLTSSPDSSPS
PITVSFGQTTSANMIATTQSGQPPVLQIVGTGGNPTSNNLVPSSDHKLWGTANSTTPSPNSISPKAFPFDGVSLPHGSSGG
PGNSSSSSCNKISPMIRDFISSVDDREWQNSLFGLLQNQTYNQCEVDLFELMCKVLDQNLFSQVDWARNSIFFKDLKVDD
QMKLLQHSWSDMLVLDHLHQRMHNNLPDETTLPNGQKFDLLCLGLLGVPTLADHFTDLTNKLQDLKFDVSDYICVKFLLL
LNPDVRGISNRKHVQEGHEQVLQALLDYTLTIYPHIQDKFTKLQQVLPEIHQMASRGEDHLYHKHCSGGAPTQTLLMEML
HAKRK*
>PHUM468660_knirps_related_protein
MESESSSPGIGSALGRWWGQSPQSSSPSTDVMNQLCRVCGEPAAGFHFGAFTCEGCKSFFGRTYNNLGSITECKNNGECV
INKKNRTACKACRLRKCLLVGMSKSGSRYGRRSNWFKIHCLLQEQQQAGAHPLKDKSSAIALWGESYKNLTTPPHFDTDSK
VSPTEESILLNNNNVTKDRERSPNPVLPSYSAEAAAALWAARHSLFPMQLPHHPMPSIPMPFMPPFANFSPPTHSQSQRN
ILLPFMQTLNSSSSSSSPPVRPNHHPPMMMSSSSSRSSEGSICSQSPSPNRNNQRNNDDVSPSTIRFTEKTETTVEKRSSLSS
SSSPPSDKSPTNNNRLSAVNSIKVKNPPLLLDNPIEITNERVDDVASSKKIQTEEHSLALLRSLGPVQDQPMDLSVRVIEIKSRR
KSSKVNHRIAKNEYYHHHHRDDDKNNESSGEEGDAVNDVKNDVLDSIENNDENFNKRKNPLDLTKSTKLEI*
>PHUM467460_Ecdysone_receptor
MFRGGASAMKPIGNCEEDLVLTQVKSEPRVHSPCEKDMTPSASSNQSNSLFSTISSSSKRPRTDDWLSPPSPGPPMGSAPL
TPSPGPHNQYTVISNGYSSPMSSGSYDPYSPNGKLGREDLSPPGSLNGYSVDSSDSKKKKGPTPRQQEELCLVCGDRASGY
HYNALTCEGCKGFFRRSITKNAVYQCKYGDSCEIDMYMRRKCQECRLKKCLSVGMRPECVVPEIQCEVKRREKKAQREKGK
PSSTTNGSPELIMADTPAIKTEPNTTQSHNEKSSANGVKPISPEQEELIHRLVYFQNEYEQPSDEDLKRISNAPSEGEDQSDL
NFRHITEITILTVQLIVEFAKRLPGFDKLLREDQIALLKACSSEVMMLRMARRYDVGSDSILFANNQPYTRDSYSLAGMGETV
DDLLRFCRQMYGMKVDNAEYALLTAIVIFSERPSLIEGWKVEKIQEIYLEALKVYVDNRRKPRSGTIFAKLLSVLTELRTLGNLN
SEMCFSLKLKNKKLPPFLAEIWDVIP*
>PHUM460980_Vitamin_D3_receptor
MNSRASSRELNDAERAKLNELIVANKALLAPLEDDLSGDDFGFNVNAMAGCGPSVLVDVLNLTAVAIRRLIKMSKKINAFK
NMCQDDQIALLKGGCTEMMILRSAMTYDPDRDSWKIPQSKEKLMNIKVDILKEAGGRVFEEHQRFLQTFDQKWRNDENI
MLILSAIALFTPDRPRVVHFDVIKLEQNSYYYLLRRYLESVYQGCEARLTFLKLIHKISELHKLNEDHVRVYLDANPREIEPLLIEIF
DLKPH*
>PHUM411130_retinoid_X_receptor
MPRGGFGNAMHVIWVVAPFGSSTFAELLSAPYSEDVGGETLDPFPEVVFHSQPLSPAPLPSFQEIYSPRFPTRQTKSPELNN
IGIKMDEDYVVYHHHHPSPAMPTTPIQSPYDFHSQHFSPYYQQQPHHPNQGYESSGGPQDSYSLPHFPTSELHINTTLNLR
QRRASLPLQRSESTNSSDSPKLRMGMMTGPPPSASSSASSSPGCGVPETVAPSLLPRPAPQSPSQLCAVCGDTAACQHYG
VRTCEGCKGFFKRTVQKGSKYVCLADKSCPVDKRRRNRCQFCRFQKCLAVGMVKEVVRTDSLKGRRGRLPSKPKSPQESPP
SPPVSLITALVRAHVDTTPDLANLDFSQYREPIPGEPPMSEAEKIQQFYSLLTTSVDVIRHFADKIPGFTELTKEDQDLLFQSAS
LELFVLRLAYRTKADDTKLTFCNGVVLDKHQCQRSFGDWLHAILDFCQSLHAMDIDISAFACLCALTLITERHGLKDPHKVE
QLQMKIIGSLRDHVTYNAEAQRKQHYFSRLLGKLPELRSLSVQGLQRIFYLKLEDLVPAPPLIENMFVASFGKKKTKITRNEDE
TREEPTESTAEIYLFTILGR*
>PHUM331460_Orphan_nuclear_receptor_NR2E1
MMMTMFTGRILYDIPCKVCQDHSSGKHYGIFACDGCAGFFKRSIRRSRQYLCKAKSEGSCTVDKTHRNQCRACRLKKCVE
AGMNKDAVQHERGPRNSTLRRQMAFYFKEPCDTTSDIGCGPSVSPILQHSQTHVLDLALPKIPTSVSPRSESNGSPLMQPT
MAAAFLFGQTLIPKLTPNFTLTNAIPLGSVEALCESAARLLFMNVTWAKNVPAFTRLNYKDQLLLLEESWRELFVLSASQFM
LPLELVDLLTAHTTVTANSEKALTIAQEVKQFQETLIKFKQLHVDIHEYACLRAIVLFKTSFDPSNNTVTSSTAPSSSGEGKTLH
DLAEVAAVQDHTQLTLNKYISAAHPTQPFRFGKLLLLLPSLRSVSNNTIEELFFRRTIGNIPIERIICDMYKASD*
>PHUM318510_Ecdysone_receptor
MDATATSTSWTQSASTTRLASQNSPTKGVVSVVVDSRNESPNVFAVKEEHIVGHRGTSITCGGVTVTSINVPQNEEQDFSS
QNVKNTLHFGANERAEIGIKCEPLSSNFNDDDSDVHLRIKKKRSNKNDEKSQDILKIEEPERPMSWEGELSDTEMSIVTDGR
RMRDDIKDELMSDVQMSPQSIPKSEIGDAIIMESGRLKQEPETCNETMSGYSSSSRELPLLVDKLLGGYSTPSQSPILQQRQ
QHHSLTKQHSLLGALPPNPSPDSAIHSAYSYSSPAASPAASRHLPSSGFSSPFTPSLSRNNSDASQYGGSQHSSCYSYNSENF
SSPTHSPIQSRHLLGYSKVDGSPIHSVTGGQLDYQQQHLDNERDEEKDPCLEEKMSCLASDFSQHSALATPPGISRQQLINS
PCPICSDKISGFHYGIFSCESCKGFFKRTVQNRKNYVCLRGSQCPVTIATRKKCPACRFEKCLKMGMKLEAIREDRTRGGRST

YQCSYSVPSSLLQNSDSIHKLSNSLTAGETLVKLETPHLGSPNSNLDSRLLPVPQLLQEIMDVEHLWHCNEREMLGSPMNN
HRHSPSSTPSSPSPSQSQTNSESNPDFLSNLCNIADHRLYKIVKWCKSLPLFKNISIDDQISLLINSWCELLLFSCCFRSMSSPG
EIRVSLGKSISLSKAKELGLGPCIERMLNFTQHLRRLRLDKYEYVAMKVIVLLTSDVNDLREPEKVRASQEKALQALQHYTLAH
YPDMPSKFGELLLRIPELQRTCQVGKEMLSIKNREGEEGPSFNLLMELLRGDH*
>PHUM301950_Orphan_nuclear_receptor_TR4
MTAKYRSNDLEADIKMEQPPLNCAIKMEQPHLNYGIKMEQALDLNISSGLKVEQALDLNMSSNLKIEQAMDLNMSSDKV
RSLCLAVELCVVCGDRASGRHYGAISCEGCKGFFKRSIRKQLGYQCRGTKNCEVTKHHRNRCQYCRLQKCLEKGMRSDSVQ
HERKPISEKNSRDLSSLIASNNNSQPFLPTDINSKIYLRKDLENAPSAFAAAFGLADLGLLTATTLGKSSCKFIEIHLLIKYNMFV
HLGNEIVKRESPHMSDEEDSAESDVNEESGDALSWAREKTLKESLDLLSKLLNENYNANESATEEDRKPLDCDESLLQDYHS
TFNLQTPSPVPVHLNVHYVCESACRLLFLSIHWARSISAFQNLSHDLQVALVRRCWPELFALGLSQCSQILPLRAMLAPIIAYL
RTAVTQDTMAAQRLIDITDHIFKLHEYVTRMVSLQIDNHEYAYLKALVLFSWHLPGLPLALKRQIDRLQEKALQELKSYVAK
NNSEEDNRSSKLLLRLLPLRFLQPQIMEELFFAGLIGNVQIDSVIPYILRMEADEYRVSLLNEQASQGFET*
>PHUM234430_Nuclear_hormone_receptor_E75
MSRDSVRYGRVPKRSRERSTEEQSRVSTSDAEQSDSDTKQLAVYDVILTVSQAHHANCGYTEEHTRNLVRKPITIPPLNGSS
STDGPEVASSTAESLEQERIWLWQQFASHVTPSVQRVVEFAKRVPGFCDLSQDDQLILIKVGFFEIWLGHVSRQTCDSSMT
FTDGTYITKQQMDLIYDPEFVSSLFHFAASFNALALNDTELGLFSAVVLLSADRPGVADIKVIEQHQDRLIEALKVQVGRNHA
SDPQLFSSLLMKLPELRNLGAKHSAHLDWFRINWQLLRLPPLFAEIFDIPKCEEDLQ*
>PHUM233230_ecdysone-induced_protein_75B
MAFIPAKVARNETMQENRFPNNEFQNLSLDEKKGASRTTSSSSSSGDEGDRSSSSSSSDDEKCPHEFCDNDYDYDVDDAV
RFGRVPKREKARILAAMQQSTNSRSQEKAVAAELEDEQRLLGTVIRAHLDTCDFTKDKIEPMIARAREQPSYTACPPTLACP
LNPNPQPLSGQQELLQDFSKRFSPAIRGVVEFAKRIPGFSLLPQDDQVTLLKAGVFEVLLVRLACMFDSQSNSMICLNGQVL
KKESIHNTSNARFLMDSMFDFAERINSLGLSDAEIGLFCSVVVIAADRPGLRNTDLIERMHNKLRNSLQAVVTQNHPGHTDI
CNELMKKIPDLRTLNTLHSEKLLAFKMTEQQQLQQQQQLQQQEMWTEEEVHSKSPAGSSWSSSSDVTMDEAVKSPLGS
VSSTESVCSGEVAAITDHQGGPASTAPLLAATLAGGICPHRHRANSGSTSGDDDIMGSHLLQNGLTITAVNNPSKSYNQSH
RFHRKLDSPSDSGIESGTEKIDKVTNSVNSAPTSLCSSPRSSLEDKDEDKQHTHIVEDMPVLKRVLQAPPLYDTNSLMDEAYK
PHKKFRALRNKDSAEAEPMIVVSPVPSVNSSEHHSSTQLHRHLTATTSAPPSLTGSSLSSTHSTLAKSLMEGPRMTAEQVKR
QEIIHNYIMRGDPPAEFNQSSSSSSCPAPRPNNNNLLNCGGNWSQPQTGYHYLPTQPQPSRWHSNGASVITTSRPFSLTP
NSTSSPPPVLQKALEQPNAEFSRIYLQPQSPHLASSSTSPSPIARNVSPSSCSPASRSPSVAGGPQMTLPSPKMMELQVDIA
DSNQQPLNLSKKSPSPSPRPLSAPSTAVQKALTLEA*
>PHUM195250_orphan_nuclear_receptor_nr6a1
MTLTRNPCELDTMSLFQDLKLKRRKVDSRCSSDGESVAETSTSSPDMAGPPSPCPKLTSPSPEPETPRNRVRLSPEQPRIRSS
PEQRIRLSPEHPRVTSEQLKLRLSLEQSRIRLSPDQQPRVRLSPDQQPRLRLSPDQQPRVRLSPEQQQRLSPLIQKGSPVIQRS
MSGGEQNTSGGVLNMISPGAFWVNSAASGRINGVRPELIGGNVNMNFPVNSQNSDMKPPRPVSTPGTPTMTRQTPTVI
MGEAGGVRTMIWSQPNPSSPADHQNQNMAASTSAINWSTSPQSSAVIVVKNQRRNFYLISVNPNPLNMERLWAGDLT
QLPVSQQNQALNLALPPGDWPRDGRPQGITIIGEPKHDPEDDEQPMICMICEDRATGLHYGIITCEGCKGFFKRTVQNRR
VYTCVADGNCEITKAQRNRCQYCRFKKCIEQGMVLQAVREDRMPGGRNSGAVYNLYKVLMRERDNKFLNVIFNNNEKG
NQQKSSISSEQTLPSHIVNGTILKTALTNPSEVVHLRQRLDNTVSSSRDRTFPLDTILNMIQALVDCDEFQDIATLRNLEDLLD
HKSDLSEKLCQIGDSIVYKLVQWTKRLPFYLELPVEVHTRLLTHKWHELLVLTTSAYQAIHGVHKLSSTSTAGQEADFTQEVS
NNLVTLQTCLTSMMGRPITMDQLRQDVGLMVEKITHVTLMFRRVKLRMEEYVCLKVITMLNQGRGGSNELEAIQDRYM
MCLKSFVEHSFPQQPARFHELLLRLPEIQSAASLLLESKMFYVPFLLNSAIQR*
>PHUM164330_retinoid_X_receptor
MMKKDKPMLSVAAIIQGVQVQHWSRGKTNVMMLLSVELGGPQSPLDMKPDTATLLGGNFSPNGGPNSPNSFTMGHS
SLIGNNSNNKMASYPPNHPLSGSKHLCSICGDRASGKHYGVYSCEGCKGFFKRTVRKDLTYACREERNCIIDKRQRNRCQFC
RYNKCLAMGMKREAVQEERQRTKEREQGEVESSGAMQADMPVERILEAEKRVECKVENQNEYENAVANICQATNTQLY
QLVEWAKHIPHFSSLPIEDQVLLLRAGWNELLIAAFSHRSVEVRDGIVLGAGITVHRNSAHQAGVGTIFDRVLTELVAKMRD
MNMDRTELGCLRSIILFNPEVRGLKSGQEVELLREKVYAALEEYTRVTRPEEPGRFAKLLLRLPALRSIGLKCLEHLFFFRLIGDI
PIDTFLMDMLGSSSDS*
>PHUM053030_hormone_receptor_hr3
MFDMWNSVSPKLETSQPGTTSNSQLPQTSTNAAGSIKAQIEIIPCKVCGDKSSGVHYGVITCEGCKGFFRRSQSSVVNYQC
PRNKNCVVDRVNRNRCQYCRLQKCLRLGMSRDAVKFGRMSKKQREKVEDEVRFHRAQQRQATESTPDSSVFDQQTPSS
SDQLHYNGYSYNSSDVGSYTNYNYTPHHIQYDISADYVDSTTAYDPRPGLDSLTADSNLMTSVVTTGGNCHSGNNTVENQ
SNQRINSLNNQTTGNDRTERNGSNERTGLEGAQRQGLDRLVTVKQENVDNIIGGGGFVDSTTLLQPMSPSSQASQNLSPR
SQVSQTMTSPSVKLKEEEDINQGCPSKMYPMQISELLSKTIADAHSRTCVYTLEHIHEMFHKPQDLSKLIYYKNMAHEELWL
ECAQKLTTVIQQIIEFAKMIPGFMKLSQDDQIVLLKAGSFELAVLRMSRYFDLSQNCVLYADTLLPQDAFFTTDTSEMKLVTS

VFEFAKSVAELKLTETELALYSAVVLLSGDRPGLKGTTEINRLRHAVARALRFEMDRNHILPLKGDVTVYDHIMAKIPALRELS
LYHMDALAKFKRSTPLLEFPALHKELFSVDS*
>PHUM039420
MGSSPECCSAGSSPAHPLPLCLGSCTPHHQTVVQAQRSPPSPFLLGHLPVNKGKILGLSCVVCGDTSSGKHYGILACNGCSG
FFKRSVRRKLIYRCQAGTGRCVVDKAHRNQCQACRLKKCLQMGMNKDAVQNERQPRNTATIRPEALVEMDQERALREA
AVAVGVFGVHIFLLKGSQSTKDKSINRDKSNHISISGRSVPPVSLAVGFSPVRYPQSLTSPSEVNSSKQEADEEDSIDVTNEES
LPPTPTSRTLPCTISTPTTHLPPPPLYTTTQETIYETSARLLFMAVKWAKNLPSFASLPFRDQVILLEECWSELFLLNAIQWCLP
VESSPLFSVNEHAATVPNGKSSQTAADIRVLNDMLLRYKAVGVDPAEFACLKAIVLFKSETRGLKDPLQVENLQDQAQVML
GQHARGQHPTQPARFGRLLLMIPLLKHVPTQRVEHIFFQRTIGNTPMEKVLCDMYKN*
>CLEC000589_HR96
MDNEERRSHEKFCSVCGDQALGYNFNAVTCESCKAFFRRNATKTRELQCPFKQCCDITAVTRRFCQKCRLAKCFRVGMKR
ELIMTEADKERKKRKILANKARMVTAKAKDSIGNSKSVQVSMVSVEIGTQTEPLTCNCRQRYKADVPNSLLMAALPPPDKT
QITYQLIKSIPTATLFPMSCQDNELLEDLIVANKALEAPLDQELNNLPGDEFKVCAFILYSCGSKSLLDVINLTALAIRRLIKMCK
RIGGFRALCQEDQLTLLKQGCTQMMLLRSVVTFDPDRNSWRIPHSVDQMSQINVEVLKEARGNLYDAHEAFLRTFDTRAS
HDLNIMCILTAIVLFDPNRTNLVNKQLIAQQQPDILLLRQISCYFQSRYFSLLQCYLQSIYSSEESNEIYQHLICKMTELHQIIEEH
VRVYLNVNPSQVEPILIEIFDLKTH*
>CLEC002129_EcR
MWVRGYAMREEGCDQVTSSCSPGVDDLELWDLGLGPGPRGIQLSEHRGDISGREDLSPPNSLNGYSADSCDGSKKKKGT
AVRQQEELCLVCGDRASGYHYNALTCEGCKGFFRRSITKNNVYQCKYGNNCEIDMYMRRKCQECRLKKCLSVGMRPECV
VPEYQCAVKRKEKKAQKDKDKPVSTTNGSPEAIKVEPEPHRVSYTSSLFQSMIKESQTQSTEGELAKVAVNGIKQVSAEQEE
LIHRLVYFQNEYEHPSDEDVRRINTPNDEEEQSDLKFRHITQITILTVQLIVEFAKRLPGFDKLLREDQIALLKACSSEVMMLR
MARRYDAQSDSILFANNQPYTRDSYNMAGMGDVVEGLLRFCRQMYNMKVDNAEYALLTAIVIFSERPSLTEGWKVEKIQ
EIYLEALKSYVDNRARPRSPTIFAKLLSVLTELRTLGNQNSEMCFSLKLQNRKLPPFLAEIWDVNP*GRQGQSPLFPIRARRRF
LAESAQSRSV*TVTSTPCLTKPTWFKTINFSMCET*N*S*QYVLKMYKRFCFVFFKYFLLKKAYKQWLYFCIRHCFF*KLWEV
QMTFSFF*CT*K*GKVTFGTPY*DGLCNLQRRMQ**ISFFFSVYRK*E*LIVYEKKKLLLVII*Y**KIFLIINWNLKK*IKCEIAIG
VIGHKHIHTKWQ*TQIGLATKIWFPLEGMVVNFVN*FLFYICFLIMVNFHHTRLAVSIIHTYFYLYFVQ*RIVQQIKKWKKNIL
KNCLRVFFFVNN
>CLEC003396_HR39
SMVLSGGPIVSAGLCPFIIRRSHFSVYM*LVFVLLVIPTDTLLDIRNIL*RISLFKE*WLRREGNAGSRQSGGQGAEKGWRGA
FQSSGENLRQEPASSHRAKSQSPQSTFTTAPMTKRKIITNLSAMEVLASVLLWSAENR*MSMKIHPRQTLKSATLPVSLSI*
GTRTGMNPGALLKGPCLGRASCLIRKSLRSNPSQRRIRLKIQKAAATPRAVK*WLR*KMKSL*H*RMRKLQRVRRQINSML
SLNYPFL*ISF*VAVVY*KDQDRLLAQTQPFTLVIPQLQAQSRQDIFFPRLLAFQALLHLPCQGTTVMRPSTVDHNTRVVIVT
VILLCPFHQHSFLRHILQFREDICTDLDFLLQF*IQEGFMNIQCKMKVVRWMINTHR*LPTCSNIPQGLPHLESAGNSSLTAR
VRFVGTRSPVSTMGYSHAKAVKVSLKGLFKIERITSVFEGHPALSR*RRGRNARLADLTSASIWA*S*KQ*EKIEPVGAGAPI
NVLTQCQRDLWSKSAQSLHHRPRLRPTLFLHYYRKSWK*SIYGNIMKLILNWLEEFQDHKVEIQQI*WPTFATSLITDCTRL*
SGASRFHFLKIYRSMIRRLCS*MHGVNSSCSLAVFGRCPPLEKLGFHSESAFH*LRLKTSASAHQSSECSTSPNI*EGSESTDM
NTLP*KLLSFYLQIRAI*ESLKKSGLRKRKLFRHSSIIRLPTIQTFRPNLESYF*GYQTCKEHAKLVKKCSRLKVKKEKDRALIF*WS
YFEETTELLRL*SINVPCWGAIKYVRLL*KF**LILCHWQHI
>CLEC003430_HNF4
MFVRDVEDGEMGEGSLSVGGNTPVPNFPQSCAICGDRATGKHYGASSCDGCKGFFRRSVRKNHLYNCRFSRTCVVDKDK
RNQCRYCRLRKCFKAGMKKEAVQNERDRISCRRNSYDDQTIGNSLSVNSLLNAEILSRQTGAALEMSHDYDLSNKQVATIN
DVCESMKQQLLFLVEWAKFIPAFLELQLDDQVALLRAHAGEHLLLGLARRSLHLKNVLLLGNNCIIPRHTADAGGTADLDIS
RVGTRVMDELVRPLTEVQIDDTEFACLKAIVFFDPNAKGLTESGKVKFLRYQIQINLEDYISDRQYDSRGRFGELLLTLPALQSI
TSQMIEQIQFAKLFGVAKIDSLLQEMLLGGAPDVPAANGQTNSTPMNIVNYPSSGGSPESQNGLSPVHSPPQGSNTVLRDI
STSTQDYQYSFKQEPGLDV*
>CLEC004804_tll
MDSRTKNHVNDRDDNAGRILYDIPCKVCQDHSSGKHYGIFACDGCAGFFKRSIRRNRQYVCKVKGEGGCLVDKTHRNQC
RACRLTKCLEAGMNRDAVQHERGPRNSTLRRQMCLLVKEEAGPSTPSMDLSAPNASPPHFSPFYTPQIGEVLSNPEAICES
AARLLFMNVRWAKNVPAFTGLNMKDQTTLLEESWRELFLLGWAQLLPLTDLAILVANSPRLLQEAATFQDTLAKLRSLGLD
HHEFACLRAVILFKTALEGEGKSLVDVASIAALQDQTQVTLNKYVSTAYPDQPYRFGKLLLLLPSLRLVSANTIEEMFFRRTIGP
IPIERIICDMYRSAGAI*
>CLEC005164_USP
LSLENNLALVGPQSPLDMKPDTASLLVGGFSPTGGTSGSTSPTTFNIGHSSVLGNGNKSTVVYPPNHPLSGSKHLCSICGDR
ASGKHYGVYSCEGCKGFFKRTVRKDLSYACREDKQCLVDKKQRNRCQYCRYQKCLSMGMKREAVQEERQRTKDRDQNE

VESTSSFHTDMPVERILEAERRVDCKIETPPIDFENQTNEICQATDKQLFQLVDWAKHIPHFTSLPLEDQILLLNAGWNELLIA
GFSHRSVTIRDGLVLGSGVTINRNNAHQAGMGTIYDRVLTELIAKMREMKMDKTELGCLRTIILYNPEVRGLKSVQDVEML
REKVYAALEEYSRMSHPDEPGRFAKLLLRLPSLRSIGLKCQEHLFFYRAIGDVSVDTFLLQMLESPERL*
>CLEC005658_HR38
MKATRPNIITSKQNKTARIGGSSSSDSPKLRQGTTPCPSASVSSSASSSPTERPNAISPSQLCAVCGDTAACQHYGVRTCEGC
KGFFKRTVQKGSKYVCLAEKSCPVDKRRRNRCQFCRFQKCLSVGMVKEVVRTDSLKGRRGRLPSKPKSPQESPPSPPVSLIT
ALVRAHLDTSPDISNLDYSQYREPNEEDVQMSETEKTQQFYNLLTSSVDVIRHFAEKIPGFSELCREDQDLLFQSASLELFVLR
LAYRSRVVDMKMTFCNGVVLSRNQCQRSLGDWLHPILEFCQSLHAMEIDISSFACLCALTLITVSMSLNLY*
>CLEC008419_PNR
MLSMQEEGKEAYCKVCGDKASGKHYGVASCDGCRGFFKRSIRRNLDYVCKEEGRCVVDVTRRNQCQACRFSKCLRVNMK
KDAVQHERSPRSNSATPSLSFVYQQFLPYPRAYLHPASIPGLQYMGCYMRSECNENEVSSSQESTESRLEPSVSLLSEDIYSTS
SRLLSLTVQCVRAIPSFQQLSPQDRDSLLEDSWKDLFILTLAQWVTYLAPIQFLSGFFVLFMTFIISLLVLLEKLIRDTGIDDASR
LDELRKDLKVVSYAIARLTQLRADHTEFACLKALVLFNPESVRKEEVEVLQEQTHVMLAEYSGVRSAKLLLLLLLNTSRTSLKNIL
LLFFNRHAPVRLEGLLASK*
>CLEC009113_HR4
GLSPECGGRKGGEVRVKEELVFDGGGPAPATSGSGYWPPQQQQQQQPQQHQAQQQNPRVNGVRHELIGNEMRPPQ
APTPRPAPTVLMGEVGGVRTMVWSQPAPAEQPTTSSWPPPNQEETAAQLLLTLGQETSPNTTGTRALNMERLWAGDLS
QLPGAQQITALNLSWAKQQPTKPDDHEDDEQPMICMICEDKATGLHYGIITCEGCKGFFKRTVQNRRVYTCVADGVCEIT
KAQRNRCQYCRFKKCIEQGMVLQAVREDRMPGGRNSGAVYNLYKVKYKKHKKSVRNGQIKSVNEKGKPAISPEHSIPPHL
VNGTILKTALTNPSEFMCTMVVHLRQRLDNAVSSSRDWALPIDATLSMIQTLIDCDEFQDIATLRNLDDLLEHKSDLSDKLM
QIGDSIVYKLVQWTKRLPFYLELPVEVHTRLLTHKWHELLVLTTSAYQAIHGAHKLASTGSDGTEAHFTQEVSNNLCTLQTCL
TSMMGRPITMDQLRQDVGLMVEKITHVTLMFRRIKLTMQEYVCLKVIIMLNPARGGSNELEAIQERYMTCLRTYVEHNSP
NNPNRFHDLLVRLPEVQSAASLLLESKMFYVPFLLNSTIQRX
>CLEC025065_Knirps1
MPKENEKKKERKREREKRKQFTFFNYVHKSKSVDRQARNYTTAKLDEPRMNQQCKVCNEPAAGFHFGAFTCEGCKSFFGRT
YNNLSSISECKNNGECVINKKNRTSCKACRLRKCLLVGMSKSGSRYGRRSNWFKIHCLLQEQGGLPAVKKEDEEEEAQKERR
FTPPPPFYPQPRRHFLFPQEPWQPWPPTPPEAPQDQPIDLSLKPRTDSEVLDLTKRQALAGWSS*
>CLEC025067_HR3
MVKHPSGTYNRRNDSTPLECPGCDPSLLEYAFPNSQWSASRGTVKGRTTPVATPRSHPAAAVPSLEQSVGGVIGVISWVR
RRQKLVKMAKCRRNAQAQIEIIPCKVCGDKSSGVHYGVITCEGCKGFFRRSQSSVVNYQCPRSKACLVDRVNRNRCQYCRL
QKCLRLGMSRDAVKFGRMSKKQREKVEDEVRYHRAQLRAQVEQTPDSSVFDQAQTPSSTDQLHNYTGYSAYGSDVGSYS
YNYTGQVTATMQYDISADFVDSTTTAYDPRPSIDQMSESSMMSANNVSTGGGVGKVVQGQQRLAIKLESEPISDSIVNTF
VDSTTSRQSEDVSPKESESNSNSYTSSLIDPAQISELLSKTIADAHARTCLYSIEHIHNMFRKPQDLSKLIFYKNMAHEELWLEC
AQKLTTVIQQIIEFAKMVPGFMKLSQDDQIVLLKAGSFELAILRMSRYFDLSSGWVLYGDTMLPQDAFYTTDTAEMKLVTL
AFEVSSGVAELKLTETELALYSACVLLSADRPGLKGLAEIGRLGQAVLRALRIELERNHTTPIKGDVTVCDALLAKIPTLRELSML
HMEALGKFKRSTPHLEFPALHKELFSVDS*
>CLEC025112_dsf
MVLIDPFVQGDRLLDIPCKVCGDRSSGKHYGIYSCDGCSGFFKRSIHRNRVYTCKAQGDLKGRCPIDKTHRNQCRACRLTKC
FQSAMNKDGMGGTIAVQHERGPRKPKQSKDGESPVRQVSGNNGVCLESSQHSPLKLTSPAYQQAPMQQADSPERRPQ
PLFVSHQPPGLLQLLMSAEKCQEIVWNGKLDVINGGNSGPGSPSATAIPLSLSPSWEVLQGSICSQETTARLLFMAVRWVR
CLAPFQTLSKRDQFLLAVNLTSMCIYLQLLLLQESWKELFLLHLAQWSIPWDLSPVLGGPKARERLPLDDTLVPNELNAIQEIL
ARFRQLSPDGSECGCMKAVILFTPETPGLVDVQPVEMLQDQAQCILNDYVRGRYARQPTRFGRLLLMIPGLKVIRQSTIERL
FFRETIGDIPIHRLLGDMYVMEKSYS*
>CLEC025114_E75
MLFDTRVPQYHGQYFAPGLPPQLPEAQPQQADHQLNIEFDGTTVLCRVCGDKASGFHYGVHSCEGCKMGCLCDQGFFR
RSIQQKIQYRPCTKNQQCAILRINRNRCQYCRLKKCIAVGMSRDEQFEPTRTHCFTNVSLDLFPKRVDNNTKDFQQPPFNK
KRPVMMTEELPILKGILNGVVNYHNAPVRFGRVPKREKARILAAMQQSTNSKCHEKALAAELEDDQRLLRTVIRAHLDTCD
FTRDKVEPMILRAREQPSFTASPPTLACPLNPNPQPLTGQQELLQDFSKRFSPAIRGVVEFAKRIPGFGILSQDDQVTLLKAG
VFEVLLVRLACMFDTQNNSMICLNGQVLKRESIHSGSNARFLMDSMFDFAERLNSLRLTDAEIGLFSSIVVIAPDRPGLRNTE
LIERMQNKLKAGLQLMMSQNHPNQPCLAQELMKKIPDLRTLNTLHSEKLLAFKMTEQQHLAEQQQMWGTVVSEDESKS
PSGSTWSSSSDVAMEEVKSPLGSVSSTESMCSGEVEYHSSHAASAPLLAATLAGGCPMRHRVGVEDNKDIKPIIIHRTFRKL
DSPSDSGIESGTEKVDKLTSSAPTSLCSSPRSSVEDKEDHQIEDMPVLKRVLQAPPLYDTNSLMDEAYKPHKKFRALRKECGD
EPDNTTSTSSLSSTHSTLAKSLMEGPRMTAEQMKRTDIIHNYIMRGEGCGWQQQQSVITTSSAARGGAYVIVNNSSPGPSI
YVPAGSPCPSSTSLVELQVGTQPLNLSKKTPPPSPRPKVLSLEA*DHPPSHVFFTCE*YKVVSAAMWYKSKPLL*SLPPCKYLS

ISLL*IGLSIVSE*IFCIIIVLNELF*WCHF*TVLRRKTK*FIIG*SIFHSLSIEFYYYDDSLLFF*DDYMCIRSEEEKKLISFFSFPILLYK
CF*NDDVVVC*SLLCILGIL*RNFNFILGRVYWPIADLNYGHRP*SLPFKTFFCFKF*KRI*IYIFINVIYFDIVGLAFYLCVYI*SHI
SLIKKKLKLKTKKKSYTS*HL**SVPRVHIMSRFLRLMEGTPASESRA*NKASIYLSPPTFKKNLFKQPKKRQKKNKCNIYIY*LL
KLRK*LQSLLFLLILKIFVHNKHNIYFAVYIF*QMGVASDAYYIISNCRPMFYLDFGYVCSSKGRGTRSVQPADLQFCFLILQKK
EKKKEPAIKX
>CLEC025160_seven-up
MEMIDDKCVLREHDKKLLKYSVGGVLCGGGASSGAPPYHHVPPEFPPRAIVPWRDPSLVTLTQEDTLVSQAIRGDTPGTNS
TQSGGSQADKNQNIECIVCGDKSSGKHYGQFTCEEEKTLPDEFYLIKLRTIIKMLLCFGFAAVQRGRVPPSQPPSLPGQFALT
NGDAMSAAAAAAAAAAGFNGHSYLSSYISLLLRAEPYPTSRYGQCMQPNNIMGIDNICELAARLLFSAVEWARNIPFFPDL
QVTDQVALLRLVWSELFVLNASQCSMPLHVAPLLAAAGLHASPMAADRVVAFMDHIRIFQEQVEKLKALHVDSAEYSCLK
AIDMKKLDISSERVPVSILLESVLVKVSLTVSAHGPVAKRGLDACGLSDVAHIESLQEKSQCALEEYCRTQYPNQPTRFGKLLL
RLPSLRTVSSQVIEQLFFVRLVGKTPIETLIRDMLLSGNNFNWPYTMX
>CLEC025178_Ftz-F1
PLLIESFLPILVESAVPRADGESKTVADRVPSSCGDRRLLLGRRPSLENRPVGHVCSFNLYPPRICPSSLTPVPDRGSTCVGPEP
AKRSQTSREAEMISSQTL*FHPRPSSVEWGSTVQVAYRPTFVARSIFTPRPAPAETCGALA*KNRPLFSRACSDLGSRFSRPF
FREAKNNGKWATMT*R*ACRRAPLRRPTRKSNRSTLTTLKLKSSKYHSTIILQVPASGWNCLLAAPLPSSSSRRCCSKPCPTC
RTPRKGSRSSVPCAATRFQATTTDSSPVNRAKDSLKGPSRIRKCTPVLQKGVVISIKHKGSDALFVDFRSALKSA*N*KRFVPI
E*EEVETNSVLCIRETELESCR**GRDSWQPRPSGTAAAIARVASETQ*PSVTSRAPTSRASISNKRSKSHKSRHSPPPRKAH
QVPSRWPSARPRTETSPTTTCAPSCPDLRLSPLCRSSTTKGSPTKASRLSNTPNSGRATRRPRLRKRSNTKVLPPRASPKYRP*
FGTSSKPSTTTNGKTPYITYYKIRRIISAKWTYSNSCAKCWIKTFFRKWTGQGIPYSSKTSRLMTR*SYSSTRGRTCSCSTTCTK
GCTIHCPTKRHYQMDRSSIC*ASASWAFHPSPSPLQRSLPNSRTSNLT*VTTSASNSSSYSTQK*EA**IGNTSKKDMNKFN
NPFMTTV*QHIHKCRTSSTNSCRSFPKSIDWRQRVRNTCT*NTVTEVHRHRHSSWRCFTQRGSNNINRYVKGLIANLQYSF
RQKHTGVPASKCQLIRPVCQF*RSRITIKRILL*KEQQSPMSNVQYSFKEEKKKIRRKVRLQVSCVHKLISSYLLCALTTVYQAH
KTTV*IIKNSL*ILLCYGYSFKCVMCKTEGLFDSFLDLIV**KREMPSKYI*NIKKKIKNKI*KK*KQKKYKAHYKLLIFLLKKGII*T
KKYIFHLAGPTPTISLFFFLK**IEFYHILLFVV*KKNCYRPIACRFLFLIIF*MYKHSKIYKRIFF*RRLLNYCYTFQFYKRVLFWKG
YFGHHKTKIYLCIINYKSLSSQNC*KHVRWIHMACKKRKNSCD*ILLIYFLIKMYKNNL*KKSL*KKLYVCKSA*TIRNTNCNNL
HSKTKVSTFIFPFAIVK*LILFFFLFGYG*FPLDIAHFLLSFLSTNFVSLLVNTFGVMVLVHVLILTF*FFKFFFLENRKEK*FLKASV
CCFK*GNLNKSWPLTI**LF*KN**LLTVLVDIFGMHVSC*DILLSFKKKCVFCF*FEIFLWF*SVKKR*TNKSYQL*HGLIVKIG
PNCDQSPLL*HVT*IKSKWYF*SL*QNMHDFVCIL*FLLIQEYIIV*KIIYQLQINCFNKFFIRRVRELNVRERVIE*VRLIKKNRC
RKTLKRFPQEIFHTFFSI*FITVKLLCEWFCVEFEFFFIIVRVCFFCHSYLLELTIR*KSC*KKIYNFKFYNPRPKLASTVSAVLWRIR
KRREKNLHSIAYF*L*NALCIITNICLRCQQIDLFLSKSMILFKKSALLKKHFLRSPLAHI*LSSLITQLSLQEQGLSLVTFYNVYLKII
WIFLFKSGEYRGSSRTC*NFMKCSNEKLLY**NEIYFYLPKLCSLNDFYNFYRTRFF*IAGTIFSHLKIKYRSMFKFMRVLKTFT
QNV*NTR*ICLRNRGLFKYKRRFFFPLKKFVNMHNFLKLYFCLLVFCSF*TAYVCAMYKKNNNVNEI*IMYIFCLMLSIFYTAR
LFK*SVT*LK*WKH*SIQYNMKKCTFFYVCCKFCDD*KKEIGILV*EYIIYFYGKCYIERRSKGEQLLEFLLIGKKRK**LKKVYCD
*REFIFKKDIKLL*EFLRTSVKILIPVIKIR*KKIHIFYLLLHLSRIL*L*II**TSHLFFSGLMFFKILLN*EKKIISSKIDTMNTSLHIILY
NMHINK*EIT*IFYSTYFCTQFWKCLVLHNKQKTQNKKLLCIFFNCFFFYYKCIHFSLSVKIFIKQNI*IIICI*G**KINNKYLKQS
ESTI*YIVMFDYLCQITKISCDDFVIICF*TVTDFLCPVTYHSLFLNIVY*DMENCYLNLYS*LVFLPF*QVSITMLIINGGYCCIM
SVYIYYRHLLCRCYCFFISG*KMASLCQ*LVCI*S*VLYCRVK*KIIKGVKSKSFIIISIIMIMIIIIIVLNLL*FLYYQIIKW*TVIN*HG
ETVLLPVX
>CLEC025263_HR51
MTTRWRVEDYSSRWNCGPQTVVRATRSPPPPQLGGGKLKSLGLSCVVCGDTSSGKHYGILACNGCSGFFKRSVRRKLIYRP
SSSMSLGLSMQYKNMISANASPGQEEINAVQPRSQYSQPSPPKSSRDENEDEDTIDVTNEEPPATGTYSMSASAPPLPPEY
VFYPTMPETVYEVAARLLLMGVKWAKNLPSFASLPFRDQVILLEECWAELFLLNAIHWCLPLETCKLFSVPEHVMAAPSGKE
GLVAHEVRALNDALQKFKAIRVDAGEFACLKAIVLFKSETRGLKDPMQVENFQDQAQVMLCQHSRTNHPDQGQRFGRLL
LMLPLLKVVPTNRIEDIFFQPTIGNTPMEKVLCDMYKG*
>CLEC001111_HR78
MEKHAESSGIMNANSDKIHSLCLGVEICVVCGDRASGRHYGAISCEGCKGFFKRSIRKRLGYQCRGNQQCEVTKHHRNRC
QYCRLQKCLTMGMRTVQHERKPISVKKEFPNHSSPFYKSLTQPSNIGQSNYGVYNSLPYNLYSDGYIKQESSNIIGNYDYSNY
DESSDSDLIESIGPGQDSKSMINSAMEIATKLGMNGNFGNSDDEEMENLQFDGRLLDDNAFIFQIQSPSPMPAYTDVHYIY
ESASRLLFLSVHWVRNVPAFQILNMDTQVSLIKGCWSELFTLGLSQCAQVLALPTIILSIINHLQSSVAQQKISSSKVKAVTEHI
FSLQDFVGSMVALNVDDHEYAYLKLIALFSPDNPSLHIRRQLSELQEKAFQELREQIGDVNSNRFARLLLRLPPLRALNRHIM
EQIFFPGLGDQCDIDNIIPFILKMESIDFVNEQNN
>CLEC001266_E78C

MSRDSVRYGRVPKRSRERSGGEERVSTSDSSSGATPPDPETTSTPAYDLIVLEQQRIWLWQQFATHITPSVQRVVEFAKRV
PGFCELSQDDQLILIKVGFFELWLSHAARLTTDTTVTFSDGTFVTRQQMELMYDVSPTQSEFVTSMFEFTSSFNSLLLGDTEL
GLFSSIVLLSPDRPGVTDVKAVEHHQDRRSTNGTDSSSVLAKLPELRALGAKHALMLEWFRLNWDKLRLPPLFAEIFDIPKSE
EDL
