## Supplementary material info for "Transcriptomics supports local sensory regulation in the antenna of the kissing bug *Rhodnius prolixus*"

**DATABASES**

**Database S1 –** Protein sequences of all target genes in fasta format.

**Database S2 –** Edited Generic Feature Format (GFF) file of the *R. prolixus* genome used for read mapping and gene expression analysis.

**Database S3 -** FPKM values of target genes in the three libraries.

**Database S4** – Fasta sequences from different insects used in the CT/DH – CRF/DH and nuclear receptor phylogenetic analyses.

**FIGURES**

**Figure S1 - Molecular phylogenetic analyses of calcitonin diuretic (CT) and corticotropin-releasing factor-related (CRF) like diuretic hormone (DH) receptors of *R. prolixus* and other insects.**

The evolutionary history of *R. prolixus* CT/DH and CRF/DH receptors was inferred by using the Maximum Likelihood method in PhyML v3.0. The support values on the bipartitions correspond to SH-like P values, which were calculated by means of aLRT SH-like test. The CT/DH receptor 3 clade was highlighted in red. The CT/DH and CRF/DH *R. prolixus* receptors were displayed in blue. The LG substitution amino-acid model was used. Species abbreviations: Dmel, *Drosophila melanogaste*r; Aaeg, *Aedes aegytpi*; Agam, *Anopheles gambiae*; Clec, *Cimex lecturiaus*; Hhal, *Halomorpha halys;* Rpro, *Rhodnius prolixus*; Amel, *Apis mellifera*, *Acyrthosiphon pisum*; and Tcas, *Tribolium castaneum.* The glutamate receptor sequence from *the D. melanogaster* (FlyBase Acc. N° GC11144) was used as an out-group. The sequences used from other insects are in Supplementary Database S4).

**Figure S2 – Molecular phylogenetic analysis of nuclear receptor genes of *R. prolixus* and other insects.**

The evolutionary history of *R. prolixus* nuclear receptors was inferred by using the Maximum Likelihood method in PhyML v3.0. The support values on the bipartitions correspond to SH-like P values, which were calculated by means of aLRT SH-like test. The *R. prolixus* nuclear receptors were displayed in blue. LG substitution amino-acid model was used. *Drosophila melanogaster* sequences were obtained from Velarde et al. (2006); and subsequently used as query in BLASTp searches against *P. humanus* and *C. lecticularius* transcript databases from VectorBase. Species abbreviations: Dmel, *Drosophila melanogaste*r; Phum, *Pediculus humanus*; Clec, *Cimex lectularius*. The *RproEip75b* sequence used was from isoform B (from our antennal transcriptome) because the sequence of isoform A is considered incomplete. The sequences used from other insects are in Supplementary Database S4).

**Figure S3 - Alignment of *R. prolixus* takeout protein sequences**. Sequences were aligned with CLUSTAL X v2.0 (Thompson et al., 1997). Asterisks indicate identical amino-acids, double points show conserved exchanges and single points show homologous amino acids. The *Drosophila melanogaster* takeout protein sequence was obtained from Justice et al. (2003). The two conserved cysteine residues defining the Takeout family in many insects (Touhara et al., 1993) are marked with white boxes. The position of the conserved motifs 1 and 2 (So et al., 2000) is indicated with grey boxes. Predicted signal peptides are underlined. Species abbreviations: Rpro, *Rhodnius prolixus*; and Dmel, *Drosophila melanogaster*.

**Figure S4 – Structure and organization of *takeout* gene clusters**

Scaffold IDs are presented on the left. White arrows represent each *Takeout* gene and its position on the scaffold.

**Table S1. Details of neuropeptide and neurohormone precursor genes**. Columns are: Gene – the gene and protein name we are assigning; VectorBase code – the official gene number in the RproC3 genome assembly, prefix is RPRC; Scaffold – the RproC3.3 genome assembly supercontig ID; AAs – number of encoded amino acids in the protein; Comments – comments on the OGS gene model and repairs performed in the genome assembly based on Blast against *de novo* antennal assemblies. NTE: Amino terminal region; CTE: Carboxyl terminal region; VB: VectorBase; GB: GenBank.

| Gene | Gene symbol | VectoBase code | Scaffold | Isoforms | Aas. | Hit against the antennal *de novo* assemblies | Comments |
|---|---|---|---|---|---|---|---|
| **Adipokinetic hormone/corazonin-related peptide** | *ACP* | - | KQ035347 | - | 126 | Yes | New gene model created based on GB sequence Acc. N° KM975505 (Zandawala et al., 2015a) |
| **Adipokinetic hormone** | *AKH* | RPRC000416 | KQ034546 | A | 71 | No | VectorBase prediction identical to GB sequence Acc. N° KM283242 (Zandawala et al., 2015b) |
| | | | | B | 70 | Yes | New isoform identified based on antennal assemblies and included in the edited genome GFF |
| **AstA** | *AstA* | - | KQ034293 | - | 203 | Yes | New gene model created based on GB sequences Acc. N° GQ856315 and JN559385 (Ons et al., 2011) |
| **Allatostatin B** | *MIP* | - | KQ034158 | - | 254 | Yes | New gene model created based on Ons *et al.* 2011 |
| **Allatostatin CC** | *AstCC* | RPRC000300 | KQ034374 | - | 117 | Yes | No changes in VB prediction |
| **Allatostatin CCC** | *AstCCC* | - | KQ034609 | - | 100 | No | New gene model was created based on Ons *et. Al.* 2011 Previously, it was annotated as AstC |
| **Allatotropin** | *AT* | - | KQ034313 | - | 119 | Yes | New gene model created was created based on GB sequence Acc. N° GQ162783 (Ons et al., 2011) |
| **Bursicon alpha** | *Burs-alfa* | RPRC000797 | KQ034200 | - | 169 | Partial | No changes in VB prediction |
| **Bursicon beta** | *Burs-beta* | - | KQ034059 | - | 107 | No | Identified in this work using *T. castaneum* GB sequence Acc. N° NM_001114308.1 as query. |
| **Diuretic hormone 31** | *Dh31* | RPRC000977 | KQ034472 (5´UTR); KQ034594 (rest of the gene model) and KQ037272 (last exon) | A | 146 | Yes | Identical to GB sequences Acc. N° GQ856316 and AEA51300 (Ons et al., 2011). Last 42 amino acids are located in the KQ037272 supercontig |
| | | | | B | 109 | No | Identical to GB sequences Acc. N° GQ856317 and AEA51301 (Ons et al., 2011) |
| | | | | C | 206 | Yes | Identical to GB sequence Acc. N° HM030714.1 (Zandawala et al., 2011) |
| **Cardioacceleratory peptide (CAPA/CAP2b)** | *CAPA* | RPRC000639 | KQ034830 | A | 158 | No | VB prediction identical to GB sequence Acc. N° ABS17680 (Paluzzi et al., 2008). In VB classified as non-translating CDS |
| | | RPRC000563 | KQ034830 | B | 158 | No | VB prediction was identical to GB sequence Acc. N° ACH70295. In VB classified as non-translating CDS |

| Gene | Gene symbol | VectoBase code | Scaffold | Isoforms | Aas. | Hit against the antennal *de novo* assemblies | Comments |
|---|---|---|---|---|---|---|---|
| **Crustacean cardiactive peptide** | *CCAP* | RPRC000466 | KQ034330 | - | 129 | Yes | VB prediction was identical to GB sequence Acc. N° GQ888668 (Ons et al., 2011) |
| **CCHamide peptide** | *CCHa* | - | KQ034137 | - | 104 | Partial | New gene model created based Ons *et al*. 2011 and included in the edited genome GFF |
| **CNMamide peptide** | *CNMa* | RPRC010893 | KQ034609 | - | 150 | No | No changes in VB prediction |
| **Corazonin** | *CZ* | - | KQ034239 | - | | No | New gene model created based on Ons *et al*. 2011 and included in the edited genome GFF |
| **Diuretic hormone 44** | *Dh44* | RPRC000596 | KQ034102 | - | 151 | Yes | No changes in VB prediction, which is identical to GB sequence Acc. N° HM153808 (Te Brugge et al., 2011b), annotated as corticotropin releasing factor-like protein |
| **Eclosion hormone** | *EH* | RPRC014242 | KQ034677 | - | 241 | No | Partial sequence. Initial methionine is still missed |
| **Elevenin-1** | *Elevin-1* | RPRC003083 | KQ034317 | - | 66 | No | No changes in VB prediction |
| **Elevenin-2** | *Elevin-2* | RPRC003084 | KQ034317 | - | 87 | Yes | No changes in VB prediction |
| **Ecdysis triggering hormone** | *ETH* | RPRC014486 | KQ034462 | - | 146 | Yes | No changes in VB prediction |
| **FLP** | *FMRFamida* | RPRC014988 | KQ035274 | - | 273 | Partial | No changes in VB prediction |
| **Glycoprotein hormone alpha 2** | *GPA2* | RPRC007092 | KQ034094 | - | 122 | Partial | No changes in VB prediction |
| **Glycoprotein hormone beta 5** | *GPB5* | - | KQ034094 | - | 156 | No | New gene model was created based on *C. lectularius* sequence XP_014244389.1. Methionine is missed. |
| **Kinin** | | RPRC000022 | KQ034106 | - | 398 | Yes | VB prediction was identical to GB sequence Acc. N° BK007870 (Te Brugge et al., 2011a) |
| **IDLSRF-like peptide** | - | RPRC000351 | KQ034112 | - | 168 | Yes | No changes in VB prediction |
| **Insulin-like peptide** | *Ilp* | RPRC007020 | KQ034142 | - | 126 | Yes | VB prediction was identical to GB sequence Acc. N° AMS34841.1 (Defferrari et al., 2016) |
| **ITG-like** | - | - | KQ034255 | - | 214 | Yes | New gene model created based on Ons *et. al* 2011 and included in the edited genome GFF |
| **Ion transport peptide** | *ITP* | RPRC000519 | KQ034208 | A | 111 | No | Annotated as sulfakinin (GB sequence Acc. N° GQ2539210). Last 33 amino acids are not in the genome |
| | | | | B | 117 | Yes | VB prediction was identical to GB sequence Acc. N° GU207866 (Ons et al., 2011) |

| Gene | Gene symbol | VectoBase code | Scaffold | Isoforms | Aas. | Hit against the antennal *de novo* assemblies | Comments |
|---|---|---|---|---|---|---|---|
| **Long Neuropeptide F** | *LNPF* | RPRC008107 | KQ034255 | No | 105 | Yes | VB prediction extended in NTE region and initial methionine fixed according to GB sequence Acc. N° KT898124.1 (Sedra and Lange, 2016) |
| **Myosuppressin** | *Ms* | RPRC000203 | KQ034384 | No | 88 | Yes | VB prediction was identical to GB sequence Acc. N° GQ344501 (Ons et al., 2011) |
| **Natalisin** | *NTL* | RPRC003680 | KQ034106 | - | 196 | Partial | No changes in VB prediction |
| **Neuroparsin** | *NP* | RPRC002095 | Q034340 | | 113 | Yes | VB prediction was identical to GB sequence Acc. N° GU207864 (Ons et al., 2011) |
| **Neuropeptide like precursor 1** | *NPLP1* | RPRC011668 | KQ034238 | - | 454 | Yes | VB prediction was identical to GB sequence Acc. N° GU207865 (Ons et al., 2011) |
| **NVP-like** | *PH2* | RPRC003052 | ACPB03040762 | - | 299 | Yes | VB prediction was shorter in NTE and CTE regions. Impossible to fix the genome model due to problem in the genome assembly. |
| **Orcokinin** | *OK* | RPRC014678 | KQ034149 | A | 165 | Yes | One exon added in NTE region of VB prediction based on GB sequence Acc. N° FJ167860 |
| | | | | B | 392 | Yes | Only first 52 amino acids present in the genome. Sequence identical to GB sequence Acc. N° FJ761320 (Sterkel et al., 2012) |
| | | | | C | 422 | No | Same problem as mentioned for isoform B. Sequence identical to GB sequence Acc. N° KF179047 |
| **Pigment dispersing factor** | *PDF* | - | KQ034061 | - | 48 | Yes | New gene model created based on Ons *et al.* 2011 and included in the edited genome GFF |
| **Proctolin** | *Proc* | RPRC000390 | KQ034188 | - | 97 | No | VB prediction was identical to GB sequence Acc. N° JN543225 (Orchard et al., 2011) |
| **Pyrokinin** | *PK-PBAN* | - | KQ034521 | - | 122 | Yes | New gene model created based on GB sequence GU230851 and included in the edited genome GFF |
| **RYamide** | RYa | RPRC000461 | KQ035177 | - | 107 | Yes | Initial methionine of VB model was fixed according to antennal assemblies |
| **Short Neuropeptide F** | *sNPF* | - | KQ034092 | - | 92 | Yes | New gene model created based on GB sequence Acc. N° GQ452380 (Ons et al., 2011) and included in the edited genome GFF |
| **SIFamide** | *SIFa* | - | KQ035590 | - | 74 | Yes | New gene model created based on GB sequence Acc. N° GQ253922 (Ons et al., 2011) |
| **Sulphakinin** | *SK* | - | KQ034228 | - | 92 | Yes | New gene model created based on GB sequence Acc. N° GQ162784 (Ons et al., 2011) |
| **Tachykinin** | *TK* | RPRC000843 | ACPB03026326 | - | 215 | Yes | VB prediction was identical to GB sequence Acc. N° GQ162785 (Ons et al., 2011) |

**Table S2. Details of G protein coupled receptor genes**. Columns are: Gene – the gene and protein name we are assigning; VectorBase code – the official gene number in the RproC3 genome assembly, prefix is RPRC; Scaffold – the RproC3 genome assembly supercontig ID; AAs – number of encoded amino acids in the protein; Comments – comments on the OGS gene model and repairs performed in the genome assembly (available on VectorBase) based on Blast against *de novo* antennal assemblies. NTE: Amino terminal region; CTE: Carboxyl terminal region; VB: VectorBase.

| Gene | Ligand | VectorBase code | Scaffold | Aas. | Hit against the antennal *de novo* assemblies | Comments |
|---|---|---|---|---|---|---|
| **BIOGENIC AMINE G PROTEIN RECEPTORS** | | | | | | |
| **Muscarinic Acetylcholine receptor A** | Acetylcholine | - | KQ034078 | 555 | No | VB predictions (RPRC001750, 001751 and 007566) fused. The last prediction located in the opposite strand |
| **Muscarinic Acetylcholine receptor B** | | RPRC010907 | KQ034218 | 879 | Yes | The first exon extended. Initial methionine is still missed |
| **Muscarinic Acetylcholine receptor C** | | RPRC010656 | KQ034118 | 315 | Yes | The NTE extended. Initial methionine fixed |
| **Dopamine 1-like receptor 1** | Dopamine | RPRC014093 | KQ034515 | 395 | Partial | New exon added in NTE region. Initial methionine is still missed |
| **Dopamine 1-like receptor 2** | | RPRC013708 | KQ034114 | 426 | Partial | NTE region extended. Initial methionine is still missed |
| **Dopamine 2-like receptor** | | RPRC011175 | KQ034171 | 99 | No | Partial sequence. The last two exons eliminated. |
| **Dopamine Ecdysone receptor** | | RPRC014528 | KQ034056 | 354 | Partial | The initial methionine is missed. No changes in VB prediction |
| **Serotonin (5-HT) receptor 1a** | Serotonin | RPRC010931 | KQ034080 | 441 | Partial | The NTE extended. The initial methionine is still missed |
| **5-HT receptor 1b** | | RPRC008923 | KQ036077 and KQ034329 | 578 | Yes | The first 176 amino acids are located in KQ036077. VB prediction RPRC008923 edited |
| **5-HT receptor 2a** | | - | KQ034057 | 373 | Partial | VB predictions RPRC005858 and RPRC001892 fused. Initial methionine is still missed |
| **5-HT receptor 2b** | | RPRC000473 | KQ034057 | 703 | Yes | Identical to GB sequence Acc. N° AKQ13312 (Paluzzi et al., 2015a) |
| **5-HT receptor 7** | | - | KQ034099 | 423 | Partial | VB predictions RPRC007788 and RPRC001792 fused. Initial methionine is missed |
| **Octopamine (Oct) beta receptor 1** | Octopamine | - | KQ034268 | 426 | Partial | VB predictions RPRC001507 and RPRC005349 fused. Initial methionine is still missed |
| **Oct beta receptor 2** | | RPRC011545 | KQ034319 | 447 | Yes | Initial methionine fixed |
| **Oct beta receptor 3** | | - | KQ034653 | 375 | Partial | VB predictions RPRC014610 and RPRC001054 fused and edited. Initial methionine is still missed |
| **α2-adrenergic-like octopamine receptor** | | RPRC015456 | KQ034169 | 330 | Partial | Initial methionine is missed. No changes on VB prediction |
| **Oct receptor in mushroom bodies (Oamb) -like** | | RPRC001341 | KQ034231 | 417 | Partial | No changes in VB prediction. Partial sequence |
| **Oct/Tyramine** | Octopamine/Tyramine | RPRC008712 | KQ034100 | 455 | Yes | The two exons of the VB prediction fused |
| **Orphan receptor 1** | Unknown | RPRC004409 | KQ034373 | 235 | Partial | No changes in VB prediction |
| **Orphan receptor 2** | Unknown | RPRC002007-8 | KQ034058 | 201 | No | Two VB predictions fused |

| Gene | Ligand | VectorBase code | Scaffold | Aas. | Hit against the antennal *de novo* assemblies | Comments |
|---|---|---|---|---|---|---|
| **NEUROPEPTIDE RECEPTORS FAMILY A** | | | | | | |
| **ACP receptor isoform A** | AKH Corazonin related peptide | - | KQ034104, KQ035406 and KQ034241 | 295 | No | New gene model created based GB sequence Acc. N° KM975506 (Zandawala et al., 2015a) |
| **ACP receptor isoform B** | | - | | 451 | No | New gene model created based on GB sequence Acc. N° KM975507 (Zandawala et al., 2015a) |
| **ACP receptor isoform C** | | - | | 430 | No | New gene model created based on GB sequence Acc. N° KM975508 (Zandawala et al., 2015a) |
| **AKH receptor** | Adipokinetic hormone | - | KQ034132 | 353 | Yes | New gene model created based on GB sequence Acc. N° AIJ49751 (Zandawala et al., 2015b) |
| **Allatostatin A receptor** | Allatostatin A | RPRC004708 | KQ034532 | 404 | No | VB prediction updated according to GB sequence Acc. N° KM283241 (Zandawala and Orchard, 2015) |
| **Allatostatin C receptor** | Allatostatin C | RPRC013486 | KQ034333 | 419 | Yes | The initial methionine fixed according antennal transcriptome |
| **Allatotropin receptor** | Allatotropin | - | KQ034097 | 306 | Partial | New gene model created, which is partially in GB sequence Acc. N° KF740716 (Alzugaray et al., 2013) |
| **Bursicon receptor** | Bursicon | RPRC001663 | KQ034113 | 688 | Yes | No changes in VB prediction |
| **CAPA receptor isoform A** | CAPA peptide | RPRC000516 | KQ034065 | 385 | No | Identical to GB sequence Acc. N° ADG27752 (Paluzzi et al., 2010) |
| **CAPA receptor isoform B** | CAPA peptide | - | KQ034065 | 354 | No | New gene model created based on GB sequence Acc. N° ADG27753 (Paluzzi et al., 2010) |
| **CCH amide receptor 1** | CCHamide peptide | RPRC007766 | KQ034099 | 331 | No | No changes in VB prediction |
| **CCH amide receptor 2** | | RPRC000608 | KQ034099 | 373 | No | No changes in VB prediction |
| **CNM amide receptor** | CNMamide peptide | RPRC001428 | KQ034058 | 140 | Partial | Partial sequence. No changes in VB prediction |
| **Crustacean cardioactive peptide receptor 1** | CCAP | RPRC001248 | KQ034056 | 374 | Yes | Identical to GB sequence Acc. N° KC004225 (Lee et al., 2013) |
| **Crustacean cardioactive peptide receptor 2** | | - | KQ034561 and KQ034059 | 188 | No | VB predictions RPRC000969 and RPRC012063 fused. Partial sequence |
| **Corazonin receptor** | Corazonin | RPRC000523 | KQ034084 | 383 | No | No changes in VB prediction |
| **Corazonin receptor alfa isoform** | | - | KQ034084 | 441 | Yes | New gene model created based on α-isoform GB sequence Acc. N° AND99324 (Hamoudi et al., 2016) |
| **Corazonin receptor beta isoform** | | - | KQ034084 | 419 | No | New gene model created based on beta isoform GB sequence Acc. N° AND99325 (Hamoudi et al., 2016) |

| Gene | Ligand | VectorBase code | Scaffold | Aas. | Hit against the antennal *de novo* assemblies | Comments |
|---|---|---|---|---|---|---|
| **Ecdysis triggering hormone receptor** | Ecdysis triggering hormone | RPRC000848, RPRC008652 | KQ034066, KQ034378 and KQ034714 | 424 | Yes | Two VB predictions were fused and edited. |
| **FMRF receptor** | FaLP | RPRC001551 | KQ034140 | 410 | Yes | No changes in VB prediction |
| **FaLPamide/Proctolin receptor** | FaLP/Proctolin | RPRC015267 | KQ034074 | 345 | Partial | No changes in VB prediction. Partial sequence, initial methionine is still missed |
| **GPA2/GPB2 receptor** | GPA2/GPB5 | RPRC007243 | KQ034109 | 591 | Partial | CTE was extended until STOP codon. Partial sequence, initial methionine is still missed |
| **Ion Transport 1 receptor** | - | RPRC004793 | KQ034083 | 611 | Yes | No changes in VB prediction |
| **Kinin receptor 1** | Kinin<br>- | RPRC000494 | KQ034056 | 414 | Yes | Two first exons and last exon eliminated from VB prediction according to our antennal transcriptome |
| **Kinin receptor 2** | | - | KQ034861 and KQ034100 | 366 | Yes | Two predictions RPRC008570 and RPRC008649 fused and edited. Described as orphan by Ons et al. (2015) as RPRC008570 |
| **Long neuropeptide F receptor 1** | Neuropeptide F | - | KQ034119 | 390 | Partial | New gene model created based on (Sedra et al., 2018) |
| **Long neuropeptide F receptor 2** | | RPRC008894 | KQ034459 | 180 | Partial | No changes in VB prediction. Partial sequence, initial methionine is still missed |
| **Myoinhibitory peptide receptor isoform A** | Allatostatin B | RPRC000605 | KQ034129 | 420 | Partial | VB prediction was identical to GB sequence Acc. N° KF958188 (Paluzzi et al., 2015b) |
| **Myoinhibitory peptide receptor isoform B** | | - | KQ034129 | 324 | No | No VB prediction |
| **Myosuppressin receptor** | Myosuppressin | - | KQ034057 | 368 | No | New gene model created based on GB sequence Acc. N° AGT02812 (Lee et al., 2015) |
| **Natalisin receptor** | Natalisin | RPRC001687 | KQ034139 | 351 | Yes | VB prediction extended in NTE region according to antennal transcriptome |
| **Pyrokinin 1 receptor** | PBAN | RPRC008528 | KQ034938 | 208 | No | No changes in VB prediction. Partial sequence. |
| **Pyrokinin 2 receptor isoform A** | | - | KQ034161 | 345 | Partial | New gene model created based on GB sequence Acc. N° AFO73269 (Paluzzi and O'Donnell, 2012). Sequence partially represented at RPRC005110 |
| **Pyrokinin 2 receptor isoform B** | | - | KQ034161 | 444 | Partial | New gene model created based on GB sequence Acc. N° AFO73270 (Paluzzi and O'Donnell, 2012). Sequence partially represented at RPRC005110 |
| **Pyrokinin 2 receptor isoform C** | | - | KQ034161 | 414 | Partial | New gene model created based on GB sequence Acc. N° AFO73271 (Paluzzi and O'Donnell, 2012). Partially represented RPRC005110 |

| Gene | Ligand | VectorBase code | Scaffold | Aas. | Hit against the antennal *de novo* assemblies | Comments |
|---|---|---|---|---|---|---|
| **Pyroglutamylate RFamide peptide receptor** | Rfamide peptides | - | KQ034100 | 395 | Yes | New gene model created. First 100 amino acids in RPRC014460 |
| **RYamide receptor** | Ryamide | - | KQ034249 and KQ034213 | 360 | Partial | New gene model created based on Ons [13]. First 133 amino acids in KQ034249. Problem between 3rd and 4th exons |
| **Short neuropeptide F receptor** | Short neuropeptide F | - | KQ034095 and KQ035872 | 448 | Yes | Three VB predictions (RPRC002266, 002268 and 002269) fused |
| **Sulfakinin receptor 1** | Sulfakinin | - | KQ035199, KQ035392 and KQ034565 | 320 | No | RPRC012816 (KQ034095) and RPRC003273 (KQ035392) represented different parts of gene model |
| **Sulfakinin receptor 2** | | RPRC012816 | KQ034565 | 138 | No | No changes in VB prediction. Partial sequence. |
| **SIFamide receptor** | SIFamide | RPRC000835 | ACPB03024746 | 451 | Yes | The first part of VB prediction eliminated. Only the last 264 amino acids were present in the genome |
| **Tachykinin receptor 86C-like** | Tachykinin | RPRC008022 | kQ035269 | 309 | Yes | No changes in VB prediction. Identified as ITP receptor by Ons (2017) |
| **Tachykinin receptor 99D-like** | | - | KQ034874 and KQ034432 | 378 | Yes | VB predictions RPRC003160 and RPRC000651 were fused |
| **Orphan receptor 3** | - | RPRC014721 | KQ034261 | 962 | Partial | No changes in VB prediction. Initial methionine is missed |
| **Orphan receptor 4** | - | RPRC004128 | KQ035493 | 381 | No | No changes in VB prediction. Initial methionine is missed |
| **Orphan receptor 5** | - | RPRC008364 | KQ034143 | 117 | No | No changes in VB prediction. Partial sequence |
| **NEUROPEPTIDE RECEPTORS FAMILY B** | | | | | | |
| **Calcitonin-like diuretic hormone receptor 1 isoform B** | Dh31 | RPRC009814 | KQ035556 and KQ034793 | 411 | Yes | First 77 amino acids located in KQ035556. Identical to GB sequence Acc. N° AHB86317 (Zandawala et al., 2013) |
| **Calcitonin-like diuretic hormone receptor 1 isoform C** | | - | KQ035556 and KQ034793 | 409 | Partial | First 77 amino acids located in KQ035556. New gene model included. Identical to GB sequence Acc. N° AHB86318 (Zandawala et al., 2013) |
| **Calcitonin-like diuretic hormone receptor 2** | | RPRC004753 | KQ034099 | 410 | No | VB prediction edited according to GB sequence Acc. N° AHB86571 (Zandawala et al., 2013) |
| **Calcitonin-like diuretic hormone receptor 3** | - | RPRC004735 | KQ034099 | 420 | Yes | Initial methionine fixed and an internal region was added to VB prediction |
| **Corticotropin-releasing factor-related like diuretic hormone receptor 1** | Dh44 | - | KQ034141 and KQ035235 | 465 | Yes | New gene model created. |

| Gene | Ligand | VectorBase code | Scaffold | Aas. | Hit against the antennal *de novo* assemblies | Comments |
|---|---|---|---|---|---|---|
| **Corticotropin-releasing factor-related like diuretic hormone receptor 2 isoform A** | Dh44 | - | KQ034325 | 385 | No | VB prediction modified according to GB sequence Acc. N° KU942308 (Lee et al., 2016). NTE is missed. CTE region was extended. |
| **Corticotropin-releasing factor-related like diuretic hormone receptor 2 isoform B** | Dh44 | RPRC000578 | KQ034325 | 485 | Yes | VB prediction modified according to GB sequence Acc. N° KJ407397 (Lee et al., 2016) |
| **PDF receptor** | PDF | RPRC009680 | KQ034059 | 318 | Partial | The NTE region extended. Initial methionine is still missed |
| **Parathyroid hormone like receptor** | - | - | KQ034058 | 498 | Yes | Predictions RPRC011083 and RPRC011086 fused and edited |
| **OPSINS** | | | | | | |
| **Long wave sensitive opsin 1 (LWS)** | - | RPRC010623 | KQ034901 | 377 | No | No changes in VB prediction |
| **UV opsin** | - | RPRC002621 | KQ034248 | 387 | Yes | No changes in VB prediction. Initial methionine is missed |
| **COpsin / Pteropsin** | - | RPRC017360 | KQ034389 | 301 | No | No changes in VB prediction |
| **Rh7** | - | RPRC015283 | KQ034074 | 361 | Partial | No changes in VB prediction. Initial methionine is missed |
| **TYROSINE KINASE AND GUANYLYL CYCLASE RECEPTORS** | | | | | | |
| **Eclosion hormone receptor** | Eclosion hormone | RPRC013306 | KQ034473 | 1160 | No | No changes in VB prediction. Initial methionine is missed |
| **NPLP receptor** | NPLP | RPRC013388 | KQ034473 | 1159 | Yes | NTE and CTE must be extended. Initial methionine is still missed |
| **Potential neuroparsin receptor*** | Ovary ecdysteroidogenic hormone | RPRC006045 | KQ034063 | 1290 | Yes | The first exon was extended. Initial methionine was fixed. In VB, initial methionine is missed due to an assembly problem |
| **Insulin receptor*** | Insulin | RPRC006251 | KQ034536 | 1198 | Yes | Initial methionine must be fixed and some internal changes are necessary |

* The sequences included in the Supplementary DataSet 1 for these genes are those obtained after the comparison to our antennal transcriptome assemblies and the adequate correction. The gene models included for these genes in our GFF file were those available in VectorBase.

CT/DH receptor 1 (variant A) and CT/DH receptor 2 (variant A) reported by Zandawala et al. (2013) are a partial sequences (143 and 122 amino acids length, respectively), and they do not have VectorBase prediction. A partial CRF/DH receptor 1 variant A sequence was reported by Lee et al. (2016), however, it is not available in GenBank.

**Table S3. Details of enzymes involved in the biogenic amines synthesis**. Columns are: Gene – the gene and protein name we are assigning; VectorBase code – the official gene number in the RproC3 genome assembly, prefix is RPRC; Scaffold – the RproC3 genome assembly supercontig ID; Aas – number of encoded amino acids in the protein; Comments – comments on the OGS gene model and repairs to be performed on the genome assembly (available on VectorBase) based on Blast searches against *de novo* antennal transcriptome assemblies. NTE: Amino terminal region.

| Gene | VectorBase code | Scaffold | Aas. | Hit against the antennal *de novo* assemblies | Comments |
|---|---|---|---|---|---|
| **Tyrosine 3-monooxigenase (ple)** | RPRC007034 | KQ034272 | 569 | Yes | Some internal problems were detected* |
| **DOPA decarboxylase (Ddc)** | RPRC005884 | KQ034063 | 629 | Yes | NTE region must be extended until initial methionine* |
| **Tyrosine decarboxylase-2 (Tdc2)** | RPRC011470 | KQ034319 | 476 | No | Fine as is |
| **Tryptophan hydroxylase (Trh)** | RPRC012297 | KQ034056 | 490 | No | Stop codon is missed |

(*) The sequences included in the Supplementary DataSet 1 for these genes are those obtained after the comparison to our antennal transcriptome assemblies and the appropriate correction.

**Table S4. Details of neuropeptide processing enzymes**. Columns are: Gene – the gene and protein name we are assigning; VectorBase code – the official gene number in the RproC3 genome assembly, prefix is RPRC; Scaffold – the RproC3 genome assembly supercontig ID; Aas – number of encoded amino acids in the protein; Comments – comments on the OGS gene model and repairs to be performed on the genome assembly (available on VectorBase) based on Blast searches against *de novo* antennal transcriptome assemblies.

| Gene | VectorBase code | Scaffold | Aas. | Hit against the antennal *de novo* assemblies | Isoforms | Comments |
|---|---|---|---|---|---|---|
| **Signal Peptidase (SP)** | RPRC009668 | KQ034208 | 375 | Yes | No | CTE is missed |
| **Amontillado (Prohormone convertase 2- PC2)** | RPRC009349 | KQ034234 | 640 | Yes | No | Initial methionine is missed |
| **Silver (Carboxypeptidase D)** | - | KQ034072 | 1129 | Yes | No | New gene model was created. RPRC011379, RPRC011383 and RPRC0011427 were fused. Partial sequence, methionine and stop codon are still missed* |
| **Prolyl endopeptidase** | RPRC006929 | KQ034086 | 690 | Yes | No | Methionine and stop codon missed. Identified using *D. melanogaster* sequence CG5355 |
| **Carboxypeptidase M (CPM)** | RPRC015124 | KQ034103 | 454 | No | A | Methionine is missed. Identified using *D. melanogaster* sequence CG4678 |
| | | | 471 | Yes | B | New isoform identified in the antennal transcriptome |
| | | | 525 | Yes | C | New isoform identified in the antennal transcriptome |
| **Peptidylglycine alfa-hydroxylating mono-oxygenase (PHM)** | - | KQ034112 | 329 | Yes | No | VectorBase predictions RPRC001216 and RPRC001217 must be fused and edited |
| **Furin (Fur) like protease-1** | RPRC006957 | KQ034094 | 810 | Yes | No | A total of 126 amino acids are missed in NTE region. Some internal problems were detected |
| **Fur-like protease 2A** | RPRC002472 | KQ034090 | 1162 | Yes | No | NTE and CTE are missed |
| **Fur-like protease 2B** | RPRC013490 | KQ034542 | 688 | Yes | No | NTE and CTE are missed |
| **Peptidyl-alpha-hydroxyglycine alpha-amidating lyase 1 (PAL1)** | - | KQ034270 | 438 | Yes | No | An internal region is missed due to problems in the genome assembly. CTE region located in the opposite strand* |
| **Peptidyl-alpha-hydroxyglycine alpha-amidating lyase 2 (PAL2)** | - | KQ034195 | 370 | Yes | No | The first two exons located in the opposite strand* |

* These models were fixed and included in the modified GFF file that was used for mapping of our RNASeq reads.

The sequences included in the Supplementary DataSet 1 for all genes are those obtained after the comparison to our antennal transcriptome assemblies and the adequate correction.

**Table S5. Details of nuclear receptor genes**. Columns are: Gene – the gene and protein name we are assigning; VectorBase code – the official gene number in the RproC3 genome assembly, prefix is RPRC; Scaffold – the RproC3 genome assembly supercontig ID; Aas – number of encoded amino acids in the protein; Comments – comments on the OGS gene model and repairs to be performed on the genome assembly (available on VectorBase) based on Blast searches against *de novo* antennal assemblies. NTE: Amino terminal region; CTE: Carboxyl terminal region; VB: VectorBase.

| Gene | VectorBase code | Scaffold | Aas. | Hit against the antennal *de novo* assemblies | Comments |
|---|---|---|---|---|---|
| ***Knirps-like1*** | RPRC003216 | KQ035852 | 325 | No | Fine as is |
| ***Knirps-like2*** | - | ACPB3007969 | 302 | Si | New gene model was created* |
| **Ecdysone-induced protein 75B isoform A** | RPRC000853 | KQ034727 | 717 | No | Methionine is missed. Exon 4th was incorrect and 5th exon in the opposite strand. |
| **Ecdysone-induced protein 75B isoform B** | RPRC000853 | KQ034727 | 653 | Si | Only exons 2nd and 3rd were predicted in VB. Exon 4th was incorrect and 5th exon in the opposite strand* |
| **Ecdysone-induced protein 78C** | RPRC009045 | KQ034642 | 344 | No | Initial methionine is missed |
| **Hormone receptor-like in 3** | RPRC003681 and RPRC0000824 | KQ034284 and KQ036430 | 579 | Si | Two VB predictions must be fused and edited. NTE region must be extended until initial methionine* |
| **Ecdysone (Ec) receptor** | RPRC014174 | KQ034515 | 475 | Si | NTE region must be extended until initial methionine* |
| **Hormone receptor-like in 96** | RPRC001794 | KQ034099 | 377 | No | Fine as is |
| **Hepatocyte nuclear factor 4** | RPRC008212 | KQ034483 | 396 | Si | CTE terminal region is incomplete. Initial methionine is missed |
| ***Ultraspiracle*** | RPRC013330 | KQ034117 | 430 | Si | NTE and CTE regions are incomplete* |
| **Hormone receptor-like in 78** | RPRC006737 | KQ034201 | 495 | Si | NTE and CTE regions are incomplete* |
| ***Tailless*** | RPRC007025 | KQ034142 | 370 | No | Initial methionine is missed |
| **Hormone receptor-like in 51 (*unfulfilled*)** | RPRC002557 | KQ034474 | 478 | No | Initial methionine is missed |
| ***Dissatisfaction*** | RPRC010625 | KQ034604 | 195 | No | Initial methionine is missed |
| ***PNR- like (*NR2E6*)*** | RPRC009755 | KQ034409 | 394 | Si | Multiple changes in VB prediction are necessary* |
| ***Seven up*** | RPRC000767 | KQ034946 | 227 | No | Fine as is |
| **Estrogen-related receptor** | - | ACPB03009538 | 426 | Si | New gene model was created* |
| **Hormone receptor-like in 38** | RPRC001680 | KQ034154 | 287 | Si | NTE is missed* |
| **Ftz transcription factor 1** | RPRC001915 | KQ034834 and KQ034274 | 618 | Si | VB predictions RPRC014120 and RPRC001915must be fused and edited* |
| **Hormone receptor-like in 39** | RPRC002968 | KQ034115 | 695 | Si | NTE region must be extended until initial methionine* |
| **Hormone receptor-like in 4** | RPRC012796 | KQ034081 | 542 | Si | NTE region must be extended until initial methionine and CTE is missed |

*Knrips-like2* and, *PNR-like*, Hormone receptor-like in 39**,** Estrogen-related receptor gene models were included in the modified GFF file that was used for mapping of our RNASeq reads.(*) The sequences included in the Supplementary DataSet 1 for these genes are those obtained after the comparison to our antennal transcriptome assemblies and the appropriate correction.

**Table S6. Details of Takeout (*to*) genes**. Columns are: Gene – the gene and protein name we are assigning; VectorBase code – the official gene number in the RproC3 genome assembly, prefix is RPRC; Scaffold – the RproC3 genome assembly supercontig ID; Aas – number of encoded amino acids in the protein; Comments – comments on the OGS gene model and repairs to be performed on the genome assembly (available on VectorBase) based on Blast searches against *de novo* antennal transcriptome assemblies. NTE: Amino terminal region.

| Gene | VectorBase code | Scaffold | Aas. | Hit against the antennal *de novo* assemblies | Comments |
|---|---|---|---|---|---|
| ***to1*** | RPRC010098 | KQ034137 | 244 | Yes | Fine as is |
| ***to2*** | RPRC010096 | KQ034137 | 242 | Yes | Annotated as *to3* |
| ***to3*** | RPRC008440 | KQ034102 | 248 | Yes | The initial methionine must be fixed |
| ***to4*** | RPRC008432 | KQ034102 | 191 | Yes | Some internal problems detected |
| ***to5*** | RPRC008451 | KQ034102 | 222 | Yes | The initial methionine must be fixed |
| ***to6*** | RPRC002313 | KQ034398 | 147 | Yes | N-terminal region must be extended until initial methionine. Annotated as *to2* |
| ***to7*** | RPRC008276 | KQ034251 | 250 | Yes | Fine as is |
| ***to8*** | RPRC009613 | KQ034205 | 250 | Yes | Fine as is |
| ***to9*** | RPRC010085 | KQ034137 | 247 | Yes | Internal problems were fixed* |
| ***to10*** | RPRC011983 and RPRC011984 | KQ034059 | 144 | Yes | Two VectorBase predictions must be fused |
| ***to11*** | RPRC005773 | KQ034137 | 248 | Yes | Fine as is |
| ***to12*** | RPRC010201 | KQ034137 | 226 | Yes | Fine as is |
| ***to13*** | RPRC010202 | KQ034137 | 227 | Yes | Fine as is |
| ***to14*** | RPRC005774 | KQ034137 | 259 | Yes | Fine as is |
| ***to15*** | RPRC005775 | KQ034137 | 255 | Yes | Fine as is |

* This model was fixed and included in the modified GFF file that was used for mapping of our RNASeq reads.

The sequences included in the Supplementary Database 1 for *to3*, *to4*, *To5*, *to6*, *to9* and *to10* genes are those obtained after the comparison to our antennal transcriptome assemblies and the adequate correction.

**Table S7. Statistical results of differentially expressed sensory genes among studied stages.** Comparison of normalized counts per million (CPM) among stages was conducted using edgeR package. LogFC: Log fold change; FDR adjusted p-value: False Discovery Rate.

| Annotation | VectorBase Code | Log FC | Log CPM | P value | FDR adjusted P value |
|---|---|---|---|---|---|
| Female vs Larvae | | | | | |
| Allatostatin-A | - | -4.1 | 2.8 | 0.0001 | 0.0196 |
| Myoinhibitory peptide | - | -4.1 | 3.0 | 0.0001 | 0.0176 |
| Adipokinetic hormone receptor | - | -3.6 | 2.3 | 0.0006 | 0.0617 |
| Calcitonin-like diuretic hormone receptor 3 | RPRC004735 | 3.1 | 6.0 | 0.0013 | 0.0978 |
| Kinin receptor 2 | - | 3.7 | 2.3 | 0.0006 | 0.0581 |
| *to11* | RPRC005773 | -4.6 | 5.9 | 2.37E-05 | 0.0054 |
| *to3* | RPRC008440 | 5.2 | 10.9 | 3.03E-06 | 0.0011 |
| Male vs Larvae | | | | | |
| Allatostatin-A | - | -5.3 | 2.9 | 0.000005 | 0.001 |
| Allatostatin-CC | RPRC000300 | -4.3 | 8.0 | 0.000062 | 0.008 |
| Myoinhibitory peptide | - | -4.4 | 3.2 | 0.000054 | 0.007 |
| Calcitonin-like diuretic hormone receptor 3 | RPRC004735 | 3.5 | 6.4 | 0.000593 | 0.041 |
| Kinin receptor 2 | - | 4.1 | 2.9 | 0.000144 | 0.014 |
| Hormone receptor-like in 3 | RPRC000824-RPRC003681 | 4.3 | 1.4 | 0.000185 | 0.017 |
| Octopamine beta receptor 3 | - | 3.4 | 1.7 | 0.001 | 0.071 |
| *to11* | RPRC005773 | -4.8 | 6.1 | 0.00001 | 0.002 |
| *to3* | RPRC008440 | 4.3 | 10.2 | 0.00006 | 0.008 |
| *to2* | RPRC010096 | -4.1 | 13.2 | 0.00011 | 0.012 |

Alzugaray, M.E., Adami, M.L., Diambra, L.A., Hernandez-Martinez, S., Damborenea, C., Noriega, F.G., Ronderos, J.R., 2013. Allatotropin: an ancestral myotropic neuropeptide involved in feeding. PLoS One 8, e77520.

Defferrari, M.S., Orchard, I., Lange, A.B., 2016. Identification of the first insulin-like peptide in the disease vector *Rhodnius prolixus*: involvement in metabolic homeostasis of lipids and carbohydrates. Insect Biochem Mol Biol 70, 148-159.

Hamoudi, Z., Lange, A.B., Orchard, I., 2016. Identification and characterization of the corazonin receptor and possible physiological roles of the corazonin-signaling pathway in *Rhodnius prolixus*. Front Neurosci 10, 357.

Justice, R., Dimitratos, S., Walter, M., Woods, D., Biessmann, H., 2003. Sexual dimorphic expression of putative antennal carrier protein genes in the malaria vector *Anopheles gambiae*. Insect molecular biology 12, 581-594.

Lee, D., Broeck, J.V., Lange, A.B., 2013. Identification and expression of the CCAP receptor in the Chagas' disease vector, *Rhodnius prolixus*, and its involvement in cardiac control. PLoS One 8, e68897.

Lee, D., James, T., Lange, A., 2015. Identification, characterization and expression of a receptor for the unusual myosuppressin in the blood feeding bug, *Rhodnius prolixus*. Insect Mol Biol 24, 129-137.

Lee, H.-R., Zandawala, M., Lange, A.B., Orchard, I., 2016. Isolation and characterization of the corticotropin-releasing factor-related diuretic hormone receptor in *Rhodnius prolixus*. Cellular Signalling 28, 1152-1162.

Ons, S., 2017. Neuropeptides in the regulation of *Rhodnius prolixus* physiology. J Insect Physiol Paris 97, 77-92.

Ons, S., Lavore, A., Sterkel, M., Wulff, J.P., Sierra, I., Barnetche, J.M., Rodriguez, M.H., Rivera-Pomar, R., 2015. Identification of G-protein coupled receptors for opsines and neurohormones in *Rhodnius prolixus*. Genomic and transcriptomic analysis. Insect Biochem Mol Biol.

Ons, S., Sterkel, M., Diambra, L., Urlaub, H., Rivera-Pomar, R., 2011. Neuropeptide precursor gene discovery in the Chagas disease vector *Rhodnius prolixus*. Insect Mol Biol 20, 29-44.

Orchard, I., Lee, D.H., Da Silva, R., Lange, A.B., 2011. The proctolin gene and biological effects of proctolin in the blood-feeding bug, *Rhodnius prolixus*. Front Endocrinol 2, 59.

Paluzzi, J.-P., O'Donnell, M.J., 2012. Identification, spatial expression analysis and functional characterization of a pyrokinin-1 receptor in the Chagas' disease vector, *Rhodnius prolixus*. Mol Cell Endocrinol 363, 36-45.

Paluzzi, J.-P., Russell, W.K., Nachman, R.J., Orchard, I., 2008. Isolation, cloning, and expression mapping of a gene encoding an antidiuretic hormone and other CAPA-related peptides in the disease vector, *Rhodnius prolixus*. Endocrinology 149, 4638-4646.

Paluzzi, J.-P.V., Bhatt, G., Wang, C.-H.J., Zandawala, M., Lange, A.B., Orchard, I., 2015a. Identification, functional characterization, and pharmacological profile of a serotonin type-2b receptor in the medically important insect, *Rhodnius prolixus*. Front Neurosci 9, 175.

Paluzzi, J.-P.V., Haddad, A.S., Sedra, L., Orchard, I., Lange, A.B., 2015b. Functional characterization and expression analysis of the myoinhibiting peptide receptor in the Chagas disease vector, *Rhodnius prolixus*. Mol Cell Endocrinol 399, 143-153.

Paluzzi, J.P., Park, Y., Nachman, R.J., Orchard, I., 2010. Isolation, expression analysis, and functional characterization of the first antidiuretic hormone receptor in insects. Proc Natl Acad Sci U S A 107, 10290-10295.

Sedra, L., Lange, A.B., 2016. Cloning and expression of long neuropeptide F and the role of FMRFamide-like peptides in regulating egg production in the Chagas vector, *Rhodnius prolixus*. Peptides 82, 1-11.

Sedra, L., Paluzzi, J.-P., Lange, A.B., 2018. Characterization and expression of a long neuropeptide F (NPF) receptor in the Chagas disease vector *Rhodnius prolixus*. PLoS One 13, e0202425.

So, W.V., Sarov-Blat, L., Kotarski, C.K., McDonald, M.J., Allada, R., Rosbash, M., 2000. Takeout, a novel *Drosophila* gene under circadian clock transcriptional regulation. Mol Cell Biol 20, 6935-6944.

Sterkel, M., Oliveira, P.L., Urlaub, H., Hernandez-Martinez, S., Rivera-Pomar, R., Ons, S., 2012. OKB, a novel family of brain-gut neuropeptides from insects. Insect Biochem Mol Biol 42, 466-473.

Te Brugge, V., Paluzzi, J.-P., Neupert, S., Nachman, R.J., Orchard, I., 2011a. Identification of kinin-related peptides in the disease vector, *Rhodnius prolixus*. Peptides 32, 469-474.

Te Brugge, V., Paluzzi, J.-P., Schooley, D.A., Orchard, I., 2011b. Identification of the elusive peptidergic diuretic hormone in the blood-feeding bug *Rhodnius prolixu*s: a CRF-related peptide. J Exp Biol 214, 371-381.

Thompson, J.D., Gibson, T.J., Plewniak, F., Jeanmougin, F., Higgins, D.G., 1997. The CLUSTAL_X windows interface: flexible strategies for multiple sequence alignment aided by quality analysis tools. Nucleic acids research 25, 4876-4882.

Touhara, K., Lerro, K.A., Bonning, B.C., Hammock, B.D., Prestwich, G.D., 1993. Ligand binding by a recombinant insect juvenile hormone binding protein. Biochemistry 32, 2068-2075.

Velarde, R.A., Robinson, G.E., Fahrbach, S.E., 2006. Nuclear receptors of the honey bee: annotation and expression in the adult brain. Insect molecular biology 15, 583-595.

Zandawala, M., Haddad, A.S., Hamoudi, Z., Orchard, I., 2015a. Identification and characterization of the adipokinetic hormone/corazonin-related peptide signaling system in *Rhodnius prolixus*. The FEBS Journal 282, 3603-3617.

Zandawala, M., Hamoudi, Z., Lange, A.B., Orchard, I., 2015b. Adipokinetic hormone signalling system in the C hagas disease vector, *Rhodnius prolixus*. Insect Mol Biol 24, 264-276.

Zandawala, M., Li, S., Hauser, F., Grimmelikhuijzen, C.J., Orchard, I., 2013. Isolation and functional characterization of calcitonin-like diuretic hormone receptors in *Rhodnius prolixus*. PLoS One 8, e82466.

Zandawala, M., Orchard, I., 2015. Identification and functional characterization of FGLamide-related allatostatin receptor in *Rhodnius prolixus*. Insect Biochem Mol Biol 57, 1-10.

Zandawala, M., Paluzzi, J.-P., Orchard, I., 2011. Isolation and characterization of the cDNA encoding DH31 in the kissing bug, *Rhodnius prolixus*. Mol Cell Endocrinol 331, 79-88.
